# Cyclic dichalcogenides extend the reach of bioreductive prodrugs to harness the thioredoxin system: applications to *seco*-duocarmycins

**DOI:** 10.1101/2022.11.11.516112

**Authors:** Jan G. Felber, Annabel Kitowski, Lukas Zeisel, Martin S. Maier, Constanze Heise, Julia Thorn-Seshold, Oliver Thorn-Seshold

## Abstract

Small molecule prodrug approaches that can activate cancer therapeutics selectively in tumors are urgently needed. Here, we developed the first antitumor prodrugs designed for activation by the thioredoxin (Trx) oxidoreductase system. This critical cellular disulfide redox axis is tightly linked to dysregulated redox/metabolic states in cancer, yet it cannot be addressed by current bioreductive prodrugs, which mainly cluster around oxidised nitrogen species. We instead harnessed Trx/TrxR-specific artificial dichalcogenides to gate the bioactivity of a series of 10 “off-to-on” reduction-activated duocarmycin prodrugs. The prodrugs were tested for cell-free and cellular activity dependent on reducing enzyme systems in 177 cell lines, to establish broad trends for redox-based cellular bioactivity of the dichalcogenides. They were well tolerated *in vivo* in mice, indicating low systemic release of their duocarmycin cargo, and *in vivo* anti-tumor efficacy trials in mouse models of breast and pancreatic cancer gave promising initial results indicating effective tumoral drug release, presumably by *in situ* bioreductive activation. This work therefore presents a chemically novel class of bioreductive prodrugs against a previously unaddressed reductase type, validates its ability to access *in vivo* compatible small-molecule prodrugs even of potently cumulative toxins, and so introduces carefully tuned dichalcogenides as a platform strategy for specific bioreduction-based release.

## 1. INTRODUCTION

Classic cancer chemotherapy, treating tumors with cytotoxic drugs against ubiquitous critical biological targets (DNA integrity, cell division, etc.), incurs severe systemic side effects from unspecific drug distribution to healthy tissues whose function also depends on these targets. Prodrug concepts can deliver an additional layer of control over drug activity beyond simple biodistribution, and tumor-preferential mechanisms for drug unmasking – e.g. small molecule^1^ or antibody-directed^2^ approaches – are intensively pursued.

Bioreductive prodrugs are enzymatically unmasked *in situ* by reduction^3^. Reductive processes are especially prominent in hypoxic environments, such as tumors, since re-oxidation of metastable intermediates is hindered. Thus, bioreductive prodrugs are sometimes termed “hypoxia-activated”. They are in active development, and several reached phase III clinical trials. However, only a small biological target space and a correspondingly restricted small chemical space have been explored for bioreductive prodrugs, which is a missed opportunity for innovative therapeutics (**Fig 1a**). The first class developed were natural product quinones, activated by ubiquitous quinone reductases, such as the mitomycins and their synthetic analogues (e.g. apaziquone^4^). Later, oxidised nitrogen species that can be reduced by a broader range of enzyme classes came to dominate designs: including (i) aliphatic *N*-oxides that are reduced to basic amines, which can trigger DNA binding (AQ4N/banoxantrone^5^); (ii) aromatic *N-*oxides that are reduced to bioactive nitrogenous bases (tirapazamine^6^); (iii) nitroaryls, that after reduction can eliminate a drug (TH-302/Evofosfamide^7^), or nucleophilically assist reaction mechanisms (PR-104^8^), or dock to targets (nitracrine^9^) (**Fig 1a**). However, no cancer prodrug using these oxidised nitrogen chemotypes and their targets has been approved;^10^ and novel, modular strategies to bioreductively activate drugs specifically inside tumors are required.

**Figure 1:**
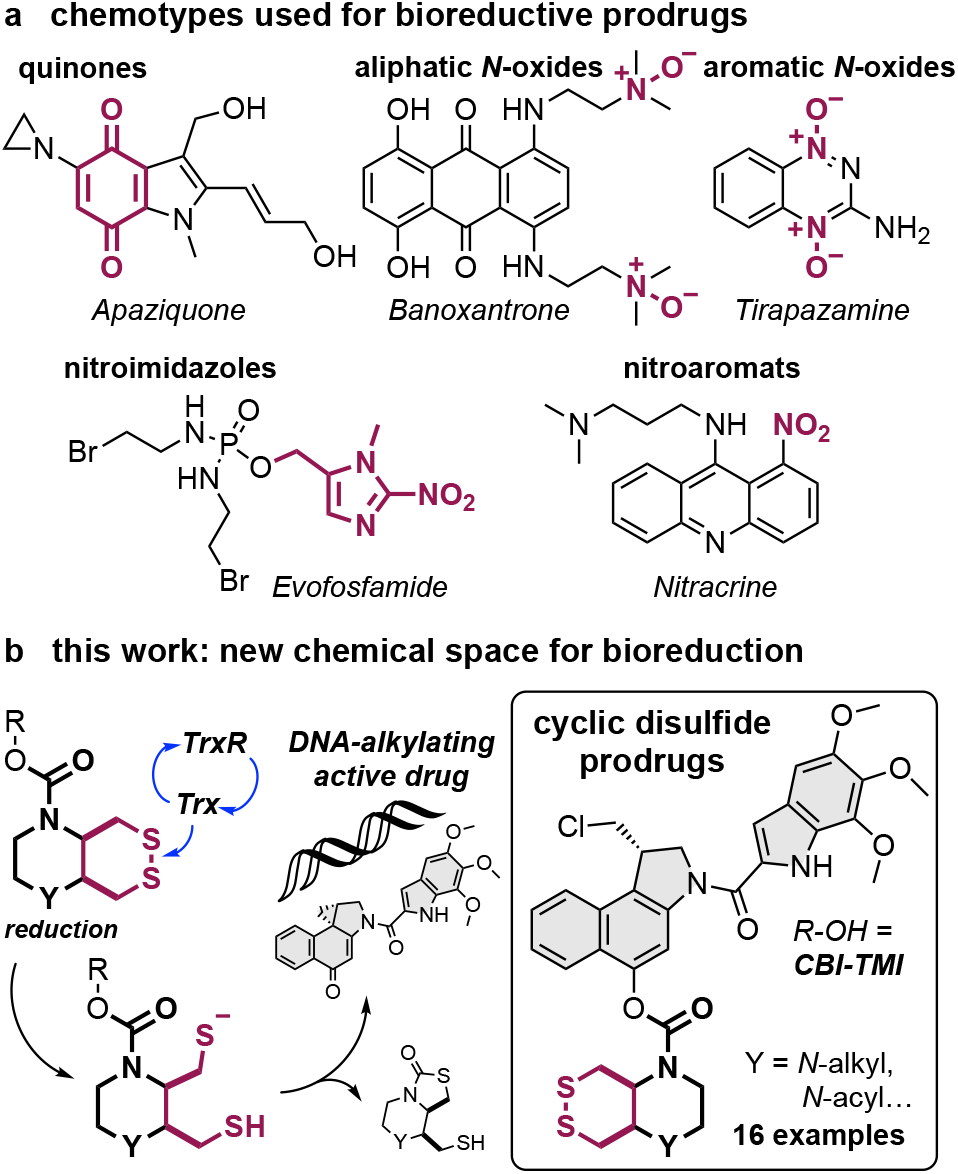
Bioreductive prodrugs. **(a)** Chemotypes used for bioreductive clinical prodrugs: i) aliphatic *N*-oxides, ii) aromatic *N*-oxides, iii) nitroimidazoles and nitroaromats. **(b)** Bicyclic disulfides targeting the Trx/TrxR system: one of the new motifs for bioreductive prodrug development introduced in this paper.

The thioredoxin system, composed of the thioredoxins (Trxs) and their NADPH-dependent reductases (TrxRs), are enzymatic reductants that are central to redox homeostasis, cellular metabolism, DNA synthesis, protein folding, and antioxidant response.^11^ Overactivity of the thioredoxin system is strongly implicated in cancer progression^12^ and Trx expression is often upregulated in tumors.^13^ This may offer a unique point of chemical attack: Trx is the cell’s strongest dithiol-based reductant, so reduction-resistant substrates, that are selectively but only slowly activated by Trx under physiological conditions, might be faster activated by Trx in diseased tissues. However, bioreductive prodrug approaches for the Trx system have not yet been designed.

Chemically, Trx/TrxR are dithiol/selenolthiol reductases that can address protein as well as small molecule disulfides. The biological target space tested by (any) disulfide-based prodrugs has remained limited, as essentially only GSH-labile nonspecific linear disulfides^14^ and 1,2-dithiolanes^15,16^ have been examined. Recently, we have developed sets of 6-membered cyclic dichalcogenides with unique reduction-resisting properties as well as reductase selectivity profiles.^17–19^ Stabilised disulfides in these series were selectively activated by Trxs, with excellent resistance to even thousand-fold higher levels of GSH and monothiols,^17^ and bifunctional bicyclic disulfides permitted drastic enhancement of their reductive activation kinetics.^19^ Cyclic selenenylsulfides instead had a selenium-preferring, regioisomer-dependent activation mechanism making them excellently selective substrates for TrxR1 in live cells.^18^ These bioreductive motifs were validated in fluorogenic probes in acute applications (minutes to hours) in cell culture. However, it is unknown whether such designs can be useful for long-term redox-selective drug delivery, particularly in the context of cancer for which their Trx/TrxR targets’ biology is most relevant (**Fig 1b**), since *in vivo* uses are more stringent: e.g. requiring them to also resist activation by hydrolytic or oxidative metabolism in the long-term.

The duocarmycins are DNA alkylators with high potency across a broad range of cell lines,^20^ that have excellent characteristics as modular payloads for cytotoxic anticancer prodrugs.^21^ Their key motif is an activated cyclopropane electrophile, which can be generated *in situ* from a biologically inactive *seco*-precursor by unmasking a phenol or aniline that then spontaneously undergoes phenylogous cyclisation (**Fig 2**).^22^ Due to this elegant off-to-on bioactivity switch, synthetic *seco-*duocarmycin-type prodrugs have been broadly developed as stimulus-responsive proagents, often harnessing the simpler **CBI**^23,24^ alkylator and the minimal **TMI**^25^ DNA binder motifs.^25^ Hydrolase-unmaskable phenolic substrates such as esters, carbamates^26,27^ and glycosides^28,29^ have been broadly applied. Bioreductive substrates aiming at increased tumor-specificity have also been tested, including nitro-*seco*-CBIs as aniline precursors^30,31^ and *N-*acyl-*O-*amines as phenol precursors.^32,33^ The modular “puzzle” design of *seco-*duocarmycins, combining a DNA binder with an electrophile segment and its unmaskable trigger, sets up a platform approach for prodrug development (**Fig 2b**).

**Figure 2:**
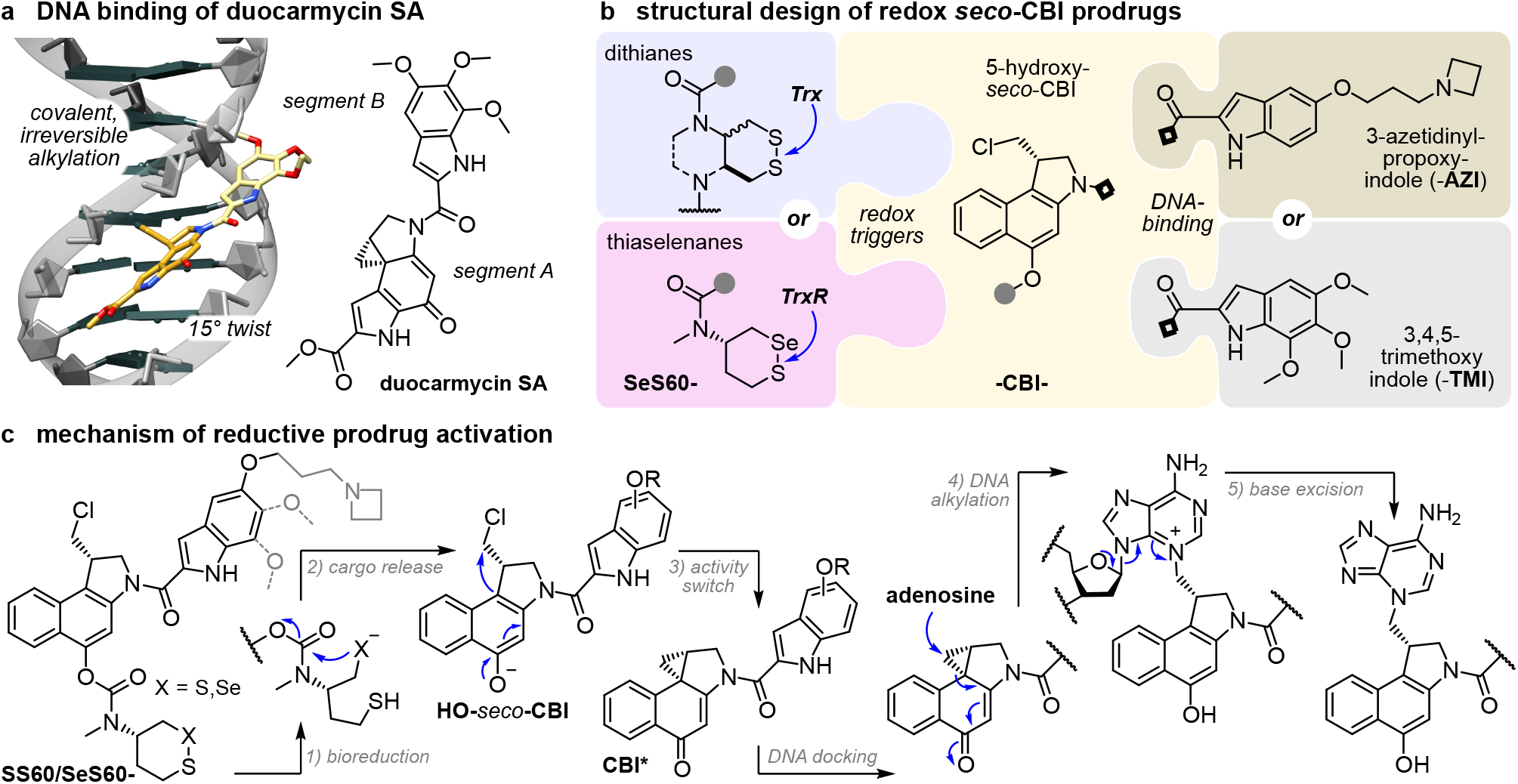
*seco*-duocarmycin prodrug design and mechanism of activation. (**a**) DNA docking by duocarmycin SA with a 15° amide twist that initiates DNA alkylation (PDB: 1DSM). **(b)** Our three-part bioreductive prodrug design: (i) an enzyme-selective dithiane/thiaselenane masks (ii) the *seco*-**CBI** phenol to which (iii) an indole segment is attached for DNA binding and/or solubility. Prodrugs are named by parts, e.g. **SeS60-CBI-AZI** (thiaselenane trigger **SeS60**, -**CBI-**, and 3-<u>az</u>etidinyl-propoxy-indole **AZI**). (**c**) Reductive activation: (1) enzymatic bioreduction; (2) the reduced chalcogenide cyclises to expel the **CBI** phenolic leaving group; (3) bioactivity switch: intramolecular Winstein cyclisation creates the activated cyclopropane; (4) DNA docking leads to DNA alkylation and (5) base excision, irreversible DNA damage, and apoptosis^34^.

Aiming to test the potential of Trx/TrxR-responsive cyclic dichalcogenides as trigger units that may open new chemical space as well as target spaces for bioreductive prodrugs, we now design and test a suite of ten redox-responsive **CBI**-based therapeutic prodrugs based on cyclic dichalcogenides (**Fig 2**).

## 2. RESULTS AND DISCUSSION

### 2.1 Modular design logic for dichalcogenide prodrugs

We aimed at modular prodrug designs, separating the Trx/TrxR-bioreducible trigger module from its duocarmycin cargo module. This design also separates the functional steps responsible for bioactivity (**Fig 2c**): (1) bioreduction of the trigger by dithiol or selenolthiol reductases e.g. Trx^17^ or TrxR^18^; (2) 5-exo-trig cyclisation by the trigger to release the (biologically inactive) duocarmycin phenolic cargo; (3) 1,4-nucleophilic Winstein cyclisation^35^ creates the activated cyclopropane CBI*; (4) CBI* docks in the DNA minor groove while twisting its aryl-aryl plane, and the now more activated cyclopropane irreversibly alkylates adenosine at the N3-position;^36^ (5) the quaternised purine base is spontaneously excised, causing DNA damage and ultimately cell death.

We expected that varying trigger and cargo motifs would clarify the separate contributions of reduction and of drug sensitivity to the overall prodrug efficacy, thus setting a rational basis both for further tuning of dichalcogenide triggers, as well as for their adaptation to alternative bioactive cargos^37^. This is because, in our model (**Fig 2c**), the trigger mainly determines the rate or degree of cargo release, by controlling reduction and cyclisation rates (steps 1-2); whereas the cargo mainly determines the expected potency for a particular degree of drug release, by controlling the speed of cyclopropane formation and likelihood of DNA binding/alkylation (steps 3-4).

We therefore planned to use a range of dichalcogenide redox triggers (e.g. dithiane **SS60-**) to covalently mask hydroxy-*seco*-CBI (**-CBI-**) by a stable tertiary carbamate linkage,^38,39^ while varying the DNA-binding indoles that complete the duocarmycins to modulate the ADME/potency of the prodrug/drug (e.g. -**AZI**). The assembled prodrugs would then be named as the combination of these abbreviations: e.g. **SS60-CBI-AZI** (**Fig 2c**).

### 2.2 Choice of modules for dichalcogenide prodrugs

Choosing trigger motifs with the right redox selectivity profile and cargo release kinetics is key to success of the prodrug design. Our redox trigger choices were based on results from short-term assays relying on phenol-releasing fluorogenic probes (**Fig 3**). Cyclic 6-membered disulfides (**SS60** or its faster-activated, solubilised analogue **P-SS60**) were used as slowly/moderately-reduced Trx-selective substrates (minor crosstalk to dithiol Grx is expected^17^). Their unstrained *cis-*annelated-piperazinyl bicyclic congeners (**P-SS66C, Me-SS66C**) have much faster, though less selective, Trx reductive activation (more crosstalk to dithiol Grx); strained *trans-*annelated congeners (**Me-SS66T**) are likewise rapidly activated by reductases, but are also moderately labile to monothiols such as GSH so were expected to produce more unspecifically toxic prodrugs. Separately to these Trx-targeted triggers, we included the specifically TrxR1-activated cyclic 6-membered selenenylsulfide **SeS60**, to probe the performance and selectivity possible by targeting the regulator (rather than effector) of the thioredoxin system (**Fig 3**).

**Figure 3:**
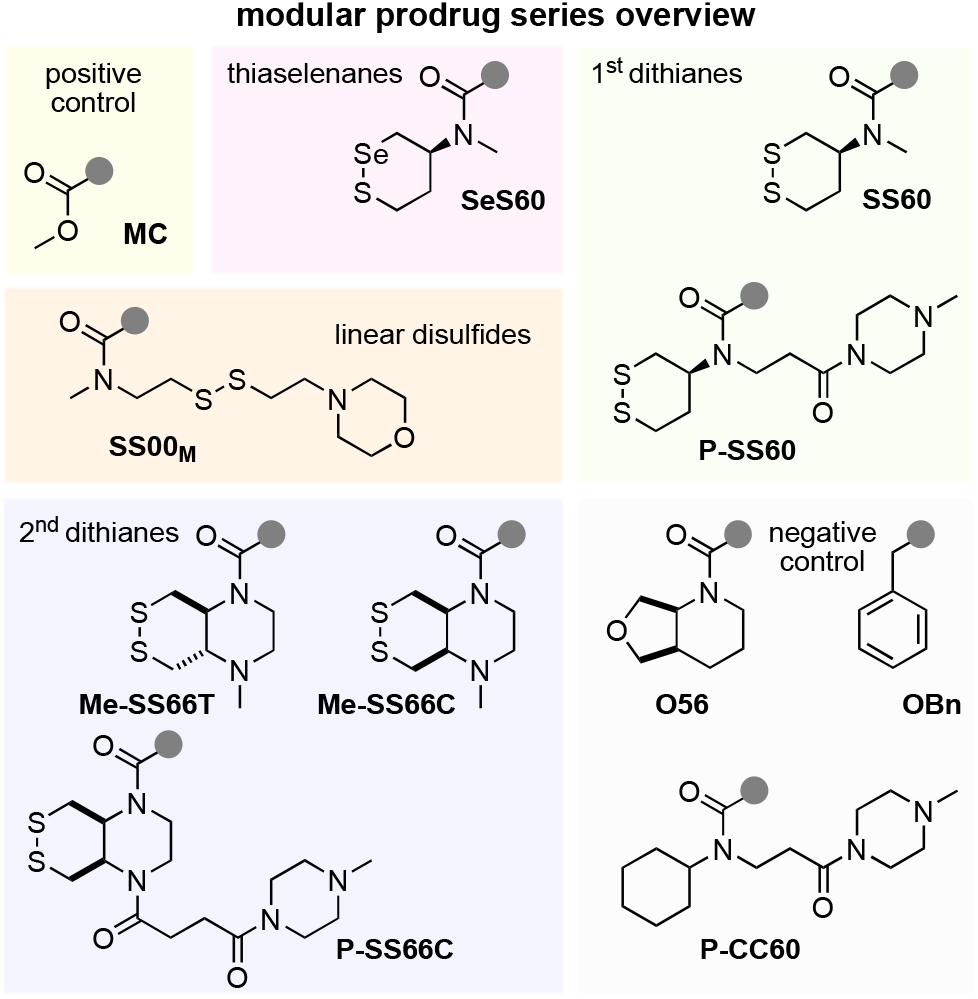
Redox triggers for modular prodrugs. 1^st^ generation Trx-triggers **SS60** and solubilised **P-SS60**;^17^ 2^nd^ generation solubilised bicyclic Trx-triggers **Me-SS66T, Me-SS66C** and **P-SS66C**;^19^ TrxR-trigger **SeS60**^18^. “Upper limit” control triggers: very labile **MC**, monothiol-labile **SS00**_**M**_. “Lower limit” control triggers: hydrolysis benchmarks **O-56** and **P-CC60**, and carbamate stability test **OBn**.

To benchmark the performance of these redox-activated triggers and to separate the contributions of reductive vs non-reductive activation (hydrolytic, oxidative, etc.), we designed “upper limit” and “lower limit” reference triggers. At the extreme end of unspecificity, we used linear disulfides (e.g. **SS00**_**M**_) which are unselectively reduced by any biological thiol,^17^ as well as a rapidly and unspecifically hydrolase-cleavable carbonate (**MC**), to test the potency expected for the fully released drug cargos and so to estimate the degree of release from the hopefully redox-selective triggers. To understand the degree of *non-reductive* drug release which contributes to the prodrugs’ overall cytotoxicity, we would compare the cytotoxicity from a non-reducible isosteric carbamate (**O56**) or a solubilised analogue (**P-CC60**), to a more-resistant ether (**OBn**; **Fig 3**). An overview of the redox and control triggers is given at **Fig S1** where their performance features are detailed.

To complete the prodrugs, we equipped hydroxy-*seco*-CBI with two types of “segment B” indole (**Fig 2b**). For a benchmark that can be compared to literature results, we used the classic trimethoxyindole **TMI** residue, as found in the original natural products duocarmycin SA/A.^22^ The lipophilic **CBI-TMI** tends to aggregate, so this cargo was only employed with the solubilised trigger motifs. To evolve the properties of synthetic duocarmycins, we also designed a novel abiotic segment B, 3-azetidinyl-propoxy-indole (**AZI**), which is the first azetidine used on a duocarmycin. Our hopes were that the basic amine would (i) add solubility so that non-solubilised triggers (e.g. **SS60, SeS60**) could be incorporated into useful and bioavailable **AZI** prodrugs; and also (ii) increase DNA association by Coulombic interactions, and so raise potency. These two features are known from e.g. dimethylamine-substituted B segments (“DEI”).^28^ However, our design also aimed to (iii) avoid undesirable metabolic attack/demethylation *in vivo*, which we expected might be useful for its *in vivo* performance. For an overview of all the resulting prodrug combinations, see the discussion at **Fig S3**.

### 2.3 Prodrug assembly by a versatile 2-step approach that avoids dangerously toxic intermediates

Commercial *O*-benzyl-*N*-Boc-(*S*)-*seco*-CBI (**BnO-CBI-Boc**) **1** served as a synthetic starting point (**Fig 4**). While the prodrug retrosynthesis is straightforward, we wished to avoid handling any directly potent DNA alkylators during synthesis, i.e. we aimed to avoid free 5-hydroxy-CBI-indoles that are otherwise the most easily diversified synthetic intermediates: so we wanted to install the triggers before coupling to the indoles. We performed *O-*debenzylation by mild heterogenous hydrogenation on Pd/C using aq. NH_4_HCO_2_ as the hydrogen source, as reported by Major^28^, to avoid the unwanted naphthalene hydrogenation and dechlorination seen with H_2 (g)_. The phenolic chloroformate produced by reaction with *in situ* generated phosgene was carbamylated^17,18^ with the eight trigger secondary amines giving good to excellent yields of trigger-CBI intermediates (e.g. trigger **H-SS66C-H** (**2**) giving intermediate **H-SS66C-CBI-Boc** (**3**), **Fig 4**). The bisamine SS66-type triggers could then be additionally functionalised by reductive amination (e.g. with formaldehyde giving **<u>Me</u>-SS66-CBI-Boc** species) or acylation (e.g. with acid anhydride **4**^40^ giving **<u>P</u>-SS66-CBI-Boc** species). The indolecarboxylic acid segments had then to be coupled. **TMI-OH** is commercial; we prepared **AZI-OH** by a 4-step literature procedure^41^ that we adapted into a 3-step sequence, using Mitsunobu-type *O*-alkylation of a 5-hydroxy-1*H*-indole-2-carboxylic ester with commercial 3-azetidinyl-propan-1-ol.^42^ Finally, the trigger-CBI intermediates were *N*-Boc-deprotected with HCl in organic solutions^43^, with TFA or with BF_3_·OEt_2_^29^, then coupled either to **AZI-OH** with similar conditions as reported by Tercel^31^ or else to the acid chloride **TMI-Cl**^44^, giving the sixteen prodrugs and controls used in this study (**Fig S3**) with moderate overall yields (**Fig S2**). Note that even traces of residual **CBI-OH, CBI\*** cyclopropane, or easily cleaved CBI byproducts, must be avoided for cellular and *in vivo* testing: their much higher potency can overpower the true performance of the major species prodrugs.

**Figure 4:**
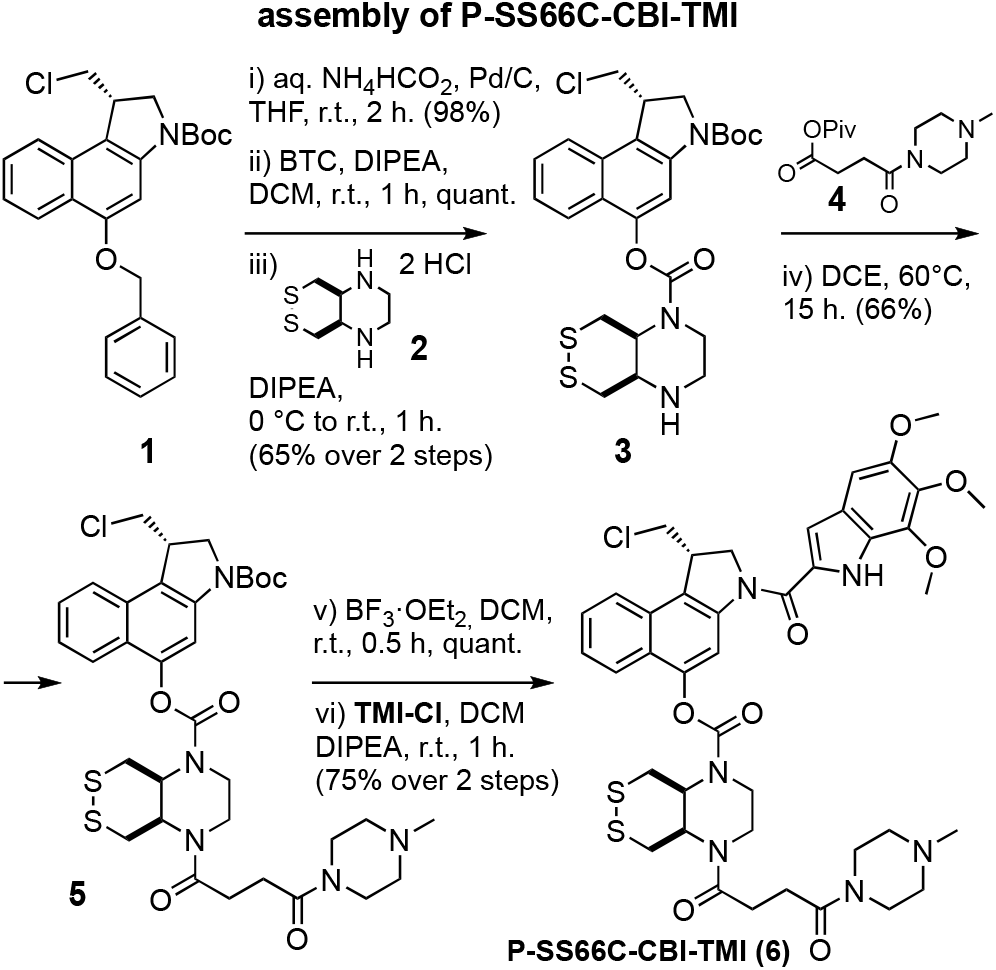
Modular *seco-*CBI prodrug synthesis. Synthesis of representative prodrug **P-SS66C-CBI-TMI** from commercial **BnO-CBI-Boc 1** (full synthesis overview in **Fig S2**-**S3**).

### 2.4 The prodrugs are not activated by monothiols, so may be specifically reducible by disulfide reductases in cells

The cyclic dichalcogenide redox triggers were previously shown to resist activation by monothiols such as glutathione (GSH), but to allow reduction/cyclisation-based release of phenolic fluorophore cargos when treated with disulfide reductase enzymes (e.g. Trx, and/or TrxR, Grx, etc.).^17,18^ These performance features had not been tested with naphthols, so we aimed to use HPLC to confirm them while also testing their reduction-based activation sequence (**Fig 2c** and **Fig S4-S6**).

In brief, when prodrugs were treated with the quantitative reductant TCEP, we could confirm that the evolution of species over time matches the activation sequence we expected (**Fig 2c**): dichalcogenides are reduced to dichalcogenols, then giving the naphthols (plus the monothiol cyclisation byproducts), then the activated cyclopropanes (**Fig 5**). Also matching expectations, the redox triggers which previously resisted monothiol-reductant-based release of phenolic cargos (**SS/SeS60** and **SS66C** types) likewise gave no detectable release of 5-hydroxy-*seco*-CBI or cyclopropane CBI* when challenged with 5 mM GSH (50 eq.) over 24 h, though partial activation by GSH was indicated for the strained **SS66T** (**Fig S6**). The reference compounds performed as expected, either being sensitive to GSH (linear disulfide **SS00**_**M**_) or else resistant to all reduction conditions (non-chalcogenide **O56** or solubilised **P-CC60**).

**Figure 5:**
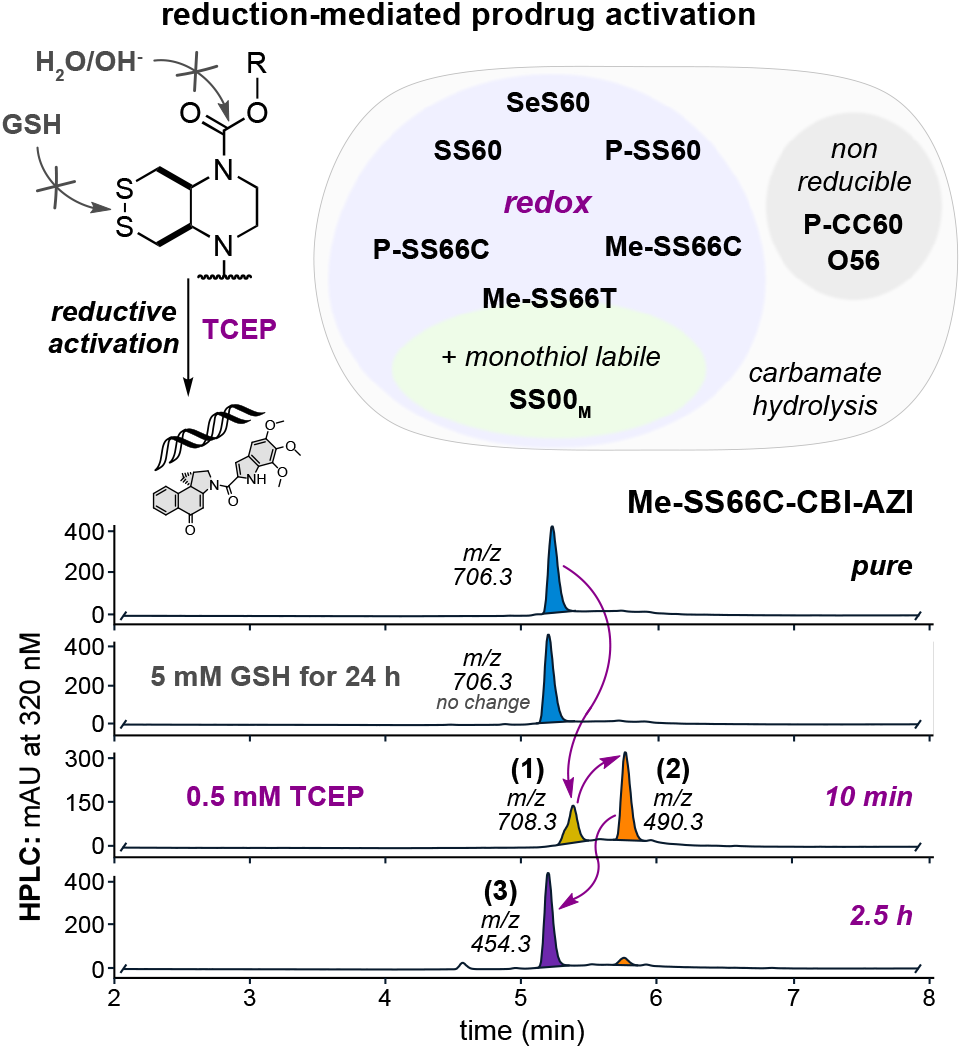
Reduction-triggered prodrug activation. HPLC-MS analysis of activation of **Me-SS66C-CBI-AZI**: (1) TCEP reduction gives the dithiol, (2) the phenolate is expelled, (3) Winstein cyclisation gives the activated cyclopropane (see **Fig S4-S6**).

Taken together, this indicated that the **(P/Me)**-**SS60, -SeS60, -SS66C** prodrugs would indeed be monothiol-resistant, giving potential for them to act as prodrugs that are selectively activated by dithiol reductases of the Trx system in cells; and that the reference prodrugs could be used as intended, to estimate the high potency expected if activated by all thiol reductants (**SS00**_**M**_), or the low potency expected if only activated by non-reductive mechanisms, e.g. hydrolysis (**O56, P-CC60**) (**Fig 5**).

### 2.5 Tunable reduction-based activity allows rational selection of reducible motifs for future *in vivo* applications

While cell lines vary in their intrinsic sensitivity to duocarmycin drugs, the *relative* toxicity of duocarmycin-type *prodrugs, within* any single cell type, should reflect their degrees of cellular activation. We were particularly interested in “low potency” prodrugs, with low activation in usual cell cultures. As a counterexample, prodrugs that are fully activated in all cell types will have high cell culture potencies, like their free duocarmycin cargo: but they are also likely to be activated in all tissues during systemic treatment *in vivo*, so causing dose- and therapy-limiting toxic side-effects.^45,46^ Instead, a prodrug that is little activated in most or all 2D cell cultures (lower potency than its free cargo) may escape such broad activation *in vivo* and be well-tolerated; and if it is distributed to tumors with a suitable bioreduction profile, it may be selectively activated there: so delivering therapeutic benefits. Comparing the relative potencies of duocarmycin prodrugs across cell cultures may give valuable indications about their likely tolerance in therapeutic settings.

For example, free CBIs (cellular IC_50_ ca. 5-30 pM^24^) and hydroxy-*seco*-CBIs (5-50 pM^34,47^) have low tolerated dosages *in vivo*; but their prodrugs are increasingly tolerated, as their cellular activation is reduced from high (esters for esterases: 100-500 pM^26^), to moderate (tertiary carbamates for oxidative and/or peptidolytic processing: 50-300 nM^26^), to low (glycosides for glycosidases: 5-10 μM^28^). Amino-*seco*-CBIs are less efficient alkylators than the phenols (100-500 pM^30^); their prodrugs can also be tuned for low nonspecific release (nitro prodrugs for metabolic reduction: 5-50 μM^31^). **Table 1** summarises these approximate average IC_50_s over a range of cell lines, highlighting how duocarmycin prodrug potency can be tuned over ca. 7 orders of magnitude by the choice of bioactivation strategy.

**Table 1:**
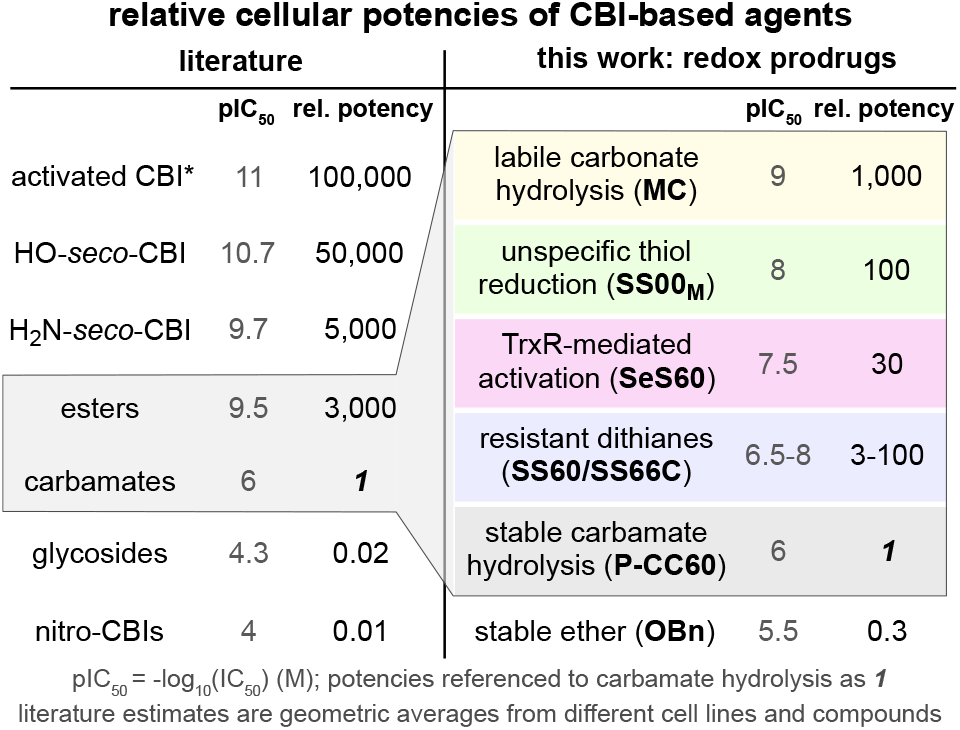
Benchmark potencies of CBI prodrug classes in cell culture. Averaged literature potencies for CBIs and their prodrugs, to put the potencies of the tertiary carbamate redox prodrugs of this work into perspective. Recall that potency maximisation within the redox prodrug series is *opposed to* the aim of this work (section 2.5).

We wished to understand the degree of bioreduction-mediated activation across the series of novel dichalcogenide prodrugs by studying their cellular potencies relative to “minimum/maximum-potency” reference compounds. Our redox-activatable prodrugs are tertiary carbamates, which will have some activation by hydrolysis, as well as by bioreduction. We thus expected their cell culture potencies to fall between a minimum for the non-reducible hydrolysis-only carbamates **O56** and **P-CC60** (ideally: low potency, anticipating low systemic release), and a maximum for the rapidly enzyme-hydrolysed carbonate **MC** (bioreductive activation unlikely to be faster than this): within which range, variations of potency would report on their relative reductive stability or lability. We also tested nonspecifically-reducible linear disulfide **SS00**_**M**_ prodrugs, to probe our expectation that only the monothiol-resistant dichalcogenides such as those we recently introduced^17,18^ can access a different performance space than prior, linear disulfides.

We used A549 lung cancer and HeLa cervical cancer cell lines for initial screening of the prodrugs. These had a >1000-fold range of potencies: rapidly hydrolysed **MC** set an enzymolytic maximum (IC_50_ ca. 0.5 nM; **Fig 6a**) that can be approached by linear disulfide **SS00**_**M**_ (nonspecific redox activation by thiols), while the non-reducible carbamates set a hydrolysis-only minimum (ca. 300 nM for **O56-CBI-AZI**, >1 μM for **P-CC60-CBI-TMI**) that was similar to ether **OBn-CBI-AZI** (>1 μM). We thus estimated that the tertiary carbamate design, with its excellent cell culture stability, offers a window of up to 1000-fold toxicity enhancement for the dichalcogenide prodrugs according to how much reductive release they undergo (**Fig 6a**). (The 3-10× higher potency and better reproducibility of our novel, solubilised **AZI-** as compared to **TMI**-series prodrugs are additional positive features).

**Figure 6:**
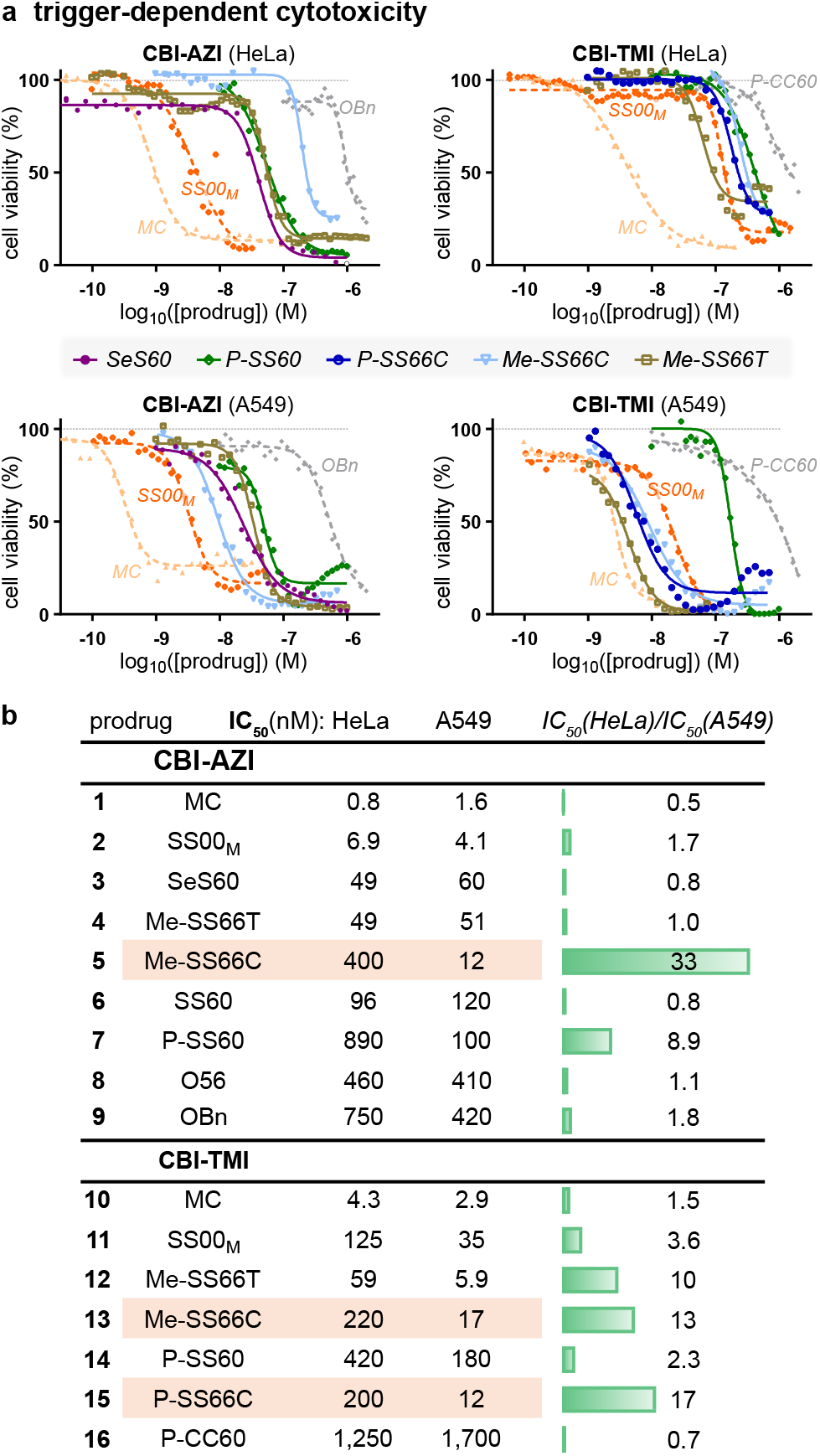
Trigger-dependent cellular cytotoxicity. **(a-b)** Potency in A549 / HeLa cells. Note e.g. consistent 13-to-30-fold greater toxicities in HeLa than in A549 cells for **SS66C** triggers (**AZI** or **TMI** cargos, **P-** or **Me-** substituents) (see also **Table S1** and **Fig S7-S8**).

The toxicities of the novel cyclic dichalcogenide prodrugs (**Fig 6b**) revealed increasing redox-based release in the order **SS60**≈**SS66C** (Trx-activated) **< SS66T** (Trx-activated plus some monothiol sensitivity) < **SeS60** (TrxR-activated with high turnover) < **SS00**_**M**_ (monothiol-activated limit): matching the order in acute assays of their corresponding fluorogenic probes^17,18^. This supports our hypothesis that engineered dichalcogenides can resist non-specific thiol-mediated release even in long-term assays (IC_50_s below **SS00**_**M**_). We consider that by avoiding substantial activation in usual cellular conditions (ca. 30-300-fold less release than references **MC**/**SS00**_**M**_), they keep alive the possibility of enhanced release selectively in tumors, as long as their bioreduction sensitivity profile suits the potentially increased expression and/or activity of reductases in tumor environments.

### 2.6 Redox activation is tied to thioredoxin system activity

Testing whether the cellular activation of the cyclic disulfide prodrugs is due to their intended target, thioredoxin, is a non-trivial task: since there are no stable Trx knockouts, nor are pharmacologically clean cellular Trx inhibitors available. A TrxR1 knockout (ko) in mouse embryonic fibroblast (MEF) cells is however accessible. We compared its prodrug sensitivity to that of its parental line^48^ (wt), on the basis that TrxR1-ko should leave most Trx1 in its oxidised state (only a fraction of it can be maintained in the reduced state by alternative reductants such as Grx2)^49^, so prodrugs relying on the cytosolic thioredoxin system (Trx1/TrxR1) for most of their bioreductive activation in long-term assays will have lower potency in TrxR1-ko cells.

The overall MEF potency trends were similar as in HeLa/A549 cells (pIC_50_: **MC > MeSS66T** > **Me-SS66C** ≈ **P-SS66C** > **P-SS60** > **P-CC60**; **Fig 5c** and **Fig S10**). Excitingly, all five prodrugs based on the bicyclic **SS66**-type trigger, which have the lowest reduction potential of the disulfide series, were many fold less potent against TrxR1-ko than wt cells (**Fig 7**). This is consistent with reductive processing of **SS66** strongly requiring thioredoxin system activity. That is not an obvious result: since, the multi-day assay provides plenty of time for dithiol reductases outside the TrxR1/Trx1 system (e.g. Grxs) to perform reductive activation instead. Indeed, the three less-stabilised monocyclic **SS60**-type prodrugs showed no such fold-change of potency (see **Table S1** and **Fig S9**), suggesting that in long term assays, cellular activation of simple dithianes can proceed through multiple redox paths. We also find it satisfying that there is such a clear division between the **SS66** and **SS60** structural classes. No matter which cargo (**AZI** or **TMI**) and what variable substituents (**P-** or **Me-** type) are employed, it is the core chemical nature of the redox trigger that dictates cellular performance. Although some redox papers have resisted the idea that SAR rules operate for reducible probes^16^, just as they do for drugs, we believe that such consistent patterns will continue to emerge and to enlighten the field, wherever design and testing are performed comprehensively.

**Figure 7:**
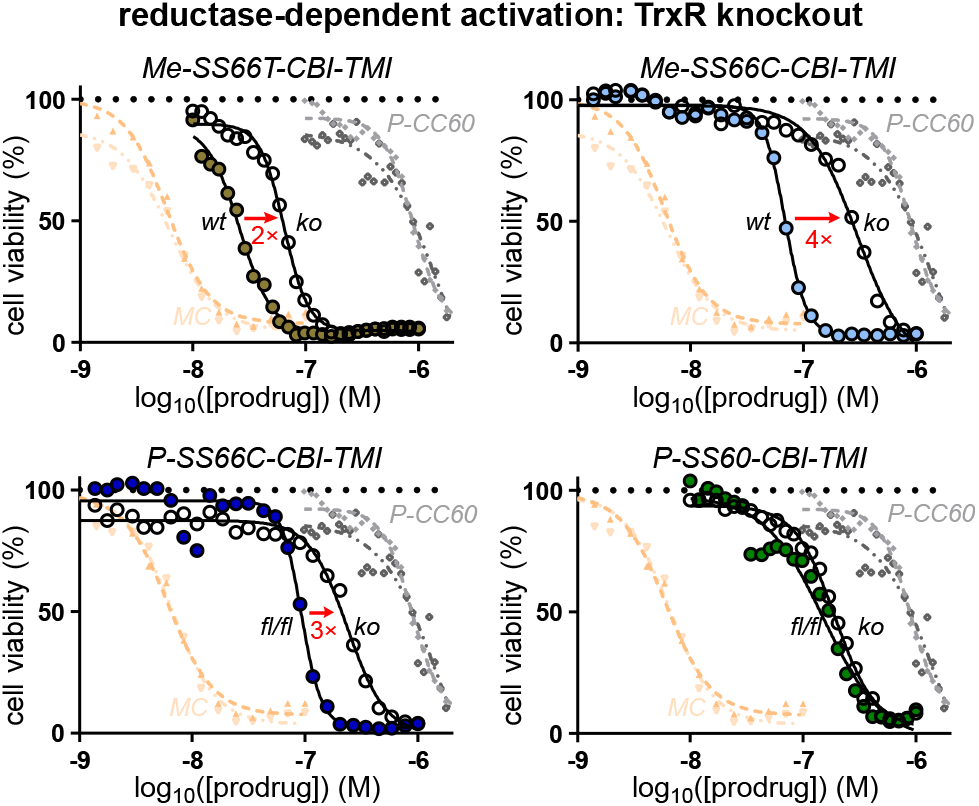
Target-dependent cellular cytotoxicity. Comparing prodrug toxicities in intact (*wt*) vs TrxR1-knockout (*ko*) MEF cells, to test whether the major cellular activation route is rate-limited by the cytosolic thioredoxin couple (TrxR1/Trx1). That is indicated for bicyclic -**SS66C** and **-SS66T**, but not monocyclic -**SS60** (see also **Table S1** and **Fig S9**).

Taken together, these assays had validated the hypothesis of tunable redox-based cellular activation of dichalcogenide produgs. Consistent with our aims, the **SS66**-type strongly depend on a specific, key enzyme pair: the thioredoxin system; even in long-term assays (**Fig 7**). This validates the first bioreductive prodrug design capable of selectively targeting a disulfide reductase in cells.

### 2.7 Profiling thioredoxin- and redox-activated prodrugs in 176 cancer cell lines suggests their *in vivo* potential

We wished to study the *in vivo* suitability of the dichalcogenide strategies, while benchmarking the performance of the different triggers in a thorough and reproducible manner that can be used to guide rational future design of other prodrug families. We therefore planned to screen the prodrugs’ redox-dependent bioactivity across large numbers of validated cancer cell lines with standard and objective assay conditions.

We initially focused on **P-SS66C** (reducible by thioredoxin) and **P-SS60** (reducible by thioredoxin and unknown other actors). Both designs will also undergo non-reductive release by carbamate cleavage (hydrolases and/or oxidative metabolism), at rates that are likely to vary with cell type. Other factors strongly depending on cell type include intrinsic cellular sensitivity to their duocarmycin cargo, prodrug rates of cellular entry, and degree of prodrug cellular accumulation. To control for all these cell-line-dependent features, we included the closely similar but non-reducible carbamate analogue, **P-CC60**. We think this is a vitally important step. Rather than focusing on the *absolute* potencies of a redox prodrug in a certain cell line, we could then examine the *fold difference* of potency between reducible **P-SS66C** or **P-SS60** vs. non-reducible **P-CC60**, to focus on the degree of bioreductive processing that the prodrugs undergo in cell culture (see below).

Note that while expression levels of some reductases have been measured on mRNA and protein levels, their actual *activity* levels are unknown - both in clinically relevant samples (e.g. patient tumors), but also even in simple 2D cultures of cell lines - due to the lack of suitable tools to quantify them.

We therefore obtained a first high-throughput automated screen of the antiproliferative potencies of reducible **P-SS66C-CBI-TMI** and **P-SS60-CBI-TMI** alongside **P-CC60-CBI-TMI** as a hydrolysis-only reference, over a panel of 140 standard cancer cell lines with diverse tissue origins (skin, ovary, lung, colon, breast, pancreas, prostate, kidney, brain, etc), conducted by commercial provider Reaction Biology (“ProliFiler-140” screen). The topoisomerase inhibitor doxorubicin was included as a benchmark for assay validation, and to illustrate general trends in cell line drug sensitivity. Incubations were performed for 72 hours before automated viability readout with CellTiterGlo® (**Fig 8**).

**Figure 8:**
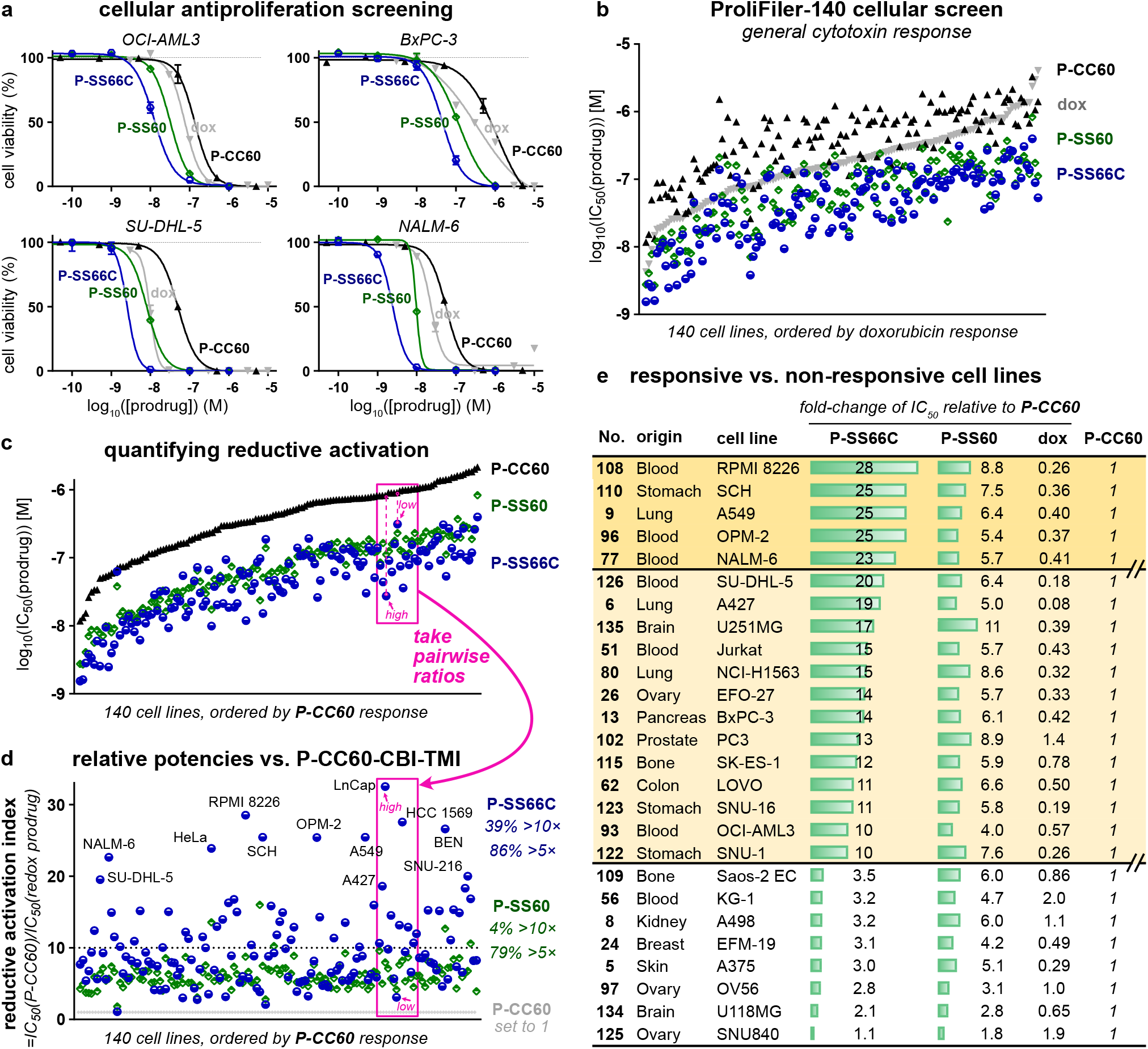
Screening bioreductive activity patterns for P-SS66C-CBI-TMI and P-SS60-CBI-TMI redox prodrugs, by benchmarking against hydrolysis control P-CC60-CBI-TMI, across 140 cancer cell lines (“ProliFiler-140”). (**a**) Sample cell lines, of the 140 tested (see too **Table S2** and **Fig S10-S11**). (**b**) Rank ordering by sensitivity to the mechanistically unrelated drug doxorubicin. (**c**) Rank ordering by sensitivity to hydrolysis control **P-CC60**. (**d**) IC_50_s as ratios relative to that of **P-CC60** hydrolysis control. (**e**) Sets of cell lines suggest different redox activity patterns.

All 140 cell lines gave well-formed and complete sigmoidal dose-response curves with steep and uniform Hill slopes, consistent with excellent assay technical performance, including compounds being well soluble at all concentrations (selection in **Fig 8a**, full data in **Table S2** and **Fig S10**). Antiproliferative IC_50_ values range from ca. 1-100 nM for reducible **P-SS60** and **P-SS66C**, and ca. 10-1000 nM for non-reducible **P-CC60** (**Fig 8b**). Prodrug potencies were slightly correlated to those of doxorubicin (**Fig 8b**), but very much better correlated to **P-CC60** (**Fig 8c**), matching expectations from their overall similar structures and shared mechanism of action. Noteworthily, **P-SS60** potency correlated more tightly than **P-SS66C** to that of **P-CC60** (green vs. blue data, **Fig 8c**). This difference now gave broad support to the indication from the single cell line TrxR knockout assay (**Fig 7**), that their bioreductive activating mechanisms in long term assays are substantially differentiated.

Matching our model, the potencies of reducible **P-SS60** and **P-SS66C** were greater than of non-reducible **P-CC60** *within* each one of the 140 cell lines, even though their absolute potencies vary over more than 100-fold *between* the different cell lines. This supports **P-CC60** being a suitable predictor of minimal, purely non-reductive release levels in any given cell line: compared to which, any additional reductive release is reflected in increased potency of the reducible prodrugs (**Fig 8a,c**).

This additional reductive release is, in our opinion, the most important data delivered by the screening. To analyse it, we define a prodrug’s “reductive activation index” in a given cell line to reflect how many fold more potent it is than the hydrolytic control: i.e. index = IC_50_[(**P-CC60**)/IC_50_(prodrug)] (**Fig 8d**). This is a qualitative indicator of how much reductive activation a prodrug experiences, benchmarked to unknown but variable level of hydrolytic cleavage, in a given cell line.

Trends emerge when the reductive index is viewed across so many cell lines (selection in **Fig 8e**, full list in **Table S2**): (1) The index of **P-SS66C** is variable with some cell lines reaching up to 30-fold; and there is no relation between which cell lines have a high index, and which are most sensitive to **P-SS66C** in an absolute sense (**Fig 8d**). This contrasts to **P-SS60** whose index remains in a narrow band between 5 and 8 over nearly all cell types. **Fig 7** had indicated that TrxR1 activity strongly impacts the bioreductive activation of **P-SS66C**, but not of **P-SS60**. Thus, it is tempting to interpret the index of **P-SS66C** as reporting substantially on variations of specific TrxR1 activity, and to see the index of **P-SS60** as reporting on other bioreductive actors which seem more constant. (2) The cell lines can be grouped by suggestive trends in the reductive index. For example: (i) *high index for* ***P-SS66C*** *but not* ***P-SS60*** (**Fig 8e**, top bracket): suggesting cell lines where comparatively high reductive activity driven by TrxR1 could be harnessed with novel bioreductive prodrug strategies such as the modular Trx/TrxR-dichalcogenides we present; (ii) *index for* ***P-SS60*** *is similar to that of* ***P-SS66C*** (**Fig 8e**, middle bracket): suggesting significant disulfide bioreduction activity outside the Trx1/TrxR1 couple, that may be exploitable by future prodrugs tuned towards other reductases e.g. Grxs; or, (iii) *low indices for both redox prodrugs* (**Fig 8e**, bottom bracket): bioreduction is outweighed by other cleavage mechanisms, so dichalcogenide prodrugs may be unsuitable for addressing these cell types.

We also used this large dataset to check for biological features that might correlate usefully to reductive activation. First, we tested whether reductive performance was clustered according to the tissue of origin of the cell lines, since any clustering could be suggestive of e.g. cancer indications that might be promising for selective targeting by reductive prodrugs. However, the tissue of origin played no role in either the reductive index or the absolute potency of the compounds tested (**Fig S11**). Second, we wished to examine whether reductive performance was clustered according to gene expression patterns, since such clustering could suggest biomarkers predictive for response to redox prodrugs, and so orient their therapeutic opportunities. However, mRNA-based gene expression analysis did not return results that could be confidently interpreted; future testing in this direction should probably rely instead on fluorogenic probes (notes at **Table S2**).

We next wished to test the reproducibility and validity of these results by standardised screening in a different lab and location, while scrutinising in more detail the bicyclic **SS66** structure which had indicated tantalising TrxR/Trx-selectivity. We obtained a second high-throughput automated screen of antiproliferative potencies through the non-commercial NCI Developmental Therapeutics Program (DTP), over the NCI-60 standardised human tumor cell line panel.^50^ 19 of the NCI cell lines overlap with the ProliFiler-140 screen so could be used for benchmarking, though differences in assay setup, run time, and readout introduce systematic shifts in absolute potencies. In a series of **-CBI-TMI** prodrugs, we newly tested **Me-SS66C-** and its more reduction-prone diastereomer **Me-SS66T-** (basic amines at the redox site), in comparison to previously tested **P-SS66C-, P-SS60-**, and non-reducible **P-CC60-** (all amides; **Fig 3**).

The new prodrugs showed dramatically how stereochemistry can rule reductive lability. *Trans-*dithiane **Me-SS66T** was consistently far more potent than any other drug, with a mean reductive index ca. 70; while *cis-*dithiane **Me-SS66C** was roughly similar in potency to **P-SS60** with mean index ca. 7 (**Fig 9a**).

**Figure 9:**
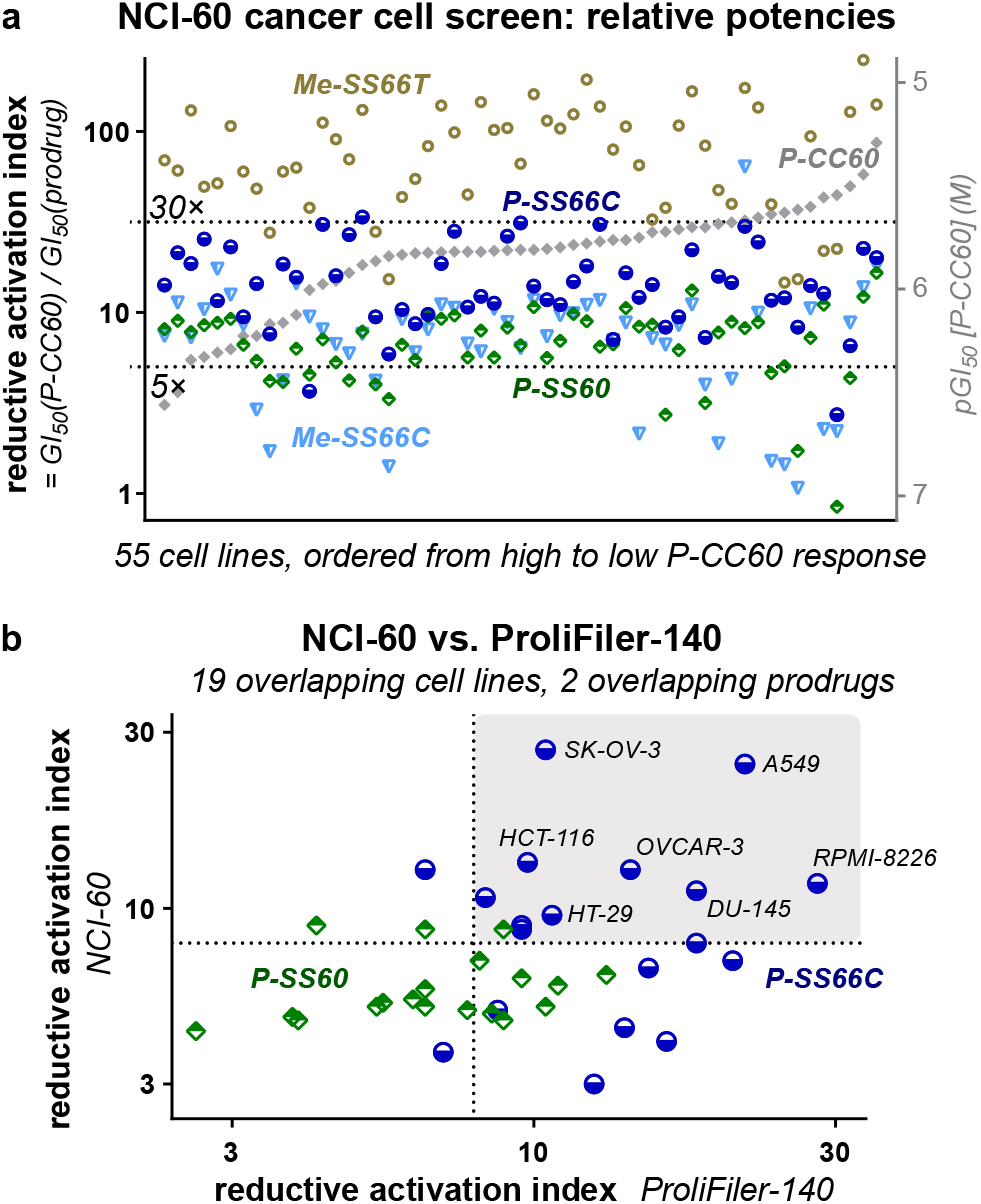
Independent screen of bioreductive activity for all redox prodrugs across the standard “NCI-60” panel. (a) Selected NCI results (full data in **Table S3**). (b) NCI-60 and ProliFiler screens give similar reductive activation indices (details in **Table S4**).

The benchmarking results of reductive index matched between NCI and ProliFiler screens: e.g. (1) variable **P-SS66C** index (average 14, maximum 30), but with nearly all **P-SS60** index in a narrow range around 6-8 (**Fig 9a**); and (2) no trends from the cell lines’ tissue of origin (**Table S3**). As expected, the *absolute* potencies of a prodrug in a given cell line differed between the screens (ProliFiler: average 4-fold higher, standard deviation 3.2-fold). Pleasingly though, despite these differences, the reductive index was well conserved between the screens (**Fig 9b**), particularly for **P-SS66C** (average 1.8-fold higher, standard deviation 1.1-fold) (**Table S4**). This suggests that the index reports reproducible aspects of reductive release, which in turn supports that the results of these screens (NCI-60: tested more prodrugs, ProliFiler: tested more cell lines) can be combined: building an unprecedented, predictive, modular overview of trigger- and cell line-dependent bioreductive performance for novel dichalcogenide bioreductive prodrugs.

Taken together, these first-ever high-throughput screens for reductase-targeting disulfide prodrugs showed outstanding performance features. Analysis referenced to the key hydrolytic control normalises out variable aspects of cell entry, non-reductive prodrug activation, and intrinsic cell line sensitivity, to show how redox SAR robustly predicts prodrug performance in these longterm assays, across 176 cell lines from many tissue types of origin. The reductive activation indices are reproducible in different labs and setups; they follow clear trends across cell lines; and above all they match the molecular understanding gained from simple cell-free and cellular assays^17^ of how the trigger structures (*trans-* or *cis-* fused, monocyclic or bicyclic) influence acute kinetic lability and reductase promiscuity. The modular tertiary carbamate design is crucial for reaching this intercomparability of results; and combines with the choice of the unmaskable, irreversibly DNA-alkylating cargo **CBI** as cargo, to ensure robust and reproducible data quality can be obtained by machine screens. This sets solid foundations for rational design or selection, and stringent validation, of reducible dichalcogenides as redox prodrug triggers in future work, even on vastly different cargos: which has been one goal of our methodological research.

The second goal of these screens was to test if any reducible triggers generally displayed the moderately activation-resisting performance that we sought, for systemically well-tolerated prodrugs, with potential for stronger tumoral bioreductive activation, towards anticancer use *in vivo*. **Me-SS66C, P-SS60** and **P-SS66C** appeared suitable for this (indices usually ca. 5-20).

It is important to clarify that although this screening identified some cancer cell lines with low nanomolar sensitivity to duocarmycin prodrugs in general (e.g. SK-MEL-28), this study did not aim to perform dubious “therapeutic proof of concept” assays by implanting *cargo-*sensitive cell lines *in vivo*. In our opinion, such experiments do not deliver useful information, since they are biased to ‘succeed’ in a way that is not replicable in uncontrolled clinical settings (cf. **section 2.9**). By focusing instead on the properties conferred by the *triggers*, we hoped to identify modular motifs that would be generally tolerated for high repeated dose administration, no matter the cargo. We therefore moved on to test these prodrugs *in vivo* in mice.

### 2.8 *In vivo* pharmacokinetics and prodrug tolerance

We first examined the *in vivo* pharmacokinetics of representative compound **SS60-CBI-AZI** in Balb/c mice after i.v. administration at 5 mg/kg (3 animals per timepoint, four timepoints from 5-90 minutes post injection). Compound plasma halflife was ca. 20 min, by HPLC-MS/MS (**Fig S12a**). Matching expectations for a low-release prodrug, no released **HO-CBI-AZI**, or activated cyclopropane, or adducts, were detected. This gave confidence that the redox prodrugs might give low systemic exposure of the activated CBI, and therefore be tolerable *in vivo*.

To test if low prodrug activation could enable *in vivo* use, we performed dosing and toxicity studies in Balb/c mice, comparing the toxicity and the tolerated dosing of low-reducible **SS60-CBI-AZI** or substantially-reducible **SeS60-CBI-AZI** and to non-reducible carbamate **O56-CBI-AZI**. Single dose toxicity was studied over the range 0.1-10 mg/kg. Dosing at ≤3 mg/kg was typically tolerated, which should be compared to the toxicity limit for fully activatable duocarmycins: typically ca. 0.1 mg/kg for *total, cumulative* dose.^45^ However, 10 mg/kg of any **AZI** carbamate was lethal in the week following treatment, though the ether control was not lethal at this dose and not even body weight variations were noted for it: suggesting that at this dose, even hydrolytic carbamate release passes the threshold for toxicity.

The potential for toxicity under low repeated dosing was then studied, comparing **SS60**- and **SeS60-** to **O56-CBI-AZI** (injections once per week over three weeks; 7 animals per condition). No adverse bodyweight losses were seen, but two toxicity features were identified. First, liver damage was indicated by statistically significant, ca. 10% increases of liver weight for the reducible probes, often with increased alanine aminotransferase levels and decreased alkaline phosphatase levels (liver damage markers); these changes were much lower for hydrolytic **O56**. Second, anemia was indicated by statistically significant decreases of typically 4-12% in hemoglobin, hematocrit, and red blood cell count, for all **CBI**s (**Fig S12b**). All other organ weights and gross pathology features were normal. Given the small statistical power of the assay, these results should be taken cautiously; but they are consistent with the liver as a site of primarily reductive activation of the monocyclic dichalcogenides.

Next, a moderate repeated dosing study in Balb/c mice (three animals per group) tested the importance of solubilising the prodrugs. For example, we have seen elsewhere that a solubilising piperazinamide **P-** sidechain near the dichalcogenide greatly increases reproducibility of cellular results^18^. Now *in vivo*, **P-SS60-CBI-TMI** was tolerated at once weekly 3 mg/kg dosing in all animals over 3 injections, without adverse effects at the end of observation 2 weeks later. By contrast, at 3 mg/kg, the still monobasic but less soluble analogues **SS60-CBI-AZI** and **SeS60-CBI-AZI** lead to significant loss of body weight (**Fig S12c**) with visible adverse effects at the injection site potentially arising from local aggregation or precipitation, since they were avoided by lower dosing at e.g. 1 mg/kg. This suggested solubilisation near the reducible trigger could indeed be beneficial for tolerability.

A larger study was run in fifty athymic nu/nu NMRI mice (ten animals per group), inoculated with BxPc3 pancreatic cancer cells since the study had been intended as a therapeutic efficacy trial (see **section 2.9**). Similarly solubilised **SS66C** derivatives were now included. In this study, the tumor growth rates in all animals were much lower than in any previous or subsequent trials (**Fig S13c**): so, we could only extract data about drug tolerability, not antitumor efficacy. Still, the study confirmed that the solubilised **P-SS60-, Me-SS66C-**, and **P-SS66C-** prodrugs of **-CBI-TMI**, and the corresponding carbamate hydrolysis control **P-CC60-CBI-TMI**, were tolerated with twice weekly injections at 3 mg/kg, over a course of five injections, in all animals: since no distinct differences of body weight or animal health were seen as compared to the vehicle control (**Fig S13a**).

Finally, another large study intended for antitumor efficacy also had to be examined only in terms of tolerability. This assay in immunocompetent Balb/c mice (ten animals per group), inoculated with 4T1 murine breast cancer cells, also treated animals with reducible **P-SS60-, Me-SS66C-, P-SS66C-CBI-TMI**, or carbamate control **P-CC60-CBI-TMI** (3 mg/kg), once or twice per week. Here, tumors grew well in all groups, but since even the technical positive control irinotecan failed to slow tumor growth (**Fig S13d**), no efficacy conclusions could be drawn. Repeated dosing at 3 mg/kg was however tolerated also in this mouse strain, again in mice bearing the additional burden of tumors (**Fig S13b**). This excellent tolerability supported our aim to develop solubilised, low-release, reducible carbamate prodrugs for *in vivo* use with duocarmycin-type cargos.

### 2.9 Dichalcogenide CBI prodrugs show anticancer efficacy in murine cancer models

We then moved to *in vivo* anticancer efficacy trials of these prodrugs. We anticipated that the *in vivo* growth environments significantly regulate the tumoral redox/metabolic biochemistries which can activate the prodrugs: so, we did not expect cell lines’ reductive index from 2D cell culture to be reproduced *in vivo*, but selected tumor models for their technical reproducibility and for their value as biologically meaningful or medically relevant models (**Fig 10a**). As one model, we chose the syngeneic murine breast cancer 4T1, implanted orthotopically (at the native tissue site) into the fat pad of immunocompetent Balb/c mice (i.e. offering a realistic immune system and tumor microenvironment). This model gives rapid, aggressive tumor growth.^51^ As a second model, we chose to xenograft human BxPC-3 pancreatic adenocarcinoma cells into immunodeficient hosts. This is a slower-growing model, with low metastasis, that can be more resistant to traditional antimitotic therapy.^52,53^ Mice were implanted, randomized once tumors reached 100-150 mm^3^ volume, and treated once or twice weekly with prodrug or control compounds typically at 3 mg/kg (8-12 animals per condition, **Fig 10a**). Tumors were measured by caliper during the assay and weighed at termination.

**Figure 10:**
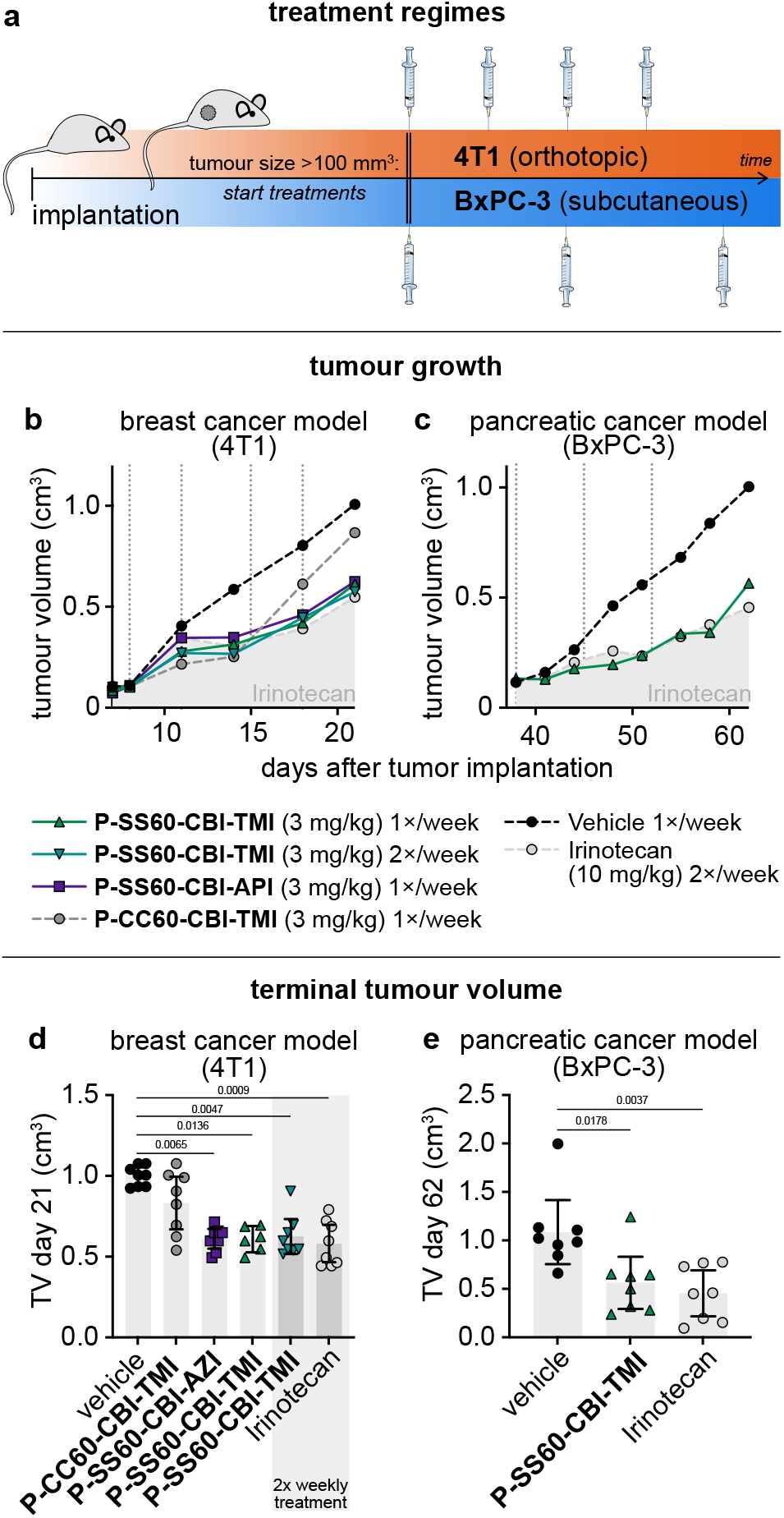
*in vivo* anticancer efficacy. (**a**) Designs for mouse anticancer efficacy assays: murine breast cancer (4T1) implanted orthotopically in the mammary fat pad of immunocompetent Balb/c mice; human pancreatic cancer (BxPC-3) implanted subcutaneously in athymic nu/nu NMRI mice. (**b**,**c**) Tumor volume (TV) over the course of the studies (median values). (**d**,**e**) TV at study termination (raw values with means; days 21 and 62 respectively; *p-*values from Kruskal-Wallis test indicated where *p* < 0.05). See also **Fig S14**.

The 4T1 efficacy study compared reducible **P-SS60-CBI-TMI** to non-reducible **P-CC60-CBI-TMI** (3 mg/kg). This time, the technical positive control irinotecan showed the expected tumor-slowing effect. Both reducible prodrugs and non-reducible control delayed tumor growth in the first week of treatment. However, only the reducible prodrugs maintained statistically significant tumor suppression until study termination (**Fig 10b,d** and **Fig S14**). Although other interpretations are possible, this is highly suggestive that reductive activation in tumors indeed delivers CBI at effective tumor-suppressing levels, that are also significantly above those provided by systemic or tumoral carbamate hydrolysis.

Finally, we tested **P-SS60-CBI-TMI** in the slower-growing pancreatic cancer model BxPC-3 in athymic NMRI mice (8 animals per group). The response to once weekly **P-SS60** treatment (3 mg/kg) was outstanding, almost identically tumor-supressive as the technical control irinotecan (10 mg/kg), with consistent ca. 70% tumor growth rate suppression over four weeks, and high statistical significance (**Fig 10c,e** and **Fig S14**). As the **P-CC60** non-reductive control had not been included, we cannot estimate how much of the therapeutically beneficial effect in this model stems from reductive or non-reductive CBI release: but at least in cell culture, BxPC-3 had reductive release six-fold higher than the non-reductive level alone (**Fig 8e**), so we expect that similarly, tumoral reductive release may be a significant factor.

After these promising studies showing efficacy and supporting the importance of reductive release, the critical question is now: is reductive CBI release higher in tumors than in healthy tissues? This cannot be answered from efficacy data alone, since acute tumoral response to CBI is different from that of healthy tissues; and exposure to released CBI cargo is also challenging to track in tissue by typical HPLC methods, due to its low (∼subnanomolar) levels that irreversibly alkylate DNA. To tackle this question and thus to estimate the potential of these dichalcogenide strategies to provide tumor-selective drug delivery, we have now begun a new *in vivo* assay program, with a different set of bioreductive prodrugs that allow sensitive tracking and quantification of release. Results will be reported in due course.

## 3. SUMMARY AND CONCLUSIONS

We have developed a novel, modular chemical space of bioreductive dichalcogenide prodrugs, for the as-yet unaddressed target space of disulfide reductases such as the thioredoxin system (**Fig 1**). The ten redox-sensitive triggers and controls allowed us to resolve contributions of reductive vs. non-reductive prodrug activation, and the 16 prodrugs (including a novel CBI azetidine) allowed us to test a modular “redox SAR”-based design hypothesis, relying on minimal and maximal activation controls (**Fig 2-4**). We have confirmed their reductive activation mechanisms (**Fig 5**) and used cellular knockout assays to quantify triggers that can reach up to ≥80% cellular selectivity for activation by the thioredoxin system (**Fig 6-7**). Two independent, automated, high-throughput cellular screens in 176 cancer cell lines confirmed the redox SAR principle of rationally tuned prodrug potency; they deliver an unprecedentedly comprehensive body of data indicating actual disulfide reductase activity, rather than reductase gene expression or overall protein levels, across this broad range of cell types (**Fig 8-9**). Several of the solubilised prodrugs were well tolerated *in vivo* over multiple trials, supporting their design principle of high metabolic and hydrolytic robustness which we believe is a prerequisite for effective tumor-selective reductive release. Tolerability is a stringent hurdle for duocarmycin prodrugs, due to their severe, cumulative toxicity; so, this success is encouraging for future applications (**Table 2**).

**Table 2.** Tolerability for repeated dose administration *in vivo*. The table summarizes dosages that are well-tolerated under weekly or twice-weekly administration, as tested in multiple study settings (PK: pharmacokinetics; MTD: maximum tolerated dose).

| *in vivo* tolerability under repeated dosing (mouse) |  |  |  |
| --- | --- | --- | --- |
| prodrug | studies | dose recommended (mg/kg) |  |
| **SS60-CBI-AZI** | PK, MTD | 0.3-1 | *adverse effects* |
| **SeS60-CBI-AZI** | MTD | 0.3-1 | *adverse effects* |
| **O56-CBI-AZI** | MTD | 1 |  |
| **P-SS60-CBI-AZI** | MTD, efficacy | 3 | *well tolerated* |
| **P-SS60-CBI-TMI** | MTD, efficacy | 3 | *well tolerated* |
| **P-SS66C-CBI-TMI** | MTD, efficacy | 3 | *well tolerated* |
| **Me-SS66C-CBI-TMI** | MTD, efficacy | 3 | *well tolerated* |
| **Me-SS66T-CBI-TMI** | MTD, efficacy | 3 | *well tolerated* |
| **P-CC60-CBI-TMI** | MTD, efficacy | 3 | *well tolerated* |

Most importantly, *all these aspects*, from synthesis to systemic robustness *in vivo*, are modular features of the dichalcogenide prodrug strategy: so, the same principles and performance can be expected from *any* cargo that is used with this (stabilised dichalcogenide plus tertiary carbamate) prodrug approach.

By combining tolerability with efficacy, this first set of CBI prodrugs also indicates that using the dichalcogenide prodrug strategy (even with the historically difficult-to-tame CBI cargo) can be a promising route for *in vivo* anticancer applications. In particular, piperazinamide **P-SS60-CBI-TMI** gave high antitumor efficacy in the relatively resistant BxPC-3 pancreatic cancer model, on the same level as a 3-fold higher dose of FDA-approved irinotecan; and it likewise gave good efficacy in the aggressive syngeneic 4T1 breast cancer model (**Fig 10**).

More broadly, we expect that while the tolerability is a general feature of the redox prodrug platform which can benefit any chemical cargo or animal model, efficacy within the tolerability window will only be reliably achieved by matching the choices of model, redox trigger, and cargo type. This multi-variable problem requires a much deeper knowledge level around disulfide manifold bioreduction than currently available. However, by separating the features of prodrug performance that are based on redox reactivity of the trigger, from those that are based on trigger hydrolysis, as well as separating the model-dependent and cargo-dependent contributions to results, the systematic modular approach we present makes significant advances towards reaching this level.

The ideal goal for redox prodrugs is to develop a platform approach that can maximise the ratio of drug exposure in tumoral vs. in healthy tissues, rather than relying only on differences of their intrinsic sensitivities to a given cargo. Testing this exposure ratio, with directly quantifiable redox reporter prodrugs based around another cargo than the CBIs, is our aim in ongoing work.

Quantifying exposure with reporters, and developing increasingly effective prodrug-based therapeutics, are mutually reinforcing advances for testing the potential of bioreductive prodrugs. We believe that both will be required, over multiple interleaved cycles of refinement, in order to face this multi-variable problem with a quantitative, SAR-based understanding. We anticipate that the systematic body of predictive data in this study, which complements previous *in vitro* development steps,^16–18^ should prove vital to enable and orient such *in vivo* followup cycles; and we believe that direct reporter methods will at last start to reveal the selectivity that engineered, synthetic dichalcogenide redox substrates can deliver by harnessing oxidoreductase-based release in the disulfide/dithiol manifold.

That challenge should not be underestimated: bioreductive release prodrug systems based on oxidised nitrogen species have taken several *decades* to reach their current, and still incomplete, level of predictive or SAR-based understanding.^3,31^ However, we believe that by emphasising a high volume of comparative SAR-based data, this work will help quantitative *in vivo* investigations to succeed rather more rapidly.

A reliable, actionable understanding of the disease indications in which redox dysregulation can be exploited, and to what degree it may provide selective therapeutic benefits, would resolve several decades of tantalising observations and theoretical deadlock. These dichalcogenides are bringing a new biochemical target space into play: time will tell if they can be used as straightforwardly as in this study, modularly retrofitting existing cargos to turn them into powerful bioreductive diagnostics and prodrugs.

## AUTHOR INFORMATION

### Funding

This research was supported by funds from the German Research Foundation (DFG: SFB 1032 project B09 number 201269156, SFB TRR 152 project P24 number 239283807, SPP 1926 project number 426018126, and Emmy Noether grant 400324123 to O.T.-S.); LMUExcellent (Junior Researcher Fund to O.T.-S.); the Munich Centre for NanoScience initiative (CeNS to O.T.-S).; and Federal Ministry of Education and Research GO-Bio 161B0632 to O.T.-S. and J.T.-S..

### Notes

J.G.F., L.Z., J.T.-S. and O.T.-S. are inventors on patent applications PCT/EP2022/057483 and PCT/EP2022/059280 filed by the Ludwig-Maximilians-University (LMU) Munich in 2021, covering compound structures reported in this paper. The authors declare no competing financial interests.

Where quantities are averaged over non-comparable conditions, the geometric mean (logarithmic) is used. For example, IC_50_ values 0.25 nM and 1 nM in two cell lines are averaged to 0.5 nM to give a single indicative value; *idem* for mean reductive index values over e.g. 55 NCI cell lines.

## Author Contributions

J.G.F. performed synthesis, analysis, cell-free prodrug activation and cellular assays; coordinated screening, analysed screening data, performed data assembly. A.K. performed cell-free prodrug activation, cellular assays, coordinated screening and *in vivo* studies, and performed data analysis. L.Z. and M.S.M. performed synthesis and analysis. C.H. performed cell-free prodrug activation and cellular assays. J.T.-S. designed, coordinated, and analysed *in vivo* studies, and performed data assembly. O.T.-S. designed the concept and experiments, supervised experiments, performed screening data analysis, and coordinated data assembly. J.G.F. and O.T.-S. wrote the manuscript with input from all co-authors.

## Acknowledgements

J.G.F. thanks the Studienstiftung des deutschen Volkes for support through a PhD scholarship; L.Z. thanks the Fonds der Chemischen Industrie (FCI) for support through a PhD scholarship; J.T.-S. thanks the Joachim Herz Foundation for fellowship support. We thank the NCI DTP for NCI60 screening; Ramon Messeguer at Leitat for initial *in vivo* proof of concept studies; Diana Behrens and team at EPO Berlin for followup *in vivo* studies; Anna Kondratiuk, Yuliia Holota, Sergey Zozulya and the other team members at Bienta (Kyiv, Ukraine) for *in vivo* PK and initial MTD studies; Bettina Stahnke and Philipp Metzger and team at Reaction Biology for further *in vivo* studies.

