## Supporting Information for "Cyclic dichalcogenides extend the reach of bioreductive prodrugs to harness the thioredoxin system: applications to *seco*-duocarmycins"

##### Author Contributions

J.G.F. performed synthesis, analysis, cell-free prodrug activation and cellular assays; coordinated screening, analysed screening data, performed data assembly, and wrote the manuscript. A.K. performed cell-free prodrug activation, cellular assays, coordinated screening and *in vivo* studies, and performed data analysis. L.Z. and M.S.M. performed synthesis and analysis. C.H. performed cell-free prodrug activation and cellular assays. J.T.-S. designed, coordinated, and analysed *in vivo* studies, and performed data assembly. O.T.-S. designed the concept and experiments, supervised experiments, performed screening data analysis, coordinated data assembly, and wrote the manuscript, with input from all co-authors.

#### Table of Contents

**1. Previous chemical tuning of dichalcogenide-based units and prodrug design S2**

**2. CBI-based redox prodrug assembly overview ........ S3**

**3. Reduction-mediated prodrug release monitored by HPLC ........ S6**

**4. Cellular proliferation assays ........ S9**

**5. ProliFiler-140 cancer cell screening: cellular cytotoxicity ($IC_{50}$ values) ........ S13**

**6. NCI-60 cancer cell screening: growth inhibition ($GI_{50}$ values) ........ S22**

**7. *in vivo* studies ........ S24**

7.1. in vivo PK and prodrug tolerance (mouse studies) ........ S24

7.2. in vivo efficacy mouse studies ........ S26

**8. General methods ........ S27**

8.1. Cell-free HPLC protocols ........ S27

8.2. Cell culture methods ........ S27

8.3. Methods for in vivo animal studies ........ S28

8.4. Synthetic techniques ........ S29

**9. Chemical synthesis ........ S31**

9.1. General synthetic protocols ........ S31

9.2. Precursor syntheses ........ S34

9.3. CBI-AZI prodrug series ........ S42

9.4. CBI-TMI prodrug series ........ S47

**10. NMR Spectra ........ S51**

**11. Supporting References ........ S86**

#### 1. Previous chemical tuning of dichalcogenide-based units and prodrug design

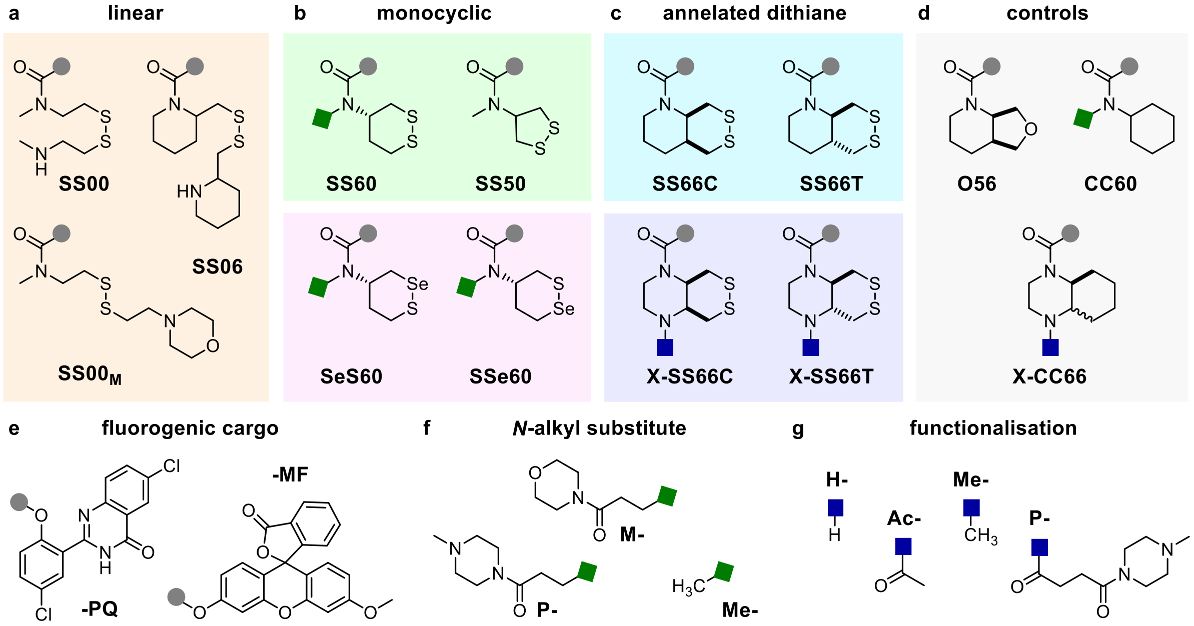


**Figure S1: Summary of previous studies.** Overview of redox triggers, molecular designs and fluorescent cargos to create fluorogenic probe systems.

(**a**) Linear disulfide-based redox trigger benchmark the basal thiol-mediated reduction in a system: The simple freely-rotatable **SS00**, the preorganised piperidine-derivative **SS06** and the asymmetric morpholino **SS00$_{\mathbf{M}}$**.[1]

(**b**) Monocyclic dichalogenides show the impact of ring-strain and redox potential and the impact of a heterogenous S-Se bond at either of both positions. These studies revealed that 1,2-dithiolane **SS50** is labile under any thiol-mediated reduction and sensitive to ring-opening polymerisation, while 1,2-dithiane **SS60** is highly reduction resistant and activated only by strong bioreductants as e.g. the redox effector protein Thioredoxin.[2] 1,2-thiaselenanes **SeS60** and **SSe60** are highly reactive with selenol nucleophiles (Sec) as e.g. found in the active site of the selenoenzyme TrxR with kinetic reversibility being the key factor to differentiate both regioisomers.[3]

(**c**) Annellating 1,2-dithianes to create more complex ring structures of the **SS66**-type, with the trans-fused **SS66T**, against expectations, being thermodynamically less stable and labile to monothiol reduction as compared to the cis-fused **SS66C** being the most reduction-resistant motif of the whole series.[1] Fused **SS66**-type piperazine triggers instead gave **X-SS66C** and **X-SS66T** with similar selectivity profiles, but highly improved release kinetics optional diverse functionalisation.[4]

(**d**) Control motifs show the robustness of the molecular designs: **O56** to control for unspecific degradation or hydrolysis of the **SS66**-type annelated dithianes; **CC60** to control for **SS60**, **SeS60** and **SSe60** with optional *N*-alkyl substitution; and the annelated cyclohexane motif **X-CC66** to control for **X-SS66**-type annelated dithianes with optional functionalisation attached.

(**e**) Fluorescence of the molecular cargos **PQ-OH** (hydroxyphenylquinazolinones) and **MF-OH** (3-*O*-methyl-fluorescein) was masked by freely combining trigger and cargo by mono-*O*-carbamoylation creating fluorogenic probes. Probes are activated by reduction of the dichalcogenide leading to 5-exo-trig cyclisation, that expels the cargo and re-establishes the molecular features key to its fluorescence properties.[1]

(**f**) Several different monocyclic trigger designs have been improved by N-alkyl substitution with either *N*-methylpiperazinamide (**P**) or morpholinamide (**M**) instead of the simple methyl (**Me**)-derivatives.[3]

(**g**) Annelated 1,2-dithianes of the **X-SS66C**-type were optionally substituted by methyl (**Me**) instead of primary amines (**H**), acetyl (**Ac**) to differentiate *N*-alkylated vs. *N*-acylated versions and *N*-methylpiperazinamide (**P**) attached by acylation. Functionalisation did not affect the redox resistance and enzyme-selectivity profiles.[4]

#### 2. CBI-based redox prodrug assembly overview

**a trigger amine precursor**

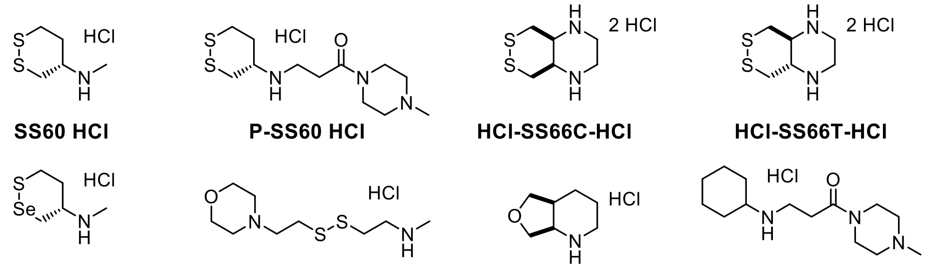

**SS60 HCl** **P-SS60 HCl** **HCl-SS66C-HCl** **HCl-SS66T-HCl**

**SeS60 HCl** **SS00$_M$ HCl** **O56 HCl** **P-CC60 HCl**

**b prodrug assembly**

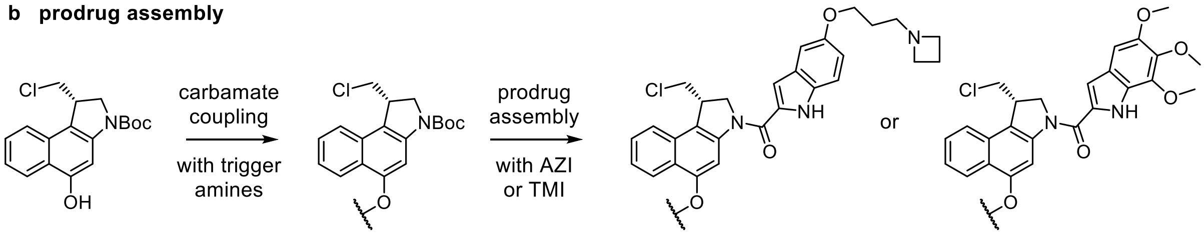


**HO-CBI-Boc** **xx-CBI-Boc** **xx-CBI-AZI** **xx-CBI-TMI**

| *synthetic scales and yields* | carbamate coupling | *additional derivatisation step* | amide coupling (AZI) | amide coupling (TMI) |
|---|---|---|---|---|
| **SS60** | 123 mg (96%) | | 44 mg (49%) | *not made* |
| **P-SS60** | 191 mg (58%) | | 17 mg (65%) | 42 mg (82%) |
| **H-SS66C** (*precursors*) | 97 mg (67%) | → **Me-SS66C** | 2 mg (8%) | 16 mg (44%) |
| | | → **P-SS66C** | *not made* | 11 mg (65%) |
| **H-SS66T** (*precursors*) | 42 mg (35%) | → **Me-SS66T** | 2 mg (14%) | 19 mg (78%) |
| **SeS60** | 85 mg (65%) | | 27 mg (51%) | *not made* |
| **SS00$_M$** | 206 mg (99%) | | 30 mg (21%) | 9 mg (37%) |
| **MC** | 72 mg (99%) | | 21 mg (75%) | 8 mg (15%) |
| **O56** | 116 mg (95%) | | 50 mg (20%) | *not made* |
| **P-CC60** | 72 mg (63%) | | *not made* | 29 mg (78%) |
| **OBn** | commercial | | 90 mg (57%) | *not made* |

**Figure S2:** (**a**) Trigger amine precursor used in this study. *For synthetic procedures see 8.1. General synthetic protocols.* (**b**) Prodrug assembly: synthetic steps and isolated yields. Carbamate coupling of **HO-CBI-Boc** was accomplished using *in situ* amine-activated triphosgene *via* chloroformate **Cl(CO)O-CBI-Boc** that was quenched with the various trigger amine HCl salts to give **xx-CBI-Boc** intermediates in good to excellent yields (35-99%). *See General Procedure A and synthetic procedures for further details.* Prodrug assembly of **xx-CBI-Boc** intermediates with **AZI-OH** or **TMI-OH** was accomplished either using EDCI (General Procedure B, *Method D*) or else **TMI-OH** pre-activated with oxalyl chloride/DMF (**General Procedure B,** *Method E*). The final 16 prodrugs and control compounds were prepared in mostly good yields (8-78%).

CBI-AZI series

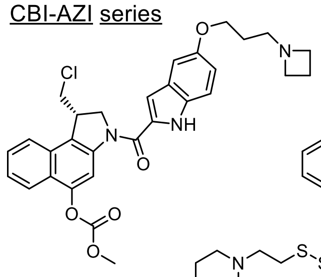

**MC-CBI-AZI**

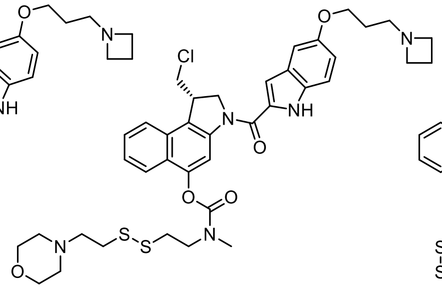

**SS00$_M$-CBI-AZI**

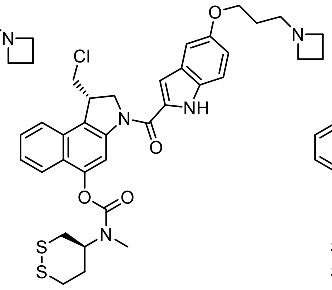

**SS60-CBI-AZI**

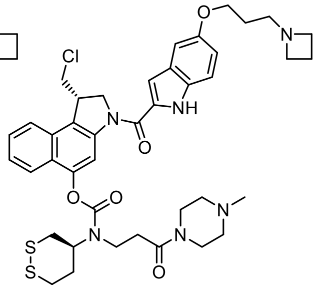

**P-SS60-CBI-AZI**

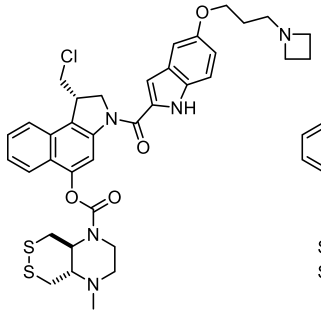

**Me-SS66T-CBI-AZI**

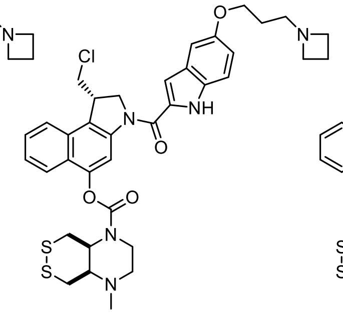

**Me-SS66C-CBI-AZI**

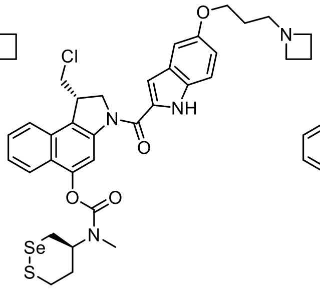

**SeS60-CBI-AZI**

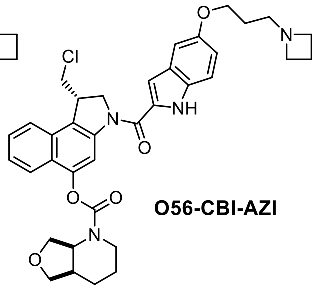

**O56-CBI-AZI**

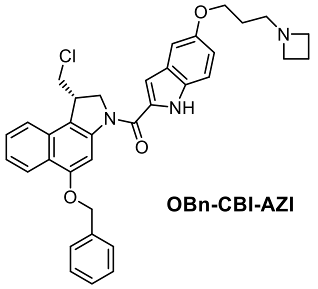

**OBn-CBI-AZI**

CBI-TMI series

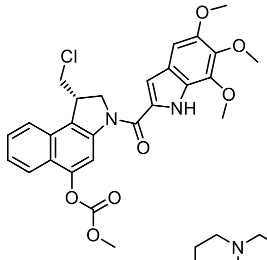

**MC-CBI-TMI**

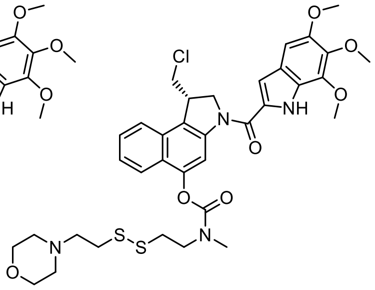

**SS00$_M$-CBI-TMI**

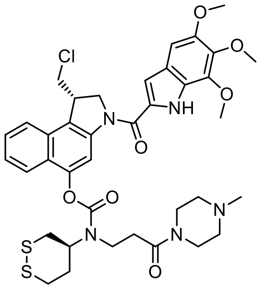

**P-SS60-CBI-TMI**

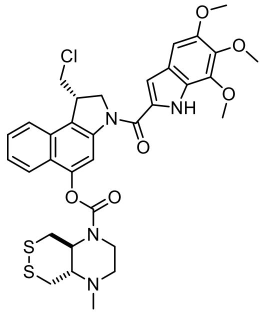

**Me-SS66T-CBI-TMI**

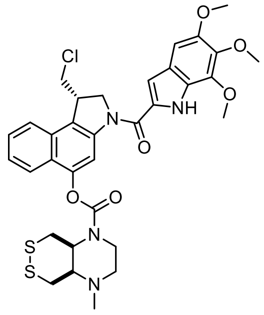

**Me-SS66C-CBI-TMI**

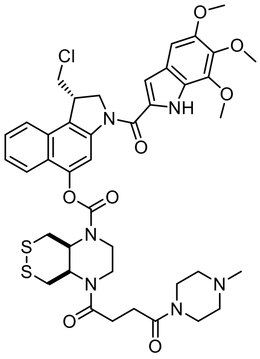

**P-SS66C-CBI-TMI**

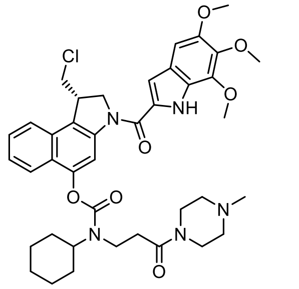

**P-CC60-CBI-TMI**

**Figure S3: Prodrug overview**. The CBI-AZI series consists of the monocyclic 1,2-dithiane-based redox prodrugs **SS60-CBI-AZI** and **P-SS60-CBI-AZI**, the 1,2-thiaselenane-based redox prodrug **SeS60-CBI-AZI**, the two annellated 1,2-dithiane-based prodrugs **Me-SS66C-CBI-AZI** and **Me-SS66T-CBI-AZI** controlled by methylcarbonate **MC-CBI-AZI**, linear disulfide reduction control **SS00$_M$-CBI-AZI**, and non-reducible stable carbamate **O56-CBI-AZI** and benzylether **OBn-CBI-AZI**. The CBI-TMI series contains the monocyclic 1,2-dithiane-based redox prodrug **P-SS60-CBI-TMI,** and the three annellated dithiane-based prodrugs **Me-SS66C-CBI-TMI** and **Me-SS66T-CBI-TMI** and **P-SS66C-CBI-TMI** controlled by methylcarbonate **MC-CBI-AZI**, linear disulfide reduction control **SS00$_M$-CBI-TMI** and the non-reducible **P-CC60-CBI-TMI**.

Example: **Full Synthesis of P-SS60-CBI-TMI**

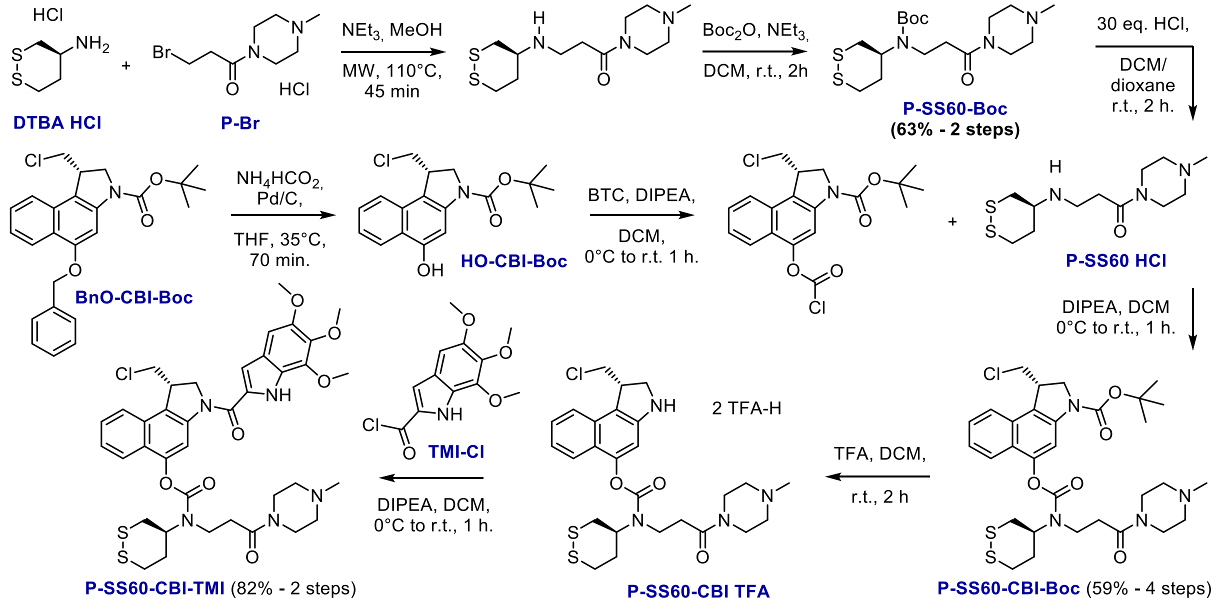


**Step 1**: To a solution of the **DTBA-HCl** (335 mg, 1.95 mmol, 1.3 eq.) in MeOH (0.3 M) was added **P-Br** (407 mg, 1.50 mmol, 1.0 eq) and $NEt_3$ (281 µL, 2.03 mmol, 1.35 eq.). The clear solution was heated in a sealed tube using a laboratory microwave (110 °C, 25 W, 45 min), was then cooled to r.t. and concentrated under reduced pressure.

**Step 2**: The material obtained in step 1 was suspended in DCM (0.1 M) and charged with $NEt_3$ (520 µL, 3.75 mmol, 2.5 eq.) and $Boc_2O$ (655 mg, 3.00 mmol, 2.0 eq.) at r.t., and the mixture was stirred at for 2h. The reaction was then concentrated under reduced pressure. Purification by flash column chromatography yielded **P-SS60-Boc** as a crystalline, colourless solid (367 mg, 0.94 mmol, 63%).

**Step 3**: **P-SS60-Boc** (234 mg, 0.60 mmol) was dissolved in DCM (0.1 M), and HCl (3.0 ml of 4 M in dioxane, 20 equiv.) was added at r.t.. The reaction mixture was stirred at r.t. for 2h, precipitation was observed and all volatiles were removed under reduced pressure to obtain **P-SS60-HCl** as a colourless solid.

**Step 4**: **OBn-CBI-Boc** (424 mg, 1.00 mmol, 1.0 eq.) and dissolved in THF (60 mL, 0.02 M). The mixture was heated to 35 °C and Pd/C (118 mg, 10% on charcoal, 1.0 eq.) and $NH_4HCO_2$ (1.0 mL of 4 M aq. Solution, 4.0 eq.) were added. The mixture was stirred for 70 min at 35 °C until quantitative turnover was confirmed by TLC, was then filtered through Celite, the filter cake was washed with ethyl acetate (40 mL), and the combined filtrates were concentrated under reduced pressure to obtain **HO-CBI-Boc** as a colourless solid.

**Step 5**: The material obtained in step 4 was dissolved in anhydrous DCM (0.016 M) and the resulting solution was cooled to 0 °C. Triphosgene (350 mg, 1.18 mmol, 1.18 eq., 0.05 M solution in anhydrous DCM) and DIPEA (212 µL, 1.2 mmol 1.2 eq, 0.05 M solution in anhydrous DCM) were added. The resulting mixture was stirred at 0 °C for 15 min, was then allowed to warm to r.t. and was further stirred for 1 h. All volatiles were removed using an external liquid nitrogen trap to afford the corresponding chloroformate derivative as a colourless solid.

**Step 6**: The material obtained in step 3 containing **P-SS60-HCl** was suspended in anhydrous DCM (0.05 M) and DIPEA (239 µL, 1.35 mmol, 2.7 eq) was added to yield a colourless solution. To the mixture was added dropwise at 0 °C a solution of the material obtained in step 5 (0.05 M in anhydrous DCM). The resulting mixture was further stirred at 0°C for 15 min, was then allowed to warm to r.t. and was further stirred for 1 h before being concentrated under reduced pressure. Purification by flash column chromatography yielded **P-SS60-CBI-Boc** as a crystalline, colourless solid (191 mg, 0.29 mmol, 59%).

**Step 7**: **P-SS60-CBI-Boc** (43.0 mg, 0.066 mmol, 1.0 eq.) was dissolved in anhydrous DCM (0.1 M), trifluoroacetic acid (230 µL, 3.91 mmol, 60.0 eq.) was added and the resulting mixture stirred at r.t. for 2 h. All volatiles were removed under reduced pressure to yield **P-SS60-CBI-TFA** as a colourless solid.

**Step 8**: The material obtained in step 7 was dissolved in anhydrous DCM (0.05 M), cooled to 0 °C and DIPEA (90 µL, 0.53 mmol 8.0 eq.) was added to give a clear solution. Then, a solution of **TMI-Cl** (pre-activated from **TMI-**0.2 M in DCM, 6.0 eq.) was added sequentially until full conversion of the aniline confirmed. The mixture was diluted with DCM, washed with water (2×) and the aq. layer was extracted with DCM (2×). The combined organic layers were dried over $Na_2SO_4$ and concentrated to give a coloured solid. Purification was achieved by FCC (DCM/MeOH) to give **P-SS60-CBI-TMI** (42.0 mg, 0.054 mmol, 82%) as a colourless solid.

#### 3. Reduction-mediated prodrug release monitored by HPLC

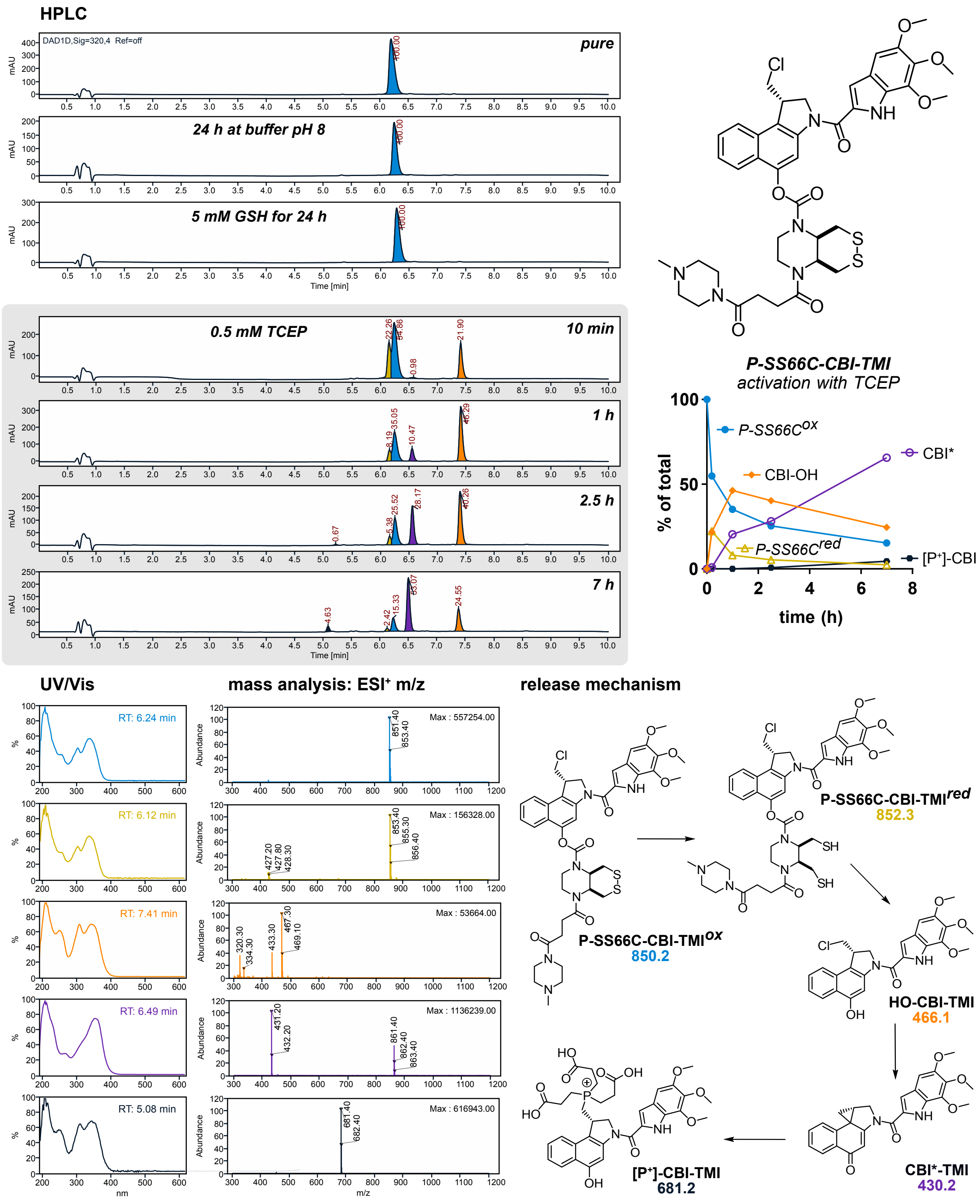


**Figure S4: Reduction mediated prodrug activation - HPLC-based reduction assay**. Phosphine-reductant-mediated cargo release of **P-SS66C-CBITMI$^{ox}$** monitored in cell-free buffer over 7 h showing: Reduction to **P-SS66C-CBI-TMI$^{red}$**, then intramolecular cargo expulsion giving **HO-CBI-TMI**, then formation of the biologically active species **CBI*-TMI**. With an excess of phosphine-based reduction-agent tris(2-carboxyethyl)-phosphine (TCEP) the corresponding adduct **[P$^+$]-CBI-TMI** was detected. *Compare the very slow kinetics of phosphine-reduction of this particular prodrug, to the much faster kinetics of phosphine-reduction of e.g.* **P-SS60-CBI-AZI** *(see next page).*

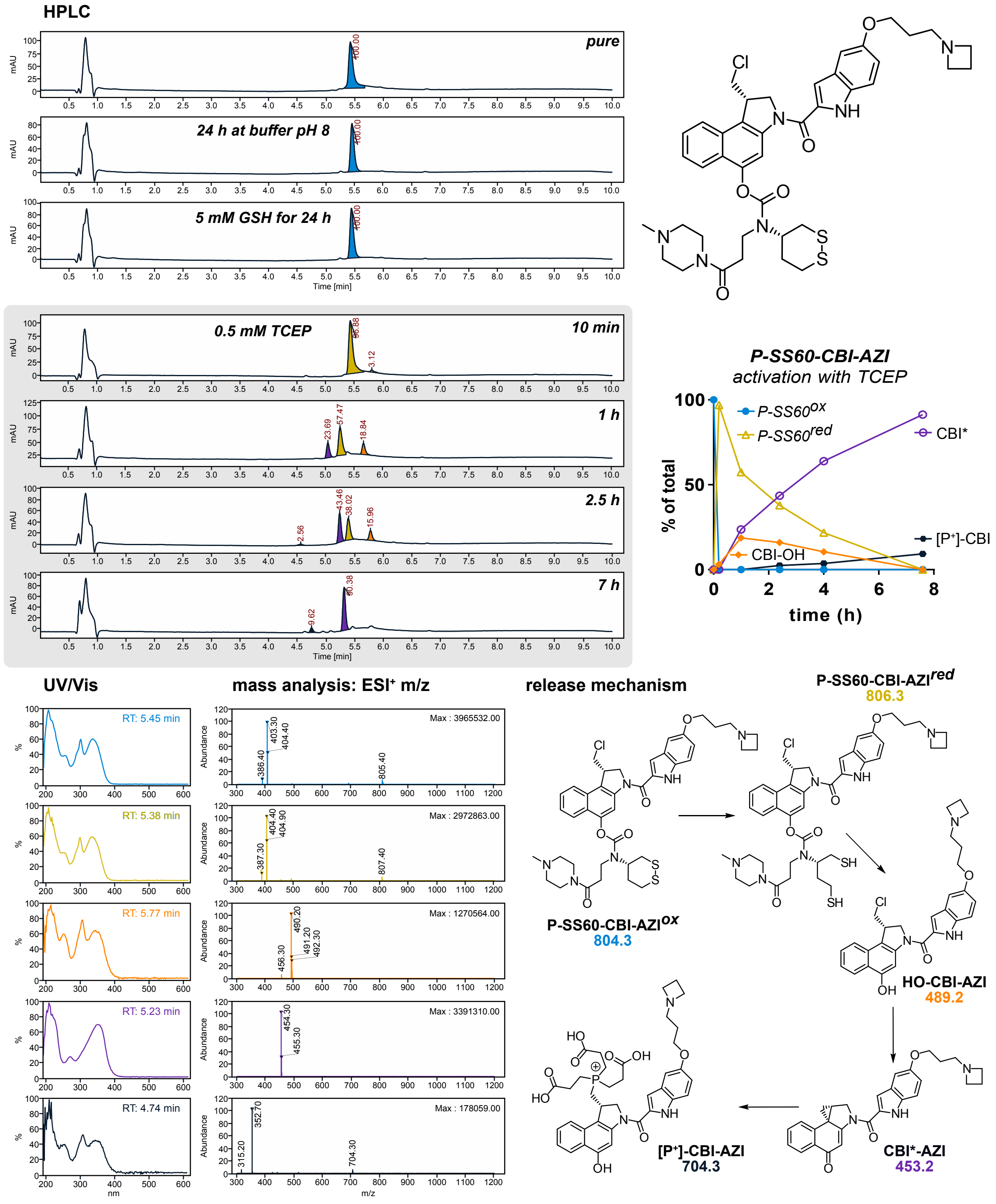


**Figure S5: Reduction mediated prodrug activation - HPLC-based reduction assay**. Phosphine-reductant-mediated cargo release of **P-SS60-CBI-AZI$^{ox}$** has been monitored at over 7 h showing: Reduction to **P-SS60-CBI-AZI$^{red}$** then intramolecular cargo expulsion giving **HO-CBI-AZI**, then formation of the biologically active species **CBI*-AZI**. Again, the phosphine adduct with TCEP **[P$^+$]-CBI-AZI** was detected. *Compare the relatively slow cargo expulsion rate for this prodrug, to the fast rates for annelated bicyclic Me-SS66C/T prodrugs (see next page).*

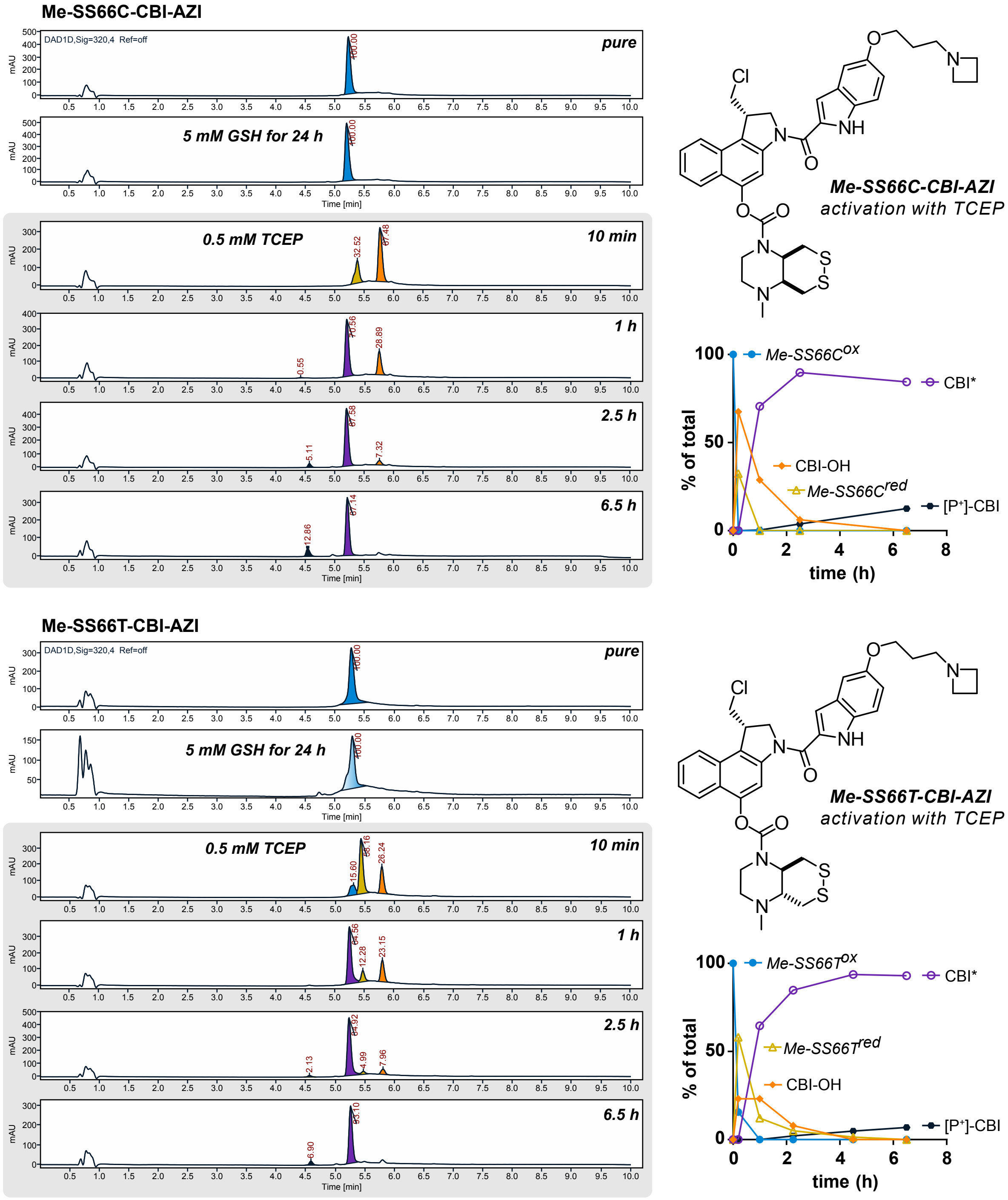


**Figure S6: Reduction mediated prodrug activation - HPLC-based reduction assay**. Reduction-mediated cargo release of **P-SS60-CBI-AZI^ox** has been monitored at over 7 h showing: Phosphine-reduction to **P-SS60-CBI-AZI^red** occurs rapidly and to high proportions, followed by rapid intramolecular cargo expulsion (**HO-CBI-AZI**) and Winstein activation (**CBI*-AZI**). *Note that the reduction and cyclisation rates are both improved compared to the examples on the previous pages. For* **Me-SS66C-CBI-AZI** *no disulfide reduction/prodrug release was indicated after challenge with GSH (5 mM) for 24 h (single UV-peak with corresponding m/z), whereas partial reduction of* **Me-SS66T-CBI-AZI** *by GSH under the same conditions was indicated (two overlaying UV-peaks with the corresponding m/z 706.2 found for the intact prodrug and m/z 454.2 found for CBI*)*

### 4. Cellular proliferation assays

#### a cellular potencies in Hela-HRE vs. A549

| cellular response | HeLa-HRE | | | A549 | | | $IC_{50}(H^*)/IC_{50}(A^*)$ | $log_{10}[IC_{50}(H^*)/IC_{50}(A^*)]$ |
|---|---|---|---|---|---|---|---|---|
| | *mean* | *SD* | *n* | *mean* | *SD* | *n* | | |
| MC-CBI-AZI | 8.09E-10 | 1.4E-10 | 3 | 1.13E-09 | 1.2E-09 | 3 (1) | 0.7 | -0.145 |
| SS00$_M$-CBI-AZI | 6.85E-09 | 5.6E-09 | 4 | 4.12E-09 | 1.6E-10 | 5 | 1.7 | 0.427 |
| SeS60-CBI-AZI | 4.89E-08 | 2.9E-08 | 3 | 6.02E-08 | 6.5E-08 | 4 | 0.8 | -0.090 |
| Me-SS66T-CBI-AZI | 4.88E-08 | 1.2E-08 | 3 (1) | 5.06E-08 | 1.3E-08 | 3 | 1.0 | -0.016 |
| Me-SS66C-CBI-AZI | 3.95E-07 | 2.0E-07 | 2 (1) | 1.23E-08 | 3.5E-09 | 3 | 32.2 | 1.508 |
| SS60-CBI-AZI | 9.56E-08 | 5.0E-08 | 5 | 1.20E-07 | 1.1E-07 | 4 | 0.8 | -0.097 |
| P-SS60-CBI-AZI | 8.88E-07 | 5.1E-07 | 3 | 1.04E-07 | 2.1E-08 | 2 (1) | 8.6 | 0.933 |
| O56-CBI-AZI | 4.62E-07 | 5.5E-07 | 3 | 6.02E-08 | 6.5E-08 | 3 | 1.1 | |
| OBn-CBI-AZI | 7.45E-07 | 1.7E-07 | 2 (1) | 4.16E-07 | 8.6E-08 | 4 | 1.8 | 0.253 |
| MC-CBI-TMI | 4.32E-09 | 1.8E-09 | 3 | 2.89E-09 | 1.5E-09 | 4 | 1.5 | 0.174 |
| SS00$_M$-CBI-TMI | 1.25E-07 | 1.4E-08 | 3 | 3.51E-08 | 1.7E-08 | 4 | 3.6 | 0.551 |
| Me-SS66T-CBI-TMI | 5.89E-08 | 1.6E-08 | 3 | 5.92E-09 | 2.6E-09 | 3 | 10.0 | 0.998 |
| Me-SS66C-CBI-TMI | 2.19E-07 | 5.7E-08 | 3 | 1.72E-08 | 1.0E-08 | 3 | 12.7 | 1.105 |
| P-SS60-CBI-TMI | 4.19E-07 | 1.3E-07 | 5 | 1.84E-07 | 1.2E-08 | 3 | 2.3 | 0.357 |
| P-SS66C-CBI-TMI | 1.99E-07 | 7.0E-08 | 4 | 1.17E-08 | 6.9E-09 | 3 | 17.0 | 1.230 |
| P-CC60-CBI-TMI | 1.35E-06 | 3.9E-07 | 3 (2) | ~1.6E-06 | / | 3 | 0.8 | -0.075 |

#### b Trx/TrxR-dependent activity: MEF TrxR1 ko vs. wt

| MEF dose response | ko | | | wt | | | $IC_{50}(ko)/IC_{50}(wt)$ | $log_{10}[IC_{50}(ko)/IC_{50}(wt)]$ |
|---|---|---|---|---|---|---|---|---|
| | *mean* | *SD* | *n* | *mean* | *SD* | *n* | | |
| SeS60-CBI-API | 1.73E-07 | 2.5E-08 | 3 | 1.26E-07 | 1.3E-08 | 3 | 1.4 | 0.137 |
| Me-SS66T-CBI-API | 2.82E-07 | 5.2E-08 | 2 (1) | 1.05E-07 | 4.2E-08 | 3 | 2.7 | 0.429 |
| Me-SS66C-CBI-API | 1.36E-06 | 8.6E-07 | 2 (2) | 3.86E-07 | 1.6E-07 | 4 | 3.5 | 0.547 |
| SS60-CBI-API | 3.29E-07 | 4.6E-08 | 3 | 4.09E-07 | 7.2E-08 | 3 | 0.8 | -0.094 |
| P-SS60-CBI-API | >1E-06 | | 4 | >1E-06 | | 4 | *1* | *0* |
| O56-CBI-API | >1E-06 | | 2 (1) | >1E-06 | | 2 (1) | *1* | *0* |
| OBn-CBI-API | >1E-06 | | 2 | >1E-06 | | 2 | *1* | *0* |
| MC-CBI-TMI | 6.02E-09 | 1.6E-10 | 3 | 6.10E-09 | 1.1E-09 | 3 | 1.0 | -0.006 |
| SS00$_M$-CBI-TMI | 1.23E-07 | 4.1E-08 | 3 | 1.69E-07 | 4.8E-08 | 2 | 0.7 | -0.137 |
| Me-SS66T-CBI-TMI | 6.42E-08 | 5.2E-09 | 3 | 2.58E-08 | 3.1E-09 | 3 | 2.5 | 0.396 |
| Me-SS66C-CBI-TMI | 3.00E-07 | 2.2E-08 | 3 | 6.20E-08 | 9.3E-09 | 3 | 4.8 | 0.684 |
| P-SS60-CBI-TMI | 1.92E-07 | 1.9E-08 | 3 | 1.30E-07 | 5.4E-08 | 3 | 1.5 | 0.171 |
| P-SS66C-CBI-TMI | 2.44E-07 | 2.4E-08 | 6 | 9.30E-08 | 7.0E-09 | 6 | 2.6 | 0.419 |
| P-CC60-CBI-TMI | 9.11E-07 | 7.3E-08 | 3 | 1.13E-06 | 2.6E-07 | 3 | 0.8 | -0.095 |

**Table S1: Summary of cellular resazurin-based cytotoxicity assays.** (**a**) Tabulated $IC_{50}$ (SD, n) values calculated from independent assays. HeLa-HRE (4 h resazurin after 48 h treatment) and A549 (4 h resazurin after 72 h treatment). Potency was compared between A549 and HeLa-HRE by fold difference: $IC_{50}$(A549)/$IC_{50}$(HeLa-HRE) and $log_{10}$[$IC_{50}$(A549)/$IC_{50}$(HeLa-HRE)]. *Compounds with a >10× fold-difference are highlighted in yellow.* (**b**) Tabulated $IC_{50}$ (SD, n) values calculated from independent assays using MEF cells (fl/fl vs. -/-) (4 h resazurin after 72 h treatment). Potency was compared between fl/fl (wild type - wt) and -/- (knockout - ko) by fold difference: $IC_{50}$(ko)/$IC_{50}$(wt) and $log_{10}$[$IC_{50}$(ko)/$IC_{50}$(wt)]. *Compounds with a >2.5× fold-difference are highlighted in yellow.*

**cellular potency I: HeLa-HRE cells (48 h treatment)**

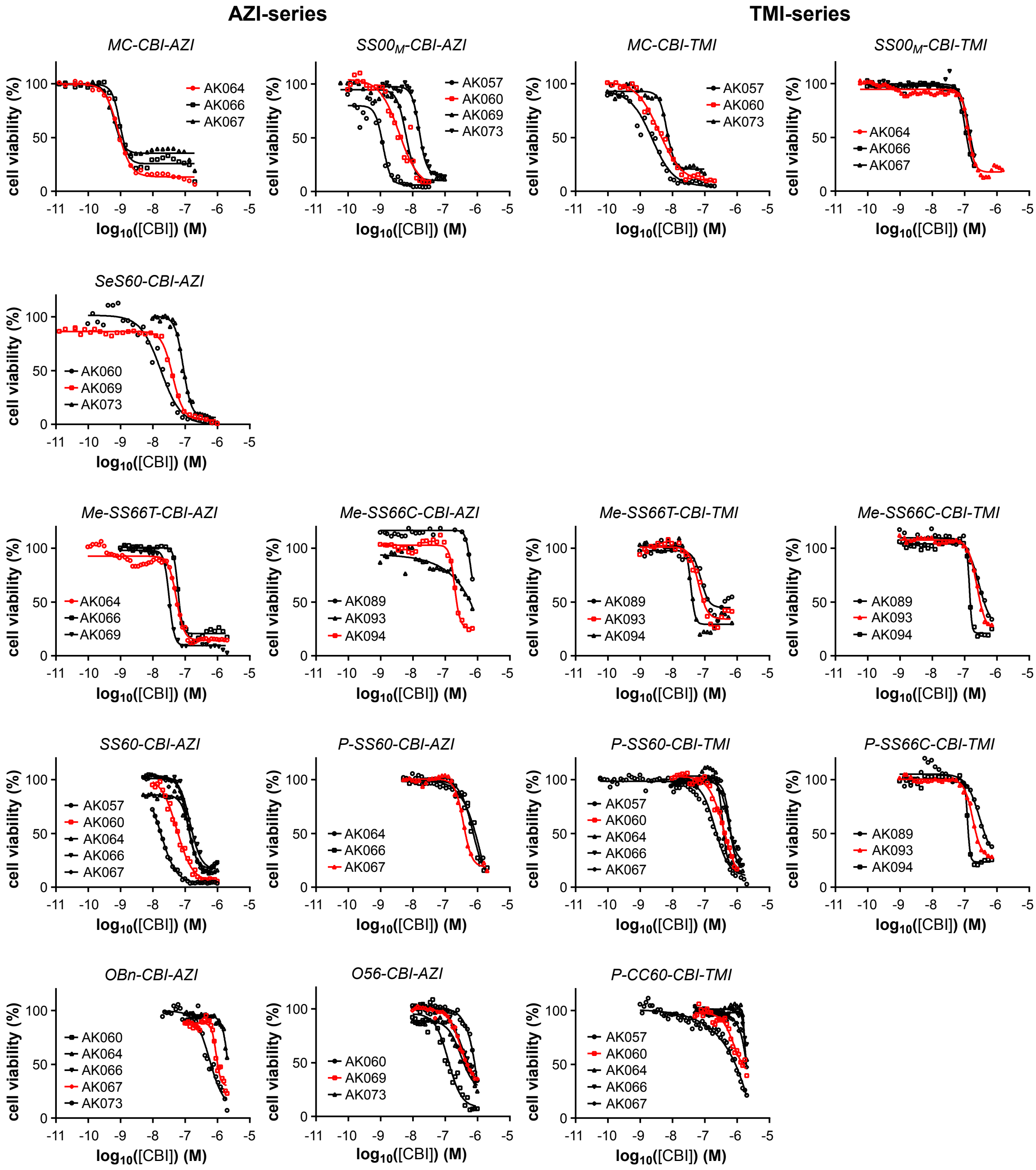


**Figure S7: Cellular resazurin-based cytotoxicity assay.** HeLa-HRE cells were treated with prodrugs of the **CBI-TMI** and **CBI-AZI** series for 48 h at concentrations from 10 pm to 30 µM depending on expected potencies of the constructs. Dose-response curves were calculated from at least 27 incremental single concentrations over a range of at least three log units. Individual experiments with annotated experiment numbers are shown for each prodrug with the representative example shown in **Figure 6** highlighted in red. $IC_{50}$ values are tabulated and correlated to dose response in A549 cells in **Table S1**.

**cellular potency II: A549 cells (72 h treatment)**

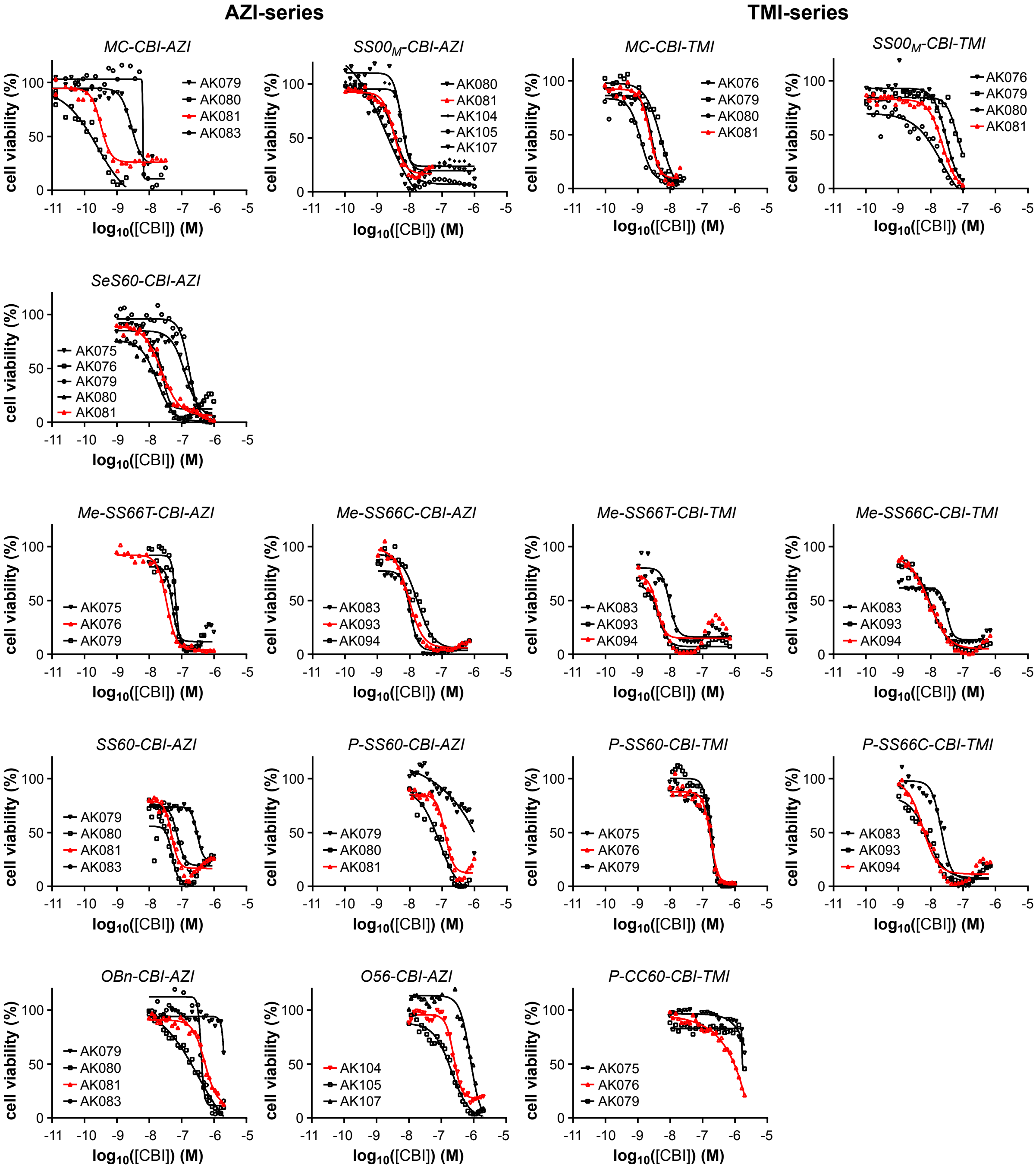


**Figure S8: Cellular resazurin-based cytotoxicity assay.** A549 cells were treated with prodrugs of the **CBI-TMI** and **CBI-AZI** series for 72 h at concentrations from 10 pm to 30 µM depending on expected potencies of the constructs. Dose-response curves were calculated from at least 27 incremental single concentrations over a range of at least three log units. Individual experiments with annotated experiment numbers are shown for each prodrug with the representative example shown in **Figure 6** highlighted in red. $IC_{50}$ values are tabulated and correlated to dose response in HeLa-HRE cells in **Table S1**.

**TrxR1/Trx1-dependent activation: MEF TrxR1 ko vs. wt cells (72 h treatment)**

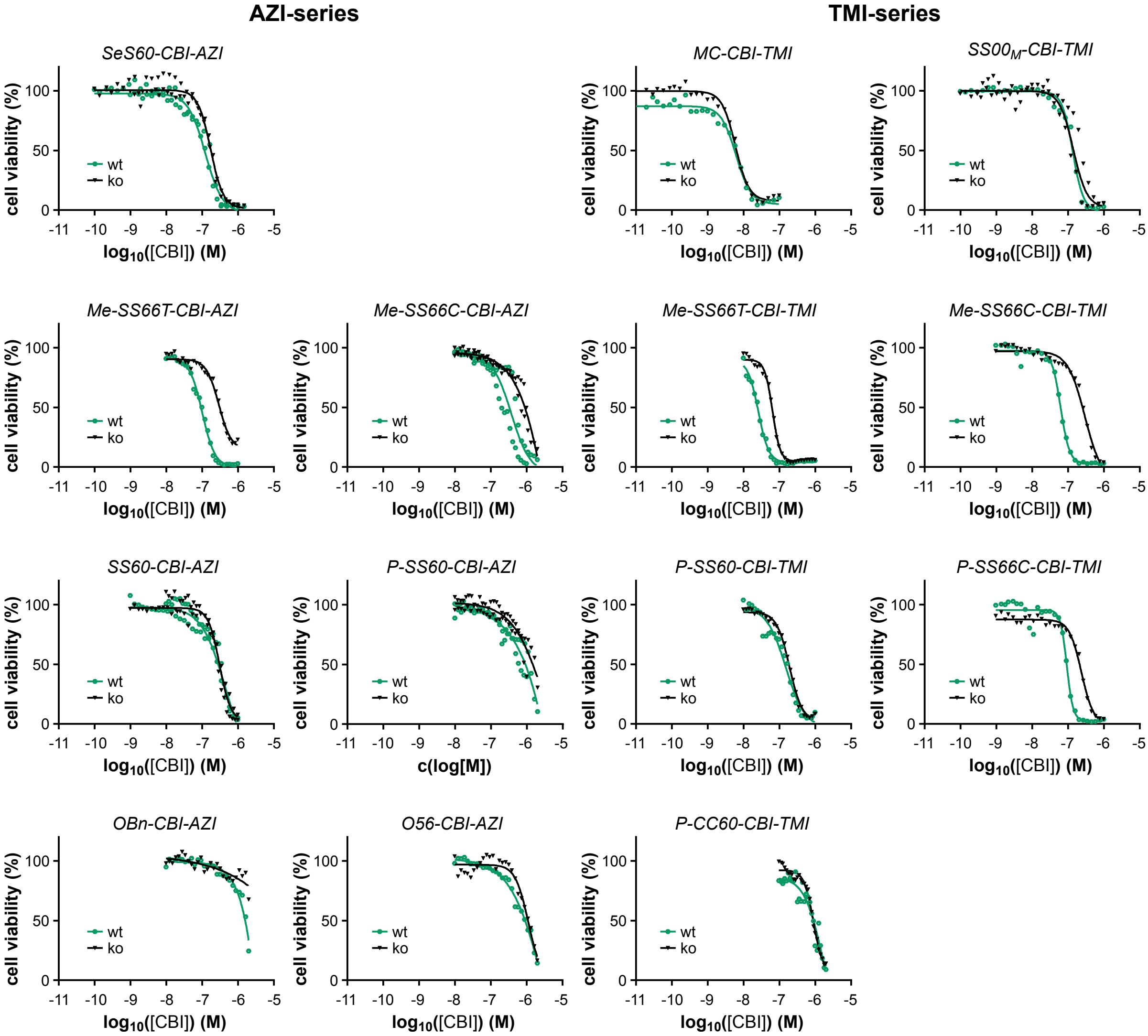


**Figure S9: Evaluation of reduction-activated prodrugs – Resazurin-based cytotoxicity assay in stable TrxR1 knockout cells.** MEF cells (*ko* vs. *wt*) were treated with prodrugs of the **CBI-TMI** and **CBI-AZI** series for 72 h at concentrations from 10 pm to 30 µM at 27 incremental single concentrations depending on expected potencies of the constructs. Data from at least 3 individual experiments was combined (no error bars shown for clarity) for each cell line. Representative compounds with strong cellular TrxR1-dependence are shown in **Figure 7**. $IC_{50}$ values (SD, n) are tabulated and correlated in **Table S1**.

#### 5. ProliFiler-140 cancer cell screening: cellular cytotoxicity ($IC_{50}$ values)

| | | | $IC_{50}$ [M] | | | | Reductive Activation Index | | |
|---|---|---|---|---|---|---|---|---|---|
| **Nr.** | **Entity** | **Cell line** | **P-SS66C** | **P-SS60** | **P-CC60** | **Dox** | **P-SS66C** | **P-SS60** | **P-CC60** |
| **76** | Blood | MV4-11 | 1.5E-09 | 2.8E-09 | 1.2E-08 | 4.3E-09 | 7.7 | 4.3 | 1 |
| **119** | Brain | SK-N-MC | 1.6E-09 | 2.7E-09 | 1.3E-08 | 1.5E-08 | 8.3 | 4.9 | 1 |
| **95** | Blood | OCI-LY19 | 2.8E-09 | 4.1E-09 | 1.5E-08 | 5.5E-09 | 5.4 | 3.7 | 1 |
| **48** | Endometrium | Ishikawa | 3.8E-09 | 7.1E-09 | 2.6E-08 | 7.7E-08 | 6.8 | 3.7 | 1 |
| **52** | Blood | JVM-3 | 3.4E-09 | 8.7E-09 | 3.1E-08 | 3.0E-08 | 9.3 | 3.6 | 1 |
| **115** | Bone | SK-ES-1 | 2.7E-09 | 5.7E-09 | 3.4E-08 | 2.6E-08 | 12 | 5.9 | 1 |
| **94** | Blood | OCI-AML5 | 6.0E-09 | 8.6E-09 | 3.4E-08 | 3.1E-08 | 5.7 | 3.9 | 1 |
| **126** | Blood | SU-DHL-5 | 2.6E-09 | 7.8E-09 | 5.0E-08 | 9.1E-09 | 19 | 6.4 | 1 |
| **31** | Lung | H460 | 5.5E-09 | 7.1E-09 | 5.1E-08 | 2.6E-08 | 9.3 | 7.2 | 1 |
| **47** | Duodenum | Hutu 80 | 9.6E-09 | 9.9E-09 | 5.4E-08 | 1.9E-08 | 5.6 | 5.4 | 1 |
| **77** | Blood | NALM-6 | 2.5E-09 | 9.8E-09 | 5.6E-08 | 2.3E-08 | 23 | 5.7 | 1 |
| **78** | Lung | NCI-H1048 | 4.0E-09 | 8.0E-09 | 5.9E-08 | 3.2E-08 | 15 | 7.4 | 1 |
| **74** | Blood | MOLM-13 | 5.3E-09 | 9.1E-09 | 6.1E-08 | 2.1E-08 | 12 | 6.6 | 1 |
| **125** | Ovary | SNU840 | 6.3E-08 | 3.8E-08 | 7.0E-08 | 1.3E-07 | 1.1 | 1.8 | 1 |
| **18** | Lung | COR-L279 | 7.2E-09 | 1.4E-08 | 7.4E-08 | 7.9E-08 | 10 | 5.1 | 1 |
| **139** | Blood | U-937 | 1.2E-08 | 1.6E-08 | 7.4E-08 | 2.6E-08 | 6.4 | 4.5 | 1 |
| **75** | Blood | MOLT-4 | 8.6E-09 | 1.3E-08 | 7.6E-08 | 2.6E-08 | 8.8 | 5.9 | 1 |
| **30** | Brain | H4 | 1.4E-08 | 1.5E-08 | 8.1E-08 | 8.2E-08 | 5.8 | 5.4 | 1 |
| **70** | Blood | MEC-1 | 1.1E-08 | 1.5E-08 | 8.9E-08 | 7.3E-08 | 8.2 | 5.9 | 1 |
| **4** | Ovary | A2780 | 8.1E-09 | 1.2E-08 | 8.9E-08 | 1.8E-08 | 11 | 7.4 | 1 |
| **23** | Lung | DV90 | 1.2E-08 | 1.3E-08 | 9.0E-08 | 1.9E-08 | 7.8 | 6.9 | 1 |
| **51** | Blood | Jurkat | 6.2E-09 | 1.6E-08 | 9.4E-08 | 4.0E-08 | 15 | 5.7 | 1 |
| **42** | Blood | HL-60 | 1.5E-08 | 1.8E-08 | 1.0E-07 | 2.6E-08 | 6.7 | 5.6 | 1 |
| **3** | Skin | A2058 | 1.1E-08 | 1.5E-08 | 1.0E-07 | 3.7E-08 | 9.4 | 6.8 | 1 |
| **103** | Lung | PC-9 | 1.4E-08 | 1.7E-08 | 1.1E-07 | 1.9E-07 | 8.1 | 6.7 | 1 |
| **19** | Ovary | COV434 | 1.5E-08 | 2.2E-08 | 1.1E-07 | 7.0E-08 | 7.7 | 5.1 | 1 |
| **1** | Kidney | 786-O | 2.9E-08 | 2.3E-08 | 1.2E-07 | 1.1E-07 | 4.1 | 5.2 | 1 |
| **87** | Lung | NCI-H2286 | 1.7E-08 | 1.9E-08 | 1.2E-07 | 4.4E-08 | 7.5 | 6.5 | 1 |
| **137** | Bone | U2OS | 3.2E-08 | 2.4E-08 | 1.3E-07 | 1.1E-07 | 4.1 | 5.4 | 1 |
| **93** | Blood | OCI-AML3 | 1.3E-08 | 3.3E-08 | 1.3E-07 | 7.6E-08 | 10 | 4.0 | 1 |
| **64** | Blood | M07e | 3.2E-08 | 1.6E-08 | 1.4E-07 | 4.6E-08 | 4.3 | 8.6 | 1 |
| **136** | Blood | U-266 | 9.4E-09 | 2.2E-08 | 1.4E-07 | 1.1E-07 | 15 | 6.3 | 1 |
| **86** | Lung | NCI-H2110 | 1.8E-08 | 2.7E-08 | 1.4E-07 | 3.6E-08 | 8.0 | 5.4 | 1 |
| **69** | Breast | MDA-MB-468 | 1.9E-08 | 3.1E-08 | 1.5E-07 | 4.7E-08 | 7.9 | 4.8 | 1 |
| **111** | Lung | SCLC-21H | 2.6E-08 | 3.2E-08 | 1.6E-07 | 2.9E-07 | 6.2 | 4.9 | 1 |
| **116** | Lung | SK-LU-1 | 2.3E-08 | 2.7E-08 | 1.6E-07 | 2.7E-07 | 7.1 | 6.0 | 1 |
| **5** | Skin | A375 | 5.5E-08 | 3.2E-08 | 1.6E-07 | 4.8E-08 | 3.0 | 5.1 | 1 |
| **34** | Breast | HCC38 | 1.4E-08 | 1.9E-08 | 1.7E-07 | 5.2E-08 | 12 | 8.7 | 1 |
| **22** | Prostate | DU-145 | 1.6E-08 | 3.6E-08 | 1.8E-07 | 1.5E-07 | 11 | 5.1 | 1 |
| **100** | Blood | P31/FUJ | 2.3E-08 | 3.7E-08 | 1.9E-07 | 1.0E-07 | 8.3 | 5.1 | 1 |
| **99** | Ovary | OVK18 | 4.6E-08 | 4.3E-08 | 2.0E-07 | 3.4E-08 | 4.4 | 4.7 | 1 |
| **97** | Ovary | OV56 | 7.2E-08 | 6.5E-08 | 2.0E-07 | 2.0E-07 | 2.8 | 3.1 | 1 |
| **66** | Breast | MCF-7 | 3.1E-08 | 4.5E-08 | 2.1E-07 | 2.3E-07 | 7.0 | 4.8 | 1 |
| **62** | Colon | LOVO | 1.9E-08 | 3.4E-08 | 2.2E-07 | 1.1E-07 | 12 | 6.6 | 1 |
| **122** | Stomach | SNU-1 | 2.3E-08 | 3.0E-08 | 2.3E-07 | 5.9E-08 | 10 | 7.6 | 1 |
| **109** | Bone | Saos-2 EC | 6.6E-08 | 3.9E-08 | 2.3E-07 | 2.0E-07 | 3.5 | 6.0 | 1 |
| **40** | Ovary | HeLa | 9.9E-09 | 1.7E-08 | 2.4E-07 | 8.0E-08 | 24 | 14 | 1 |
| **88** | Lung | NCI-H292 | 3.8E-08 | 4.8E-08 | 2.6E-07 | 7.2E-08 | 6.9 | 5.4 | 1 |
| **7** | Skin | A431 | 2.0E-08 | 4.0E-08 | 2.6E-07 | 1.7E-07 | 13 | 6.5 | 1 |
| **89** | Lung | NCI-H441 | 5.1E-08 | 5.4E-08 | 2.9E-07 | 3.5E-07 | 5.8 | 5.4 | 1 |
| **106** | Colon | RKO | 2.1E-08 | 4.5E-08 | 3.0E-07 | 9.9E-08 | 14 | 6.6 | 1 |

***Start of Table S2 (part 1 of 3):*** *ProliFiler-140 potencies and reductive index.* *Table continues on next page.*

| Nr. | Entity | Cell line | IC$_{50}$ [M] P-SS66C | P-SS60 | P-CC60 | Dox | Reductive Activation Index P-SS66C | P-SS60 | P-CC60 |
|---|---|---|---|---|---|---|---|---|---|
| 8 | Kidney | A498 | 9.4E-08 | 5.0E-08 | 3.0E-07 | 3.2E-07 | 3.2 | 6.0 | 1 |
| 79 | Lung | NCI-H1437 | 1.8E-08 | 5.2E-08 | 3.0E-07 | 1.4E-07 | 16 | 5.8 | 1 |
| 61 | Lung | LOU-NH91 | 3.5E-08 | 7.5E-08 | 3.1E-07 | 2.9E-07 | 8.9 | 4.1 | 1 |
| 20 | Head/Neck | Detroit562 | 1.9E-08 | 4.3E-08 | 3.2E-07 | 1.5E-07 | 17 | 7.4 | 1 |
| 44 | Fibrosarcoma | HT-1080 | 2.1E-08 | 5.0E-08 | 3.2E-07 | 7.6E-08 | 15 | 6.4 | 1 |
| 14 | Kidney | Caki-1 | 6.3E-08 | 6.3E-08 | 3.3E-07 | 3.4E-07 | 5.2 | 5.2 | 1 |
| 128 | Colon | SW480 | 5.5E-08 | 6.5E-08 | 3.3E-07 | 1.2E-07 | 6.0 | 5.1 | 1 |
| 108 | Blood | RPMI 8226 | 1.2E-08 | 3.9E-08 | 3.4E-07 | 8.8E-08 | 28 | 8.8 | 1 |
| 16 | Lung | Calu-6 | 5.9E-08 | 5.2E-08 | 3.4E-07 | 7.7E-07 | 5.8 | 6.6 | 1 |
| 58 | Blood | KMS-12-BM | 3.3E-08 | 6.6E-08 | 3.4E-07 | 1.6E-07 | 10 | 5.2 | 1 |
| 114 | Breast | SK-BR-3 | 4.2E-08 | 7.4E-08 | 3.7E-07 | 1.3E-07 | 8.8 | 5.0 | 1 |
| 71 | Pancreas | Mia PaCA 2 | 8.6E-08 | 6.8E-08 | 3.8E-07 | 7.4E-08 | 4.4 | 5.5 | 1 |
| 113 | Bone | SJSA-1 | 9.8E-08 | 2.4E-08 | 3.8E-07 | 4.0E-07 | 3.9 | 16 | 1 |
| 110 | Stomach | SCH | 1.5E-08 | 5.2E-08 | 3.9E-07 | 1.4E-07 | 25 | 7.5 | 1 |
| 134 | Brain | U118MG | 1.9E-07 | 1.4E-07 | 3.9E-07 | 2.6E-07 | 2.1 | 2.8 | 1 |
| 29 | Lung | H2228 | 4.5E-08 | 5.7E-08 | 4.0E-07 | 5.7E-08 | 8.8 | 7.0 | 1 |
| 85 | Lung | NCI-H2009 | 2.7E-08 | 6.2E-08 | 4.2E-07 | 1.4E-07 | 16 | 6.8 | 1 |
| 82 | Lung | NCI-H1581 | 4.7E-08 | 4.2E-08 | 4.3E-07 | 1.5E-07 | 9.3 | 10 | 1 |
| 121 | Ovary | SK-OV3 | 3.7E-08 | 6.2E-08 | 4.4E-07 | 1.8E-07 | 12 | 7.1 | 1 |
| 36 | Colon | HCT116 | 3.2E-08 | 6.9E-08 | 4.4E-07 | 1.1E-07 | 14 | 6.4 | 1 |
| 105 | Lung | RERF-LC-MS | 3.1E-08 | 6.4E-08 | 4.6E-07 | 1.7E-07 | 15 | 7.1 | 1 |
| 127 | Brain | SW-1783 | 4.8E-08 | 6.7E-08 | 4.8E-07 | 3.0E-07 | 9.9 | 7.2 | 1 |
| 73 | Stomach | MKN-45 | 8.2E-08 | 9.1E-08 | 4.9E-07 | 3.2E-07 | 5.9 | 5.4 | 1 |
| 27 | Lung | EPLC-272H | 4.6E-08 | 7.9E-08 | 5.1E-07 | 2.0E-07 | 11 | 6.4 | 1 |
| 102 | Prostate | PC3 | 4.0E-08 | 5.8E-08 | 5.2E-07 | 7.4E-07 | 13 | 8.9 | 1 |
| 37 | Colon | HCT-15 | 1.4E-07 | 1.0E-07 | 5.5E-07 | 6.4E-07 | 3.9 | 5.2 | 1 |
| 57 | Blood | KG-1 a | 1.1E-07 | 1.0E-07 | 5.5E-07 | 4.9E-07 | 5.2 | 5.3 | 1 |
| 2 | Brain | A172 | 1.1E-07 | 1.2E-07 | 5.7E-07 | 9.4E-08 | 5.3 | 4.9 | 1 |
| 112 | Ovary | SiHa | 1.1E-07 | 1.1E-07 | 6.1E-07 | 1.5E-07 | 5.4 | 5.5 | 1 |
| 56 | Blood | KG-1 | 1.9E-07 | 1.3E-07 | 6.2E-07 | 1.2E-06 | 3.2 | 4.7 | 1 |
| 46 | Liver | HuH7 | 1.0E-07 | 1.1E-07 | 6.5E-07 | 2.1E-07 | 6.5 | 5.7 | 1 |
| 133 | Brain | T98G | 8.1E-08 | 8.6E-08 | 6.6E-07 | 1.2E-06 | 8.2 | 7.6 | 1 |
| 96 | Blood | OPM-2 | 2.6E-08 | 1.2E-07 | 6.6E-07 | 2.4E-07 | 25 | 5.4 | 1 |
| 13 | Pancreas | BxPC-3 | 4.6E-08 | 1.1E-07 | 6.6E-07 | 2.8E-07 | 14 | 6.1 | 1 |
| 130 | Colon | SW948 | 1.4E-07 | 1.3E-07 | 6.8E-07 | 3.5E-07 | 4.8 | 5.3 | 1 |
| 118 | Brain | SK-N-FI | 1.4E-07 | 1.2E-07 | 6.8E-07 | 1.4E-06 | 4.9 | 5.7 | 1 |
| 129 | Colon | SW620 | 1.5E-07 | 1.4E-07 | 7.0E-07 | 3.1E-07 | 4.6 | 5.0 | 1 |
| 54 | Blood | KARPAS 299 | 1.1E-07 | 1.6E-07 | 7.1E-07 | 7.3E-08 | 6.5 | 4.4 | 1 |
| 123 | Stomach | SNU-16 | 6.2E-08 | 1.2E-07 | 7.1E-07 | 1.4E-07 | 11 | 5.8 | 1 |
| 63 | Blood | LP-1 | 1.0E-07 | 1.7E-07 | 7.2E-07 | 6.6E-07 | 6.9 | 4.2 | 1 |
| 49 | Bladder | J82 | 9.6E-08 | 9.9E-08 | 7.2E-07 | 8.0E-07 | 7.5 | 7.3 | 1 |
| 59 | Brain | LN229 | 1.3E-07 | 1.4E-07 | 7.2E-07 | 2.2E-07 | 5.4 | 5.2 | 1 |
| 28 | Lung | H1299 | 5.4E-08 | 7.9E-08 | 7.3E-07 | 2.2E-07 | 14 | 9.3 | 1 |
| 67 | Breast | MDA MB 231 | 1.4E-07 | 1.7E-07 | 7.4E-07 | 2.8E-07 | 5.4 | 4.2 | 1 |
| 107 | Endometrium | RL95-2 | 1.3E-07 | 1.5E-07 | 7.4E-07 | 2.0E-07 | 5.9 | 4.8 | 1 |
| 120 | Brain | SK-N-SH | 1.1E-07 | 1.5E-07 | 7.5E-07 | 9.8E-08 | 6.5 | 5.1 | 1 |
| 90 | Lung | NCI-H82 | 8.8E-08 | 1.5E-07 | 7.5E-07 | 6.2E-07 | 8.5 | 5.0 | 1 |
| 91 | Lung | NCI-H838 | 7.4E-08 | 1.2E-07 | 7.5E-07 | 1.5E-07 | 10 | 6.2 | 1 |
| 17 | Colon | Colo 205 | 8.6E-08 | 1.6E-07 | 7.6E-07 | 1.5E-07 | 8.8 | 4.9 | 1 |
| 9 | Lung | A549 | 3.0E-08 | 1.2E-07 | 7.7E-07 | 3.1E-07 | 25 | 6.4 | 1 |
| 104 | Lung | RERF-LC-Ad2 | 1.4E-07 | 1.9E-07 | 7.8E-07 | 4.6E-07 | 5.5 | 4.2 | 1 |
| 21 | Colon | DLD-1 | 1.3E-07 | 1.7E-07 | 7.9E-07 | 1.4E-07 | 6.3 | 4.7 | 1 |
| 83 | Lung | NCI-H1703 | 5.1E-08 | 1.2E-07 | 8.1E-07 | 9.2E-08 | 16 | 6.7 | 1 |
| 65 | Ovary | MCAS | 1.4E-07 | 1.3E-07 | 8.2E-07 | 1.1E-07 | 5.9 | 6.5 | 1 |
| 84 | Lung | NCI-H1838 | 7.7E-08 | 8.5E-08 | 8.2E-07 | 2.5E-07 | 11 | 9.6 | 1 |

***Middle of Table S2 (part 2 of 3):*** *ProliFiler-140 potencies and reductive index. Table continues on next page.*

| | | | IC$_{50}$ [M] | | | | Reductive Activation Index | | |
|---|---|---|---|---|---|---|---|---|---|
| Nr. | Entity | Cell line | P-SS66C | P-SS60 | P-CC60 | Dox | P-SS66C | P-SS60 | P-CC60 |
| 6 | Lung | A427 | 4.4E-08 | 1.6E-07 | 8.2E-07 | 6.6E-08 | 19 | 5.0 | 1 |
| 60 | Prostate | LnCap | 2.7E-08 | 1.1E-07 | 8.9E-07 | 1.9E-07 | 33 | 8.0 | 1 |
| 33 | Breast | HCC202 | 9.1E-08 | 1.3E-07 | 9.0E-07 | 7.6E-07 | 9.8 | 7.1 | 1 |
| 26 | Ovary | EFO-27 | 6.2E-08 | 1.6E-07 | 9.0E-07 | 2.9E-07 | 15 | 5.7 | 1 |
| 132 | Colon | T84 | 1.6E-07 | 1.8E-07 | 9.1E-07 | 1.4E-06 | 5.7 | 5.1 | 1 |
| 24 | Breast | EFM-19 | 3.1E-07 | 2.3E-07 | 9.6E-07 | 4.7E-07 | 3.1 | 4.2 | 1 |
| 43 | Stomach | Hs 746T | 7.5E-08 | 1.3E-07 | 9.9E-07 | 4.0E-07 | 13 | 7.7 | 1 |
| 32 | Breast | HCC 1569 | 3.6E-08 | 1.2E-07 | 9.9E-07 | 7.0E-07 | 28 | 8.2 | 1 |
| 45 | Colon | HT-29 | 1.1E-07 | 1.9E-07 | 1.0E-06 | 3.7E-07 | 9.5 | 5.3 | 1 |
| 98 | Ovary | OVCAR-3 | 7.9E-08 | 1.2E-07 | 1.0E-06 | 3.7E-07 | 13 | 8.7 | 1 |
| 117 | Skin | SK-MEL-3 | 1.6E-07 | 1.8E-07 | 1.0E-06 | 5.1E-07 | 6.6 | 5.8 | 1 |
| 131 | Breast | T-47D | 9.8E-08 | 2.3E-07 | 1.0E-06 | 4.3E-07 | 11 | 4.5 | 1 |
| 81 | Lung | NCI-H1573 | 9.0E-08 | 1.8E-07 | 1.1E-06 | 3.0E-06 | 12 | 6.1 | 1 |
| 68 | Skin | MDA MB 435 | 1.3E-07 | 1.8E-07 | 1.1E-06 | 4.8E-07 | 8.6 | 6.1 | 1 |
| 53 | Blood | K562 | 1.3E-07 | 2.1E-07 | 1.1E-06 | 4.4E-07 | 8.6 | 5.3 | 1 |
| 12 | Breast | BT-20 | 1.2E-07 | 2.4E-07 | 1.1E-06 | 2.3E-07 | 9.1 | 4.6 | 1 |
| 92 | Stomach | NCI-N87 | 7.5E-08 | 1.4E-07 | 1.1E-06 | 2.0E-07 | 15 | 8.1 | 1 |
| 138 | Brain | U87MG | 1.5E-07 | 2.3E-07 | 1.2E-06 | 1.6E-07 | 8.0 | 5.4 | 1 |
| 41 | Liver | Hep3B2.1-7 | 1.7E-07 | 2.4E-07 | 1.3E-06 | 5.3E-07 | 7.8 | 5.6 | 1 |
| 101 | Pancreas | PANC-1 | 2.0E-07 | 2.5E-07 | 1.4E-06 | 8.0E-07 | 6.9 | 5.4 | 1 |
| 35 | Lung | HCC827 | 9.1E-08 | 1.9E-07 | 1.4E-06 | 3.5E-07 | 15 | 7.3 | 1 |
| 140 | Breast | ZR-75-1 | 3.1E-07 | 2.2E-07 | 1.4E-06 | 1.3E-06 | 4.6 | 6.4 | 1 |
| 11 | Lung | BEN | 5.3E-08 | 1.1E-07 | 1.4E-06 | 4.1E-06 | 27 | 13 | 1 |
| 15 | Kidney | Caki-2 | 2.7E-07 | 2.1E-07 | 1.5E-06 | 1.0E-06 | 5.6 | 7.2 | 1 |
| 72 | Stomach | MKN-1 | 1.4E-07 | 2.1E-07 | 1.5E-06 | 5.2E-08 | 11.1 | 7.3 | 1 |
| 50 | Breast | JIMT-1 | 1.0E-07 | 1.9E-07 | 1.6E-06 | 1.3E-06 | 15 | 8.4 | 1 |
| 55 | Stomach | Kato III | 2.2E-07 | 3.7E-07 | 1.6E-06 | 9.0E-07 | 7.4 | 4.3 | 1 |
| 80 | Lung | NCI-H1563 | 1.1E-07 | 2.0E-07 | 1.7E-06 | 5.4E-07 | 15 | 8.6 | 1 |
| 25 | Breast | EFM-192A | 2.7E-07 | 3.0E-07 | 1.8E-06 | 7.3E-07 | 6.6 | 5.9 | 1 |
| 38 | Endometrium | HEC-1-A | 1.0E-07 | 2.8E-07 | 1.8E-06 | 3.3E-07 | 18 | 6.5 | 1 |
| 124 | Stomach | SNU-216 | 9.2E-08 | 2.7E-07 | 1.8E-06 | 3.9E-07 | 20 | 6.8 | 1 |
| 135 | Brain | U251MG | 1.2E-07 | 1.9E-07 | 2.0E-06 | 7.8E-07 | 17 | 11 | 1 |
| 10 | Pancreas | AsPC-1 | 2.6E-07 | 2.6E-07 | 2.1E-06 | 1.2E-06 | 8.2 | 8.2 | 1 |
| 39 | Endometrium | HEC-1-B | 4.0E-07 | 8.3E-07 | 3.3E-06 | 2.4E-06 | 8.3 | 4.0 | 1 |

***End of Table S2 (part 3 of 3):*** *ProliFiler-140 potencies and reductive index.*

**The first four data columns** show the sensitivities of 140 cancer cell lines to bioreductive prodrugs **P-SS60-CBI-TMI** and **P-SS66C-CBI-TMI**, and to **P-CC60-CBI-TMI** (hydrolysis control) and doxorubicin (assay control, reference compound). Cell lines were sorted by increasing IC$_{50}$ values for **P-CC60-CBI-TMI**.

The **final four columns** show the ratios for the IC$_{50}$ values divided by those of **P-CC60-CBI-TMI** (***reductive activation index***): aiming to identify in which lines one or both prodrugs show significantly higher potency than in others, which may reflect greater degrees of reductive rather than only hydrolytic activation in those lines. *Cell lines with a >10× fold-index for **P-SS66C-CBI-TMI** are highlighted in yellow*.

For both mathematical and biological reasons, the index itself is not a quantitative measurement of bioreduction, and directly comparing indices across very different situations (e.g. for one prodrug in one cell line, compared to a different prodrug in a different cell line) is not an appropriate way to quantify differences in bioreductive activation. Quantitative analysis of bioreduction would instead require screening with a direct absolute readout - such as fluorogenic probes - not with the indirect and cell-line-dependent readout of cell death given here. We believe that such quantitative data would also be more appropriate for testing whether biological features (e.g. tissues of origin, gene expression patterns) might correlate usefully to reductive activation.

**ProliFiler-140 cell line panel part I (lines 1-28)**

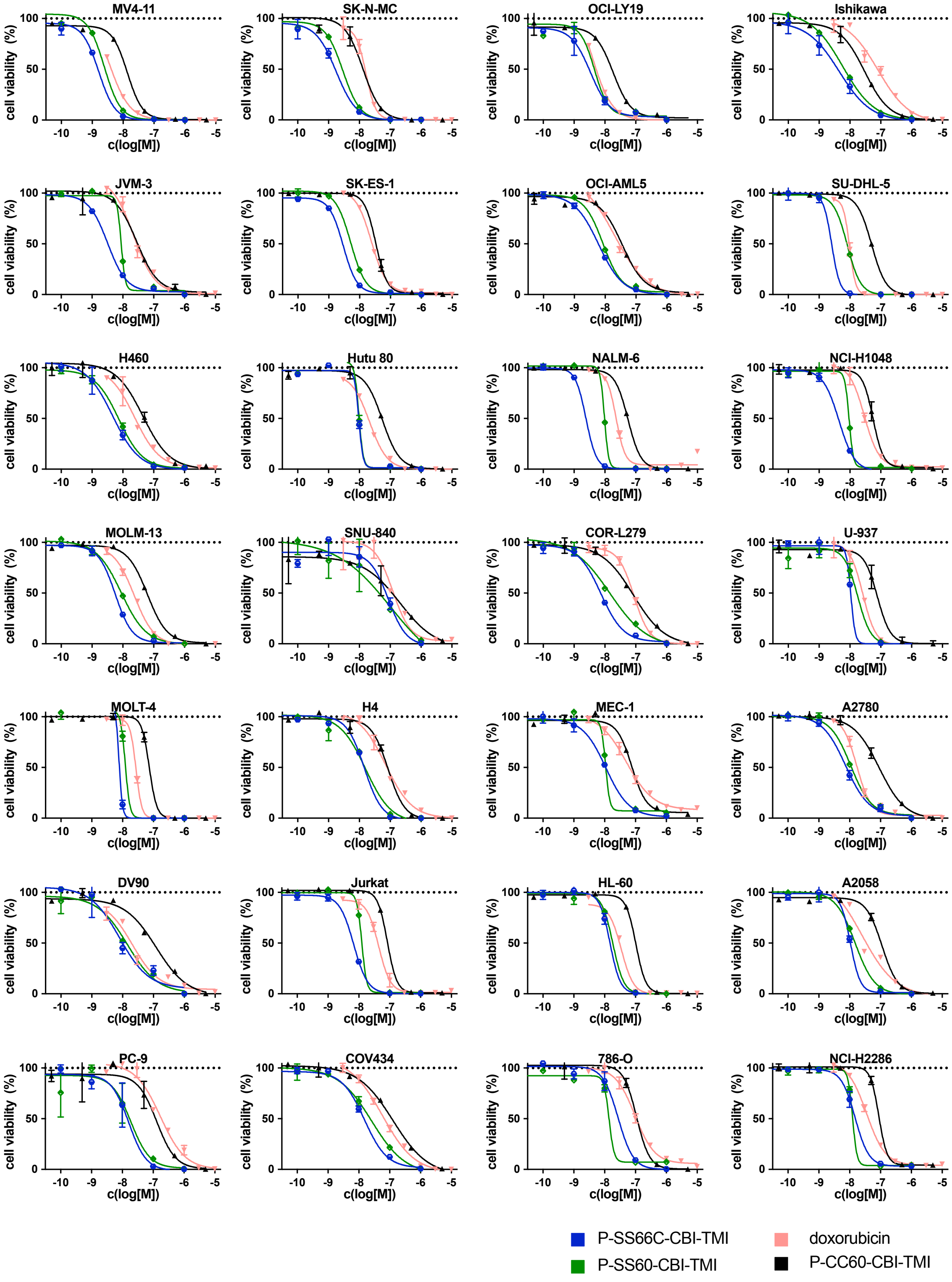


***Start of Fig S10 (part 1 of 5: figure continues on next page): ProliFiler-140 screen for cytotoxicities***

**ProliFiler-140 cell line panel part II (lines 29-56)**

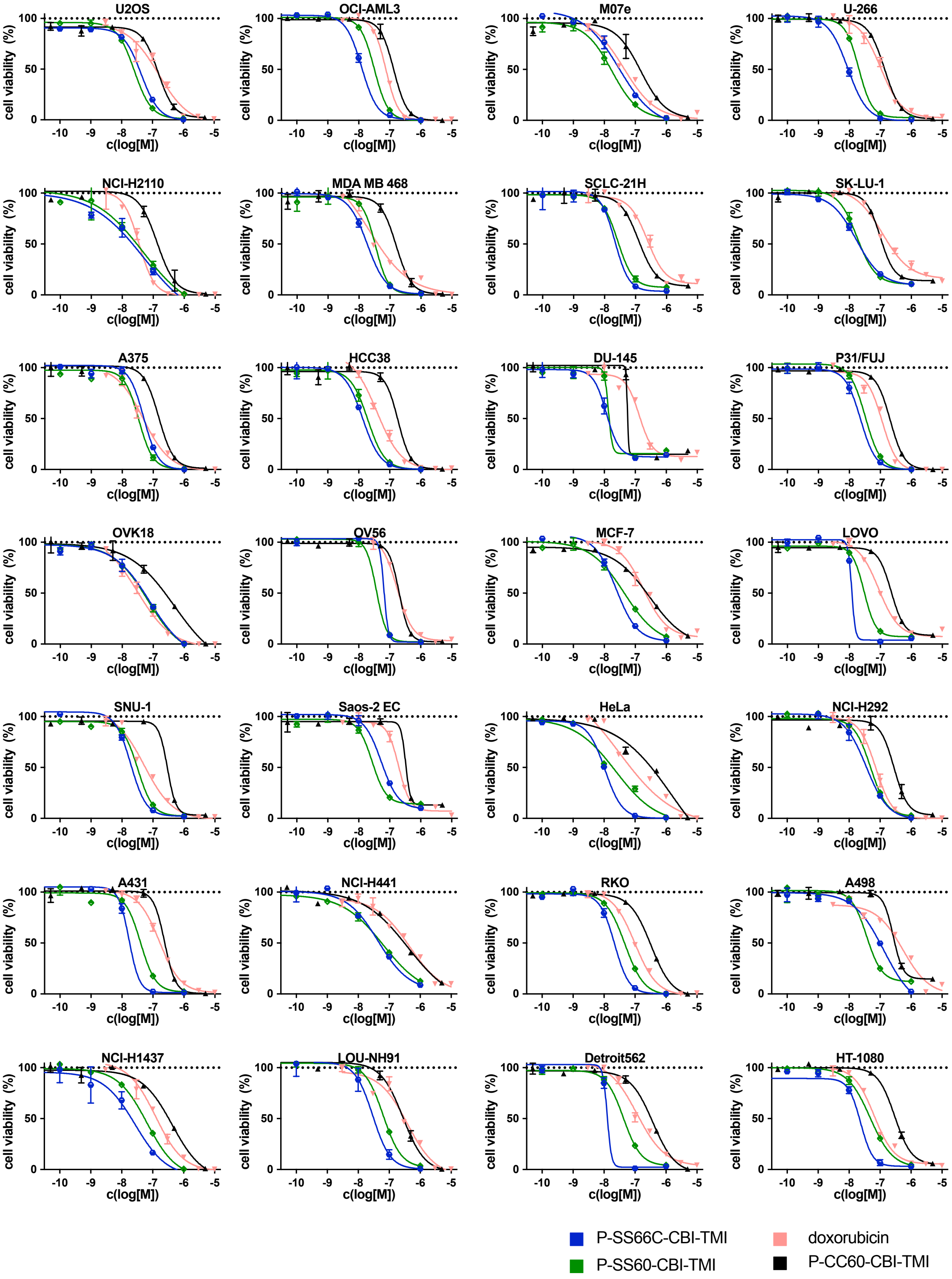


***Continuation of Fig S10 (part 2 of 5: figure continues on next page): ProliFiler-140 screen for cytotoxicities***

**ProliFiler-140 cell line panel part III (lines 57-84))**

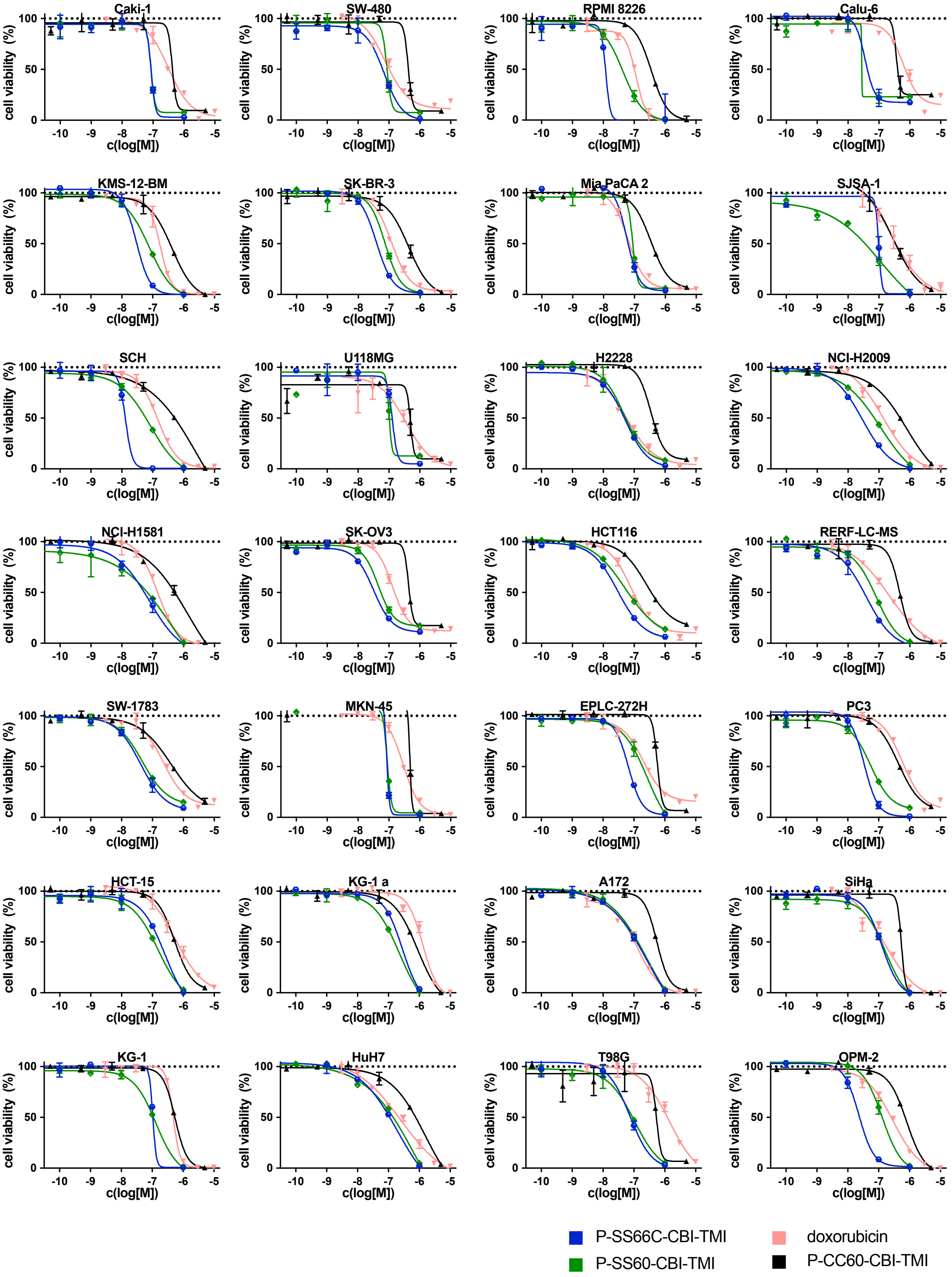


***Continuation of Fig S10 (part 3 of 5: figure continues on next page): ProliFiler-140 screen for cytotoxicities***

**ProliFiler-140 cell line panel part IV (lines 85-112)**

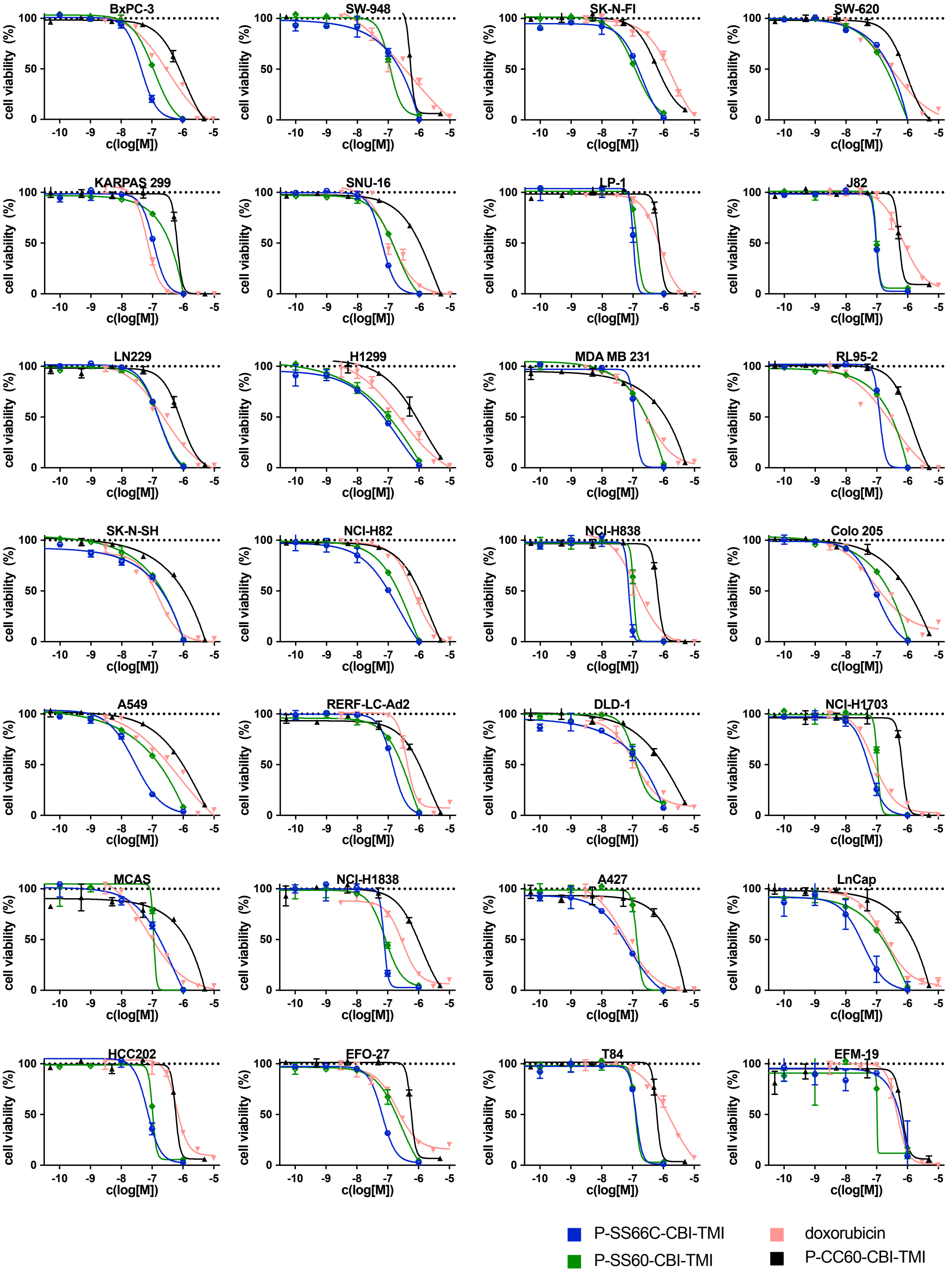


***Continuation of Fig S10 (part 4 of 5: figure continues on next page): ProliFiler-140 screen for cytotoxicities***

**ProliFiler-140 cell line panel part V (lines 113-140)**

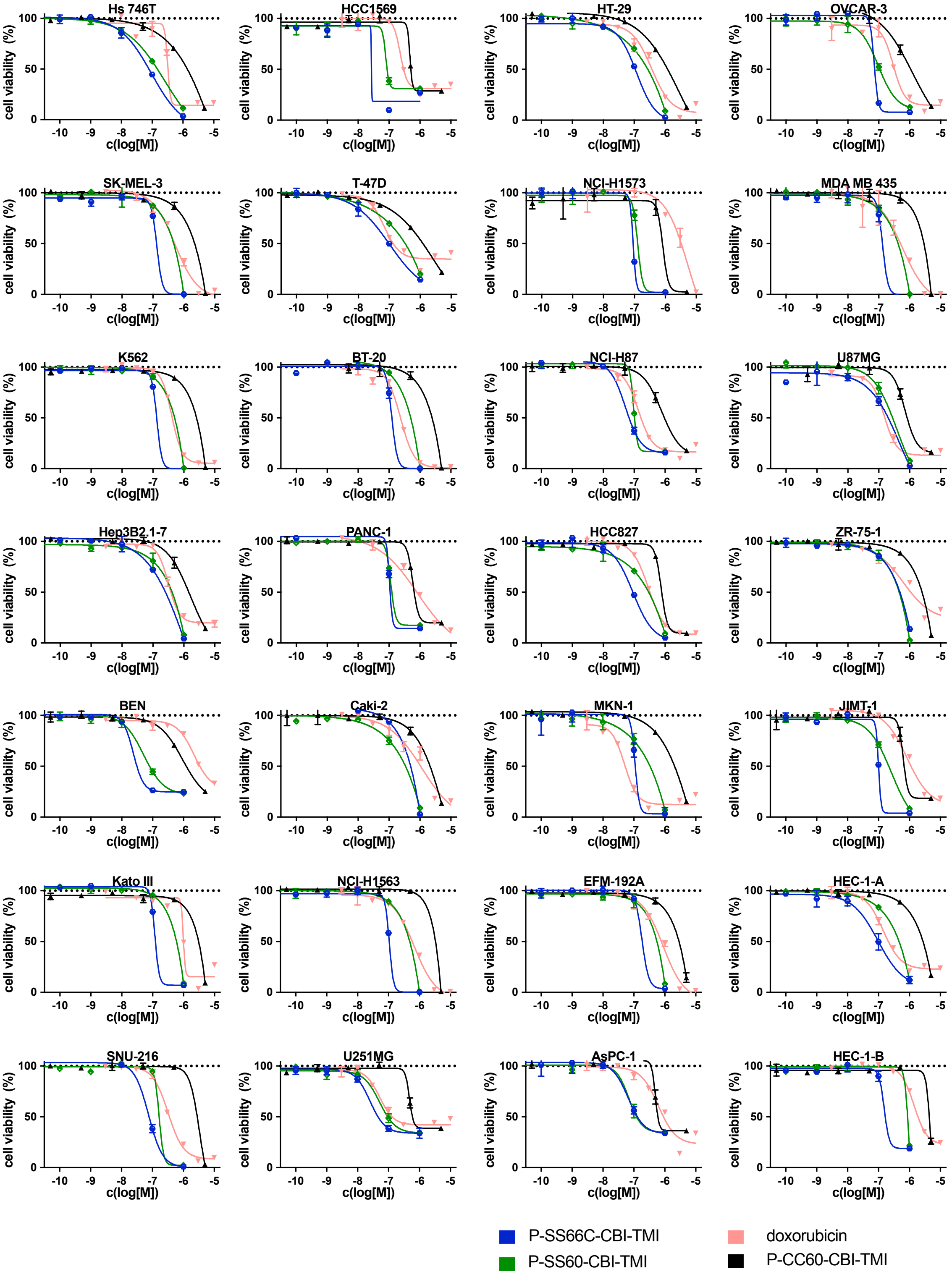


**End of Fig S10 (part 5 of 5):** ProliFiler-140 cancer cell line screening for cytotoxicity of the prodrugs. 8-data-point dose response curves from CellTiter-Glo® viability readout. All graphs within this figure have been presented in order of decreasing potency of **P-CC60-CBI-TMI**.

#### ProliFiler-140 cancer cell screen: Summary and population analysis

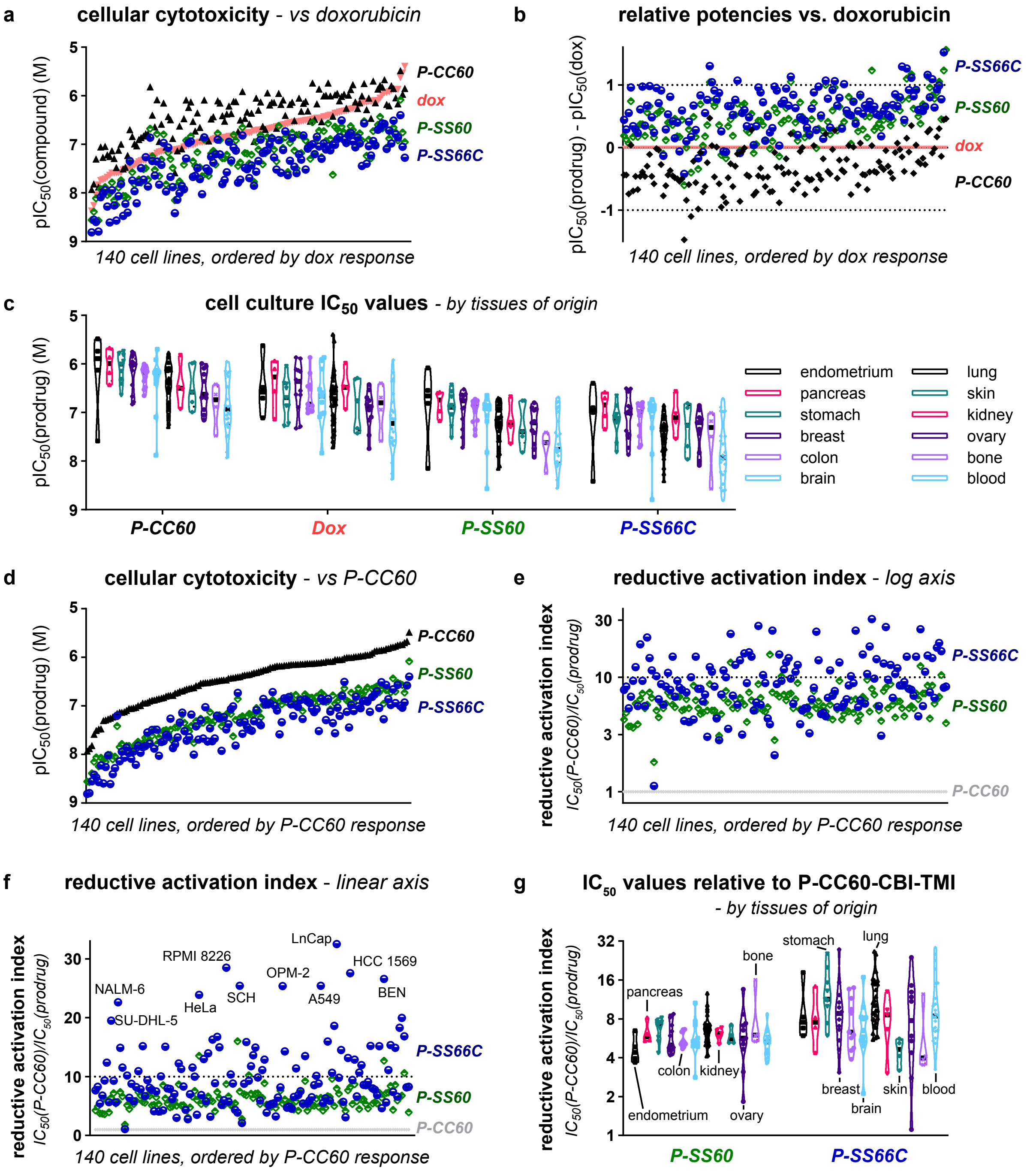


**Figure S11: ProliFiler-140 cancer cell screening – population analysis.** (**a**) $IC_{50}$ values for **P-SS60-CBI-TMI** (**P-SS60**), **P-SS66C-CBI-TMI** (**P-SS66C**) and **P-CC60-CBI-TMI** (**P-CC60**) are shown for 140 cell lines sorted by decreasing response to **doxorubicin** (**dox**). (**b**) $pIC_{50}$ values (= $-\log_{10}(IC_{50})$ (M)) compared to those for **dox**, sorted by decreasing response to **dox**. (**c**) $IC_{50}$ values sorted by tissue of origin. (**d**) $IC_{50}$ values sorted by decreasing response to **P-CC60**. (**e**,**f**) Reductive activation index (= $IC_{50}$(**P-CC60**)/$IC_{50}$(prodrug)) values, sorted by decreasing response to **P-CC60**. (**g**) Reductive activation index values for **P-SS66C** and **P-SS60**, grouped by tissues of origin.

#### 6. NCI-60 cancer cell screening: growth inhibition ($GI_{50}$ values)

| | $GI_{50}$ [M] | | | | | Reductive Activation Index | | | |
|---|---|---|---|---|---|---|---|---|---|
| **Cell line** | **P-CC60** | **P-SS60** | **P-SS66C** | **Me-SS66C** | **Me-SS66T** | **P-SS60** | **P-SS66C** | **Me-SS66C** | **Me-SS66T** |
| SR | 2.8E-07 | 3.4E-08 | 2.0E-08 | 3.7E-08 | 4.0E-09 | 8 | 14 | 7 | 69 |
| MCF7 | 3.2E-07 | 3.5E-08 | 1.5E-08 | 2.8E-08 | 5.2E-09 | 9 | 21 | 11 | 61 |
| DU-145 | 4.6E-07 | 5.9E-08 | 2.5E-08 | 6.3E-08 | 3.5E-09 | 8 | 19 | 7 | 131 |
| NCI-H23 | 4.7E-07 | 5.5E-08 | 1.9E-08 | 4.5E-08 | 9.5E-09 | 9 | 25 | 10 | 49 |
| ACHN | 4.9E-07 | 5.6E-08 | 4.2E-08 | 2.8E-08 | 9.6E-09 | 9 | 12 | 17 | 52 |
| NCI-H460 | 5.1E-07 | 5.6E-08 | 2.2E-08 | 4.1E-08 | 4.8E-09 | 9 | 23 | 12 | 107 |
| MOLT-4 | 5.9E-07 | 9.0E-08 | 6.3E-08 | 7.0E-08 | 9.9E-09 | 7 | 9 | 9 | 60 |
| HOP-62 | 5.9E-07 | 1.1E-07 | 4.1E-08 | 2.1E-07 | 1.2E-08 | 5 | 14 | 3 | 48 |
| K-562 | 6.8E-07 | 1.6E-07 | 8.9E-08 | 4.0E-07 | 2.5E-08 | 4 | 8 | 2 | 28 |
| MDA-MB-468 | 6.9E-07 | 1.7E-07 | 3.7E-08 | 1.6E-07 | 1.2E-08 | 4 | 18 | 4 | 60 |
| HL-60(TB) | 7.5E-07 | 1.2E-07 | 4.8E-08 | 5.2E-08 | 1.2E-08 | 6 | 16 | 14 | 63 |
| CCRF-CEM | 9.9E-07 | 2.2E-07 | 2.7E-07 | 1.1E-07 | 2.6E-08 | 5 | 4 | 9 | 38 |
| RXF 393 | 1.1E-06 | 1.5E-07 | 3.5E-08 | 1.3E-07 | 9.5E-09 | 7 | 31 | 8 | 111 |
| LOX IMVI | 1.1E-06 | 2.1E-07 | 6.9E-08 | 1.7E-07 | 1.2E-08 | 5 | 16 | 7 | 90 |
| SN12C | 1.2E-06 | 2.9E-07 | 4.5E-08 | 2.0E-07 | 1.7E-08 | 4 | 27 | 6 | 70 |
| SF-539 | 1.3E-06 | 1.7E-07 | 3.9E-08 | 1.7E-07 | 1.0E-08 | 8 | 34 | 8 | 131 |
| COLO 205 | 1.4E-06 | 3.4E-07 | 1.5E-07 | 3.3E-07 | 4.9E-08 | 4 | 9 | 4 | 28 |
| NCI-H226 | 1.4E-06 | 4.3E-07 | 2.5E-07 | 1.0E-06 | 9.5E-08 | 3 | 6 | 1 | 15 |
| RPMI-8226 | 1.5E-06 | 2.2E-07 | 1.4E-07 | 1.6E-07 | 3.4E-08 | 7 | 10 | 9 | 43 |
| CAKI-1 | 1.5E-06 | 2.7E-07 | 1.7E-07 | 2.5E-07 | 2.7E-08 | 5 | 9 | 6 | 55 |
| HCT-116 | 1.5E-06 | 1.6E-07 | 1.5E-07 | 1.9E-07 | 1.8E-08 | 9 | 10 | 8 | 83 |
| U251 | 1.5E-06 | 1.6E-07 | 8.1E-08 | 1.4E-07 | 1.1E-08 | 9 | 19 | 11 | 139 |
| HCC-2998 | 1.5E-06 | 1.6E-07 | 5.4E-08 | 1.4E-07 | 1.5E-08 | 10 | 28 | 11 | 99 |
| HT29 | 1.5E-06 | 2.7E-07 | 1.4E-07 | 2.3E-07 | 3.4E-08 | 6 | 11 | 7 | 45 |
| M14 | 1.5E-06 | 1.9E-07 | 1.2E-07 | 2.5E-07 | 1.1E-08 | 8 | 12 | 6 | 145 |
| UACC-62 | 1.5E-06 | 2.7E-07 | 1.4E-07 | 1.5E-07 | 1.5E-08 | 6 | 11 | 10 | 101 |
| SF-268 | 1.5E-06 | 1.9E-07 | 5.8E-08 | 1.8E-07 | 1.5E-08 | 8 | 26 | 9 | 104 |
| NCI-H322M | 1.6E-06 | 2.4E-07 | 5.0E-08 | 2.4E-07 | 2.3E-08 | 7 | 31 | 6 | 66 |
| SF-295 | 1.6E-06 | 1.5E-07 | 1.1E-07 | 1.3E-07 | 9.7E-09 | 11 | 14 | 12 | 160 |
| MALME-3M | 1.6E-06 | 2.8E-07 | 1.3E-07 | 2.1E-07 | 1.4E-08 | 6 | 12 | 7 | 115 |
| BT-549 | 1.6E-06 | 2.3E-07 | 1.4E-07 | 1.7E-07 | 1.5E-08 | 7 | 11 | 10 | 103 |
| SK-MEL-5 | 1.6E-06 | 1.7E-07 | 1.1E-07 | 1.7E-07 | 1.3E-08 | 10 | 15 | 10 | 124 |
| OVCAR-8 | 1.7E-06 | 1.9E-07 | 9.2E-08 | 1.5E-07 | 8.6E-09 | 9 | 18 | 11 | 193 |
| NCI-H522 | 1.7E-06 | 2.6E-07 | 5.6E-08 | 1.5E-07 | 1.2E-08 | 6 | 31 | 12 | 137 |
| HCT-15 | 1.7E-06 | 2.6E-07 | 2.4E-07 | 2.5E-07 | 2.2E-08 | 7 | 7 | 7 | 79 |
| 786-0 | 1.7E-06 | 1.6E-07 | 1.1E-07 | 2.0E-07 | 1.6E-08 | 11 | 16 | 9 | 106 |
| MDA-MB-435 | 1.8E-06 | 2.1E-07 | 1.5E-07 | 8.4E-07 | 2.7E-08 | 8 | 12 | 2 | 65 |
| SW-620 | 1.9E-06 | 2.2E-07 | 1.3E-07 | 2.6E-07 | 5.8E-08 | 9 | 14 | 7 | 33 |
| T-47D | 1.9E-06 | 7.0E-07 | 2.3E-07 | 2.9E-07 | 5.0E-08 | 3 | 8 | 7 | 38 |
| EKVX | 2.0E-06 | 3.2E-07 | 2.1E-07 | 2.3E-07 | 1.8E-08 | 6 | 9 | 8 | 107 |
| A549/ATCC | 2.0E-06 | 1.5E-07 | 9.0E-08 | 1.8E-07 | 1.2E-08 | 13 | 22 | 11 | 166 |
| UACC-257 | 2.0E-06 | 6.3E-07 | 2.7E-07 | 5.0E-07 | 2.4E-08 | 3 | 7 | 4 | 83 |
| KM12 | 2.1E-06 | 2.7E-07 | 1.3E-07 | 1.1E-06 | 4.4E-08 | 8 | 16 | 2 | 47 |
| OVCAR-3 | 2.1E-06 | 2.4E-07 | 1.4E-07 | 4.9E-07 | 5.3E-08 | 9 | 15 | 4 | 40 |
| SK-OV-3 | 2.2E-06 | 2.6E-07 | 7.2E-08 | 3.4E-08 | 1.2E-08 | 8 | 30 | 64 | 173 |
| SNB-19 | 2.2E-06 | 2.5E-07 | 9.1E-08 | 2.2E-07 | 1.6E-08 | 9 | 24 | 10 | 135 |
| IGROV1 | 2.3E-06 | 5.0E-07 | 2.0E-07 | 1.5E-06 | 5.8E-08 | 5 | 12 | 2 | 40 |
| OVCAR-4 | 2.3E-06 | 4.6E-07 | 1.9E-07 | 1.6E-06 | 1.6E-07 | 5 | 12 | 1 | 15 |
| HS 578T | 2.4E-06 | 1.4E-06 | 2.9E-07 | 2.3E-06 | 1.6E-07 | 2 | 8 | 1 | 15 |
| HOP-92 | 2.5E-06 | 3.5E-07 | 1.8E-07 | 2.4E-07 | 2.7E-08 | 7 | 14 | 11 | 94 |
| A498 | 2.8E-06 | 2.5E-07 | 2.2E-07 | 1.2E-06 | 1.3E-07 | 11 | 13 | 2 | 22 |
| TK-10 | 2.9E-06 | 3.4E-06 | 1.1E-06 | 1.3E-06 | 1.3E-07 | 1 | 3 | 2 | 22 |
| PC-3 | 3.1E-06 | 7.2E-07 | 4.8E-07 | 3.6E-07 | 2.5E-08 | 4 | 7 | 9 | 128 |
| SK-MEL-28 | 3.6E-06 | 2.9E-07 | 1.6E-07 | 2.6E-07 | 1.5E-08 | 12 | 23 | 14 | 247 |
| OVCAR-5 | 5.1E-06 | 3.1E-07 | 2.6E-07 | 2.8E-07 | 3.6E-08 | 16 | 20 | 18 | 141 |

**Table S3: NCI-60 cancer cell screening.** 55 standard cancer cell lines, tested for growth inhibition (GI) caused by **-CBI-TMI** redox prodrugs (controlled by the **P-CC60-** hydrolytic prodrug), screened by the National Cancer Institute (NCI) developmental therapeutics program (DTP). Cell lines were sorted by increasing $GI_{50}$ values for **P-CC60-CBI-TMI**. Final four columns plot the reductive indices, i.e. potencies referenced against **P-CC60-CBI-TMI**, to reveal qualitatively where reductive activation occurs at significantly higher levels than hydrolysis.

| Cell line | *NCI-60* IC$_{50}$ [M] P-CC60 | *ProliFiler* GI$_{50}$ [M] P-CC60 | GI$_{50}$/IC$_{50}$ | *NCI-60* IC$_{50}$ [M] P-SS60 | *ProliFiler* GI$_{50}$ [M] P-SS60 | GI$_{50}$/IC$_{50}$ | *NCI-60* IC$_{50}$ [M] P-SS66C | *ProliFiler* GI$_{50}$ [M] P-SS66C | GI$_{50}$/IC$_{50}$ |
|---|---|---|---|---|---|---|---|---|---|
| MCF7 | 2.1E-07 | 3.2E-07 | 1.5 | 4.5E-08 | 3.5E-08 | 0.8 | 3.1E-08 | 1.5E-08 | 0.5 |
| DU-145 | 1.8E-07 | 4.6E-07 | 2.5 | 3.6E-08 | 5.9E-08 | 1.6 | 1.6E-08 | 2.5E-08 | 1.5 |
| MOLT-4 | 7.6E-08 | 5.9E-07 | 7.8 | 1.3E-08 | 9.0E-08 | 6.9 | 8.6E-09 | 6.3E-08 | 7.3 |
| MDA-MB-468 | 1.5E-07 | 6.9E-07 | 4.6 | 3.1E-08 | 1.7E-07 | 5.4 | 1.9E-08 | 3.7E-08 | 2.0 |
| HL-60 | 1.0E-07 | 7.5E-07 | 7.4 | 1.8E-08 | 1.2E-07 | 6.5 | 1.5E-08 | 4.8E-08 | 3.2 |
| COLO 205 | 7.6E-07 | 1.4E-06 | 1.8 | 1.6E-07 | 3.4E-07 | 2.2 | 8.6E-08 | 1.5E-07 | 1.7 |
| RPMI-8226 | 3.4E-07 | 1.5E-06 | 4.4 | 3.9E-08 | 2.2E-07 | 5.8 | 1.2E-08 | 1.4E-07 | 12.0 |
| CAKI-1 | 3.3E-07 | 1.5E-06 | 4.5 | 6.3E-08 | 2.7E-07 | 4.3 | 6.3E-08 | 1.7E-07 | 2.7 |
| HCT-116 | 4.4E-07 | 1.5E-06 | 3.4 | 6.9E-08 | 1.6E-07 | 2.3 | 3.2E-08 | 1.5E-07 | 4.8 |
| HT29 | 1.0E-06 | 1.5E-06 | 1.5 | 1.9E-07 | 2.7E-07 | 1.4 | 1.1E-07 | 1.4E-07 | 1.3 |
| HCT-15 | 5.5E-07 | 1.7E-06 | 3.2 | 1.0E-07 | 2.6E-07 | 2.5 | 1.4E-07 | 2.4E-07 | 1.7 |
| 786-0 | 1.2E-07 | 1.7E-06 | 14.7 | 2.3E-08 | 1.6E-07 | 7.3 | 2.9E-08 | 1.1E-07 | 3.7 |
| SW-620 | 7.0E-07 | 1.9E-06 | 2.7 | 1.4E-07 | 2.2E-07 | 1.6 | 1.5E-07 | 1.3E-07 | 0.9 |
| T-47D | 1.0E-06 | 1.9E-06 | 1.8 | 2.3E-07 | 7.0E-07 | 3.0 | 9.8E-08 | 2.3E-07 | 2.3 |
| A549 | 7.7E-07 | 2.0E-06 | 2.6 | 1.2E-07 | 1.5E-07 | 1.3 | 3.0E-08 | 9.0E-08 | 3.0 |
| OVCAR-3 | 1.0E-06 | 2.1E-06 | 2.1 | 1.2E-07 | 2.4E-07 | 2.0 | 7.9E-08 | 1.4E-07 | 1.8 |
| SK-OV-3 | 4.4E-07 | 2.2E-06 | 4.9 | 6.2E-08 | 2.6E-07 | 4.2 | 3.7E-08 | 7.2E-08 | 1.9 |
| A498 | 3.0E-07 | 2.8E-06 | 9.2 | 5.0E-08 | 2.5E-07 | 5.0 | 9.4E-08 | 2.2E-07 | 2.3 |
| PC-3 | 5.2E-07 | 3.1E-06 | 6.1 | 5.8E-08 | 7.2E-07 | 12.4 | 4.0E-08 | 4.8E-07 | 11.9 |
| **Mean** | | | **4.6** | | | **4.0** | | | **3.5** |
| **SD** | | | **3.2** | | | **2.8** | | | **3.3** |

| **Reductive activation index** Cell line | *NCI-60* P-SS60 | *ProliFiler* P-SS60 | ratio | *NCI-60* P-SS66C | *ProliFiler* P-SS66C | ratio |
|---|---|---|---|---|---|---|
| MCF7 | 9 | 6.1 | 1.5 | 21 | 7.0 | 3.0 |
| DU-145 | 8 | 3.1 | 2.5 | 19 | 11.3 | 1.6 |
| MOLT-4 | 7 | 0.8 | 7.8 | 9 | 8.8 | 1.1 |
| MDA-MB-468 | 4 | 0.9 | 4.6 | 18 | 7.9 | 2.3 |
| HL-60 | 6 | 0.9 | 7.4 | 16 | 6.7 | 2.3 |
| COLO 205 | 4 | 2.2 | 1.8 | 9 | 8.8 | 1.1 |
| RPMI-8226 | 7 | 1.5 | 4.4 | 10 | 28.5 | 0.4 |
| CAKI-1 | 5 | 1.2 | 4.5 | 9 | 5.2 | 1.7 |
| HCT-116 | 9 | 2.8 | 3.4 | 10 | 13.6 | 0.7 |
| HT29 | 6 | 3.7 | 1.5 | 11 | 9.5 | 1.1 |
| HCT-15 | 7 | 2.1 | 3.2 | 7 | 3.9 | 1.8 |
| 786-0 | 11 | 0.7 | 14.7 | 16 | 4.1 | 4.0 |
| SW-620 | 9 | 3.2 | 2.7 | 14 | 4.6 | 3.1 |
| T-47D | 3 | 1.5 | 1.8 | 8 | 10.7 | 0.8 |
| A549 | 13 | 5.1 | 2.6 | 22 | 25.4 | 0.9 |
| OVCAR-3 | 9 | 4.3 | 2.1 | 15 | 12.9 | 1.1 |
| SK-OV-3 | 8 | 1.7 | 4.9 | 30 | 11.8 | 2.5 |
| A498 | 11 | 1.2 | 9.2 | 13 | 3.2 | 4.0 |
| PC-3 | 4 | 0.7 | 6.1 | 7 | 12.9 | 0.5 |
| **Mean** | | | **4.6** | | | **1.8** |
| **SD** | | | **3.2** | | | **1.1** |

**Table S4:** Comparison of absolute potencies and "Reductive Activation Indices" in the 19 overlapping cell lines between NCI-60 and ProliFiler-140 screens.

### 7. *in vivo* studies

#### *7.1. in vivo PK and prodrug tolerance (mouse studies)*

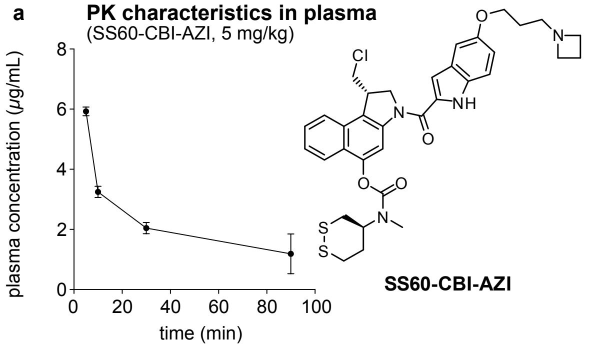


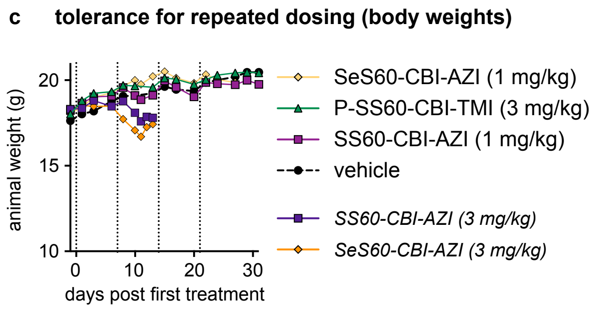


**b signs of toxicity, repeated dosing**

| | vehicle | | SS60-CBI-AZI | | | SeS60-CBI-AZI | | | O56-CBI-AZI | | |
|---|---|---|---|---|---|---|---|---|---|---|---|
| | mean | SD | mean | SD | p(t) | mean | SD | p(t) | mean | SD | p(t) |
| BW (body weight) | 19.4 | 1.3 | 19.7 | 0.8 | 0.586 | 19.5 | 1.5 | 0.59 | 19.4 | 1.25 | 0.47 |
| LW (liver weight) | 40.9 | 2.0 | 46.1 | 1.3 | **0.0001** | 46.6 | 2.8 | **0.001** | 42.9 | 4.5 | 0.301 |
| ALT/AAT | 28.6 | 5.1 | 33.7 | 7.1 | 0.147 | 40.5 | 11.4 | **0.03** | 30.9 | 2.9 | 0.312 |
| ALP | 110 | 7.5 | 93 | 6.4 | **0.001** | 91 | 17.5 | **0.022** | 97 | 7.0 | **0.006** |
| HGB/HB | 141 | 3.8 | 131 | 9.3 | **0.025** | 128 | 4.9 | **0.0002** | 132 | 3.1 | **0.0004** |
| HCT | 62.6 | 1.8 | 58.7 | 3.0 | **0.001** | 57.9 | 3.0 | **0.005** | 58.2 | 1.4 | **0.0002** |
| RBC | 10.3 | 0.3 | 9.8 | 0.5 | 0.081 | 9.55 | 0.4 | **0.002** | 9.8 | 0.3 | **0.05** |

**Fig S12:** (**a**) plasma pharmacokinetics of initial (non-solubilised) prodrug **SS60-CBI-AZI** following *i.v.* injection in 100 µL of 20:80 DMSO:saline [3 mice per timepoint; see Section 8.3]. (**b**) investigation of potential toxicity parameters for initial prodrugs [seven mice per group; see Section 8.3]. Dosing regimes: three injections, once per week; **SS60-CBI-AZI**: 0.3 then 0.1 then 0.1 mg/kg; **SeS60-CBI-AZI**: 0.3 then 0.2 then 0.2 mg/kg; **O56-CBI-AZI**: 0.3 then 0.2 then 0.2 mg/kg. (**c**) bodyweights during repeated dosing [3 mice per group; weekly injections in 10:90 DMSO:saline repeated up to up to three times; see Section 8.3]. BW: animal body weight (g). LW: liver weight (mg/g body weight). ALT/AAT: alpha-1 antitrypsin (U/L). ALP: alkaline phosphatase (U/L). HGB/HB: hemoglobin (g/L). HCT: hematocrit (%). RBC: red blood cell count (×$10^{12}$/L). p(t): p value compared to the vehicle group; Mann-Whitney U-test without correction for multiple comparisons.

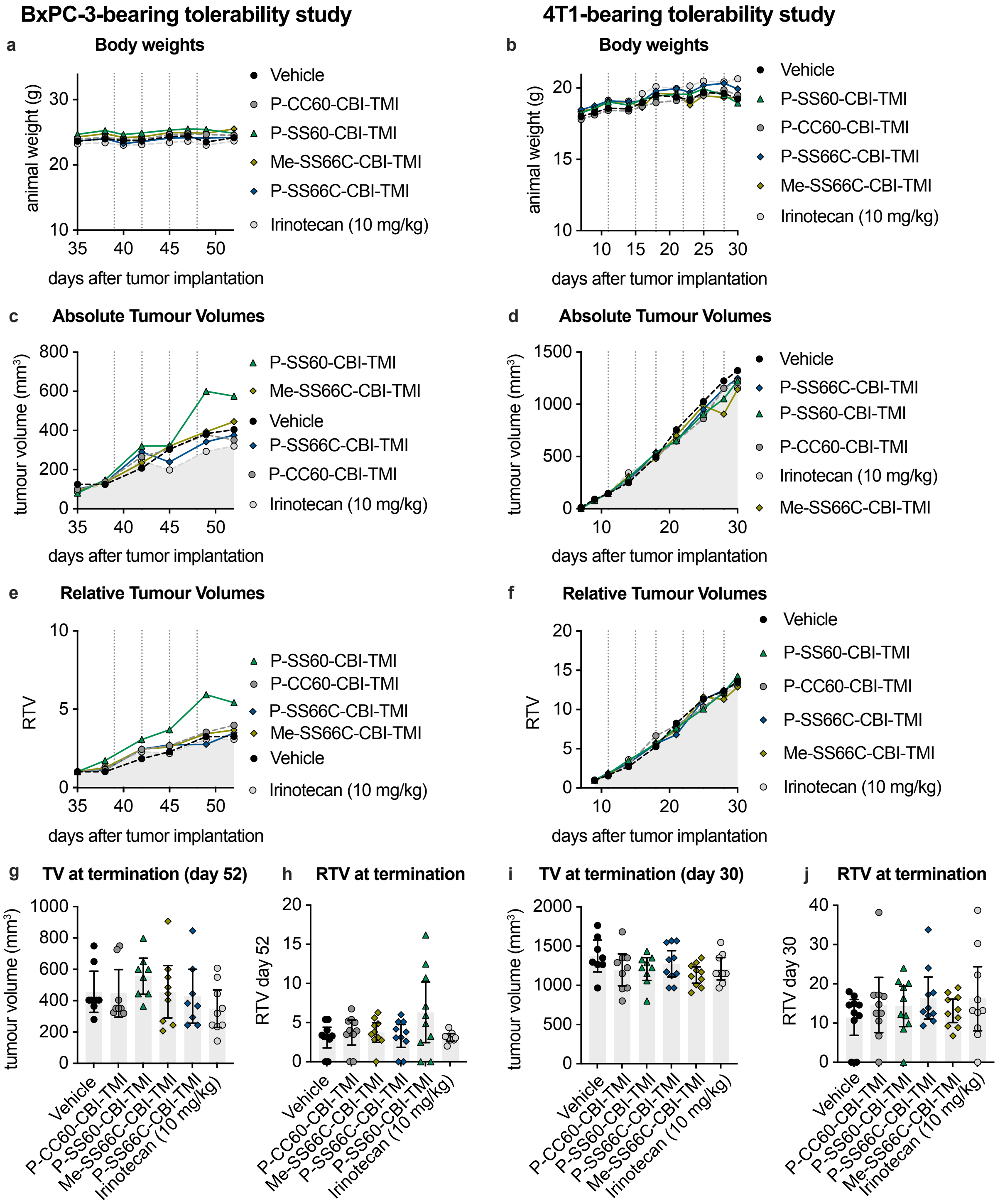


**Fig S13: Repeated dosing studies in tumor-bearing mice.** (**a,c,e,g,h**): BxPC-3 subcutaneous xenograft in NMRi mice, 10 mice per group; (**b,d,f,i,j**): 4T1 murine breast cancer syngeneic orthotopic implant in Balb/c mice, 10 mice per group**.** *Note that reference therapeutic irinotecan failed to slow tumor growth in either model, indicating that these studies are not to be interpreted as efficacy assays but only for tolerability.* (**a,b**) Bodyweights (means); (**c,d**) absolute tumor volumes (medians); (**e,f**) tumor volumes relative to individual volumes at start of treatment (medians); (**g,h,i,j**) absolute and relative tumor volumes at final day of study (datapoints with means and 95% confidence interval). All tumor measurements are *in vivo* caliper measurements. Dotted vertical lines indicate days of treatment. All CBI groups dosed at 3 mg/kg *i.v.,* ca. 100 µL injection volume in vehicle (10:90 DMSO:saline).

#### *7.2. in vivo efficacy mouse studies*

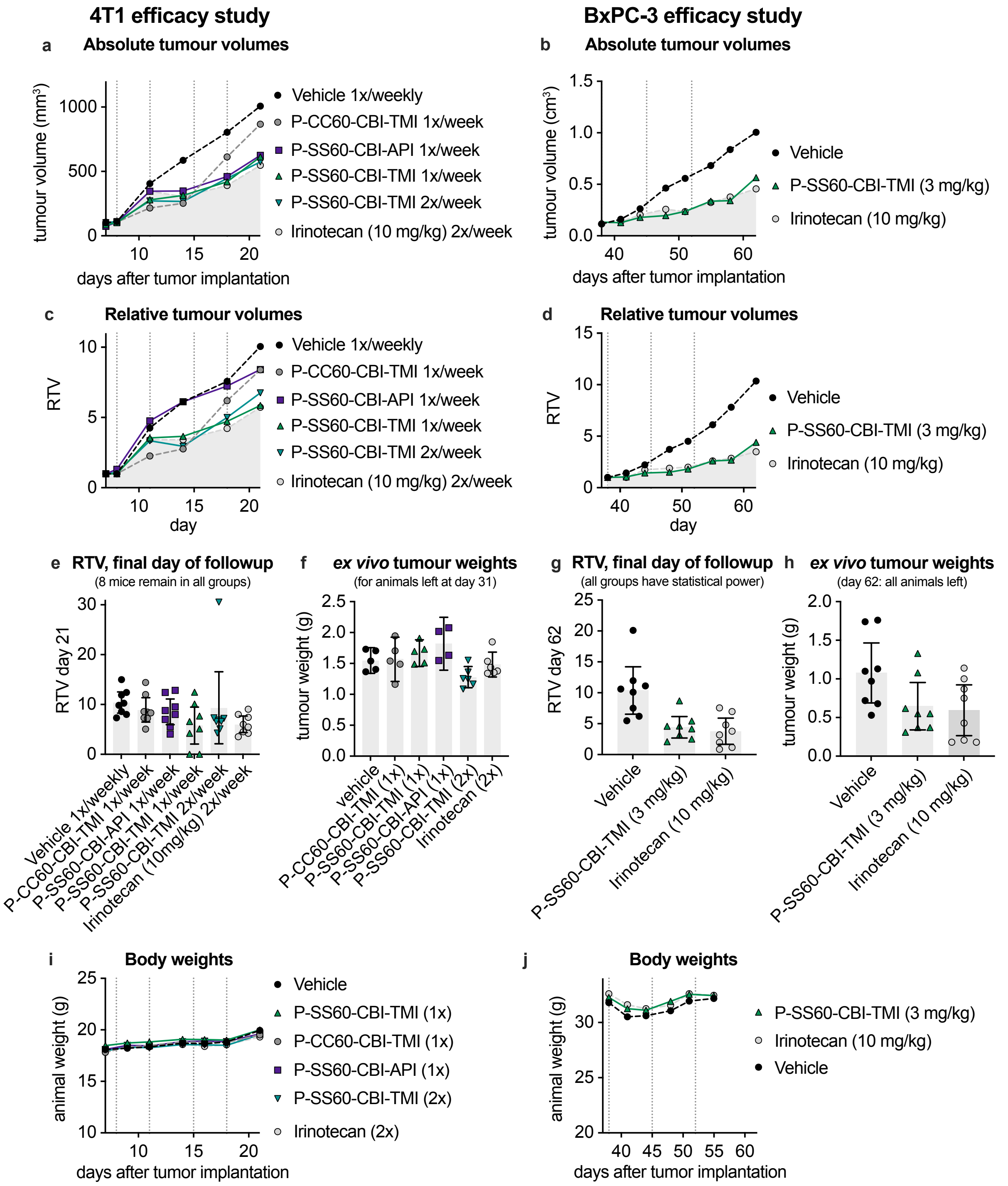


**Fig S14: Efficacy studies in tumor-bearing mice.** (**a,c,e,f,i**): 4T1 murine breast cancer syngeneic orthotopic implant in Balb/c mice, 8 mice per group; (**b,d,g,h,j**): BxPC-3 subcutaneous xenograft in NMRi mice, 8 mice per group. (**a,b**) absolute tumor volumes (median); (**c,d**) tumor volumes relative to individual volumes at start of treatment (median); (**e,g**) *in vivo* caliper measurement of relative tumor volumes at final day of followup [last day when all mice in all groups remain in the study] (datapoints with means and 95% confidence interval); (**f,h**) *ex vivo* tumor weights at termination of study (only mice not terminated on ethical grounds before the indicated days were harvested); (**i,j**) Bodyweights (means). All tumor measurements are *in vivo* caliper measurements except panels f and h. All CBI groups dosed at 3 mg/kg *i.v.* either once or twice weekly as annotated (4T1 study), or once weekly (BxPC-3 study); dotted vertical lines indicate days of [potential] treatment.

### 8. General methods

#### 8.1. *Cell-free HPLC protocols*

HPLC studies were conducted on an *Agilent 1200 SL* system *Agilent Technologies Corp.,* Santa Clara (USA) equipped with a DAD detector, a *Hypersil Gold* HPLC column from *ThermoFisher Scientific GmbH*, Dreieich (Germany) and consecutive low-resolution mass detection using a LC/MSD IQ mass spectrometer applying ESI from *Agilent Technologies Corp.,* Santa Clara (USA). The mobile phase consists of eluent A: water (analytical grade, 0.1% formic acid) and eluent B: MeCN (analytical grade, 0.1% formic acid). The analysis was performed using a gradient of 10% B to 90% B over 10 min.

Reduction assay using TCEP and GSH

10 µL of a solution of the prodrug (1 mM in DMSO) was added to 80 µL of TE-buffer (50 mM Tris-HCl, 20 mM EDTA, pH = 8) in HPLC vials with micro-insert, followed by addition of either 10 µL of a solution of tris(2-carboxyethyl)phospine) (TCEP, 5 mM in TE-buffer), or reduced glutathione (GSH, 50 mM in TE-buffer) or else blank TE-buffer. Final concentrations: Prodrug (0.1 mM), TCEP (0.5 mM), GSH (5 mM). The blank sample was measured after 10 min ("pure") and 24 h ("$OH^-$") to control for basic hydrolysis. The GSH sample was measured after 24 h to control for monothiol-based reductive release. The TCEP sample was measured at several timepoints (e.g. 10 min, 1 h, 2.5 h, 4 h, 7 h) to control for release kinetics of irreversible reduction-based activation. Individual species were identified using ESI-MS analysis. Separate UV/Vis spectra and mass spectra were extracted for each of the intermediate species and final release products. Peak areas were calculated relative to the sum of all peak areas at 320 nm to quantify individual species at each timepoints. As UV/Vis absorption at 320 nm is not equal for all species (especially CBI*-TMI/CBI*-AZI have slightly different absorption profiles) we expect the accuracy of this calculation within an error of ±5%, which is sufficient for the purpose of this experiment.

#### 8.2. *Cell culture methods*

General cell cultivation

HeLa, A549 or MEF cells were cultured in Dulbecco's modified Eagle's medium (DMEM: L-glucose (1 g/L), *L*-pyruvate, phenol-red, $NaHCO_3$ (2.7 g/L), purchased from *Merck GmbH*, Darmstadt, Germany), which was supplemented with 10% heat-inactivated fetal bovine serum, penicillin (100 U/mL), streptomycin (100 µg/mL), *L*-glutamine and $Na_2SeO_3$ (100 nM). Dulbecco PBS buffer (*Merck GmbH*, Darmstadt, Germany) was used for washing and resuspension steps, TrypLE™ Express (*gibco Life Technologies Inc.*, Massachusetts, USA) was used for trypsination. Centrifugation steps were performed for 5 min at a speed of 500 × g, by using an *Eppendorf centrifuge 5810R* (*Eppendorf GmbH*, Hamburg, Germany). Cells were maintained at 37 °C under 5% $CO_2$ and cell growth and morphology was monitored using a *Nikon Eclipse Ti* microscope from *Nikon Corp*., Minato (Japan).

HeLa cells used in this study were HeLa-HRE (14-118ACL) cells, purchased from *abeomics Inc.* (Antibodies & Engineered Cell Lines™, San Diego, USA). Briefly, HeLa-HRE is a stably transfected HeLa cell line that expresses the renilla luciferase reporter gene under the transcriptional control of the hypoxia response element (HRE), which can be used in reporter studies of hypoxic response. A549 cells (DSMZ ACC107) were purchased from the German Collection of Microorganisms and Cell Cultures. TrxR1 knockout (*$TrxR^{-/-}$* or *TrxR-ko*) and reference (*$TrxR^{fl/fl}$* or *TrxR-wt*) mouse embryonic fibroblasts (MEF) are a kind gift from Marcus Conrad. Briefly, MEFs isolated from conditional TrxR1 knockout mouse embryos, were immortalised by lentiviral transduction. *In vitro* deletion of TrxR1 was achieved by Tat-Cre induced recombination and verified by PCR and Immunoblotting for TrxR1.[5] All cell lines are tested regularly for mycoplasma contamination and only mycoplasma negative cells were used in assays.

Cellular toxicity assays – resazurin-based cell viability assay

For evaluation of cellular toxicity, Hela-HRE, A549 or MEF cells were seeded in 96-well plates (Microplates, 96-well; F-bottom, transparent, *Greiner bio-one GmbH*, Kremsmünster, Austria), with a density of $10^4$ cells/well and in 100 µL (HeLa-HRE, MEF) or 150 µL (A549) of medium. Cells were allowed to attach over night before compound treatment was initiated the next morning. In general, compounds were applied from DMSO stock solutions, with a maximum final DMSO level of 2% per well and covering concentration ranges from 0.01 nM up to 30 µM. All compounds were measured at least three times in independent experiments.

After treatment, cell plates were incubated at 37 °C under 5% $CO_2$ atmosphere for 48 h (HeLa-HRE) or 72 h (A549, MEF), before a resazurin solution (150 mg/L resazurin sodium salt in PBS) was added. Resazurin was left on cells for 4 h at 37 °C and 5% $CO_2$, before fluorescence of its metabolite resorufin was measured at 590 nm (excitation 544 nm) using a *FLUOstar Omega* microplate reader (*BMG Labtech GmbH*, Ortenburg, Germany). Data were normalized to data from DMSO treated control cells, as "100% viability", wherease signal from wells without cells served as "0% viability" control. Three different concentrations of the linear disulfide-based control compound **$SS00_M$-CBI-AZI** served as interrun controls on every plate. A minimum of three experiments was performed per compound. Analysis and visualization were performed in *GraphPad Prism 8.4.2* using a nonlinear regression curve (log (inhibitor) vs. response – variable solpe, four parameter) to calculate "cellular $IC_{50}$ values".

##### 8.3. *Methods for* in vivo *animal studies*

All *in vivo* studies were conducted as contract research together with *ReactionBiology Europe GmbH* (Freiburg, Germany), *EPO – Experimental Pharmacology & Oncology GmbH* (Berlin, Germany); *Bienta/Enamine Ltd.* (Kyiv, Ukraine) and *Leitat – Managing Technologies* (Barcelona, Spain), adhering to the respective animal welfare protocols.

Mice were kept in individually ventilated cages at constant temperature (22±2 °C, air-conditioned) and humidity (45 – 65%), under optimum hygienic conditions with 10 – 15 air changes per hour. A cycle of 12 h artificial fluorescent lightning and 12 h darkness was applied, and animal behavior was monitored daily throughout the studies. The animals received food and water *ad libitum*.

Toxicity studies

To evaluate toxicity of selected compounds, dose level selection studies within dose range from 0.1 mg/kg to 10 mg/kg were performed by intravenous administration into Balb/c mice. Each dose for each tested compound was administered in one mouse. After drug administration, the animals were observed for mortality, body weight changes and clinical signs of gross toxicity three times per week for 21 consecutive days of post-dosing period. After the first week of observation without adverse events, two doses for each test compound were selected as tolerated. These two doses were tested in two additional mice for confirmation of tolerability. Additional animals were observed for mortality, body weight changes, and clinical signs of gross toxicity three times per week for 21 consecutive days of post-dosing period. Toxicity of repeated administrations was tested with three repeated pre-selected doses at a dosing interval of 7 days. Mortality, body weight changes, and clinical signs of gross toxicity were monitored after each drug administration and three times per week for 21 consecutive days after the first compound administration. Bleeding with subsequent hematological tests, clinical chemistry parameters analysis (ALT, AST, ALP, LDH, creatine kinase, creatinine, urea, cholesterol, and triglycerides), and necropsy with macroscopic inspection were performed on the terminal day of the repeated dosing-regime.

Pharmacokinetics of **SS60-CBI-AZI**

Following single dose intravenous administration, levels of **SS60-CBI-AZI** in blood plasma over time were determined by HPLC-MS/MS. Treatment groups were three Balb/c mice (randomly assigned) per time point (total: four termination time points, at 5, 10, 30 and 90 min). Blood collection was performed from the orbital sinus in microtainers containing $K_2$EDTA. All samples were immediately processed, flash-frozen and stored at -70 °C until analysis. Briefly, plasma samples (50 µL) were mixed with 200 µL of internal standard (**IS**) solution (IS-7639, 400 ng/mL in ethanol). After mixing by pipetting and centrifuging for 4 min at 6000 rpm, 2 µL of each supernatant was injected into LC-MS/MS (Shimadzu LC: 2 isocratic pumps LC-10ADvp, autosampler SIL-20AC, sub-controller FCV-14AH, degasser DGU-14A; AB Sciex MS: API 3000 (triple-quadrupole), ESI). The data acquisition and system control was performed using *Analyst 1.5.2 software* (*AB Sciex Ltd.*, Canada).

***Chromatographic Conditions:***
*Column: Discovery HS C18 (50 x 2.1 mm, 5 µm)*
*Mobile phase A: Acetonitrile : Water : Formic acid = 50 : 950 : 1*
*Mobile phase B: Acetonitrile : Formic acid = 100 : 0.1*
*Linear gradient: 0 min 5 % B, 1.1 min 100% B, 1.11 min 5 % B, 2.5 min stop.*
*Elution rate: 400 µL/min. A divert valve directed the flow to the detector from 1.2 to 2.3 min.*
*Column temperature: 30 °C*

***MS/MS Detection:***
*Scan type: Positive MRM, Ion source: Turbo spray, Ionization mode: ESI*
*Nebulizer gas: 15 L/min, Curtain gas: 8 L/min, Collision gas: 4 L/min.*
*Ion spray voltage: 5000 V, Temperature: 400 °C*

Tolerability studies

To test tolerability of selected compounds *in vivo*, female BALB/c mice were randomized by weight into seven groups of three animals. On days 0 and 7, the first 3 groups were treated with 3 mg/kg of either **SS60-CBI-AZI, SeS60-CBI-AZI** or **P-SS60-CBI-TMI** and mice were inspected 30 min, 1 h, 2 h, 3 h and 4 h after the first treatment. If a treatment was well tolerated, it was taken into a confirmation cycle with another set of three mice; if a treatment was not tolerated, the applied dose was reduced and tested again in three mice. **P-SS60-CBI-TMI** was tolerated at 3 mg/kg over two weeks and 4 treatments, which was further confirmed by another group of 3 animals. Severe adverse effects (weight loss and signs of inflammation at injection site) were observed for both groups treated with 3 mg/kg **SS60-CBI-AZI** or **SeS60-CBI-AZI**, which led to a dose reduction to 1 mg/kg in a second cohort of each 3 mice (treated twice a week). 1 mg/kg **SS60-CBI-AZI** was tolerated in two groups of each three mice with only mild adverse effects, while **SeS60-CBI-AZI** still caused adverse effects leading to discontinuation of this compound.

Efficacy studies

***Syngeneic breast cancer model: Balb/c:***

4T1 murine breast cancer cells were cultured in RPMI-1640 high Glutamax1 with 10 % fetal calf serum, 100 units penicillin/mL and 100 µg streptomycin/mL, until they reach a confluency of 70 – 90 %, before being split routinely using trypsin/EDTA. 100 µL $10^5$ 4T1 tumor cell suspension was orthotopically injected into the left mammary fat pad of each mouse.

***Pancreatic cancer xenograft model in NMRI:nu/nu or athymic nude (crl: NU(NCr)-Foxn1$^{nu}$):***

BxPC-3 human pancreas carcinoma cells were cultured in RPMI-1640 high Glutamax1 with 10 % fetal calf serum, 100 units penicillin/mL and 100 µg streptomycin/mL, until they reach a confluency of 70 – 90 %, before being split routinely using trypsin/EDTA. 100 µL 5 x $10^6$ BxPC-3 tumor cell suspension was subcutaneously injected into the left flank of each mouse.

Mice were randomized after a mean tumor volume of ~100 $mm^3$ was reached, and treatment was started on the same day. In general, compounds were dissolved in DMSO to obtain a 10× stock solution, from which all further working solutions in 0.9% NaCl with a final concentration of 1% DMSO were prepared freshly. Treatment was performed in the indicated dosages, once or twice weekly, by intravenous injection. After start of the therapy, animal weight was determined three times weekly and tumor monitoring twice weekly by calipering. Deviation of the health status of the animals were documented and animals were euthanized individually before study termination when ethical abortion criteria were reached (e.g., body weight loss ≥ 20%, signs of sickness; after s.c. implantation: tumor diameter ≥ 2 cm, tumor ulceration). Single groups were terminated early if four animals of tht group reached ethical abortion criteria.

#### *8.4. Synthetic techniques*

All solvents, reagents and building blocks were purchased from standard commercial sources. Anhydrous solvents obtained in septum-capped bottles and analytical grade, or higher quality solvents were used without purification. Industrial grade solvents were distilled prior to use. Unless otherwise stated, reactions were performed at room temperature without precautions regarding air or moisture and were stirred using a magnetic Teflon®-coated stir bars. Air or moisture sensitive reactions were conducted in dry Schlenk glassware.

The term "column chromatography" refers to either (a) manual flash column chromatography conducted under positive nitrogen pressure using *Ceduran® Si 60* silica gel (40−63 µm) from *Merck GmbH* as stationary phase, or (b) purification on the automated column chromatography system *Biotage® Select* using prepacked silica cartridges purchased from *Biotage GmbH*, Uppsala (Sweden). All eluent and solvent mixtures are given as volume ratios unless otherwise specified. Unless stated otherwise, and thin layer chromatography (TLC) to monitor reactions and determine $R_f$ values was performed on silica coated aluminium sheets with fluorescent indicator (*TLC Silica gel 60 F254* from *Merck GmbH*, Darmstadt, Germany) with visualisation by UV irradiation (254 nm/360 nm) or staining with $KMnO_4$ solution (3.0 g $KMnO_4$, 20 g $K_2CO_3$, 0.30 g KOH, 0.30 L $H_2O$).

**Analytical methods for chemical synthesis**

High resolution mass spectrometry (**HRMS**) was conducted either using a *Thermo Finnigan LTQ FT Ultra FourierTransform* ion cyclotron resonance spectrometer from *ThermoFisher Scientific GmbH*, Dreieich (Germany) applying electron spray ionisation (ESI) with a spray capillary voltage of 4 kV at temperature 250 °C with a method dependent range from 50 to 2000 u or a *Finnigan MAT 95* from *Thermo Fisher Scientific*, Dreieich (Germany) applying electron ionisation (EI) at a source temperature of 250 °C and an electron energy of 70 eV with a method dependent range from 40 to 1040 u. All reported *m/z* values refer to positive ionization mode, unless stated otherwise.

Nuclear magnetic resonance (**NMR**) spectroscopy was performed using a *Bruker Avance* (600/150 MHz, with *TCI cryoprobe) or a Bruker Avance III HD Biospin* (400/100 MHz, with BBFO cryoprobeTM) from Bruker Corp., Billerica (USA) either at 400 MHz or 500 MHz. $^{19}F$-NMR spectra were recorded on a *Bruker Avance III* spectrometer (400 MHz for $^1H$; 377 MHz for $^{19}F$). NMR-spectra were measured at 298 K, unless stated otherwise, and were analysed with the program *MestreNova 12* developed by *MestreLab Ltd.*, Santiago de Compostela (Spain). $^1H$-NMR spectra chemical shifts (δ) in parts per million (ppm) relative to tetramethylsilane (δ = 0 ppm) are reported using the residual protic solvent ($CHCl_3$ in $CDCl_3$: δ = 7.26 ppm, DMSO-$d_5$ in DMSO-$d_6$: δ = 2.50 ppm, $CHD_2OD$ in $CD_3OD$: δ = 3.31 ppm) as an internal reference. For $^{13}C$-NMR spectra, chemical shifts in ppm relative to tetramethylsilane (δ = 0 ppm) are reported using the central resonance of the solvent signal ($CDCl_3$: δ = 77.16 ppm, DMSO-$d_6$: δ = 39.52 ppm, $CD_3OD$: δ = 49.00 ppm) as an internal reference. For $^1H$-NMR spectra in addition to the chemical shift the following data is reported in parenthesis: multiplicity, coupling constant(s), number of hydrogen atoms and, if available, assignment. The abbreviations for multiplicities and related descriptors are s = singlet, d = doublet, t = triplet, q = quartet, p = pentuplet, hept = heptet or combinations thereof, m = multiplet and br = broad. The numbering scheme used for the assignments is specified in each case in a figure depicting the respective molecular structure and does not follow any convention. The reported assignments are supported by 2D-NMR experiments (HMBC, HSQC, COSY). Where known products matched literature analysis data, only selected data acquired are reported.

Analytical high-performance liquid chromatography (**HPLC**) analysis was conducted either using an *Agilent 1100* system from *Agilent Technologies Corp.,* Santa Clara (USA) equipped with a DAD detector and a *Hypersil Gold* HPLC column from *ThermoFisher Scientific GmbH*, Dreieich (Germany) or a *Agilent 1200 SL* system *Agilent Technologies Corp.,* Santa Clara (USA) equipped with a DAD detector, a *Hypersil Gold* HPLC column from *ThermoFisher Scientific GmbH*, Dreieich (Germany) and consecutive low-resolution mass detection using a LC/MSD IQ mass spectrometer applying ESI from *Agilent Technologies Corp.,* Santa Clara (USA). For both systems mixtures of water (analytical grade, 0.1% formic acid) and MeCN (analytical grade, 0.1% formic acid) were used as eluent systems.

Single crystal **X-Ray** crystallography was conducted using a *Bruker D8 Venture TXS system* from *Bruker Corp*., Billerica (USA) equipped with a multilayer mirror monochromator and a Mo Kα rotating anode X-ray tube ($\lambda$ = 0.71073 Å). The frames were integrated with the *Bruker SAINT software package* (Bruker, 2012. SAINT. Bruker AXS Inc., Madison, Wisconsing, USA). Data were corrected for absorption effects using the Multi-Scan method (SADABS) (Sheldrick, G. M., 1996. SADABS. University of Goettingen, Germany). The structures were solved and refined using the *Bruker SHELXTL Software Package*.[6] The figures have been drawn at the 50% ellipsoid probability level.[7] Crystallographic data have been deposited with the *Cambridge Crystallographic Data Centre, CCDC*, 12 Union Road, Cambridge CB21EZ, UK. Copies of the data can be obtained free of charge on quoting the depository number CCDC 2214471 (https://www.ccdc.cam.ac.uk/structures/).

### 9. Chemical synthesis

#### *9.1. General synthetic protocols*

##### BnO-CBI-Boc

*N*-Boc-(5-(benzyloxy)-(*S*)-1-(chloromethyl)-1,2-dihydro-3*H*-benzo[*e*]indole

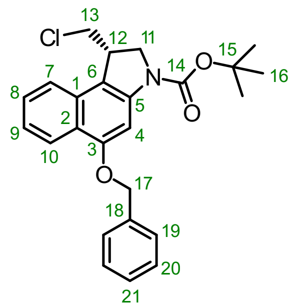


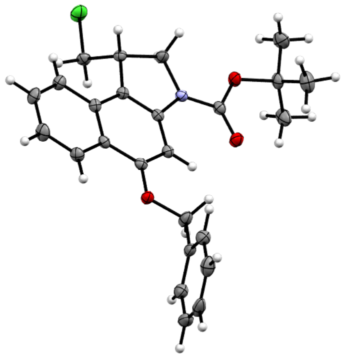

For the synthesis of *seco*-CBI derived prodrugs commercial **BnO-CBI-Boc** was purchased as a colourless, crystalline solid from *ArchBioscience Inc.*, West Chester, PA (USA). The identity and purity of the material was determined by NMR spectroscopy, MS and X-ray crystallography; in particular, the (*S*)-configuration at C1 which is fundamental for high-potency activity was confirmed by determination of the specific rotation[8], and by acquiring an X-Ray crystallography structure (**Table S5**) matching literature[9]. Analytical data matched literature reports.

**TLC** $R_f = 0.38$ (isohexane:EtOAc, 5:1). **HRMS** (ESI$^+$) for $C_{18}H_{21}ClNO_3^+$: calcd. *m/z* 334.12045, found *m/z* 334.12068. **$[\alpha]^{25}_D$** = -15.2 (c = 4.66 g/L, $CH_2Cl_2$). Unambigous confirmation of the absolute configuration was derived from a single-crystal X-Ray analysis. **$^1$H-NMR** (500 MHz, $CDCl_3$): δ (ppm) = 8.30 (dd, *J* = 8.7, 1.2 Hz, 1H, 10-H), 7.88 (br-s, 1H, 4-H), 7.65 (d, *J* = 8.3 Hz, 1H, 7-H), 7.55 (d, *J* = 7.5 Hz, 2H, 19-H), 7.51 (ddd, *J* = 8.2, 6.8, 1.3 Hz, 1H, 9-H), 7.46 – 7.41 (m, 2H, 20-H), 7.39 – 7.36 (m, 1H, 21-H), 7.34 (ddd, *J* = 8.2, 6.8, 1.2 Hz, 1H, 8-H), 5.27 (s, 2H, 17-H), 4.27 (s, 1H, 11-H), 4.20 – 4.10 (m, 1H, 11'-H), 4.05 – 3.89 (m, 2H, 12-H+13-H), 3.44 (t, *J* = 10.4 Hz, 1H, 13'-H), 1.62 (s, 9H, 16-H). **$^{13}$C-NMR** (126 MHz, $CDCl_3$): δ (ppm) = 156.2 (C3), 152.8 (C=O, C14), 142.0 ($C_{Ar}$, C5), 136.9 ($C_{Ar}$, C18), 130.4 ($C_{Ar}$, C2), 128.7 ($C_{Ar}$H, C20), 128.2 ($C_{Ar}$H, C21), 127.8 ($C_{Ar}$H, C9), 127.7 ($C_{Ar}$H, C19), 123.8 ($C_{Ar}$H, C10), 123.3 ($C_{Ar}$H, C8), 122.6 ($C_{Ar}$, C6), 121.9 ($C_{Ar}$H, C7), 114.5 ($C_{Ar}$, C1), 96.6 ($C_{Ar}$, C4), 81.5 (*C*$(CH_3)_3$, C15), 71.7 ($CH_2$, C17), 54.9 ($CH_2$, C13), 48.3 ($CH_2$, C11), 41.3 (CH, C12), 29.2 (C(*C*$H_3)_3$, C16).

##### HO-CBI-Boc

*N*-Boc-(*S*)-1-(chloromethyl)-5-hydroxy-1,2-dihydro-3*H*-benzo[*e*]indole

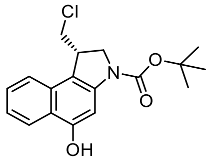


Commercial **BnO-CBI-Boc** (591 mg, 1.39 mmol) was dissolved in THF (75 mL, 0.02 M). The mixture was heated to 35 °C and Pd/C (297 mg, 10% on charcoal) and $NH_4HCO_2$ (1.4 mL of 4 M aq. solution) were added. The mixture was stirred for 70 min at 35 °C until quantitative turnover was confirmed by TLC, was then filtered through Celite and the filter cake was washed with EtOAc (40 mL). The combined filtrates were concentrated under reduced pressure to obtain **HO-CBI-Boc** as a colourless solid (460 mg, 1.38 mmol, 99%). Optional purification by FCC (isohexane/EtOAc) gave **HO-CBI-Boc** as a colourless solid. Analytical data machted literature reports.[10,11]

**TLC** $R_f = 0.59$ (isohexane:EtOAc, 10:1). **HRMS** (ESI$^-$) for $C_{18}H_{21}ClNO_3^-$: calcd. *m/z* 332.10589, found *m/z* 332.10639. **$^1$H-NMR** (400 MHz, $CDCl_3$): δ (ppm) = 8.18 (d, *J* = 8.4 Hz, 1H), 7.74 (s, 1H), 7.63 (d, *J* = 8.4 Hz, 1H), 7.50 (ddd, *J* = 8.3, 6.8, 1.3 Hz, 1H), 7.34 (ddd, *J* = 8.1, 6.8, 1.1 Hz, 1H), 6.32 (s, 1H), 4.24 (d, *J* = 11.9 Hz, 1H), 4.15 – 4.06 (m, 1H), 4.03 – 3.88 (m, 2H), 3.51 – 3.38 (m, 1H), 1.61 (s, 9H).

#### Summary of trigger precursor HCl salt syntheses

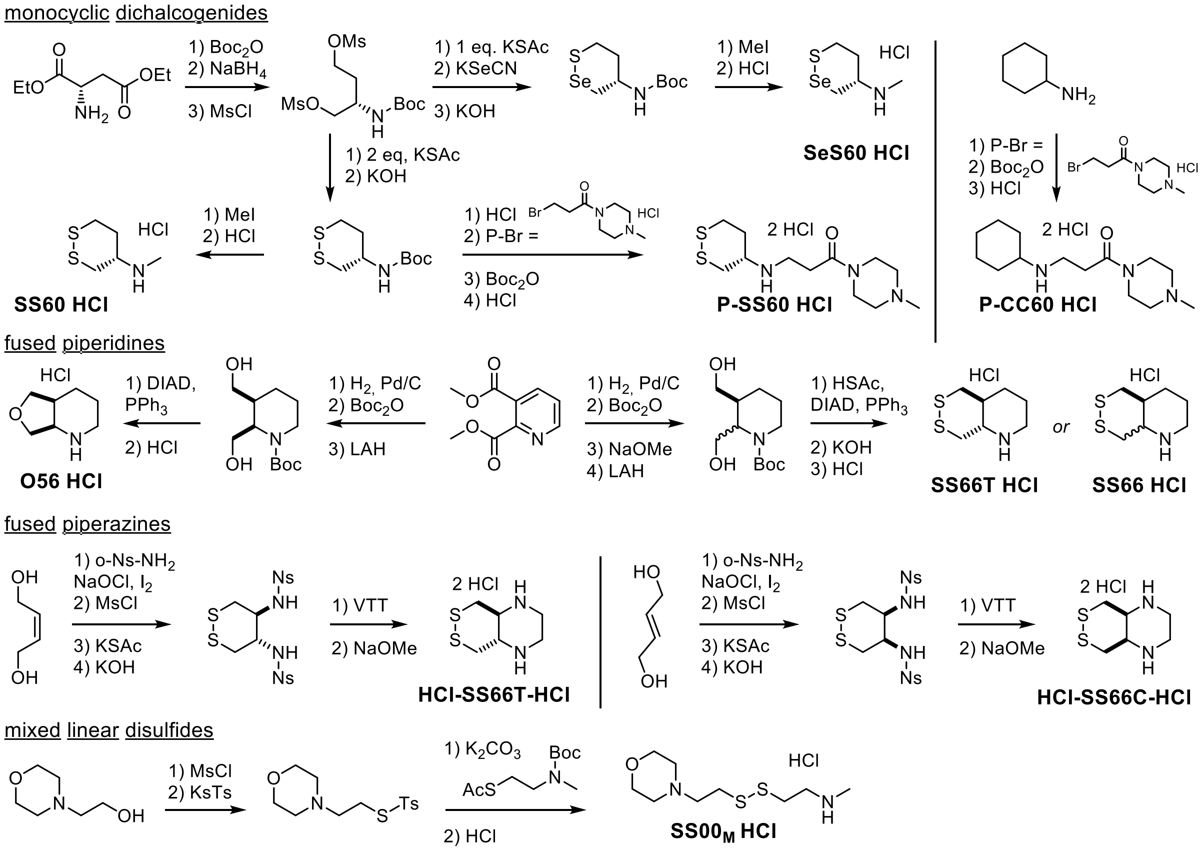


**Scheme 1 Trigger Synthesis.** (**a**) monocyclic dichalcogenides were prepared according to *JACS* **2021**[1], *Chem* **2022**[3] and *ChemRxiv* **2022**[12]; (**b**) fused piperidines according to *JACS* **2021**[1]; (**c**) fused piperazines according to WO 2022/200347[13] and *paper in preparation* **2022**[4]; (**d**) mixed linear disulfides according to *JACS* **2021**.[1]

##### **General protocol A:** Carbamate coupling using triphosgene

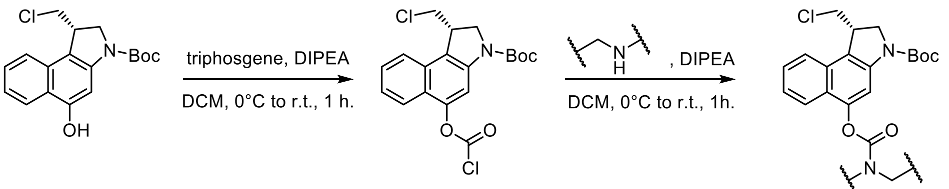


**HO-CBI-Boc** **ClC(O)O-CBI-Boc**

**Step 1: HO-CBI-Boc** (1 eq.) was dissolved in anhydrous DCM (0.1 M) and the resulting solution was cooled to 0 °C. Triphosgene (1.2 eq., 0.2 M solution in anhydrous DCM) and DIPEA (4.0 eq, 0.4 M solution in anhydrous DCM) were subsequently added dropwise. The resulting mixture was stirred at 0 °C for 0.5 h, was then allowed to warm to r.t. and was further stirred for 0.5 h. All volatile compounds were removed under reduced pressure to afford the corresponding chloroformate **ClC(O)O-CBI-Boc** as a solid.
*Note: Residual phosgene was collected using an external liquid nitrogen trap. The trapped mixture was carefeully thawed and quenched by addition of an aq. solution of NaOH (2 M) and diisopropylamine (10 eq.).*

**Step 2:** A secondary trigger amine hydrochloride (1.2 eq.) was suspended in anhydrous DCM (0.02 M). DIPEA (2.4 eq., 0.2 M solution in anhydrous DCM) was added to afford a clear solution of the corresponding free amine. A solution (0.02 M in anhydrous DCM) of **ClC(O)O-CBI-Boc** (1.0 eq) was added dropwise to the reaction flask at 0 °C and the resulting mixture was further stirred for 0.5 h, then allowed to warm to r.t. and further stirred for 0.5 h. Purification was achieved using FCC to give the title compounds as colourless solids.

**General protocol B:** Prodrug assembly (Amide coupling)

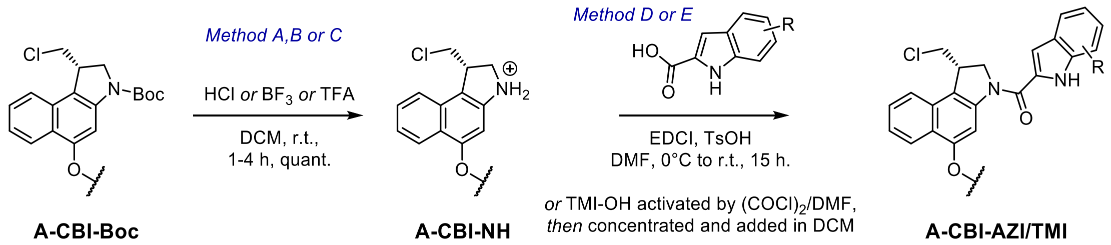


Step 1:

*Method A*: The *N*-Boc protected *O*-alkyl or *O*-acyl *seco*-CBI derivative **A-CBI-Boc** (1.0 eq.) was dissolved in anhydrous DCM (0.05 M). The solution was cooled to 0 °C and a solution of HCl (80 eq., 4 M in dioxane) was added. The mixture was stirred at r.t. for 4 h. All volatiles were removed under reduced pressure to afford the corresponding aniline **A-CBI-NH** without further purification as a coloured solid.

*Method B:* The *N*-Boc protected *O*-alkyl or *O*-acyl *seco*-CBI derivative **A-CBI-Boc** (1.0 eq.) was dissolved in anhydrous DCM (0.05 M). The solution was cooled to 0 °C and $BF_3 \cdot OEt_2$ (5.0 eq.) was added in one portion. The mixture was stirred at 0 °C for 1 h. All volatiles were removed under reduced pressure to afford the corresponding aniline **A-CBI-NH** without further purification as a coloured solid.

*Method C*: The *N*-Boc protected *O*-alkyl or *O*-acyl *seco*-CBI derivative **A-CBI-Boc** (1.0 eq.) was dissolved in anhydrous DCM (0.1 M). To the solution was added trifluoroacetic acid (60 eq.) and the resulting mixture was stirred at r.t. for 2 h. All volatiles were removed under reduced pressure, the residue was co-evaporated with toluene (1×) to afford the corresponding aniline **A-CBI-NH** without further purification as a coloured solid.

Step 2:

*Method D:* The free aniline *O*-alkyl or *O*-acyl *seco*-CBI derivative **A-CBI-NH** (1.0 eq.) was dissolved in anhydrous DMF (0.03 M). The solution was cooled to 0 °C and **AZI-OH** (1.1 eq., 68 wt%) or **TMI-OH**, EDCI (4.0 eq.) and *p*-TsOH (1.1 eq.) were added as solids. The mixture was stirred for 0.5 h, then allowed to warm to r.t. and further stirred for 15 h. All volatile compounds were removed under reduced pressure after co-evaporation with *n*-heptane to afford the crude mixture as a coloured solid. Purification by preparative HPLC ($H_2O$/MeCN/$HCO_2H$) afforded the title compounds **A-CBI-AZI** or **A-CBI-TMI** as colourless solids.

*Method E:* **TMI-OH** (2.0 eq.) was dissolved in anhydrous DCM (0.05 M), oxalyl chloride (6.0 eq.) was added and the mixture was cooled to 0 °C. A catalytic amount of DMF (1.0 eq.) was added and the mixture was allowed to warm to r.t. and was further stirred for 1.5 h. All volatiles were removed under reduced pressure, the residue was co-evaporated with toluene (1×) to afford the corresponding **TMI-Cl** without further purification as a brown solid. (*Comment: Full conversion to the carbonyl chloride was detected by trapping with piperazine; the carbonyl chloride can be stored as a solid at r.t. for 1-2 weeks*). The free aniline *O*-alkyl or *O*-acyl seco-CBI derivative **A-CBI-NH** (1.0 eq.) was dissolved in anhydrous DCM (0.05 M), cooled to 0 °C and DIPEA (8.0 eq.) was added to give a clear solution. Then, a solution of **TMI-Cl** (0.2 M in DCM, 2.0 eq.) was added dropwise, the mixture was allowed to warm to r.t. and was further stirred for 30 min. Conversion of the aniline was checked by LC-MS and further carbonyl chloride equivalents were added if required to ensure full conversion to the desired amide. The mixture was diluted with DCM, washed with water (2×) and the aq. layer was extracted with DCM (2×). The combined organic layers were dried over $Na_2SO_4$ and concentrated to give a coloured solid. Purification was achieved by FCC (DCM/MeOH or isohexane/EtOAc) to give the title compounds **A-CBI-TMI** as colourless solids. Further purification for cellular and *in vivo* application was achieved by preparative HPLC ($H_2O$/MeCN/0.1% formic acid).

*9.2. Precursor syntheses*

**SS60-CBI-Boc**

*N*-Boc (*S*)-5-((((*S*)-1,2-dithian-4-yl)(methyl)carbamoyl)oxy)-1-(chloromethyl)-1,2-dihydro-3*H*-benzo[*e*]indole

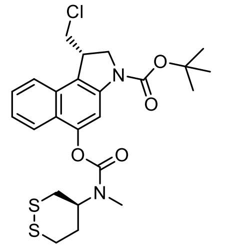

Carbamate coupling of **HO-CBI-Boc** (84 mg, 0.25 mmol) and (*S*)-*N*-methyl-1,2-dithian-4-amine hydrochloride (58 mg, 0.31 mmol) was accomplished according to the **General Protocol A** to give **SS60-CBI-Boc** as a colourless solid (123 mg, 0.24 mmol, 96%).

**TLC** $R_f$ = 0.64 (isohexane:EtOAc, 2:1). **HRMS** ($ESI^+$) for $C_{24}H_{30}ClN_2O_4S_2^+$: calcd. *m/z* 526.15955, found *m/z* 526.15972. NMR experiments at r.t. showed two rotameric species: **$^1$H-NMR** (400 MHz, $CDCl_3$): δ (ppm) = 8.05 (*br*-s, 1H), 7.78 (dd, *J* = 15.2, 8.4 Hz, 1H), 7.70 (d, *J* = 8.4 Hz, 1H), 7.50 (t, *J* = 7.6 Hz, 1H), 7.41 – 7.32 (m, 1H), 4.61 – 4.31 (m, 1H), 4.27 (d, *J* = 10.9 Hz, 1H), 4.13 (dd, *J* = 11.7, 8.8 Hz, 1H), 4.08 – 3.97 (m, 1H), 3.86 (dd, *J* = 11.2, 3.2 Hz, 1H), 3.47 (t, *J* = 10.8 Hz, 1H), 3.32 – 3.16 (m, 2H), 3.18 – 2.91 (m, 4H), 2.88 – 2.76 (m, 1H),, 2.44 – 2.02 (m, 2H), 1.58 (s, 9H). **$^{13}$C-NMR** (101 MHz, $CDCl_3$): δ (ppm) = 154.3 (*C*=O, major rotamer), 154.1 (*C*=O, minor rotamer), 152.5 (*C*=O), 148.6 ($C_{Ar}$), 141.3 ($C_{Ar}$), 130.3 ($C_{Ar}$), 127.7 ($C_{Ar}$H), 124.4 ($C_{Ar}$H), 124.2 ($C_{Ar}$), 122.7 ($C_{Ar}$H), 122.5 ($C_{Ar}$H), 120.2 ($C_{Ar}$), 109.4 ($C_{Ar}$H) 81.7 (*C*($CH_3$)$_3$), 56.3 (*C*H, minor rotamer), 55.3 (*C*H, major rotamer), 53.1 (*C*$H_2$), 46.0 (*C*$H_2$), 42.2 (*C*H), 36.5 (*C*$H_2$, minor rotamer), 36.3 (*C*$H_2$, minor rotamer), 36.2 (*C*$H_2$, major rotamer), 35.8 (*C*$H_2$, major rotamer), 33.4 (*C*$H_2$, minor rotamer), 32.6 (*C*$H_2$, major rotamer), 30.1 (*C*$H_3$), 28.6 (C(*C*$H_3$)$_3$).

**P-SS60-CBI-Boc**

*N*-Boc (*S*)-5-(((1,2-dithian-4-yl)(3-(4-methylpiperazin-1-yl)-3-oxopropyl)carbamoyl) oxy)-1-(chloromethyl)-1,2-dihydro-*3H*-benzo[*e*]indole

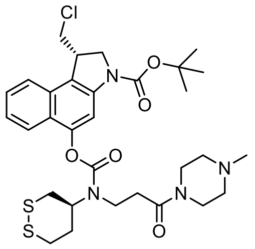

Carbamate coupling of **HO-CBI-Boc** (168 mg, 0.50 mmol) and (*S*)-3-((1,2-dithian-4-yl)amino)-1-(4-methylpiperazin-1-yl)propan-1-one dihydrochloride (199 mg, 0.55 mmol) was accomplished according to the **General Protocol A** to give **P-SS60-CBI-Boc** as a colourless solid (191 mg, 0.29 mmol, 58%).

**TLC** $R_f$ = 0.47 (DCM:MeOH, 9:1). **HRMS** ($ESI^+$) for $C_{31}H_{42}ClN_4O_5S_2^+$: calcd. *m/z* 649.22797, found *m/z* 649.22830. **$^1$H-NMR** (500 MHz, $CDCl_3$): δ (ppm) = 8.04 (s, 1H), 7.72 (dd, *J* = 16.5, 8.4 Hz, 2H), 7.51 (t, *J* = 7.1 Hz, 1H), 7.43 – 7.33 (m, 1H), 4.49 – 4.16 (m, 2H), 4.17 – 4.10 (m, 1H), 4.02 (t, *J* = 9.5 Hz, 1H), 3.93 (d, *J* = 11.1 Hz, 1H), 3.86 – 3.79 (m, 1H), 3.64 (s, 3H), 3.53 (d, *J* = 15.7 Hz, 2H), 3.50 – 3.44 (m, 1H), 3.35 (dt, *J* = 41.8, 12.0 Hz, 1H), 3.28 – 3.16 (m, 1H), 3.04 (dd, *J* = 31.0, 13.8 Hz, 1H), 2.96 (d, *J* = 12.3 Hz, 1H), 2.86 – 2.78 (m, 1H), 2.72 (ddt, *J* = 23.1, 14.7, 7.8 Hz, 1H), 2.40 (s, 5H), 2.30 (d, *J* = 3.6 Hz, 4H), 1.58 (s, 9H). **$^{13}$C-NMR** (126 MHz, $CDCl_3$): δ (ppm) = 169.1 (*C*=O), 168.7 (*C*=O), 154.1 (*C*=O), 152.6 ($C_{Ar}$), 148.3 ($C_{Ar}$), 141.2 ($C_{Ar}$), 130.4 ($C_{Ar}$), 127.8 ($C_{Ar}$H), 124.6 ($C_{Ar}$H), 124.1, 122.6 ($C_{Ar}$H), 120.6 ($C_{Ar}$), 109.4 ($C_{Ar}$H), 81.5 (*C*($CH_3$)$_3$), 58.2 (*C*H, minor rotamer), 57.8 (*C*H, major rotamer), 55.1 (*C*$H_2$), 54.6 (*C*$H_2$), 54.2 (*C*H), 53.1 (*C*$H_2$), 46.4 (*C*$H_2$), 46.0 (*C*$H_3$, minor rotamer), 45.9 (*C*$H_3$, major rotamer), 45.4 (*C*$H_2$), 42.4 (*C*H), 42.1 (*C*$H_2$), 41.5 (*C*$H_2$), 41.3 (*C*$H_2$), 37.1 (*C*$H_2$), 36.6 (*C*$H_2$), 36.4 (*C*$H_2$), 36.3 (*C*$H_2$), 34.3 (*C*$H_2$), 34.0 (*C*$H_2$), 33.4 (*C*$H_2$), 33.0 (*C*$H_2$), 28.6 (C(*C*$H_3$)$_3$).

**SeS60-CBI-Boc**

*N*-Boc (*S*)-1-(chloromethyl)-5-((methyl((*S*)-1,2-thiaselenan-4-yl)carbamoyl)oxy)-1,2-dihydro-3*H*-benzo[*e*]indole

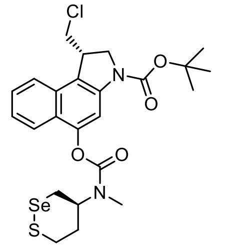

Carbamate coupling of **HO-CBI-Boc** (77 mg, 0.23 mmol) and (*S*)-*N*-methyl-1,2-thiaselenan-4-amine hydrochloride (56 mg, 0.24 mmol) was accomplished according to the **General Protocol A** to give **SeS60-CBI-Boc** as a colourless, crystalline solid (85 mg, 0.15 mmol, 65%).

**TLC** $R_f$ = 0.79 (isohexane:EtOAc, 2:1). **HRMS** (ESI$^+$) for $C_{24}H_{30}ClN_2O_4SSe^+$: calcd. *m/z* 574.10400, found *m/z* 574.10383. NMR experiments at r.t. showed two rotameric species: **$^1$H-NMR** (500 MHz, $CDCl_3$): δ (ppm) = 8.05 (s, 1H), 7.78 (dd, *J* = 12.9, 8.6 Hz, 1H), 7.70 (d, *J* = 8.3 Hz, 1H), 7.50 (t, *J* = 7.3 Hz, 1H), 7.41 – 7.32 (m, 1H), 4.54 – 4.20 (m, 2H), 4.12 (dd, *J* = 13.8, 6.6 Hz, 1H), 4.01 (t, *J* = 9.5 Hz, 1H), 3.93 (d, *J* = 10.8 Hz, 1H), 3.46 (t, *J* = 10.9 Hz, 2H), 3.32 (dt, *J* = 33.0, 12.3 Hz, 1H), 3.20 – 3.05 (m, 3H), 2.95 (d, *J* = 16.8 Hz, 2H), 2.58 – 2.16 (m, 2H), 1.58 (s, 9H). **$^{13}$C-NMR** (126 MHz, $CDCl_3$): δ (ppm) = 154.2 (*C*=O, major rotamer), 154.0 (*C*=O, minor rotamer), 152.5 (*C*=O), 148.5 ($C_{Ar}$), 141.2 ($C_{Ar}$), 130.3 ($C_{Ar}$), 127.7 ($C_{Ar}$H), 124.4 ($C_{Ar}$H), 122.6 ($C_{Ar}$H), 122.4 ($C_{Ar}$H), 120.3 ($C_{Ar}$), 109.4 ($C_{Ar}$H), 81.4 (*C*($CH_3$)$_3$), 57.7 (*C*H, major rotamer), 57.1 (*C*H, minor rotamer), 53.0 (*C*$H_2$), 46.4 (*C*$H_2$), 42.2 (*C*H), 36.0 (*C*$H_2$), 35.3 (*C*$H_2$), 34.2 (*C*$H_2$), 33.4 (*C*$H_2$), 30.2 (*C*$H_3$), 28.5 (C(*C*$H_3$)$_3$), 28.0 (*C*$H_2$), 27.9 (*C*$H_2$).

**H-SS66T-CBI-Boc**

*N*-Boc (*S*)-1-(chloromethyl)-2,3-dihydro-1*H*-benzo[*e*]indol-5-yl (*trans*)-hexahydro-[1,2]dithiino[4,5-*b*]pyrazine-1(2*H*)-carboxylate

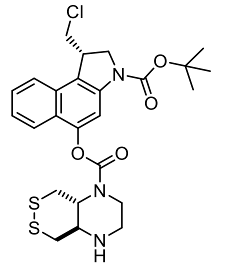

Carbamate coupling of **HO-CBI-Boc** (77.2 mg, 0.23 mmol) and *trans*-octahydro-[1,2]dithiino[4,5-*b*]pyrazine dihydrochloride (100.8 mg, 63 wt%, 0.35 mmol) was accomplished according to **General Protocol A** to give **H-SS66T-CBI-Boc** as a colourless solid (42.0 mg, 0.08 mmol, 35%).

**TLC** $R_f$ = 0.32 (EtOAc). **HRMS** (ESI$^+$) for $C_{25}H_{31}ClN_3O_4S_2^+$: calcd. m/z 536.14390, found *m/z* 536.14431. **$^1$H-NMR** (500 MHz, $CDCl_3$) δ (ppm) = 8.06 (s, 1H), 7.86 – 7.77 (m, 1H), 7.71 (d, *J* = 8.4 Hz, 1H), 7.52 (t, *J* = 7.5 Hz, 1H), 7.46 – 7.32 (m, 1H), 4.25 (s, 1H), 4.19 – 4.12 (m, 1H), 4.09 – 3.98 (m, 3H), 3.93 (d, *J* = 10.8 Hz, 1H), 3.65 (s, 1H), 3.48 (td, *J* = 11.0, 1.9 Hz, 1H), 3.40 – 3.25 (m, 5H), 3.04 (dt, *J* = 13.8, 7.3 Hz, 2H), 2.94 – 2.74 (m, 1H), 1.58 (s, 9H). **$^{13}$C-NMR** (126 MHz, $CDCl_3$) δ (ppm) = 154.0 (*C*=O), 152.6 (*C*=O), 148.1 ($C_{Ar}$), 141.2 ($C_{Ar}$), 130.3 ($C_{Ar}$), 127.8 ($C_{Ar}$H), 124.6 ($C_{Ar}$H), 124.0 ($C_{Ar}$), 122.7 ($C_{Ar}$H), 122.5 ($C_{Ar}$H), 120.6 ($C_{Ar}$), 109.3 ($C_{Ar}$H), 81.6 (*C*($CH_3$)$_3$), 63.9 (*C*H), 56.7 (*C*H), 53.1 (*C*$H_2$), 46.4 (*C*$H_2$), 44.6 (*C*$H_2$), 42.2 (*C*H), 39.9 (*C*$H_2$), 38.0 (*C*$H_2$), 28.6 (C(*C*$H_3$)$_3$).

**Me-SS66T-CBI-Boc**

*N*-Boc (*S*)-1-(chloromethyl)-2,3-dihydro-1*H*-benzo[*e*]indol-5-yl (*trans*)-4-methylhexahydro-[1,2]dithiino[4,5-*b*] pyrazine-1(2*H*)-carboxylate

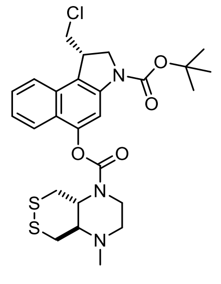

**H-SS66T-CBI-Boc** (20.0 mg, 0.037 mmol) was dissolved in MeOH (5 mM) and a solution of formaldehyde (424 µL, 0.88 M in MeOH, 0.370 mmol) was added at r.t., followed by addition of a solution of $NaCNBH_3$ (40 µL, 1 M in MeOH, 0.040 mmol). The reaction mixture was stirred at r.t. for 30 min, was then concentrated under reduced pressure; the residue was taken into DCM (10 mL) and washed with aq. NaCl (20 mL). The aq. layer was extracted with DCM (4×10 mL), the combined organic layers were dried over $Na_2SO_4$, filtered and concentrated under reduced pressure. Purification was achieved by FCC (EtOAc) to give **Me-SS66T-CBI-Boc** (12.0 mg, 0.022 mmol, 59%) as a colourless solid.

**TLC** $R_f$ = 0.53 (EtOAc). **HRMS** ($ESI^+$) for $C_{26}H_{33}ClN_3O_4S_2^+$: calcd. m/z 550.15955, found *m/z* 550.16002. **$^1$H-NMR** (500 MHz, $CDCl_3$): δ (ppm) = 8.06 (s, 1H), 7.81 (d, *J* = 8.6 Hz, 1H), 7.70 (d, *J* = 8.3 Hz, 1H), 7.55 – 7.48 (m, 1H), 7.42 – 7.35 (m, 1H), 4.27 (s, 1H), 4.13 (dd, *J* = 12.0, 8.9 Hz, 1H), 4.07 – 3.97 (m, 2H), 3.97 – 3.87 (m, 2H), 3.72 (s, 1H), 3.56 – 3.41 (m, 2H), 3.29 (d, *J* = 11.6 Hz, 1H), 3.19 (dt, *J* = 14.4, 4.8 Hz, 2H), 3.03 (ddd, *J* = 13.1, 10.6, 2.1 Hz, 1H), 2.90 (t, *J* = 10.1 Hz, 1H), 2.78 (dt, *J* = 10.8, 5.0 Hz, 1H), 2.41 (d, *J* = 0.9 Hz, 3H), 1.58 (s, 9H).**$^{13}$C-NMR** (126 MHz, $CDCl_3$): δ (ppm) = 153.8 (*C*=O), 152.5 (*C*=O), 148.3 ($C_{Ar}$), 141.2 ($C_{Ar}$), 130.4 ($C_{Ar}$), 127.7 ($C_{Ar}$H), 124.5 ($C_{Ar}$H), 124.1 ($C_{Ar}$), 122.7 ($C_{Ar}$H), 122.5 ($C_{Ar}$H), 120.4 ($C_{Ar}$), 109.4 ($C_{Ar}$H), 81.4 ($C(CH_3)_3$), 63.6 (*C*H), 62.6 (*C*H), 55.1 ($CH_3$), 53.1 ($CH_2$), 46.4 ($CH_2$), 42.2 (*C*H), 40.6 ($CH_2$), 38.9 ($CH_2$), 37.9 ($CH_2$), 28.6 ($C(CH_3)_3$).

**H-SS66C-CBI-Boc**

*N*-Boc (*S*)-1-(chloromethyl)-2,3-dihydro-1*H*-benzo[*e*]indol-5-yl (*cis*)-hexahydro-[1,2]dithiino[4,5-*b*]pyrazine-1(2*H*)-carboxylate

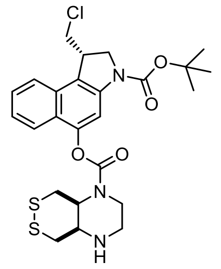

Carbamate coupling of **HO-CBI-Boc** (90.6 mg, 0.27 mmol) and *cis*-octahydro-[1,2]dithiino[4,5-*b*]pyrazine dihydrochloride (134.0 mg, 63 wt%, 0.34 mmol) was accomplished according to **General Protocol A**. **H-SS66C-CBI-Boc** was obtained as a colourless solid (97.1 mg, 0.18 mmol, 67%).

**TLC** $R_f$ = 0.35 (EtOAc). **HRMS** ($ESI^+$) for $C_{25}H_{31}ClN_3O_4S_2^+$: calcd. m/z 536.14390, found *m/z* 536.14355. Individual rotamers were observed by NMR spectroscopy at 298 K (ratio ca. 1:1). **$^1$H-NMR** (600 MHz, $CDCl_3$) δ (ppm) = 8.05 (s, 2H), 7.80 – 7.72 (m, 2H), 7.70 (d, *J* = 8.4 Hz, 2H), 7.50 (t, *J* = 7.3 Hz, 2H), 7.37 (q, *J* = 7.3 Hz, 2H), 4.56 (d, *J* = 11.8 Hz, 1H), 4.37 (d, *J* = 11.6 Hz, 1H), 4.30 – 4.09 (m, 5H), 4.06 – 3.89 (m, 5H), 3.77 (td, *J* = 12.6, 7.6 Hz, 1H), 3.66 (t, *J* = 12.3 Hz, 1H), 3.58 – 3.44 (m, 4H), 3.42 (s, 1H), 3.40 – 3.33 (m, 1H), 3.31 – 3.19 (m, 3H), 3.13 – 2.98 (m, 6H), 1.93 (s, 2H), 1.58 (s, 18H). **$^{13}$C-NMR** (151 MHz, $CDCl_3$) δ (ppm) = 153.6 (*C*=O, rotamer 1), 153.4 (*C*=O, rotamer 2), 152.5 (*C*=O), 148.3 ($C_{Ar}$), 141.2 ($C_{Ar}$), 130.3 ($C_{Ar}$), 127.7 ($C_{Ar}$H), 124.5 ($C_{Ar}$H, rotamer 1), 124.4 ($C_{Ar}$H, rotamer 2), 122.5 (2×$C_{Ar}$H), 120.6 ($C_{Ar}$, rotamer 1), 120.4 ($C_{Ar}$, rotamer 2), 109.4 ($C_{Ar}$H), 81.5 ($C(CH_3)_3$), 53.2 (*C*H, rotamer 1), 53.1 ($CH_2$), 52.9 (*C*H, rotamer 1), 52.6 (*C*H, rotamer 2), 52.0 (*C*H, rotamer 2), 46.4 ($CH_2$), 45.6 ($CH_2$, rotamer 1), 45.4 ($CH_2$, rotamer 2), 42.4 ($CH_2$, rotamer 1), 42.2 (*C*H, $CH_2$, rotamer 1), 41.1 ($CH_2$), 40.3 ($CH_2$), 30.0 ($CH_2$, rotamer 1), 29.8 ($CH_2$, rotamer 1), 29.2 ($CH_2$, rotamer 2), 29.1 ($CH_2$, rotamer 2), 28.5 ($C(CH_3)_3$).

**Me-SS66C-CBI-Boc**

*N*-Boc (*S*)-1-(chloromethyl)-2,3-dihydro-1*H*-benzo[e]indol-5-yl (cis)-4-methylhexahydro-[1,2]dithiino[4,5-*b*] pyrazine-1(2*H*)-carboxylate

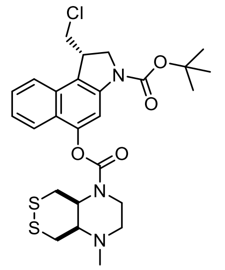

**H-SS66C-CBI-Boc** (78.0 mg, 0.145 mmol) was dissolved in MeOH (5 mM) and a solution of formaldehyde (4.96 mL, 0.88 M in MeOH, 4.36 mmol) was added at r.t., followed by addition of a solution of $NaCNBH_3$ (91.7 mg, 0.5 M solution in MeOH, 1.45 mmol). The reaction mixture was stirred at r.t. for 30 min, was then concentrated under reduced pressure; the residue was taken into DCM (10 mL) and washed with aq. NaCl (20 mL). The aq. layer was extracted with DCM (4×10 mL), the combined organic layer were dried over $Na_2SO_4$, filtered and concentrated under reduced pressure. Purification was achieved by FCC (isohexane/EtOAc) to give **Me-SS66C-CBI-Boc** (39.8 mg, 0.072 mmol, 50%) as a colourless solid. A fraction of unreacted starting material **H-SS66C-CBI-Boc** (32.7 mg, 0.061 mmol, 42 %) was recovered.

**TLC** $R_f$ = 0.47 (isohexane:EtOAc, 1:1). **HRMS** ($ESI^+$) for $C_{26}H_{33}ClN_3O_4S_2^+$: calcd. *m/z* 550.15955, found *m/z* 550.15910. Individual rotamers were observed by NMR spectroscopy at 298 K (ratio ca. 1:1). **$^1$H-NMR** (800 MHz, $CDCl_3$): δ (ppm) = 8.05 (s, 2H), 7.78 – 7.75 (m, 1H), 7.75 – 7.72 (m, 1H), 7.70 (d, *J* = 8.2 Hz, 2H), 7.51 (t, *J* = 7.4 Hz, 2H), 7.37 (q, *J* = 7.2, 6.5 Hz, 2H), 4.57 (d, *J* = 10.2 Hz, 1H), 4.38 (d, *J* = 10.8 Hz, 1H), 4.30 – 4.20 (m, 3H), 4.19 – 4.07 (m, 2H), 4.09 – 3.96 (m, 4H), 3.96 – 3.89 (m, 3H), 3.57 (s, 1H), 3.55 – 3.45 (m, 2H), 3.45 – 3.26 (m, 5H), 3.10 – 2.93 (m, 2H), 2.69 (d, *J* = 7.8 Hz, 1H), 2.63 (s, 1H), 2.60 – 2.48 (m, 3H), 2.45 (t, *J* = 11.4 Hz, 1H), 2.32 (s, 6H), 1.57 (s, 18H). **$^{13}$C-NMR** (201 MHz, $CDCl_3$): δ (ppm) = 153.3 (*C*=O, rotamer 1), 153.2 (*C*=O, rotamer 2), 152.5 (*C*=O), 148.3 ($C_{Ar}$), 141.3 ($C_{Ar}$), 130.3 ($C_{Ar}$), 127.7 ($C_{Ar}$H), 124.5 ($C_{Ar}$H, rotamer 1), 124.4 ($C_{Ar}$H, rotamer 2), 124.1 ($C_{Ar}$), 122.5 (2×$C_{Ar}$H), 120.5 ($C_{Ar}$), 109.4 ($C_{Ar}$H), 81.5 (*C*($CH_3$)$_3$), 61.5 (*C*H, rotamer 1), 61.3 (*C*H, rotamer 2), 56.3 (*C*$H_2$, rotamer 1), 56.0 (*C*$H_2$, rotamer 2), 54.8 (*C*H, rotamer 1), 53.7 (*C*H, rotamer 2), 53.1 (*C*$H_2$), 46.4 (*C*$H_2$), 42.2 (*C*H), 41.9 (*C*$H_3$), 40.9 (*C*$H_2$, rotamer 1), 40.2 (*C*$H_2$, rotamer 2), 39.2 (*C*$H_2$, rotamer 1), 38.9 (*C*$H_2$, rotamer 2), 30.7 (*C*$H_2$, rotamer 1), 29.8 (*C*$H_2$, rotamer 2), 28.6 (C(*C*$H_3$)$_3$).

**SS00$_M$-CBI-Boc**

*N*-Boc (*S*)-1-(chloromethyl)-5-((methyl(2-((2-morpholinoethyl)disulfaneyl)ethyl) carbamoyl)oxy)-1,2-dihydro-3*H*-benzo[*e*]indole

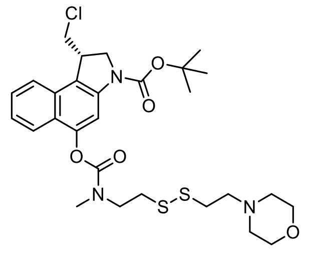

Carbamate coupling of **HO-CBI-Boc** (116.5 mg, 0.350 mmol) and *N*-methyl-2-((2-morpholinoethyl)disulfaneyl) ethan-1-amine dihydrochloride (130.0 mg, 0.42 mmol) was accomplished according to **General Protocol A** to give **SS00$_M$-CBI-Boc** as a colourless solid (206.0 mg, 0.346 mmol, 99%).

**TLC** $R_f$ = 0.58 (EtOAc:$NEt_3$, 30:1). **HRMS** ($ESI^+$) for $C_{28}H_{39}N_3O_5Cl^+$: calcd. *m/z* 596.20142, found *m/z* 596.20393. **$^1$H-NMR** (400 MHz, $CDCl_3$): δ (ppm) = 8.06 (s, 1H), 7.86 (d, *J* = 8.8 Hz, 1H), 7.70 (d, *J* = 8.4 Hz, 1H), 7.50 (t, *J* = 7.3 Hz, 1H), 7.41 – 7.34 (m, 1H), 4.25 (s, 1H), 4.13 (dd, *J* = 11.8, 8.7 Hz, 1H), 4.01 (t, *J* = 9.6 Hz, 1H), 3.93 (d, *J* = 11.1 Hz, 1H), 3.87 (t, *J* = 7.0 Hz, 1H), 3.74 – 3.59 (m, 5H), 3.46 (td, *J* = 10.9, 2.5 Hz, 1H), 3.28 (s, 2H, major rotamer), 3.10 (s, 1H, minor rotamer), 3.07 – 3.02 (m, 1H), 2.97 (t, *J* = 6.9 Hz, 1H), 2.90 – 2.83 (m, 2H), 2.68 (dd, *J* = 13.1, 7.0 Hz, 2H), 2.53 – 2.44 (m, 2H), 2.41 (s, 2H), 1.58 (s, 9H). **$^{13}$C-NMR** (101 MHz, $CDCl_3$): δ (ppm) = 155.1 (*C*=O), 154.5 (*C*=O), 152.5 ($C_{Ar}$), 148.0 ($C_{Ar}$), 141.6 ($C_{Ar}$), 130.6 ($C_{Ar}$), 127.7 ($C_{Ar}$H), 126.8 ($C_{Ar}$), 124.4 ($C_{Ar}$H), 122.8 ($C_{Ar}$H), 122.4 ($C_{Ar}$H), 109.5 ($C_{Ar}$H), 81.5 (*C*($CH_3$)$_3$), 66.9 (*C*$H_2$), 58.1 (*C*$H_2$), 53.6 (*C*$H_2$), 53.3 (*C*$H_2$), 49.2 (*C*$H_2$), 46.4 (*C*$H_2$), 42.2 (*C*H), 36.4 (*C*$H_2$), 35.9 (*C*$H_3$), 28.6 (C(*C*$H_3$)$_3$).

**P-SS66C-CBI-Boc**

*N*-Boc (*S*)-1-(chloromethyl)-2,3-dihydro-1*H*-benzo[*e*]indol-5-yl (*cis*)-4-(4-(4-methylpiperazin-1-yl)-4-oxobutanoyl)hexahydro-[1,2]dithiino[4,5-*b*]pyrazine-1(2*H*)-carboxylate

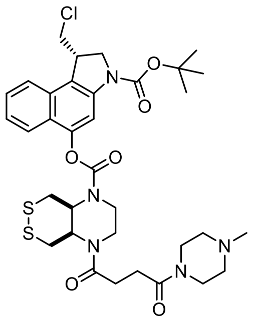

4-(4-methylpiperazin-1-yl)-4-oxobutanoic acid hydrochloride (68.0 mg, 0.29 µmol, 7.0 eq.) was dissolved in anhydrous 1,2-dichloroethane (0.15 M) and DIPEA (98 µL, 0.56 µmol, 14.0 eq.) was added followed by trimethylacetyl chloride (35 µL, 0.29 mmol, 7.0 eq.). The resulting mixture was stirred at r.t. for 2 h, before **H-SS66C-CBI-Boc** (22.0 mg, 0.041 mmol, 1.0 eq.) in 1,2-dichloroethane (0.1 M) was added in one portion. The mixture was heated to 70 °C for 15 h, was then concentrated under reduced pressure and taken into DCM (50 mL). The organic layer was washed with aq. NaCl (1 M, 4×20 mL), was then dried over $Na_2SO_4$ and concentrated under reduced pressure. Purification was achieved using FCC (DCM/MeOH) to yield **P-SS66C-CBI-Boc** as a colourless solid (26.0 mg, 0.036 mmol, 88%).

**TLC** $R_f$ = 0.53 (DCM:MeOH, 9:1). **HRMS** (ESI$^+$) for $C_{34}H_{45}ClN_5O_6S_2^+$: calcd. *m/z* 718.24943, found *m/z* 718.24928. **$^1$H-NMR** (400 MHz, $CDCl_3$): δ (ppm) = 8.06 (s, 1H), 7.77 (d, *J* = 8.5 Hz, 1H), 7.71 (d, *J* = 8.4 Hz, 1H), 7.51 (t, *J* = 7.2 Hz, 1H), 7.39 (t, *J* = 7.4 Hz, 1H), 4.75 (s, 1H), 4.44 (s, 2H), 4.35 – 4.21 (m, 1H), 4.20 – 4.09 (m, 1H), 4.01 (dd, *J* = 13.6, 8.1 Hz, 3H), 3.93 (d, *J* = 11.9 Hz, 1H), 3.89 – 3.62 (m, 6H), 3.57 – 3.40 (m, 2H), 3.20 (d, *J* = 14.1 Hz, 1H), 3.01 (d, *J* = 11.0 Hz, 1H), 2.81 – 2.54 (m, 8H), 2.46 (s, 3H), 1.57 (s, 9H). **$^{13}$C-NMR** (151 MHz, $CDCl_3$): δ (ppm) = 172.1 (*C*=O), 170.4 (*C*=O), 154.0 (*C*=O), 152.5 (*C*=O), 148.0 ($C_{Ar}$), 141.2 ($C_{Ar}$), 130.4 ($C_{Ar}$), 127.8 ($C_{Ar}$H), 124.6 ($C_{Ar}$H), 123.9 ($C_{Ar}$), 122.5 (2×$C_{Ar}$H), 120.6 ($C_{Ar}$), 109.4 ($C_{Ar}$H), 81.5 (*C*(CH$_3$)$_3$), 54.6 (*C*H$_2$), 54.4 (*C*H$_2$), 53.1 (*C*H$_2$), 51.9 (*C*H), 46.4 (*C*H$_2$), 45.4 (*C*H), 44.6 (*C*H$_2$), 42.2 (*C*H), 42.0 (*C*H$_2$), 41.0 (*C*H$_2$), 36.0 (*C*H$_2$), 28.5 (C(*C*H$_3$)$_3$), 28.0 (*C*H$_2$).

**SS66T-CBI-Boc**

*N*-Boc (*S*)-1-(chloromethyl)-2,3-dihydro-1*H*-benzo[*e*]indol-5-yl (*trans*)-hexahydro-[1,2]dithiino[4,5-*b*]pyridine-1(2*H*)-carboxylate

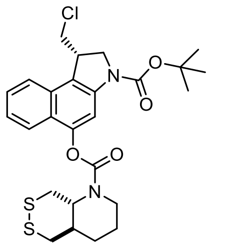

Carbamate coupling of **HO-CBI-Boc** (139.0 mg, 0.35 mmol) and *trans*-octahydro-[1,2]dithiino[4,5-*b*]pyridine hydrochloride (88.9 mg, 0.42 mmol) was accomplished according to the **General Protocol A** to give **SS66T-CBI-Boc** as a colourless, crystalline solid (162 mg, 0.30 mmol, 86%). *This compound was not further used for prodrug assembly in this study, but it is added to this report for comparison.*

**TLC** $R_f$ = 0.44 (isohexane:EtOAc, 3:1). **HRMS** (ESI$^+$) for $C_{26}H_{35}ClN_3O_3S_2^+$: calcd. m/z 552.17520, found *m/z* 552.17594. **$^1$H-NMR** (500 MHz, $CDCl_3$) δ (ppm) = 8.06 (s, 1H), 7.78 (d, *J* = 8.4 Hz, 1H), 7.70 (d, *J* = 8.4 Hz, 1H), 7.51 (t, *J* = 7.3 Hz, 1H), 7.38 (t, *J* = 7.6 Hz, 1H), 4.26 (d, *J* = 10.8 Hz, 1H), 4.13 (t, *J* = 10.2 Hz, 1H), 4.05 – 3.96 (m, 2H), 3.96 – 3.82 (m, 2H), 3.57 – 3.40 (m, 2H), 3.21 (d, *J* = 5.9 Hz, 2H), 3.03 – 2.91 (m, 1H), 2.83 (dd, *J* = 13.5, 2.6 Hz, 1H), 2.28 – 2.14 (m, 1H), 2.12 – 1.99 (m, 1H), 1.93 – 1.78 (m, 1H), 1.77 – 1.67 (m, 1H), 1.58 (s, 9H), 1.54 – 1.43 (m, 1H). **$^{13}$C-NMR** (126 MHz, $CDCl_3$) δ (ppm) = 154.0 (*C*=O), 152.5 (*C*=O), 148.4 ($C_{Ar}$), 141.5 ($C_{Ar}$), 130.3 ($C_{Ar}$), 128.1 ($C_{Ar}$), 127.7 ($C_{Ar}$H), 124.5 ($C_{Ar}$H), 122.6 ($C_{Ar}$H), 122.5 ($C_{Ar}$H), 120.4 ($C_{Ar}$), 109.4 ($C_{Ar}$H), 81.9 (*C*(CH$_3$)$_3$), 63.3 (*C*H), 53.0 (*C*H$_2$), 46.4 (*C*H$_2$), 42.2 (*C*H), 41.2 (*C*H$_2$), 40.8 (*C*H), 39.7 (*C*H$_2$), 37.2 (*C*H$_2$), 28.5 (C(*C*H$_3$)$_3$), 27.3 (*C*H$_2$), 23.3 (*C*H$_2$).

**SS66-CBI-Boc** (racemate; *cis:trans* 1:1)

*N*-Boc (*S*)-1-(chloromethyl)-2,3-dihydro-1*H*-benzo[*e*]indol-5-yl hexahydro-[1,2]dithiino[4,5-*b*]pyridine-1(2*H*)-carboxylate

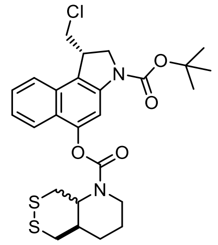


Carbamate coupling of **HO-CBI-Boc** (269 mg, 0.68 mmol) and octahydro-[1,2]dithiino[4,5-*b*]pyridine hydrochloride (racemate; *cis:trans* 1:1) (144 mg, 0.68 mmol) was accomplished according to the **General Protocol A** to give **SS66-CBI-Boc** as a colourless, crystalline solid (245 mg, 0.46 mmol, 68%). *This compound was not further used for prodrug assembly in this study, but it is added to this report for comparison.*

**TLC** $R_f$ = 0.44 (isohexane:EtOAc, 3:1, two isomers indistinguishable on TLC). **HRMS** (ESI$^+$) for $C_{26}H_{35}ClN_3O_3S_2^+$: calcd. m/z 552.17520, found *m/z* 552.17531. **$^1$H-NMR** (400 MHz, $CDCl_3$): δ (ppm) = 8.05 (s, 1H), 7.77 (dd, *J* = 10.9, 7.8 Hz, 1H), 7.70 (d, *J* = 8.5 Hz, 1H), 7.50 (tt, *J* = 8.9, 1.7 Hz, 1H), 7.41 – 7.32 (m, 1H), 4.84 – 4.50 (m, 0.5H, *cis*), 4.35 (d, *J* = 13.6 Hz, 0.5H, *cis*), 4.27 (d, *J* = 11.4 Hz, 1H, *trans*), 4.21 – 4.09 (m, 1.5H), 4.06 – 3.98 (m, 1.5H), 3.97 – 3.89 (m, 1.5H), 3.64 (td, *J* = 12.7, 3.3 Hz, 0.5H, *cis*), 3.59 – 3.36 (m, 2H), 3.21 (d, *J* = 6.9 Hz, 1H, *trans*), 3.13 (d, *J* = 13.4 Hz, 0.5H, *cis*), 3.02 – 2.92 (m, 1H), 2.92 – 2.78 (m, 1H), 2.76 – 2.50 (m, 1H, *cis*), 2.46 – 2.27 (m, 1H, *cis*), 2.27 – 2.13 (m, 0.5H, *trans*), 2.12 – 1.98 (m, 0.5H, *trans*), 1.96 – 1.80 (m, 1H), 1.77 – 1.67 (m, 1H), 1.58 (s, 9H), 1.54 – 1.43 (m, 1H). **$^{13}$C-NMR** (126 MHz, $CDCl_3$): δ (ppm) = 154.2 (*C*=O, *cis*), 154.0 (*C*=O, *trans*), 153.7 (*C*=O, *cis*), 153.5 (*C*=O, *cis*), 152.8 (*C*=O, *cis*), 152.5 (*C*=O, *trans*), 148.5 ($C_{Ar}$, *trans*), 147.3 ($C_{Ar}$, *cis*), 141.3 ($C_{Ar}$), 130.3 ($C_{Ar}$), 127.7 ($C_{Ar}$H, *trans*), 126.9 ($C_{Ar}$H, *cis*), 124.4 ($C_{Ar}$H), 122.6 ($C_{Ar}$H), 122.5 ($C_{Ar}$H), 120.2 ($C_{Ar}$), 109.4 ($C_{Ar}$H, *trans*), 109.3 ($C_{Ar}$H, *cis*), 81.4 (*C*(CH$_3$)$_3$), 63.3 (*C*H, *trans*), 57.0 (*C*H, *cis*), 54.3 (*C*H$_2$, *cis*), 53.3 (*C*H$_2$, *cis*), 53.1 (*C*H$_2$, *trans*), 46.4 (*C*H$_2$), 42.4 (*C*H, *cis*), 42.2 (*C*H, *trans*), 41.5 (*C*H$_2$, *cis*), 41.2 (*C*H$_2$, *trans*), 40.8 (*C*H, *trans*), 40.3 (*C*H, *cis*), 37.2 (*C*H$_2$, *trans*), 36.5(*C*H$_2$, *cis*), 36.0 (*C*H$_2$, *cis*), 33.8 (*C*H$_2$), 30.3 (*C*H$_2$), 29.5 (*C*H$_2$), 28.6 (C(*C*H$_3$)$_3$), 27.4 (*C*H$_2$, *trans*), 25.7 (*C*H$_2$, *cis*), 25.2 (*C*H$_2$, *cis*), 23.6 (*C*H$_2$, *cis*), 23.3 (*C*H$_2$, *trans*).

**MC-CBI-Boc**

(*S*)-1-(chloromethyl)-5-((methoxycarbonyl)oxy)-1,2-dihydro-3*H*-benzo[*e*]indole-3-carboxylate

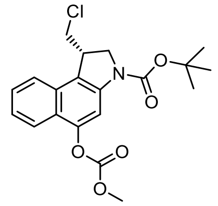


To a solution of **HO-CBI-Boc** (62.0 mg, 0.186 mmol) in anhydrous DCM (0.05 M) was added a solution of pyridine (75 µL, 0.56 mmol) and methyl chloroformate (43 µL, 0.56 mmol) at 0 °C. The mixture was stirred at r.t. for 1h, was then concentrated under reduced pressure to remove all volatiles. Purification by FCC (isohexane/EtOAc) afforded **MC-CBI-Boc** as a colourless solid (72.0 mg, 0.184 mmol, 99%).

**TLC** $R_f$ = 0.56 (isohexane:EtOAc, 6:1). **HRMS** (ESI$^+$) for $C_{20}H_{22}NNaO_5Cl^+$: calcd. *m/z* 414.10787, found *m/z* 414.10871. **$^1$H-NMR** (400 MHz, $CDCl_3$): δ (ppm) = 8.15 (s, 1H), 7.92 (d, *J* = 8.4 Hz, 1H), 7.71 (d, *J* = 8.4 Hz, 1H), 7.59 – 7.45 (m, 1H), 7.40 (ddd, *J* = 8.1, 6.9, 1.0 Hz, 1H), 4.27 (s, 1H), 4.15 (t, *J* = 10.2 Hz, 1H), 4.07 – 3.98 (m, 1H), 3.96 (s, 3H), 3.92 (s, 1H), 3.48 (t, *J* = 10.8 Hz, 1H), 1.59 (s, 9H). **$^{13}$C-NMR** (101 MHz, $CDCl_3$): δ (ppm) = 154.4 (*C*=O), 152.5 (*C*=O), 148.1 ($C_{Ar}$), 141.0 ($C_{Ar}$), 130.3 ($C_{Ar}$), 127.9 ($C_{Ar}$H), 124.6 ($C_{Ar}$H), 123.3 ($C_{Ar}$H), 122.5 ($C_{Ar}$H), 108.9 ($C_{Ar}$H), 81.6 (*C*(CH$_3$)$_3$), 56.6 (*C*H), 53.0 (*C*H$_2$), 47.0 (*C*H$_2$), 42.6 (*C*H$_3$), 28.6 (C(*C*H$_3$)$_3$).

**O56-CBI-Boc**

*N*-Boc (*S*)-1-(chloromethyl)-2,3-dihydro-1*H*-benzo[*e*]indol-5-yl (*cis*)-hexahydrofuro[3,4 *b*]pyridine-1(2*H*)-carboxylate

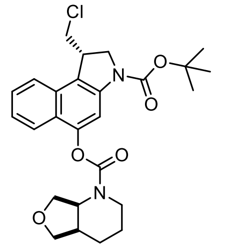

Carbamate coupling of **HO-CBI-Boc** (84 mg, 0.25 mmol) and *cis*-octahydrofuro[3,4-*b*]pyridine hydrochloride (51 mg, 0.31 mmol) was accomplished according to the **General Protocol A** to give **O56-CBI-Boc** as a colourless solid (116 mg, 0.24 mmol, 95%).

**TLC** $R_f$ = 0.32 (isohexane:EtOAc, 2:1). **HRMS** (ESI$^+$) for $C_{26}H_{32}ClN_2O_5^+$: calcd. *m/z* 504.22598, found *m/z* 504.22598. NMR experiments at r.t. showed two rotameric species. A dynamic equilibrium between both rotameric species in the $^1$H-NMR spectrum was induced at 323 K. **$^1$H-NMR** (400 MHz, $CDCl_3$, 323 K): δ (ppm) = 7.98 (s, 1H), 7.81 (d, *J* = 8.5 Hz, 1H), 7.70 (dt, *J* = 8.5, 1.0 Hz, 1H), 7.50 (ddd, *J* = 8.3, 6.8, 1.2 Hz, 1H), 7.36 (ddd, *J* = 8.2, 6.8, 1.1 Hz, 1H), 4.96 (s, 1H), 4.37 – 4.07 (m, 4H), 4.02 (ddd, *J* = 12.1, 7.7, 2.8 Hz, 1H), 3.98 – 3.91 (m, 2H), 3.87 – 3.78 (m, 1H), 3.74 (d, *J* = 8.7 Hz, 1H), 3.48 (t, *J* = 10.7 Hz, 1H), 3.10 (s, 1H), 2.44 – 2.24 (m, 1H), 1.96 – 1.79 (m, 2H), 1.59 (s, 9H), 1.56 – 1.51 (m, 2H). **$^{13}$C-NMR** (126 MHz, $CDCl_3$, 298 K): δ (ppm) 154.4 (*C*=O), 152.5 (*C*=O), 148.5 ($C_{Ar}$), 141.3 ($C_{Ar}$), 130.3 ($C_{Ar}$), 127.7 ($C_{Ar}$H), 124.4 ($C_{Ar}$H), 122.7 ($C_{Ar}$H), 122.4 ($C_{Ar}$H), 120.2 ($C_{Ar}$), 109.4 ($C_{Ar}$H), 81.4 (*C*($CH_3$)$_3$), 73.4 (*C*$H_2$), 66.5 (*C*$H_2$), 54.5 (*C*H, major rotamer), 54.1 (*C*H, minor rotamer), 53.1 (*C*$H_2$), 42.2 (*C*H), 40.9 (*C*$H_2$, major rotamer), 40.5 (*C*$H_2$, minor rotamer), 36.3 (*C*H, major rotamer), 36.1 (*C*H, minor rotamer), 28.6 (C(*C*$H_3$)$_3$), 25.1 (*C*$H_2$), 23.9 (*C*$H_2$, major rotamer), 23.6 (*C*$H_2$, minor rotamer).

**P-CC60-CBI-Boc**

*N*-Boc (*S*)-1-(chloromethyl)-5-((cyclohexyl(3-(4-methylpiperazin-1-yl)-3-oxopropyl)carbamoyl)oxy)-1,2-dihydro-3*H*-benzo[*e*]indole

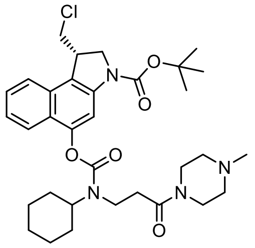

Carbamate coupling of **HO-CBI-Boc** (62.9 mg, 0.189 mmol) and 3-(cyclohexylamino)-1-(4-methylpiperazin-1-yl)propan-1-one dihydrochloride (92.6 mg, 0.284 mmol) was accomplished according to the **General Protocol A** to give **P-CC60-Boc** as a colourless solid (71.9 mg, 0.117 mmol, 63%).

**TLC** $R_f$ = 0.47 (DCM:MeOH, 9:1). **HRMS** (ESI$^+$) for $C_{33}H_{46}ClN_4O_5^+$: calcd. *m/z* 613.31512, found *m/z* 613.31534. **$^1$H-NMR** (600 MHz, MeOD-*d$_4$*): δ (ppm) = 7.94 (s, 1H), 7.91 – 7.83 (m, 2H), 7.58 (t, *J* = 7.4 Hz, 1H), 7.50 – 7.41 (m, 1H), 4.27 (t, *J* = 7.8 Hz, 1H), 4.20 (d, *J* = 7.1 Hz, 2H), 4.00 (d, *J* = 10.3 Hz, 1H), 3.91 (t, *J* = 12.5 Hz, 1H), 3.85 (d, *J* = 4.5 Hz, 1H), 3.78 – 3.67 (m, 3H), 3.66 – 3.61 (m, 1H), 2.95 (s, 1H), 2.88 (s, 1H), 2.84 – 2.77 (m, 2H), 2.74 – 2.62 (m, 2H), 2.57 (s, 2H), 2.44 (s, 1H), 2.08 (d, *J* = 10.8 Hz, 1H), 1.95 (d, *J* = 13.2 Hz, 1H), 1.90 (d, *J* = 10.6 Hz, 2H), 1.82 – 1.74 (m, 1H), 1.75 – 1.68 (m, 3H), 1.62 (s, 9H), 1.49 (q, *J* = 13.2 Hz, 1H), 1.28 – 1.19 (m, 1H). **$^{13}$C-NMR** (151 MHz, MeOD-*d$_4$*): δ (ppm) = 171.7 (*C*=O, rotamer 1), 171.5 (*C*=O rotamer 2), 156.3 (*C*=O), 153.8 (*C*=O), 149.5 ($C_{Ar}$), 131.7 ($C_{Ar}$), 128.8 ($C_{Ar}$H), 125.6 ($C_{Ar}$H), 125.4 ($C_{Ar}$), 123.8 ($C_{Ar}$H), 123.5 ($C_{Ar}$H), 122.2 ($C_{Ar}$), 121.4 ($C_{Ar}$), 110.0 ($C_{Ar}$H), 82.5 (*C*($CH_3$)$_3$), 59.0 (*C*H, major rotamer), 58.8 (*C*H, minor rotamer), 55.3 (*C*$H_2$, rotamer 1), 55.1 (*C*$H_2$, rotamer 2), 54.9 (*C*$H_2$, rotamer 3), 54.8 (*C*$H_2$, rotamer 4), 54.7 (*C*$H_2$, rotamer 5), 54.0 (*C*$H_2$), 47.7 (*C*$H_2$), 45.2 (*C*$H_2$), 45.1 (*C*$H_3$, minor rotamer), 44.8 (*C*$H_3$, major rotamer), 42.0 (*C*H), 41.7 (*C*$H_2$), 41.4 (*C*$H_2$), 35.2 (*C*$H_2$), 34.2 (*C*$H_2$), 32.6 (*C*$H_2$), 31.7 (*C*$H_2$), 28.7 (*C*$H_3$), 27.1 (*C*$H_2$), 26.5 (C(*C*$H_3$)$_3$), (*C*$H_2$).

**Methyl 5-hydroxy-1*H*-indole-2-carboxylate**

5-hydroxy-1*H*-indole-2-carboxylic acid (1.27 g, 7.15 mmol) was dissolved in MeOH (14 mL) and cooled to 0 °C. $SOCl_2$ (0.78 mL, 1.28 g, 10.7 mmol) was added dropwise and the resulting mixture was heated to 75°C for 2h, then cooled to 25°C and diluted with EtOAc (100 mL). The organic layer was washed with sat. aq. $NaHCO_3$ (1×) and sat. aq. NaCl (1×) and the combined aqueous layers extracted with EtOAc (3×50 mL). The combined organic layers were dried over $Na_2SO_4$, filtered, and concentrated under reduced pressure to give an orange crude mixture. Purification by FCC (isohexane/EtOAc) gave methyl 5-hydroxy-1*H*-indole-2-carboxylate as a colourless powder (1.32 g, 6.9 mmol, 97%).

**TLC** $R_f$ = 0.38 (isohexane:EtOAc, 5:1). **HRMS** ($ESI^+$) for $C_{10}H_{10}NO_3^+$: calcd. *m/z* 191.0582, found *m/z* 191.0578. **$^1$H-NMR** (500 MHz, $CDCl_3$): δ (ppm) = 8.75 (br-s, 1H), 7.30 (dd, *J* = 8.8, 0.8 Hz, 1H), 7.09 (dd, *J* = 2.1, 1.0 Hz, 1H), 7.06 (d, *J* = 2.4 Hz, 1H), 6.93 (dd, *J* = 8.8, 2.4 Hz, 1H), 4.60 (s, 1H), 3.93 (s, 3H). **$^{13}$C-NMR** (126 MHz, $CDCl_3$): δ (ppm) = 162.4 (*C*=O), 150.3 ($C_{Ar}$), 132.5 ($C_{Ar}$), 128.3 ($C_{Ar}$), 128.1 ($C_{Ar}$), 116.3 ($C_{Ar}$H), 112.9 ($C_{Ar}$H), 108.1 ($C_{Ar}$H), 106.1 ($C_{Ar}$H), 52.2 ($CH_3$). Analytical data matched literature reports.[14]

**AZI-OMe**

Methyl 5-(3-(azetidine-1-yl)propoxy-1*H*-indole-2-carboxylate

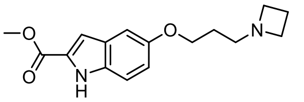

Route A:

**Step 1:** To methyl 5-hydroxy-1*H*-indole-2-carboxylate (190 mg, 1.00 mmol) in butanone (15 mL) was added $K_2CO_3$ (208 mg, 1.51 mmol) and 1-bromo-3-chloropropane (129 µL, 1.31 mmol). The mixture was stirred at 70°C for 24 h, then cooled to 25°C and poured into sat. aq. NaCl and diluted with EtOAc (50 mL). The layers were seperated and the aqueous layer was extracted with EtOAc (3×50 mL). The combined organic layers were dried over $Na_2SO_4$ and concentrated under reduced pressure. Purification by FCC (isohexane/EtOAc) gave methyl 5-(3-chloropropoxy)-1*H*-indole-2-carboxylate as a colourless, crystalline solid (190 mg, 0.71 mmol, 71%).

Intermediate: methyl 5-(3-chloropropoxy)-1*H*-indole-2-carboxylate

**TLC** $R_f$ = 0.30 (isohexane:EtOAc, 4:1). **HRMS** ($ESI^-$) for $C_{13}H_{13}NO_2Cl^-$: calcd. *m/z* 266.05894, found *m/z* 266.05929. **$^1$H-NMR** (400 MHz, $CDCl_3$): δ (ppm) = 8.81 (br-s, 1H), 7.32 (dt, *J* = 8.9, 0.8 Hz, 1H), 7.13 (dd, *J* = 2.1, 1.0 Hz, 1H), 7.10 (d, *J* = 2.4 Hz, 1H), 7.00 (dd, *J* = 8.9, 2.4 Hz, 1H), 4.15 (t, *J* = 5.8 Hz, 2H), 3.94 (s, 3H), 3.78 (t, *J* = 6.4 Hz, 2H), 2.27 (p, *J* = 6.1 Hz, 2H). **$^{13}$C-NMR** (101 MHz, $CDCl_3$): δ (ppm) = 162.4 (*C*=O), 153.9 ($C_{Ar}$), 132.4 ($C_{Ar}$), 128.0 ($C_{Ar}$), 127.8 ($C_{Ar}$), 117.5 ($C_{Ar}$H), 112.9 ($C_{Ar}$H), 108.5 ($C_{Ar}$H), 104.0 ($C_{Ar}$H), 65.1 ($CH_2$), 52.1 ($CH_3$), 41.8 ($CH_2$), 32.5 ($CH_2$).

**Step 2***:* To methyl 5-(3-chloropropoxy)-1*H*-indole-3-carboxylate (154 mg, 0.58 mmol) in MeCN (20 mL) was added $K_2CO_3$ (318 mg, 2.30 mmol), NaI (43 mg, 0.29 mmol), 15-crown-5 (76 mg, 0.35 mmol) and azetidine hydrochloride (161 mg, 1.73 mmol). The mixture was stirred at 70°C for 120 h, cooled to 25°C, filtered and washed with EtOAc (20 mL). The filtrate was concentrated giving a brown oil. Purification by FCC (EtOAc/MeOH) gave **AZI-OMe** as a colourless solid (177 mg, 0.62 mmol, 93%).

Route B:

$PPh_3$ (1.25 g, 4.78 mmol, 1.1 eq) was dissolved in anhydrous THF (0.25 M) under inert gas atmosphere, the resulting solution was cooled to 0°C and DIAD (0.95 mL, 4.78 mmol, 1.1 eq) was added. The mixture was stirred for 30 min and precipitation of a yellow solid was observed. A solution of methyl 5-hydroxy-1*H*-indole-2-carboxylate (0.99 g, 5.21 mmol, 0.5 M in anyhrous THF) and a solution of 3-(azetidine-1-yl)propanol (0.50 g, 4.34 mmol, 0.5 M in anhydrous THF) was added at 0°C. The mixture was allowed to warm to r.t. and was further stirred at r.t. for 15 h, before being concentrated under reduced pressure. Purification by FCC (EtOAc/MeOH) gave **AZI-OMe** as a colourless solid (0.98 g, 3.39 mmol, 78%).

**TLC** $R_f$ = 0.30 (isohexane:EtOAc, 4:1). **HRMS** ($ESI^-$) for $C_{16}H_{21}N_2O_2^-$: calcd. *m/z* 289.15467, found *m/z* 289.15478. **$^1$H-NMR** (400 MHz, $CDCl_3$): δ (ppm) = 8.89 (br-s, 1H), 7.29 (dt, *J* = 8.9, 0.8 Hz, 1H), 7.11 (dd, *J* = 2.1, 1.0 Hz, 1H), 7.06 (d, *J* = 2.4 Hz, 1H), 6.98 (dd, *J* = 8.9, 2.4 Hz, 1H), 4.02 (t, *J* = 6.4 Hz, 2H), 3.23 (t, *J* = 7.0 Hz, 4H), 2.61 (t, *J* = 7.2 Hz, 2H), 2.09 (p, *J* = 7.0 Hz, 2H), 1.93 – 1.79 (m, 2H). **$^{13}$C-NMR** (101 MHz, $CDCl_3$): δ (ppm) = 162.5 (*C*=O), 154.2 ($C_{Ar}$), 132.4 ($C_{Ar}$), 128.0 ($C_{Ar}$), 127.6 ($C_{Ar,}$), 117.6 ($C_{Ar}$H), 112.8 ($C_{Ar}$H), 108.5 ($C_{Ar}$H), 103.8 ($C_{Ar}$H), 66.7 ($CH_2$), 56.6 ($CH_2$), 55.4 ($CH_2$), 52.1 ($CH_3$), 27.7 ($CH_2$), 17.7 ($CH_2$).

**AZI-OH**

5-(3-(azetidine-1-yl)propoxy-1*H*-indole-2-carboxylic acid

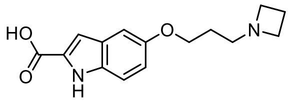


**AZI-OMe** (607 mg, 2.11 mmol) was dissolved in MeOH (100 mL) and solid KOH (167 mg, 2.53 mmol) was added. The resulting solution was heated to 75 °C for 15h, was cooled to r.t. and a solution of HCl (2.0 mL, 1.25 M in MeOH) was added. The resulting heterogenous mixture was concentrated under reduced pressure to obtain the product as a pure organic compound (quant., 68 w% calc. purity) contaminated with KCl. Purification of a fraction by RP chromatography ($H_2O$/MeCN) gave **AZI-OH** as a colourless powder (15 mg, 0.05 mmol).

**HRMS** (ESI$^-$) for $C_{13}H_{13}NO_2Cl^-$: calcd. *m/z* 275.13902, found *m/z* 275.13910. $^1$**H-NMR** (400 MHz, DMSO-*d*$_6$): δ (ppm) = 11.14 (br-s, 1H), 7.25 (d, *J* = 8.8 Hz, 1H), 7.08 (d, *J* = 2.4 Hz, 1H), 6.75 (dd, *J* = 8.8, 2.3 Hz, 2H), 4.02 (t, *J* = 6.8 Hz, 2H), 3.63 (t, *J* = 7.5 Hz, 4H), 2.91 (t, *J* = 6.5 Hz, 2H), 2.20 (p, *J* = 7.5 Hz, 2H), 1.86 (p, *J* = 6.7 Hz, 2H). $^{13}$**C-NMR** (101 MHz, DMSO-*d*$_6$): δ (ppm) = 165.1 (*C*=O), 152.6 ($C_{Ar}$), 134.1 ($C_{Ar}$), 131.7 ($C_{Ar}$H), 127.6 ($C_{Ar}$), 114.3 ($C_{Ar}$H), 112.9 ($C_{Ar}$H), 104.3 ($C_{Ar}$H), 102.6 ($C_{Ar}$H), 65.3 ($C$H$_2$), 53.2 ($C$H$_2$), 53.0 ($C$H$_2$), 25.4 ($C$H$_2$), 16.2 ($C$H$_2$).

#### *9.3. CBI-AZI prodrug series*

**SS60-CBI-AZI**

(*S*)-3-(5-(3-(azetidin-1-yl)propoxy)-1*H*-indole-2-carbonyl)-1-(chloromethyl)-2,3-dihydro-1*H*-benzo[*e*]indol-5-yl ((*S*)-1,2-dithian-4-yl)(methyl)carbamate

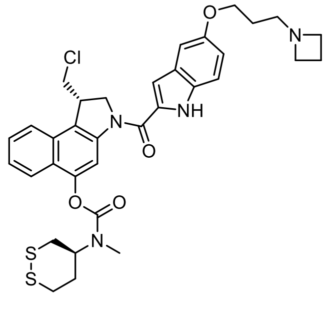


Amide coupling and was accomplished according to the **General Protocol B**. HCl (4 M in dioxane) (*Method A*) was applied for *N*-Boc deprotection of **SS60-CBI-Boc** (68.0 mg, 0.134 mmol) and **AZI-OH** (64.7 mg, 68 w%, 0.160 mmol) was activated for final prodrug assembly using EDCI (*Method D*). **SS60-CBI-AZI** was obtained as a colourless solid (43.6 mg, 0.655 mmol, 49%).

**TLC** $R_f$ = 0.50 (DCM:MeOH, 4:1). **HRMS** (ESI$^+$) for $C_{34}H_{38}ClN_4O_4S_2^+$: calcd. *m/z* 665.20175, found *m/z* 665.20156. $^1$**H-NMR** (400 MHz, DMSO-*d*$_6$): δ (ppm) = 11.59 (s, 1H), 8.21 (s, 1H), 8.04 (d, *J* = 8.3 Hz, 1H), 7.84 (d, *J* = 8.5 Hz, 1H), 7.66 – 7.58 (m, 1H), 7.55 – 7.48 (m, 1H), 7.39 (d, *J* = 8.9 Hz, 1H), 7.22 – 7.08 (m, 2H), 6.91 (dd, *J* = 8.9, 2.3 Hz, 1H), 4.88 (t, *J* = 10.1 Hz, 1H), 4.62 (d, *J* = 10.9 Hz, 1H), 4.48 – 4.36 (m, 1H), 4.20 – 4.05 (m, 2H), 4.05 – 3.92 (m, 3H), 3.26 – 3.17 (m, 3H), 3.16 (s, 2H), 3.09 (t, *J* = 6.9 Hz, 4H), 2.99 – 2.89 (m, 2H), 2.23 – 2.07 (m, 1H), 1.95 (p, *J* = 6.9 Hz, 2H), 1.71 (p, *J* = 6.6 Hz, 2H). $^{13}$**C-NMR** (126 MHz, DMSO-*d*$_6$): δ (ppm) = 160.2 (*C*=O), 153.4 (*C*=O), 153.1 ($C_{Ar}$), 147.3 ($C_{Ar}$), 141.5 ($C_{Ar}$), 131.7 ($C_{Ar}$), 130.6 ($C_{Ar}$), 129.5 ($C_{Ar}$), 127.6 ($C_{Ar}$), 127.5 ($C_{Ar}$H), 125.1 ($C_{Ar}$H), 124.3 ($C_{Ar}$), 123.4 ($C_{Ar}$H), 122.3 ($C_{Ar}$H), 116.0 ($C_{Ar}$H), 113.2 ($C_{Ar}$H), 110.7 ($C_{Ar}$H), 105.6 ($C_{Ar}$H), 103.1 ($C_{Ar}$H), 65.8 ($C$H$_2$), 55.7 ($C$H), 55.3 ($C$H$_2$), 55.0 ($C$H$_2$), 54.4 ($C$H$_2$), 47.7 ($C$H$_2$), 41.2 ($C$H), 35.6 ($C$H$_2$), 34.6 ($C$H$_2$), 31.7 ($C$H$_2$), 30.2 ($C$H$_3$), 26.7 ($C$H$_2$), 16.9 ($C$H$_2$).

**P-SS60-CBI-AZI**

(*S*)-3-(5-(3-(azetidin-1-yl)propoxy)-1*H*-indole-2-carbonyl)-1-(chloromethyl)-2,3-dihydro-1*H*-benzo[*e*]indol-5-yl ((*S*)-1,2-dithian-4-yl)(3-(4-methylpiperazin-1-yl)-3-oxopropyl)carbamate

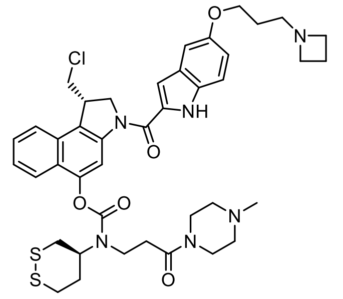


Amide coupling was accomplished according to the **General Protocol B**. $BF_3 \cdot OEt_2$ (*Method B*) was applied for *N*-Boc deprotection of **P-SS60-CBI-Boc** (22.0 mg, 0.034 mmol) and **AZI-OH** (26.5 mg, 68 w%, 0.051 mmol) was activated for final prodrug assembly using EDCI (*Method D*). **P-SS60-CBI-AZI** was obtained as a colourless solid (17.3 mg, 0.022 mmol, 65%).

**TLC** $R_f$ = 0.07 (DCM:MeOH, 5:1). **HRMS** ($ESI^+$) for $C_{41}H_{50}ClN_6O_5S_2^+$: calcd. *m/z* 805.29671, found *m/z* 805.29632. **$^1$H-NMR** (500 MHz, DMSO-*d*$_6$): δ (ppm) = 11.58 (s, 1H), 8.25 (s, 1H), 8.22 (s, 1H), 8.08 – 8.00 (m, 1H), 7.84 (t, *J* = 8.1 Hz, 1H), 7.66 – 7.59 (m, 1H), 7.56 – 7.48 (m, 1H), 7.40 (d, *J* = 8.9 Hz, 1H), 7.17 – 7.12 (m, 2H), 6.92 (dd, *J* = 8.9, 2.3 Hz, 1H), 4.87 (t, *J* = 10.0 Hz, 1H), 4.62 (d, *J* = 10.9 Hz, 1H), 4.41 (s, 1H), 4.09 (d, *J* = 11.1 Hz, 1H), 4.03 – 3.97 (m, 3H), 3.92 (s, 1H), 3.82 – 3.74 (m, 1H), 3.21 (t, *J* = 7.0 Hz, 3H), 3.15 – 3.11 (m, 1H), 3.01 (d, *J* = 10.9 Hz, 1H), 2.85 (dq, *J* = 16.0, 8.2 Hz, 1H), 2.67 (d, *J* = 12.8 Hz, 1H), 2.58 (t, *J* = 7.0 Hz, 1H), 3.33 – 3.27 (m, 2H), 2.31 – 2.21 (m, 5H), 2.21 – 2.08 (m, 5H), 2.00 (p, *J* = 7.0 Hz, 2H), 1.74 (p, *J* = 6.5 Hz, 2H). **$^{13}$C-NMR** (126 MHz, DMSO-*d*$_6$): δ (ppm) = 169.8 (*C*=O), 164.9 (*C*=O), 160.5 (*C*=O), 152.7 ($C_{Ar}$), 148.1 ($C_{Ar}$), 141.9 ($C_{Ar}$), 132.2 ($C_{Ar}$), 131.0 ($C_{Ar}$), 130.2 ($C_{Ar}$), 128.0 ($C_{Ar}$H), 125.8 ($C_{Ar}$H), 125.0 ($C_{Ar}$), 123.9 ($C_{Ar}$H), 122.8 ($C_{Ar}$H), 116.9 ($C_{Ar}$H), 115.1 ($C_{Ar}$), 113.7 ($C_{Ar}$H), 111.4 ($C_{Ar}$H), 106.7 ($C_{Ar}$H), 103.6 ($C_{Ar}$H), 66.3 ($CH_2$), 58.5 (*C*H), 55.7 ($CH_2$), 55.4 ($CH_2$), 55.2 ($CH_2$), 54.9 ($CH_2$), 54.7 ($CH_2$), 48.3 ($CH_2$), 46.5 ($CH_3$), 45.9 ($CH_2$), 42.9 ($CH_2$), 41.7 (*C*H), 41.3 ($CH_2$), 36.6 ($CH_2$), 35.6 ($CH_2$), 33.8 ($CH_2$), 32.5 ($CH_2$), 28.5 ($CH_2$), 22.5, 18.0 ($CH_2$).

**SeS60-CBI-AZI**

(*S*)-3-(5-(3-(azetidin-1-yl)propoxy)-1*H*-indole-2-carbonyl)-1-(chloromethyl)-2,3-dihydro-1*H*-benzo[*e*]indol-5-yl methyl((*S*)-1,2-thiaselenan-4-yl)carbamate

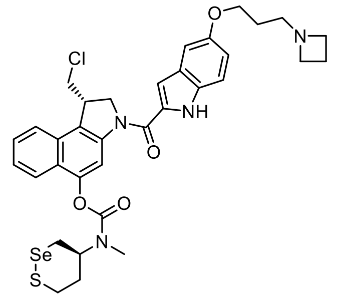


Amide coupling was accomplished according to the **General Protocol B**. HCl (4 M in dioxane) (*Method A*) was applied for *N*-Boc deprotection of **SeS60-CBI-Boc** (41.0 mg, 0.074 mmol) and **AZI-OH** (35.7 mg, 68 w%, 0.088 mmol) was activated for final prodrug assembly using EDCI (*Method D*). **SeS60-CBI-AZI** was obtained as a colourless solid (27.0 mg, 0.038 mmol, 51%).

**TLC** $R_f$ = 0.57 (DCM:MeOH, 4:1). **HRMS** ($ESI^+$) for $C_{34}H_{38}ClN_4O_4SSe^+$: calcd. *m/z* 713.14520, found *m/z* 713.14594. **$^1$H-NMR** (400 MHz, DMSO-*d*$_6$): δ (ppm) = 11.64 (s, 1H), 9.86 (s, 1H), 8.21 (s, 1H), 8.04 (d, *J* = 8.5 Hz, 1H), 7.84 (d, *J* = 8.8 Hz, 1H), 7.61 (q, *J* = 7.3 Hz, 1H), 7.52 (t, *J* = 7.8 Hz, 1H), 7.47 (d, *J* = 7.9 Hz, 1H, $TsO^-$), 7.43 (d, *J* = 9.0 Hz, 1H), 7.18 (s, 1H), 7.14 (s, 1H), 7.11 (d, *J* = 8.0 Hz, 1H, $TsO^-$), 6.95 (dd, *J* = 8.9, 2.6 Hz, 1H), 4.96 – 4.78 (m, 1.5H, major rotamer), 4.61 (d, *J* = 9.8 Hz, 1H), 4.56 – 4.51 (m, 0.5H, minor rotamer), 4.41 (s, 1H), 4.33 – 4.22 (m, 1H), 4.14 – 3.97 (m, 6H), 3.56 – 3.39 (m, 1H), 3.25 – 2.95 (m, 1H), 3.15 (s, 2H, major rotamer), 2.92 (s, 1H, minor rotamer,), 2.41 – 2.30 (m, 2H), 2.28 (s, 1.5H, $TsO^-$), 2.23 – 2.00 (m, 2H), 1.95 (p, *J* = 6.5 Hz, 2H). **$^{13}$C-NMR** (126 MHz, DMSO-*d*$_6$): δ (ppm) = 160.2 (*C*=O), 152.7 (*C*=O), 150.7 ($C_{Ar}$), 147.3 ($C_{Ar}$), 145.8 ($C_{Ar}$, $TsO^-$), 142.8 ($C_{Ar}$), 141.4 ($C_{Ar}$), 137.5 ($C_{Ar}$), 135.6 ($C_{Ar}$, $TsO^-$), 131.8 ($C_{Ar}$), 130.8 ($C_{Ar}$), 129.5 ($C_{Ar}$), 128.0 ($C_{Ar}$H, $TsO^-$), 127.7 ($C_{Ar}$), 127.5 ($C_{Ar}$H), 125.5 ($C_{Ar}$H, $TsO^-$), 124.3 ($C_{Ar}$H), 123.7 ($C_{Ar}$H), 122.4 ($C_{Ar}$H), 117.3 ($C_{Ar}$), 115.9 ($C_{Ar}$H), 113.3 ($C_{Ar}$H), 110.4 ($C_{Ar}$H), 105.5 ($C_{Ar}$H), 103.4 ($C_{Ar}$H), 64.9 ($CH_2$), 56.0 (*C*H), 54.8 ($CH_2$), 54.0 ($CH_2$), 51.8 ($CH_2$),

47.2 ($CH_2$), 41.1 ($CH$), 32.9 ($CH_2$), 30.0 ($CH_2$), 26.6 ($CH_2$), 24.3 ($CH_2$), 20.8 ($CH_3$, $TsO^-$), 15.9 ($CH_2$). 0.5 eq. $TsO^-$: counter ion, 0.5 eq identified by NMR.

**Me-SS66T-CBI-AZI**

(*S*)-3-(5-(3-(azetidin-1-yl)propoxy)-1*H*-indole-2-carbonyl)-1-(chloromethyl)-2,3-dihydro-1*H*-benzo[*e*]indol-5-yl (*trans*)-4-methylhexahydro-[1,2]dithiino[4,5-*b*]pyrazine-1(2*H*)-carboxylate

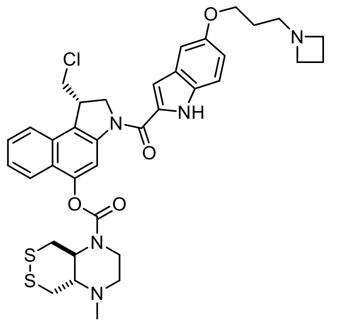

Amide coupling was accomplished according to the **General Protocol B**. $BF_3{\cdot}OEt_2$ (*Method B*) was applied for *N*-Boc deprotection of **Me-SS66T-CBI-Boc** (12.0 mg, 0.021 mmol) and **AZI-OH** (17.0 mg, 68 w%, 0.033 mmol) was activated for final prodrug assembly using EDCI (*Method D*). **Me-SS66T-CBI-AZI** was obtained as a colourless solid (2.0 mg, 0.003 mmol, 14%).

.**TLC** $R_f$ = 0.1 (DCM:MeOH 5:1). **HRMS** ($ESI^+$) for $C_{36}H_{41}ClN_5O_4S_2^+$: calcd. *m/z* 706.22830, found *m/z* 706.22814. **$^1$H-NMR** (800 MHz, DMSO-$d_6$): δ (ppm) = 11.61 (s, 1H), 8.21 (s, 1H), 8.05 (d, *J* = 8.3 Hz, 1H), 7.84 (d, *J* = 8.4 Hz, 1H), 7.65 – 7.60 (m, 1H), 7.58 – 7.51 (m, 1H), 7.47 (d, *J* = 8.0 Hz, 1H), 7.39 (d, *J* = 8.8 Hz, 1H), 7.16 – 7.12 (m, 2H), 7.11 (d, *J* = 7.8 Hz, 1H), 6.91 (dd, *J* = 8.8, 2.4 Hz, 1H), 4.87 (t, *J* = 9.4 Hz, 1H), 4.62 (dd, *J* = 10.6, 2.0 Hz, 1H), 4.42 (d, *J* = 2.7 Hz, 1H), 4.10 (dt, *J* = 11.2, 3.0 Hz, 1H), 4.00 (dt, *J* = 15.4, 6.6 Hz, 4H), 3.93 (s, 1H), 3.53 – 3.46 (m, 2H), 3.22 – 3.17 (m, 5H), 3.01 – 2.94 (m, 1H), 2.89 (d, *J* = 8.6 Hz, 1H), 2.71 – 2.65 (m, 2H), 2.58 – 2.55 (m, 1H), 2.33 – 2.27 (m, 3H), 1.99 (p, *J* = 7.0 Hz, 2H), 1.74 (p, *J* = 6.6 Hz, 2H). **$^{13}$C-NMR** (201 MHz, DMSO-$d_6$): δ (ppm) = 163.6 ($C$=O), 160.3 ($C$=O), 153.1 ($C_{Ar}$), 147.0 ($C_{Ar}$), 145.9 ($C_{Ar}$), 141.5 ($C_{Ar}$), 137.5 ($C_{Ar}$), 131.7 ($C_{Ar}$), 130.6 ($C_{Ar}$), 129.6 ($C_{Ar}$), 128.0 ($C_{Ar}$), 127.7 ($C_{Ar}H$), 127.5 ($C_{Ar}H$), 125.5 ($C_{Ar}H$), 125.2 ($C_{Ar}H$), 124.3 ($C_{Ar}$), 123.5 ($C_{Ar}H$), 122.5 ($C_{Ar}$), 122.1 ($C_{Ar}H$), 116.1 ($C_{Ar}H$), 113.2 ($C_{Ar}H$), 110.6 ($C_{Ar}H$), 105.6 ($C_{Ar}H$), 103.1 ($C_{Ar}H$), 65.8 ($CH_2$), 62.2 ($CH$), 61.9 ($CH$), 55.4 ($CH_2$), 54.9 ($CH_2$), 54.5 (2×$CH_2$), 47.7 ($CH_2$), 43.5 ($CH_2$), 41.2 ($CH$), 38.2 ($CH_2$), 36.7 ($CH_2$), 26.8 ($CH_2$), 20.8 ($CH_3$), 17.0 ($CH_2$).

**Me-SS66C-CBI-AZI**

(*S*)-3-(5-(3-(azetidin-1-yl)propoxy)-1*H*-indole-2-carbonyl)-1-(chloromethyl)-2,3-dihydro-1*H*-benzo[*e*]indol-5-yl (*cis*)-4-methylhexahydro-[1,2]dithiino[4,5-*b*]pyrazine-1(2*H*)-carboxylate

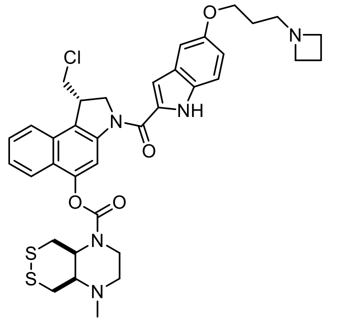

Amide coupling was accomplished according to the **General Protocol B**. $BF_3{\cdot}OEt_2$ (*Method B*) was applied for *N*-Boc deprotection of **Me-SS66C-CBI-Boc** (21.0 mg, 0.038 mmol) and **AZI-OH** (36.6 mg, 68 w%, 0.105 mmol) was activated for final prodrug assembly using EDCI (*Method D*). **Me-SS66C-CBI-AZI** was obtained as a colourless solid (2.0 mg, 0.003 mmol, 8%).

**TLC** $R_f$ = 0.1 (DCM:MeOH 5:1). **HRMS** ($ESI^+$) for $C_{36}H_{41}ClN_5O_4S_2^+$: calcd. *m/z* 706.22830, found *m/z* 706.22798. **$^1$H-NMR** (800 MHz, DMSO-$d_6$): δ (ppm) = 11.58 (s, 1H), 8.29 (s, 1H), 8.20 (s, 1H), 8.04 (d, *J* = 8.3 Hz, 1H), 7.87 – 7.80 (m, 1H), 7.62 (t, *J* = 7.5 Hz, 1H), 7.55 – 7.50 (m, 1H), 7.39 (d, *J* = 8.8 Hz, 1H), 7.16 – 7.11 (m, 2H), 6.91 (dd, *J* = 8.8, 2.4 Hz, 1H), 4.87 (t, *J* = 9.4 Hz, 1H), 4.62 (d, *J* = 10.6 Hz, 1H), 4.41 (s, 0H), 4.16 (dd, *J* = 30.6, 12.1 Hz, 1H), 4.09 (d, *J* = 11.2 Hz, 1H), 4.02 – 3.96 (m, 3H), 3.94 – 3.89 (m, 1H), 3.87 – 3.78 (m, 2H), 3.64 – 3.55 (m, 1H), 3.51 – 3.43 (m, 1H), 3.11 (t, *J* = 6.9 Hz, 4H), 2.96 (dd, *J* = 26.2, 10.8 Hz, 1H), 2.86 – 2.78 (m, 1H), 2.41 (dt, *J* = 3.6,

1.8 Hz, 2H), 2.34 – 2.25 (m, 1H), 2.21 (s, 3H), 1.99 – 1.92 (m, 2H), 1.72 (p, *J* = 6.7 Hz, 2H), 1.28 – 1.23 (m, 2H). **$^{13}$C-NMR** (201 MHz, DMSO-$d_6$): δ (ppm) = 160.8 (*C*=O), 153.1 (*C*=O), 147.1 ($C_{Ar}$), 141.5 ($C_{Ar}$), 131.7 ($C_{Ar}$), 130.6 ($C_{Ar}$), 129.5 ($C_{Ar}$), 127.7 ($C_{Ar}$H, *rotamer 1*), 127.5 ($C_{Ar}$H, *rotamer 2*), 125.2 ($C_{Ar}$H), 124.2 ($C_{Ar}$H), 123.3 ($C_{Ar}$), 122.5 ($C_{Ar}$H, *rotamer 1*), 122.2 ($C_{Ar}$H, *rotamer 2*), 116.0 ($C_{Ar}$H), 113.2 ($C_{Ar}$H), 110.7 ($C_{Ar}$H), 105.6 ($C_{Ar}$H), 103.1 ($C_{Ar}$H), 65.9 ($CH_2$), 60.6 (*C*H, *rotamer 1*), 60.4 (*C*H, *rotamer 2*), 55.7 ($CH_2$), 55.4 ($CH_2$), 55.2 ($CH_2$, *rotamer 1*), 55.0 ($CH_2$, *rotamer 2*), 54.5 ($CH_2$), 54.3 (*C*H), 53.5 (*C*H), 47.7 ($CH_2$), 41.2 ($CH_3$), 40.5 ($CH_2$), 38.1 ($CH_2$), 29.0 ($CH_2$), 27.0 ($CH_2$), 17.0 ($CH_2$).

**SS00$_M$-CBI-AZI**

(*S*)-3-(5-(3-(azetidin-1-yl)propoxy)-1*H*-indole-2-carbonyl)-1-(chloromethyl)-2,3-dihydro-1*H*-benzo [*e*]indol-5-yl methyl(2-((2-morpholinoethyl)disulfaneyl)ethyl)carbamate

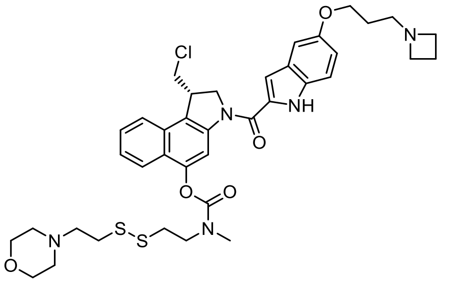

Amide coupling was accomplished according to the **General Protocol B**. HCl (4 M in dioxane) (*Method A*) was applied for *N*-Boc deprotection of **SS00$_M$-CBI-Boc** (112.0 mg, 0.188 mmol) and **AZI-OH** (90.9 mg, 68 w%, 0.225 mmol) was activated for final prodrug assembly using EDCI (*Method D*). **SS00$_M$-CBI-AZI** was obtained as a colourless solid (30 mg, 0.040 mmol, 21%).

**TLC** $R_f$ = 0.23 (DCM:MeOH, 5:1). **HRMS** ($ESI^+$) for $C_{38}H_{47}ClN_5O_5S_2^+$: calcd. *m/z* 752.27037, found *m/z* 752.27017. **$^1$H-NMR** (400 MHz, DMSO-$d_6$): δ (ppm) = 11.61 (s, 1H), 8.38 (s, 1H), 8.23 (s, 1H), 8.04 (d, *J* = 8.3 Hz, 1H), 8.01 – 7.82 (m, 1H), 7.62 (t, *J* = 7.5 Hz, 1H), 7.55 – 7.47 (m, 1H), 7.39 (d, *J* = 8.9 Hz, 1H), 7.18 – 7.08 (m, 2H), 6.91 (dd, *J* = 8.9, 2.4 Hz, 1H), 4.87 (t, *J* = 9.9 Hz, 1H), 4.62 (d, *J* = 11.0 Hz, 1H), 4.41 (s, 1H), 4.10 (dd, *J* = 11.0, 2.9 Hz, 1H), 3.99 (q, *J* = 6.5 Hz, 3H), 3.89 (s, 1H), 3.64 (d, *J* = 6.9 Hz, 1H), 3.51 (d, *J* = 4.4 Hz, 2H), 3.47 – 3.42 (m, 2H), 3.26 (s, 2H), 3.15 (t, *J* = 6.9 Hz, 1H), 3.10 (t, *J* = 6.9 Hz, 4H), 3.07 – 2.98 (m, 3H), 2.90 (t, *J* = 8.8 Hz, 2H), 2.56 (dd, *J* = 14.7, 5.5 Hz, 2H), 2.39 – 2.32 (m, 2H), 2.28 (d, *J* = 9.7 Hz, 2H), 1.95 (p, *J* = 6.9 Hz, 2H), 1.71 (p, *J* = 6.6 Hz, 2H). **$^1$H/$^{13}$C-HSQC** (400 MHz/101 MHz, DMSO-$d_6$): δ (ppm) = 127.2 ($C_{Ar}$H), 124.6 ($C_{Ar}$H), 123.3 ($C_{Ar}$H), 122.0 ($C_{Ar}$H), 115.9 ($C_{Ar}$H), 113.0 ($C_{Ar}$H), 110.4 ($C_{Ar}$H), 105.2 ($C_{Ar}$H), 102.6 ($C_{Ar}$H), 65.7 ($CH_2$), 65.4 ($CH_2$), 57.0 ($CH_2$), 55.4 ($CH_2$), 54.7 ($CH_2$), 54.1 ($CH_2$), 52.8 ($CH_2$), 47.6 ($CH_2$), 47.3 ($CH_2$), 40.8 (CH), 35.0 ($CH_2$), 43.3 ($CH_3$), 26.6 ($CH_2$), 16.5 ($CH_2$).

**MC-CBI-AZI**

(*S*)-3-(5-(3-(azetidin-1-yl)propoxy)-1*H*-indole-2-carbonyl)-1-(chloromethyl)-2,3-dihydro-1*H*-benzo[*e*]indol-5-yl methyl carbonate

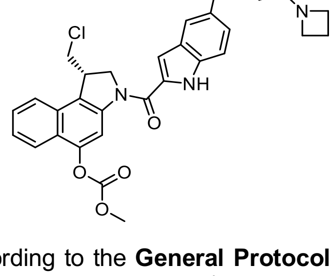

Amide coupling was accomplished according to the **General Protocol B**. $BF_3 \cdot OEt_2$ (*Method B*) was applied for *N*-Boc deprotection of **MC-CBI-Boc** (20.0 mg, 0.051 mmol) and **AZI-OH** (31.3 mg, 68 w%, 0.077 mmol) was activated for final prodrug assembly using EDCI (*Method D*). **MC-CBI-AZI** was obtained as a colourless solid (21.0 mg, 0.038 mmol, 75%).

**TLC** $R_f$ = 0.27 (DCM:MeOH, 5:1). **HRMS** ($ESI^+$) for $C_{30}H_{31}ClN_3O_5^+$: calcd. *m/z* 548.19468, found *m/z* 548.19519. **$^1$H-NMR** (500 MHz, DMSO-$d_6$): δ (ppm) = 11.64 (s, 1H), 8.34 (s, 1H), 8.26 (s, 1H), 8.07 (d, *J* = 8.3 Hz, 1H), 7.89 (d, *J* = 8.4 Hz, 1H), 7.70 – 7.61 (m, 1H), 7.58 – 7.51 (m, 1H), 7.41 (d, *J* = 8.8 Hz, 1H), 7.19 – 7.12 (m, 2H), 6.93 (d, *J* = 8.7 Hz, 1H), 4.88 (t, *J* = 10.0 Hz, 2H), 4.63 (d, *J* = 10.5 Hz, 2H), 4.43 (s, 2H), 4.10 (d, *J* = 8.9 Hz, 1H), 4.05 – 3.97 (m, 2H), 3.92 (s, 4H), 3.46 – 3.34 (m, 4H), 2.74 (d, *J* = 6.6 Hz, 2H), 2.15 – 2.01 (m, 2H), 1.89 – 1.69 (m, 2H). **$^{13}$C-NMR** (101 MHz, DMSO-$d_6$): δ (ppm) = 164.2 (*C*=O), 160.3 (*C*=O), 153.7 ($C_{Ar}$), 153.0 ($C_{Ar}$), 146.6 ($C_{Ar}$), 141.5

($C_{Ar}$), 131.8 ($C_{Ar}$), 130.5 ($C_{Ar}$), 129.6 ($C_{Ar}$), 127.9 ($C_{Ar}$), 127.5 ($C_{Ar}$H), 125.5 ($C_{Ar}$H), 123.6 ($C_{Ar}$), 123.5 ($C_{Ar}$H), 121.8 ($C_{Ar}$H), 116.1 ($C_{Ar}$H), 113.2 ($C_{Ar}$H), 110.2 ($C_{Ar}$H), 105.7 ($C_{Ar}$H), 103.2 ($C_{Ar}$H), 65.6 ($CH_2$), 55.9 ($CH_3$), 55.0 ($CH_2$), 54.4 ($CH_2$), 54.2 ($CH_2$), 47.6 ($CH_2$), 41.3 (CH), 26.1 ($CH_2$), 16.6 ($CH_2$).

**O56-CBI-AZI**

(*S*)-3-(5-(3-(azetidin-1-yl)propoxy)-1*H*-indole-2-carbonyl)-1-(chloromethyl)-2,3-dihydro-1*H*-benzo[*e*]indol-5-yl (*cis*)-hexahydrofuro[3,4-*b*]pyridine-1(2*H*)-carboxylate

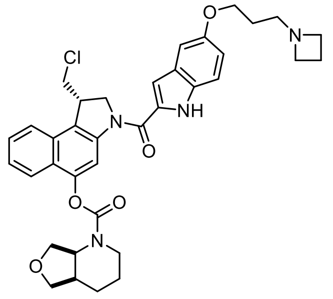


Amide coupling was accomplished according to the **General Protocol B**. HCl (4 M in dioxane) (*Method A*) was applied for *N*-Boc deprotection of **O56-CBI-Boc** (189 mg, 0.388 mmol) and **AZI-OH** (167 mg, 68 w%, 0.407 mmol) was activated for final prodrug assembly using EDCI (*Method D*). **O56-CBI-AZI** was obtained as a colourless solid (50.0 mg, 0.078 mmol, 20%).

**TLC** $R_f$ = 0.32 (DCM:MeOH, 5:1). **HRMS** ($ESI^+$) for $C_{36}H_{40}ClN_4O_5^+$: calcd. *m/z* 643.26817, found *m/z* 643.26776. **$^1$H-NMR** (400 MHz, DMSO-*d₆*): δ (ppm) = 11.65 (m, 1H), 9.95 (s, 1H), 8.20 (s, 1H), 8.04 (d, *J* = 8.4 Hz, 1H), 7.87 – 7.77 (m, 1H), 7.67 – 7.58 (m, 1H), 7.52 (t, *J* = 7.8 Hz, 1H) , 7.47 (d, *J* = 7.9 Hz, 1H, $TsO^-$), 7.43 (d, *J* = 8.9 Hz, 1H), 7.18 (d, *J* = 2.4 Hz, 1H), 7.14 (d, *J* = 2.4 Hz, 1H), 7.11 (d, *J* = 8.0 Hz, 1H, $TsO^-$), 6.95 (dt, *J* = 8.9, 2.9 Hz, 1H), 5.02 (s, 0.5H, major rotamer), 4.94 – 4.79 (m, 1H), 4.68 (s, 0.5H, major rotamer), 4.62 (dd, *J* = 10.9, 2.3 Hz, 1H), 4.41 (s, 1H), 4.34 – 4.17 (m, 1H), 4.05 (m, 8H), 3.85 (s, 3H), 3.62 (d, *J* = 8.5 Hz, 2H), 3.32 – 3.27 (m, 2H), 3.00 (s, 1H), 2.39 – 2.30 (m, 3H), 2.28 (s, 1.5H, $TsO^-$), 1.95 (p, *J* = 6.5 Hz, 2H), 1.87 – 1.75 (m, 2H), 1.67 – 1.42 (m, 2H). **$^{13}$C-NMR** (101 MHz, DMSO-$d_6$): δ (ppm) = 160.2 ($C_{Ar}$), 153.7 ($C_{Ar}$), 152.7 ($C_{Ar}$), 147.3 ($C_{Ar}$), 145.8 ($C_{Ar}$, $TsO^-$), 141.5 ($C_{Ar}$), 137.6 ($C_{Ar}$, $TsO^-$), 132.5 ($C_{Ar}$), 131.9 ($C_{Ar}$), 131.6 ($C_{Ar}$), 130.8 ($C_{Ar}$), 129.5 ($C_{Ar}$), 128.1 ($C_{Ar}$H, $TsO^-$), 127.7 ($C_{Ar}$), 127.5 ($C_{Ar}$H), 125.9 ($C_{Ar}$H, $TsO^-$), 124.8 ($C_{Ar}$H), 123.6 ($C_{Ar}$H), 122.4 ($C_{Ar}$H), 116.0 ($C_{Ar}$H), 113.3 ($C_{Ar}$H), 110.4 ($C_{Ar}$H), 105.6 ($C_{Ar}$H), 103.3 ($C_{Ar}$H), 72.5 ($CH_2$), 65.1 ($CH_2$), 64.9 ($CH_2$), 55.0 ($CH_2$), 53.9 ($CH_2$), 53.4 (CH) 51.8 ($CH_2$), 47.7 ($CH_2$), 41.3 (CH), 40.2 ($CH_2$), 35.0 (CH), 24.4 ($CH_2$), 24.3 ($CH_2$), 23.1 ($CH_2$), 20.8 ($CH_3$, $TsO^-$), 15.9 ($CH_2$). $TsO^-$: counter ion, 0.5 eq identified by NMR.

**OBn-CBI-AZI**

(*S*)-(5-(3-(azetidin-1-yl)propoxy)-1*H*-indol-2-yl)(5-(benzyloxy)-1-(chloromethyl)-1,2-dihydro-3*H*-benzo[e]indol-3-yl)methanone

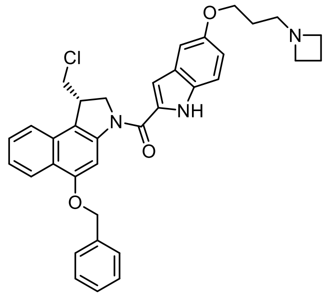


Amide coupling was accomplished according to the **General Protocol B**. HCl (4 M in dioxane) (*Method A*) was applied for *N*-Boc deprotection of **OBn-CBI-Boc** (116.0 mg, 0.274 mmol) and **AZI-OH** (110.0 mg, 68 w%, 0.274 mmol) was activated for final prodrug assembly using EDCI (*Method D*). **OBn-CBI-AZI** was obtained as a colourless solid (90.0 mg, 0.155 mmol, 57%).

**TLC** $R_f$ = 0.32 (DCM:MeOH, 5:1). **HRMS** ($ESI^+$) for $C_{35}H_{35}ClN_3O_3^+$: calcd. *m/z* 580.23615, found *m/z* 580.23615. **$^1$H-NMR** (400 MHz, DMSO-$d_6$): δ (ppm) = 11.70 (s, 1H), 10.39 (br-s, 1H), 8.22 (d, *J* = 8.4 Hz, 1H), 8.12 (br-s, 1H), 7.94 (d, *J* = 8.4 Hz, 1H), 7.65 – 7.53 (m, 3H), 7.49 – 7.31 (m, 5H), 7.18 (d, *J* = 2.4 Hz, 1H), 7.12 (d, *J* = 2.1 Hz, 1H), 6.94 (dd, *J* = 8.9, 2.4 Hz, 1H), 5.32 (s, 2H), 4.82 (t, *J* = 10.1 Hz, 1H), 4.58 (dd, *J* = 10.9, 2.2 Hz, 1H), 4.30 (td, *J* = 6.9, 3.0 Hz, 1H), 4.13 – 3.96 (m, 7H), 3.92 (dd, *J* = 11.1, 7.1 Hz, 1H), 3.28 (t, *J* = 7.5 Hz, 2H), 2.37 – 2.29 (m, 2H), 2.03 – 1.89 (m, 2H). **$^{13}$C-NMR** (101 MHz, DMSO-$d_6$): δ (ppm) = 160.2 ($C_{Ar}$), 154.4 ($C$=O), 153.2 ($C_{Ar}$), 142.0

($C_{Ar}$), 136.8 ($C_{Ar}$), 131.8 ($C_{Ar}$), 131.0 ($C_{Ar}$), 129.7 ($C_{Ar}$), 128.6 ($C_{Ar}$H), 128.0 ($C_{Ar}$), 127.7 ($C_{Ar}$H), 127.5 ($C_{Ar}$H), 125.5 ($C_{Ar}$), 124.1 ($C_{Ar}$), 123.0 ($C_{Ar}$), 122.7 ($C_{Ar}$H), 122.6 ($C_{Ar}$H), 116.7 ($C_{Ar}$), 116.0 ($C_{Ar}$H), 113.2 ($C_{Ar}$H), 105.9 ($C_{Ar}$H), 103.2 ($C_{Ar}$H), 98.3 ($C_{Ar}$H), 70.5 ($CH_2$), 64.9 ($CH_2$), 54.9 ($CH_2$), 53.8 ($CH_2$), 51.7 ($CH_2$), 47.7 ($CH_2$), 41.2 ($C$H), 25.9 ($CH_2$), 15.6 ($CH_2$).

#### *9.4. CBI-TMI prodrug series*

**P-SS60-CBI-TMI**

(*S*)-1-(chloromethyl)-3-(5,6,7-trimethoxy-1*H*-indole-2-carbonyl)-2,3-dihydro-1*H*-benzo[*e*]indol-5-yl (1,2-dithian-4-yl)(3-(4-methylpiperazin-1-yl)-3-oxopropyl)carbamate

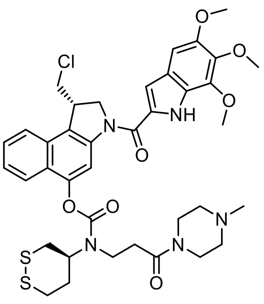

Amide coupling and was accomplished according to the **General Protocol B**. TFA (*Method C*) was applied for *N*-Boc deprotection of **P-SS60-CBI-Boc** (43.0 mg, 0.066 mmol) and **TMI-OH** (77.9 mg, 0.310 mmol) was activated for final prodrug assembly using oxaxylchloride/DMF (*Method E*). **P-SS60-CBI-TMI** was obtained as a colourless solid (42.0 mg, 0.054 mmol, 82%).

**TLC** $R_f$ = 0.58 (DCM:MeOH, 9:1). **HRMS** ($ESI^+$) for $C_{38}H_{45}ClN_5O_7S_2^+$: calcd. *m/z* 782.24434, found *m/z* 782.24486. **$^1$H-NMR** (800 MHz, MeOD-d*4*): δ (ppm) = 8.17 (s, 1H), 7.95 (dd, *J* = 8.5, 3.4 Hz, 1H), 7.89 (dd, *J* = 27.3, 8.4 Hz, 1H), 7.61 (t, *J* = 7.4 Hz, 1H), 7.56 – 7.46 (m, 1H), 7.10 (s, 1H), 6.99 (s, 1H), 4.74 (d, *J* = 7.6 Hz, 2H), 4.62 (s, 1H), 4.30 (tt, *J* = 7.5, 3.3 Hz, 1H), 4.07 (d, *J* = 3.5 Hz, 3H), 4.05 – 4.02 (m, 1H), 3.92 – 3.90 (m, 8H), 3.80 (ddt, *J* = 16.3, 12.5, 6.0 Hz, 1H), 3.72 – 3.65 (m, 1H), 3.62 (s, 4H), 3.49 – 3.38 (m, 2H), 3.29 – 3.04 (m, 2H), 2.99 – 2.90 (m, 2H), 2.84 – 2.75 (m, 1H), 2.54 – 2.49 (m, 2H), 2.48 – 2.43 (m, 2H), 2.40 – 2.34 (m, 2H), 2.34 – 2.22 (m, 3H). **$^{13}$C-NMR** (201 MHz, MeOD-d*4*): δ (ppm) = 171.5 ($C$=O), 171.4 ($C$=O), 162.7 ($C$=O), 155.7 ($C$=O), 155.5 ($C$=O), 155.1 ($C$=O), 151.2 ($C_{Ar}$), 148.9 ($C_{Ar}$), 142.6 ($C_{Ar}$), 141.7 ($C_{Ar}$), 140.4 ($C_{Ar}$), 131.4 ($C_{Ar}$), 128.9 ($C_{Ar}$H), 127.4 ($C_{Ar}$), 126.4 ($C_{Ar}$), 126.3 ($C_{Ar}$H), 125.2 ($C_{Ar}$), 124.4 ($C_{Ar}$), 124.3 ($C_{Ar}$H), 123.6 ($C_{Ar}$H), 112.1 ($C_{Ar}$H), 108.2 ($C_{Ar}$), 107.9 ($C_{Ar}$H), 99.3 ($C_{Ar}$H), 61.9 ($CH_3$), 61.7 ($CH_3$), 60.1 ($C$H), 56.8 ($CH_3$), 56.7 ($CH_2$), 55.8 ($CH_2$), 55.3 ($CH_2$), 47.5 ($CH_2$), 46.3 ($CH_2$), 45.8 ($CH_2$), 44.0 ($CH_2$), 43.5 ($C$H), 42.2 ($CH_2$), 37.6 ($CH_2$), 36.9 ($CH_2$), 35.1 ($CH_2$), 33.9 ($CH_2$).

**Me-SS66T-CBI-TMI**

(*S*)-1-(chloromethyl)-3-(5,6,7-trimethoxy-1*H*-indole-2-carbonyl)-2,3-dihydro-1*H*-benzo[*e*]indol-5-yl (*trans*)-4-methylhexahydro-[1,2]dithiino[4,5-*b*]pyrazine-1(2*H*)-carboxylate

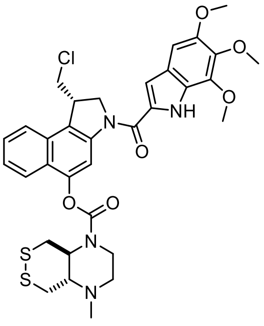

Amide coupling and was accomplished according to the **General Protocol B**. $BF_3{\cdot}OEt_2$ (*Method B*) was applied for *N*-Boc deprotection of **Me-SS66T-CBI-Boc** (20.0 mg, 0.036 mmol) and **TMI-OH** (22.8 mg, 0.091 mmol) was activated for final prodrug assembly using oxaxylchloride/DMF (*Method E*). **Me-SS66T-CBI-TMI** was obtained as a colourless solid (19.2 mg, 0.028 mmol, 78%).

**TLC** $R_f$ = 0.32 (DCM:MeOH, 20:1). **HRMS** ($ESI^+$) for $C_{33}H_{36}ClN_4O_6S_2^+$: calcd. *m/z* 683.17593, found *m/z* 683.17534. **$^1$H-NMR** (400 MHz, $CDCl_3$): δ (ppm) = 9.41 (s, 1H), 8.39 (s, 1H), 7.87 (dd, *J* = 8.3, 4.6 Hz, 1H), 7.77 (d, *J* = 8.3 Hz, 1H), 7.55 (t, *J* = 7.4 Hz, 1H), 7.50 – 7.41 (m, 1H), 7.01 (d, *J* = 2.1 Hz, 1H), 6.88 (s, 1H), 4.81 (dd,

$J$ = 10.7, 1.6 Hz, 1H), 4.67 (t, $J$ = 9.5 Hz, 1H), 4.18 (dd, $J$ = 10.8, 5.9 Hz, 1H), 4.08 (s, 3H), 4.02 – 3.96 (m, 2H), 3.95 (s, 3H), 3.92 (s, 3H), 3.92 – 3.86 (m, 1H), 3.75 (d, $J$ = 10.4 Hz, 1H), 3.57 – 3.45 (m, 2H), 3.31 (d, $J$ = 12.2 Hz, 1H), 3.28 – 3.17 (m, 2H), 3.12 – 3.01 (m, 1H), 2.96 (t, $J$ = 10.0 Hz, 1H), 2.84 (dt, $J$ = 10.3, 4.6 Hz, 1H), 2.44 (s, 3H). **$^{13}$C-NMR** (126 MHz, $CDCl_3$): δ (ppm) = 160.4 (*C*=O), 153.6 (*C*=O), 150.4 ($C_{Ar}$), 148.0 ($C_{Ar}$), 141.5 ($C_{Ar}$), 140.7 ($C_{Ar}$), 139.0 ($C_{Ar}$), 129.9 ($C_{Ar}$), 129.6 ($C_{Ar}$), 127.9 ($C_{Ar}$H), 125.8 ($C_{Ar}$), 125.4 ($C_{Ar}$H), 125.2 ($C_{Ar}$), 123.7 ($C_{Ar}$), 122.8 ($C_{Ar}$H), 122.8 ($C_{Ar}$H), 122.2 ($C_{Ar}$), 122.1 ($C_{Ar}$), 111.4 ($C_{Ar}$H), 106.7 ($C_{Ar}$H), 97.7 ($C_{Ar}$H), 63.5 (*C*H), 61.6 ($CH_3$), 61.3 ($CH_3$), 56.4 ($CH_3$), 55.2 ($CH_2$), 55.2 ($CH_2$), 46.0 ($CH_2$), 43.5 (*C*H), 40.5 ($CH_3$), 38.7 ($CH_2$), 37.9 ($CH_2$).

**Me-SS66C-CBI-TMI**

(*S*)-1-(chloromethyl)-3-(5,6,7-trimethoxy-1*H*-indole-2-carbonyl)-2,3-dihydro-1*H*-benzo[*e*]indol-5-yl (*cis*)-4-methylhexahydro-[1,2]dithiino[4,5-*b*]pyrazine-1(2*H*)-carboxylate

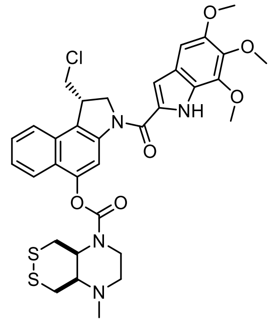

Amide coupling and was accomplished according to the **General Protocol B**. $BF_3 \cdot OEt_2$ (*Method B*) was applied for *N*-Boc deprotection of **Me-SS66C-CBI-Boc** (28.5 mg, 0.052 mmol) and **TMI-OH** (26.1 mg, 0.104 mmol) was activated for final prodrug assembly using oxaxylchloride/DMF (*Method E*). **Me-SS66C-CBI-TMI** was obtained as a colourless solid (15.5 mg, 0.023 mmol, 44%).

**TLC** $R_f$ = 0.36 (DCM:MeOH, 20:1). **HRMS** (ESI$^+$) for $C_{33}H_{36}ClN_4O_6S_2^+$: calcd. *m/z* 683.17593, found *m/z* 683.17527. Individual rotamers/conformers were observed by NMR spectroscopy at 298 K (ratio ca. 1:1). **$^1$H-NMR** (600 MHz, $CDCl_3$): δ (ppm) = 9.37 (s, 2H), 8.38 (s, 2H), 7.90 – 7.81 (m, 2H), 7.78 (d, $J$ = 8.3 Hz, 2H), 7.55 (t, $J$ = 7.5 Hz, 2H), 7.45 (t, $J$ = 7.5 Hz, 2H), 7.01 (d, $J$ = 2.1 Hz, 2H), 6.88 (s, 2H), 4.81 (d, $J$ = 10.6 Hz, 2H), 4.75 – 4.65 (m, 2H), 4.62 (dd, $J$ = 11.0, 3.3 Hz, 1H), 4.40 (d, $J$ = 11.6 Hz, 1H), 4.32 – 4.25 (m, 1H), 4.19 (t, $J$ = 8.4 Hz, 2H), 4.09 (s, 6H), 4.04 (d, $J$ = 11.7 Hz, 2H), 4.03 – 3.98 (m, 2H), 3.95 (s, 6H), 3.92 (s, 6H), 3.61 (t, $J$ = 13.0 Hz, 1H), 3.51 (t, $J$ = 11.0 Hz, 2H), 3.43 – 3.27 (m, 5H), 3.02 (dd, $J$ = 19.7, 11.7 Hz, 2H), 2.72 (dt, $J$ = 13.0, 3.1 Hz, 1H), 2.66 (s, 1H), 2.55 (dd, $J$ = 23.5, 13.2 Hz, 3H), 2.47 (t, $J$ = 11.7 Hz, 1H), 2.33 (s, 6H). **$^{13}$C-NMR** (151 MHz, $CDCl_3$): δ (ppm) = 160.4 (*C*=O), 153.3 (*C*=O, *rotamer 1*), 153.3 (*C*=O, *rotamer 2*), 153.2 ($C_{Ar}$, *rotamer 1*), 153.1 ($C_{Ar}$ , *rotamer 2*), 150.3 ($C_{Ar}$), 148.1 ($C_{Ar}$), 148.0 ($C_{Ar}$, *rotamer 1*), 148.0 ($C_{Ar}$, *rotamer 2*), 141.5 ($C_{Ar}$, *rotamer 1*), 141.5 ($C_{Ar}$, *rotamer 2*), 140.7 ($C_{Ar}$, *rotamer 1*), 139.0 ($C_{Ar}$), 129.9 ($C_{Ar}$, *rotamer 1*), 129.9 ($C_{Ar}$, *rotamer 2*), 129.6 ($C_{Ar}$, *rotamer 1*), 129.6 ($C_{Ar}$, *rotamer 2*), 128.0 ($C_{Ar}$H, *rotamer 1*), 127.9 ($C_{Ar}$H, *rotamer 2*), 125.7 ($C_{Ar}$), 125.4 ($C_{Ar}$H, *rotamer 1*), 125.3 ($C_{Ar}$H, *rotamer 2*), 125.2 ($C_{Ar}$, *rotamer 1*), 125.1 ($C_{Ar}$, *rotamer 1*), 123.7 ($C_{Ar}$), 122.9 ($C_{Ar}$H, *rotamer 1*), 122.8 ($C_{Ar}$H, *rotamer 2*), 122.6 ($C_{Ar}$H, *rotamer 1*), 122.6 ($C_{Ar}$H, *rotamer 2*), 122.2 ($C_{Ar}$, *rotamer 1*), 122.2 ($C_{Ar}$, *rotamer 2*), 122.1 ($C_{Ar}$, *rotamer 1*), 122.1 ($C_{Ar}$, *rotamer 2*), 111.4 ($C_{Ar}$H, *rotamer 1)*, 111.3 ($C_{Ar}$H, *rotamer 2*), 106.7 ($C_{Ar}$H, *rotamer 2*), 97.6 ($C_{Ar}$H, *rotamer 2*), 61.6 ($CH_3$), 61.4 (*C*H, *rotamer 1*), 61.3 ($CH_3$), 61.3 (*C*H, *rotamer 2*), 56.4 ($CH_3$), 56.2 ($CH_2$, *rotamer 1*), 56.2 ($CH_2$, *rotamer 2*), 56.0 ($CH_2$), 55.2 ($CH_2$, *rotamer 1*), 55.1 ($CH_2$, *rotamer 2*), 54.8 (*C*H, *rotamer 1*), 54.7 (*C*H, *rotamer 2*), 53.7 (*C*H, *rotamer 3*), 53.6 (*C*H, *rotamer 4*), 46.0 ($CH_2$), 43.5 (CH), 42.0 ($CH_3$, *rotamer 1*), 41.9 ($CH_3$, *rotamer 2*), 40.9 ($CH_2$, *rotamer 1*), 40.9 ($CH_2$, *rotamer 2*), 40.2 ($CH_2$), 39.1 ($CH_2$), 38.8 ($CH_2$).

**P-SS66C-CBI-TMI**

(*S*)-1-(chloromethyl)-3-(5,6,7-trimethoxy-1*H*-indole-2-carbonyl)-2,3-dihydro-1*H*-benzo[*e*]indol-5-yl (*cis*)-4-(4-(4-methylpiperazin-1-yl)-4-oxobutanoyl)hexahydro-[1,2]dithiino[4,5-*b*]pyrazine-1(2*H*)-carboxylate

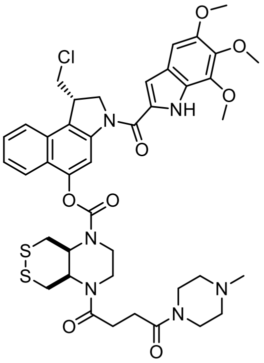

Amide coupling and was accomplished according to the **General Protocol B**. $BF_3{\cdot}OEt_2$ (*Method B*) was applied for *N*-Boc deprotection of **P-SS66C-CBI-Boc** (14.5 mg, 0.020 mmol) and **TMI-OH** (10.0 mg, 0.040 mmol) was activated for final prodrug assembly using oxaxylchloride/DMF (*Method E*). **P-SS66C-CBI-TMI** was obtained as a colourless solid (10.9 mg, 0.013 mmol, 65%).

**TLC** $R_f$ = 0.36 (DCM:MeOH, 20:1). **HRMS** (ESI$^+$) for $C_{41}H_{48}ClN_6O_8S_2^+$: calcd. *m/z* 851.26581, found *m/z* 851.26421. $^1$**H-NMR** (400 MHz, $CDCl_3$): δ (ppm) = 9.37 (s, 1H), 8.40 (d, *J* = 3.8 Hz, 1H), 7.89 – 7.82 (m, 1H), 7.78 (d, *J* = 8.3 Hz, 1H), 7.57 (t, *J* = 7.6 Hz, 1H), 7.51 – 7.43 (m, 1H), 7.01 (d, *J* = 2.2 Hz, 1H), 6.88 (s, 1H), 4.81 (d, *J* = 10.7 Hz, 1H), 4.68 (t, *J* = 9.6 Hz, 1H), 4.47 (s, 1H), 4.19 (t, *J* = 9.4 Hz, 1H), 4.09 (s, 3H), 4.07 – 3.96 (m, 3H), 3.95 (s, 3H), 3.92 (s, 3H), 3.90 – 3.82 (m, 2H), 3.64 (d, *J* = 4.7 Hz, 2H), 3.60 – 3.46 (m, 4H), 3.21 (d, *J* = 14.1 Hz, 1H), 3.04 (d, *J* = 11.2 Hz, 1H), 2.74 (d, *J* = 8.9 Hz, 4H), 2.45 (dd, *J* = 144.7, 4.7 Hz, 2H), 2.39 (t, *J* = 4.9 Hz, 2H), 2.32 (s, 3H). $^{13}$**C-NMR** (101 MHz, $CDCl_3$): δ (ppm) = 172.2 (*C*=O), 170.2 (*C*=O), 160.5 (*C*=O), 154.0 ($C_{Ar}$), 150.4 ($C_{Ar}$), 147.9 ($C_{Ar}$), 141.5 ($C_{Ar}$), 140.8 ($C_{Ar}$), 139.0 ($C_{Ar}$), 130.0 ($C_{Ar}$), 129.6 ($C_{Ar}$), 128.0 ($C_{Ar}$H), 125.8 ($C_{Ar}$), 125.6 ($C_{Ar}$H), 125.1 ($C_{Ar}$), 123.7 ($C_{Ar}$), 122.9 ($C_{Ar}$H), 122.7 ($C_{Ar}$H), 122.3 ($C_{Ar}$), 111.4 ($C_{Ar}$H), 106.8 ($C_{Ar}$H), 97.7 ($C_{Ar}$H), 61.6 ($CH_3$), 61.3 ($CH_3$), 56.4 ($CH_3$), 55.2 ($CH_2$), 55.1 ($CH_2$), 54.8 ($CH_2$), 52.0 (*C*H), 46.1 ($CH_3$), 46.0 ($CH_2$), 45.3 ($CH_2$), 43.5 (*C*H), 42.3 ($CH_2$), 41.8 ($CH_2$), 28.3 ($CH_2$).

**P-CC60-CBI-TMI**

(*S*)-1-(chloromethyl)-3-(5,6,7-trimethoxy-1*H*-indole-2-carbonyl)-2,3-dihydro-1*H*-benzo[*e*]indol-5-yl cyclohexyl(3-(4-methylpiperazin-1-yl)-3-oxopropyl)carbamate

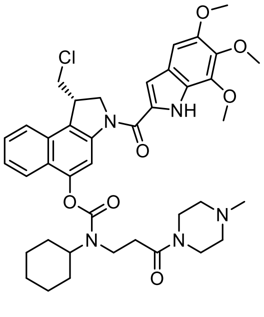

Amide coupling and was accomplished according to the **General Protocol B**. TFA (*Method C*) was applied for *N*-Boc deprotection of **P-CC60-CBI-Boc** (31.0 mg, 0.050 mmol) and **TMI-OH** (76.3 mg, 0.303 mmol) was activated for final prodrug assembly using oxaxylchloride/DMF (*Method E*). **P-CC60-CBI-TMI** was obtained as a colourless solid (29.0 mg, 0.039 mmol, 78%).

**TLC** $R_f$ = 0.58 (DCM:MeOH, 9:1). **HRMS** (ESI$^+$) for $C_{40}H_{49}ClN_5O_7^+$: calcd. *m/z* 746.33150, found *m/z* 746.33220. $^1$**H-NMR** (800 MHz, MeOD-$d_4$): δ (ppm) = 8.29 – 8.09 (m, 1H), 7.93 – 7.82 (m, 2H), 7.56 (t, *J* = 7.5 Hz, 1H), 7.52 – 7.43 (m, 1H), 7.06 (s, 1H), 6.97 (s, 1H), 4.66 (q, *J* = 11.5, 10.9 Hz, 2H), 4.30 – 4.13 (m, 2H), 4.05 (s, 3H), 4.00 (dd, *J* = 11.3, 3.2 Hz, 1H), 3.89 (s, 6H), 3.83 (dd, *J* = 13.8, 7.5 Hz, 1H), 3.79 – 3.71 (m, 1H), 3.67 – 3.56 (m, 4H), 2.92 (s, 1H), 2.83 – 2.73 (m, 1H), 2.45 (d, *J* = 18.6 Hz, 2H), 2.36 (d, *J* = 21.5 Hz, 2H), 2.30 – 2.18 (m, 3H), 2.04 (d, *J* = 10.2 Hz, 1H), 2.00 – 1.91 (m, 1H), 1.88 (d, *J* = 9.2 Hz, 2H), 1.72 (dd, *J* = 31.7, 12.9 Hz, 3H), 1.54 – 1.18 (m, 4H). $^{13}$**C-NMR** (201 MHz, MeOD-$d_4$): δ (ppm) = 171.8 (*C*=O), 171.5 (*C*=O), 162.6 (*C*=O), 151.2 ($C_{Ar}$), 149.1 ($C_{Ar}$), 142.7 ($C_{Ar}$), 141.7 ($C_{Ar}$), 140.4 ($C_{Ar}$), 131.4 ($C_{Ar}$), 128.9 ($C_{Ar}$H), 127.3 ($C_{Ar}$), 126.4 ($C_{Ar}$, $C_{Ar}$H), 125.2 ($C_{Ar}$), 124.3 ($C_{Ar}$H), 123.6 ($C_{Ar}$H), 112.1 ($C_{Ar}$H), 108.2 ($C_{Ar}$H), 99.3 ($C_{Ar}$H), 61.8 ($CH_3$), 61.7 ($CH_3$), 58.9 (*C*H), 58.8 (*C*H), 56.7 ($CH_2$), 55.8 ($CH_3$), 55.3 ($CH_2$), 47.5 ($CH_2$), 46.3 ($CH_2$), 45.8 ($CH_3$), 43.5 (*C*H), 42.2 ($CH_2$), 41.9 ($CH_2$), 35.4 ($CH_2$), 34.1 ($CH_2$), 32.6 ($CH_2$), 31.7 ($CH_2$), 30.4 ($CH_2$), 27.1 ($CH_2$), 26.4 ($CH_2$).

**SS00$_M$-CBI-TMI**

(*S*)-1-(chloromethyl)-3-(5,6,7-trimethoxy-1*H*-indole-2-carbonyl)-2,3-dihydro-1*H*-benzo[*e*]indol-5-yl methyl(2-((2-morpholinoethyl)disulfaneyl)ethyl)carbamate

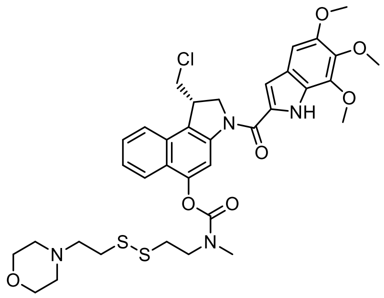

Amide coupling and was accomplished according to the **General Protocol B**. $BF_3 \cdot OEt_2$ (*Method B*) was applied for *N*-Boc deprotection of **SS00$_M$-CBI-Boc** (18.7 mg, 0.031 mmol) and **TMI-OH** (15.8 mg, 0.063 mmol) was activated for final prodrug assembly using EDCI (*Method D*). **SS00$_M$-CBI-TMI** was obtained as a colourless solid (8.5 mg, 0.012 mmol, 37%).

**TLC** $R_f$ = 0.65 (EtOAc). **HRMS** (ESI$^+$) for $C_{35}H_{41}ClN_4O_7S_2^+$: calcd. *m/z* 728.20942, found *m/z* 729.21892. **$^1$H-NMR** (500 MHz, DMSO-$d_6$): δ (ppm) = 11.47 (d, *J* = 1.6 Hz, 1H, N-*H*), 8.16 (s, 1H), 8.02 (d, *J* = 8.4 Hz, 1H), 7.91 (dd, *J* = 17.8, 8.5 Hz, 1H), 7.65 – 7.57 (m, 1H), 7.50 (t, *J* = 7.6 Hz, 1H), 7.10 (d, *J* = 2.1 Hz, 1H), 6.97 (s, 1H), 4.82 (t, *J* = 10.1 Hz, 1H), 4.54 (dd, *J* = 11.0, 2.1 Hz, 1H), 4.37 (s, 1H), 4.08 (dd, *J* = 11.2, 3.2 Hz, 1H), 3.99 – 3.95 (m, 1H), 3.94 (s, 3H), 3.91 – 3.86 (m, 1H), 3.82 (s, 3H), 3.80 (s, 3H), 3.67 – 3.61 (m, 1H), 3.54 – 3.49 (m, 2H), 3.46 – 3.42 (m, 2H), 3.25 (s, 2H), 3.14 (t, *J* = 6.9 Hz, 1H), 3.03 (t, *J* = 6.8 Hz, 1H), 3.00 (s, 1H), 2.89 (q, *J* = 6.6 Hz, 2H), 2.57 (t, *J* = 7.1 Hz, 1H), 2.54 (d, *J* = 7.6 Hz, 1H), 2.36 (d, *J* = 1.8 Hz, 2H), 2.27 (s, 2H). **$^{13}$C-NMR** (126 MHz, DMSO-$d_6$): δ (ppm) = 160.3 (*C*=O), 153.9 ($C_{Ar}$), 153.7 ($C_{Ar}$), 149.2 ($C_{Ar}$), 147.5 ($C_{Ar}$), 147.3 ($C_{Ar}$), 141.5 ($C_{Ar}$), 140.0 ($C_{Ar}$), 139.1 ($C_{Ar}$), 130.6 ($C_{Ar}$), 129.6 ($C_{Ar}$H), 127.6 ($C_{Ar}$), 125.5 ($C_{Ar}$), 125.0 ($C_{Ar}$), 124.4 ($C_{Ar}$H), 123.4 ($C_{Ar}$H), 123.2, 122.4 ($C_{Ar}$), 122.2 ($C_{Ar}$H), 110.6 ($C_{Ar}$), 110.4 ($C_{Ar}$), 106.4 ($C_{Ar}$H), 98.0 ($C_{Ar}$H), 66.1 ($CH_2$, *rotamer 1*), 66.0 ($CH_2$, *rotamer 2*), 61.1 ($CH_3$), 61.0 ($CH_3$), 57.4 ($CH_2$), 56.0 ($CH_3$), 55.0 ($CH_2$), 53.0 ($CH_2$), 52.9 ($CH_2$), 48.2 ($CH_2$), 47.9 ($CH_2$), 47.6 ($CH_2$), 41.1 (*C*H), 35.6 ($CH_2$), 35.2 ($CH_2$), 34.9 ($CH_3$, *rotamer 1*), 34.8 ($CH_3$, *rotamer 2*).

**MC-CBI-TMI**

(*S*)-1-(chloromethyl)-3-(5,6,7-trimethoxy-1*H*-indole-2-carbonyl)-2,3-dihydro-1*H*-benzo[*e*]indol-5-yl methyl carbonate

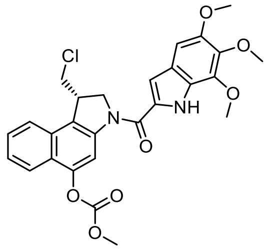

Amide coupling and was accomplished according to the **General Protocol B**. $BF_3 \cdot OEt_2$ (*Method B*) was applied for *N*-Boc deprotection of **MC-CBI-Boc** (39.2 mg, 0.100 mmol) and **TMI-OH** (25.1 mg, 0.010 mmol) was activated for final prodrug assembly using EDCI (*Method D*). **MC-CBI-TMI** was obtained as a colourless solid (7.9 mg, 0.015 mmol, 15%).

**TLC** $R_f$ = 0.55 (isohexane:EtOAc, 1:1). **HRMS** (ESI$^+$) for $C_{27}H_{26}ClN_2O_7^+$: calcd. *m/z* 525.14231, found *m/z* 525.14229. **$^1$H-NMR** (500 MHz, $CDCl_3$): δ (ppm) = 9.39 (s, 1H), 8.49 (s, 1H), 7.99 (d, *J* = 8.4 Hz, 1H), 7.79 (d, *J* = 8.3 Hz, 1H), 7.58 (t, *J* = 7.2 Hz, 1H), 7.47 (t, *J* = 7.3 Hz, 1H), 7.01 (d, *J* = 2.2 Hz, 1H), 6.88 (s, 1H), 4.83 (dd, *J* = 10.8, 1.7 Hz, 1H), 4.74 – 4.65 (m, 1H), 4.20 (t, *J* = 9.4 Hz, 2H), 4.09 (s, 3H), 4.03 – 3.99 (m, 2H), 3.98 (s, 3H), 3.95 (s, 3H), 3.92 (s, 3H), 3.52 (t, *J* = 11.0 Hz, 1H). **$^{13}$C-NMR** (126 MHz, $CDCl_3$): δ (ppm) = 160.5 (*C*=O), 154.3 ($C_{Ar}$), 150.4 ($C_{Ar}$), 147.9 ($C_{Ar}$), 141.5 ($C_{Ar}$), 140.8 ($C_{Ar}$), 139.0 ($C_{Ar}$), 130.0 ($C_{Ar}$), 129.7 ($C_{Ar}$), 128.1 ($C_{Ar}$H), 125.8 ($C_{Ar}$), 125.5 ($C_{Ar}$H), 124.5 ($C_{Ar}$), 123.7 ($C_{Ar}$), 122.8 ($C_{Ar}$H), 122.7 ($C_{Ar}$H), 122.5 ($C_{Ar}$), 110.9 ($C_{Ar}$H), 106.7 ($C_{Ar}$H), 97.8 ($C_{Ar}$H), 61.6 ($CH_3$), 61.3 ($CH_3$), 56.4 ($CH_3$), 55.9 ($CH_3$), 55.2 ($CH_2$), 45.9 ($CH_2$), 43.6 (*C*H).

#### 10. NMR Spectra

#### SS60-CBI-Boc

[1]H-NMR

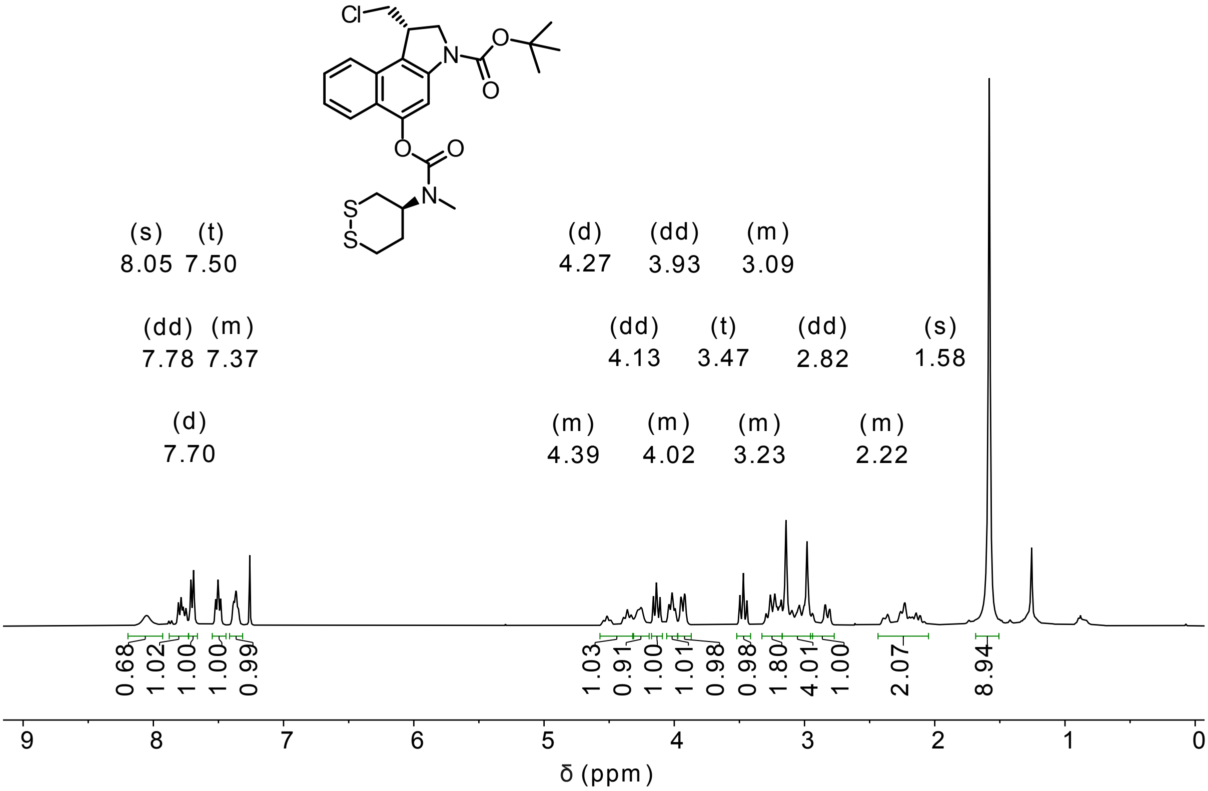


[13]C-NMR

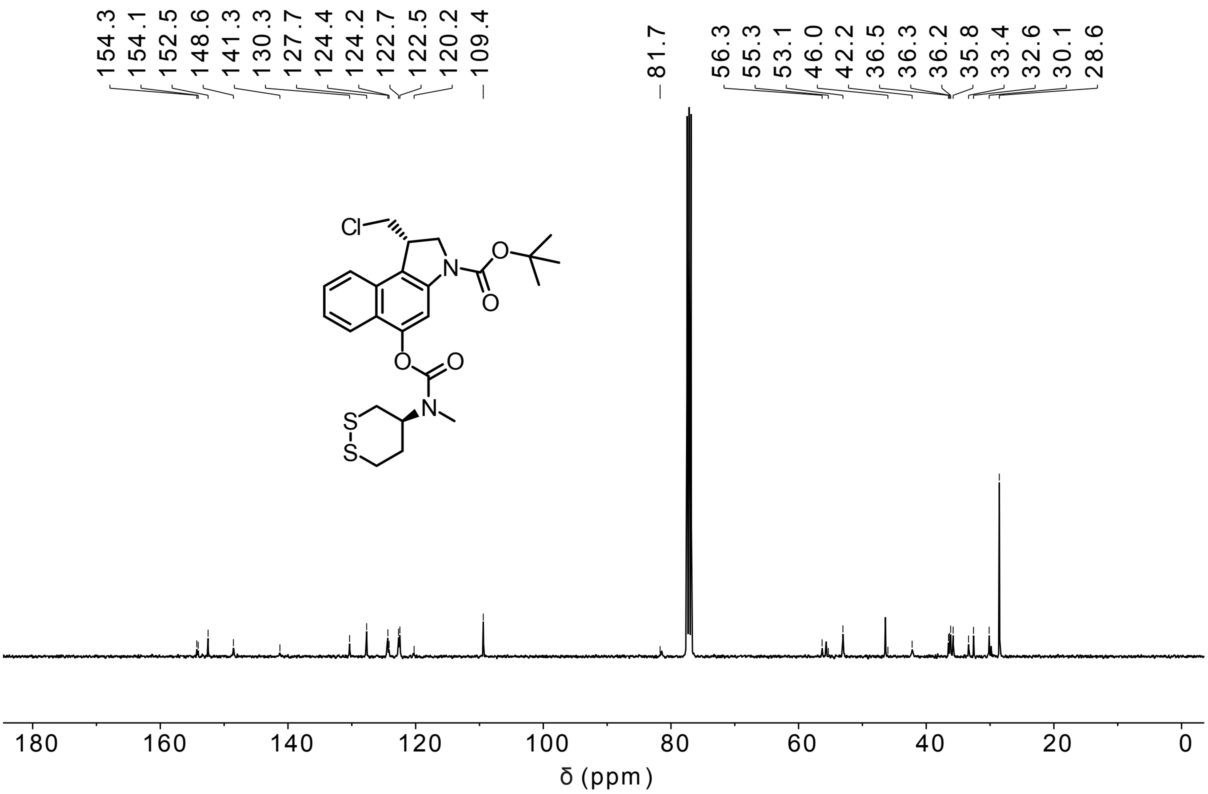

**P-SS60-CBI-Boc**

$^{1}$H-NMR

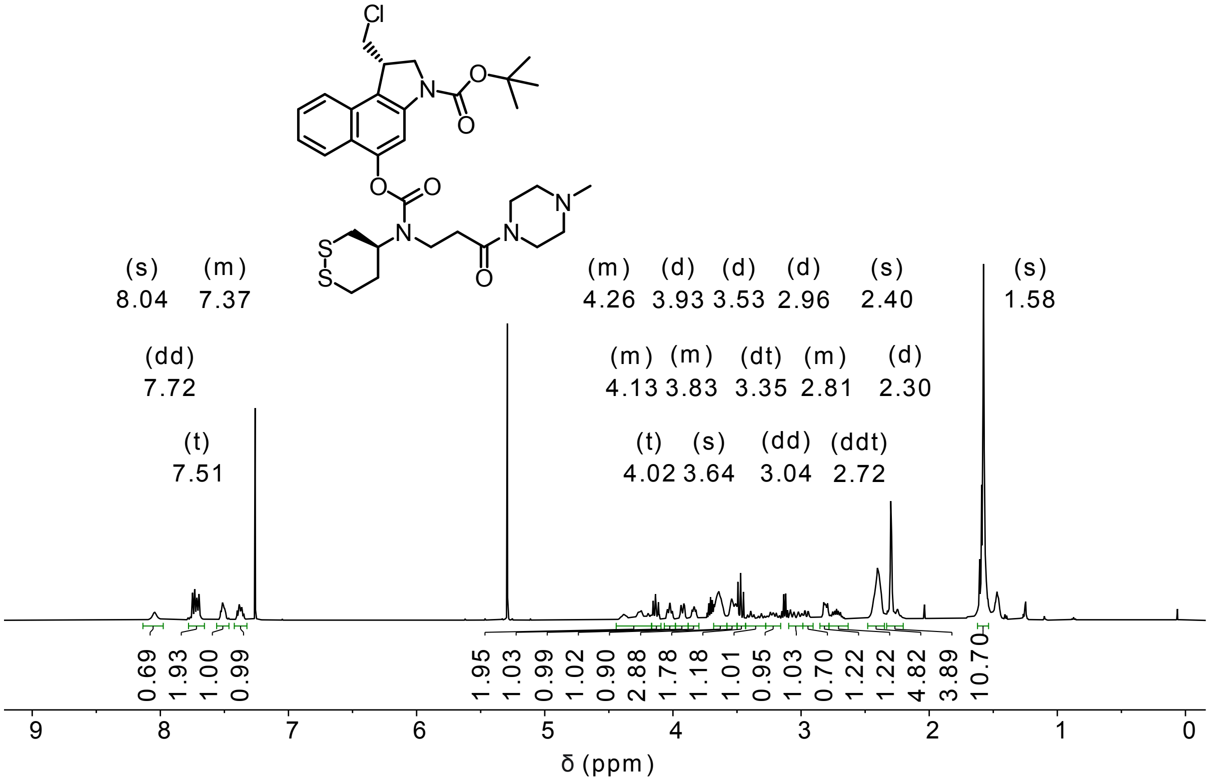


$^{13}$C-NMR

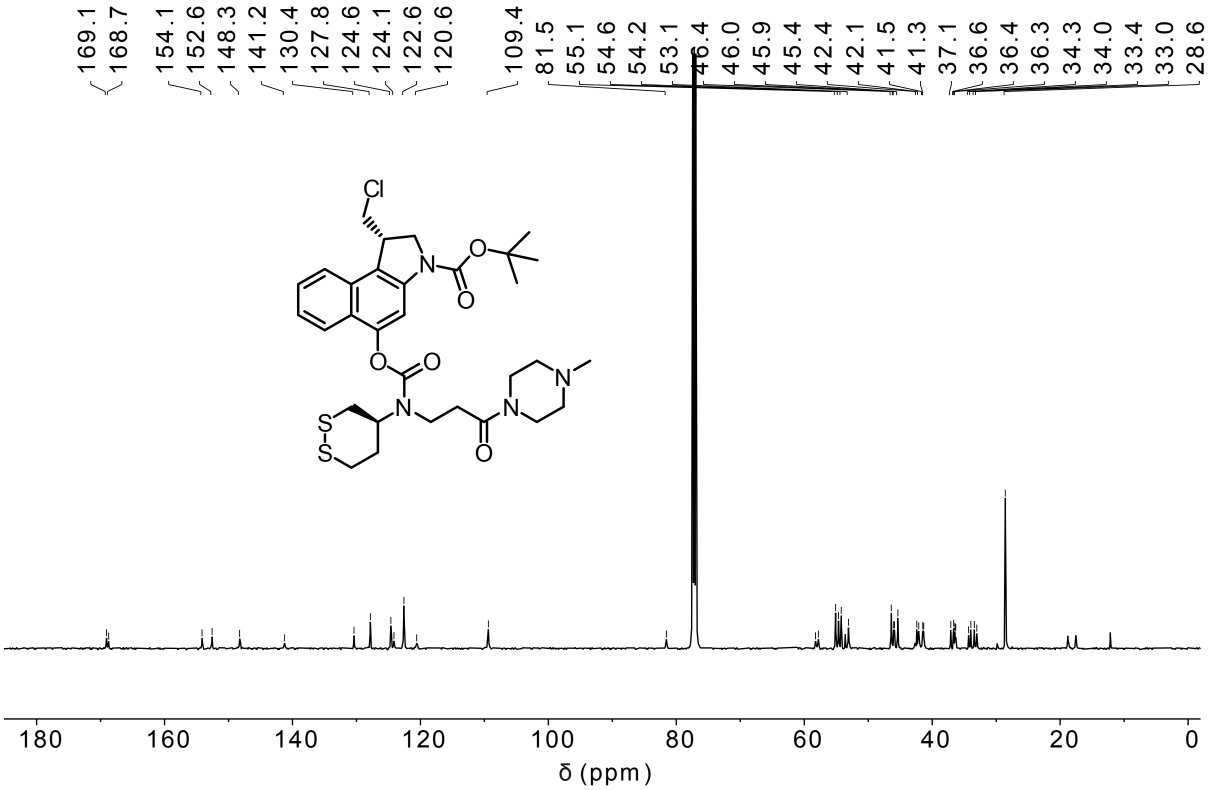

**SeS60-CBI-Boc**

$^{1}$H-NMR

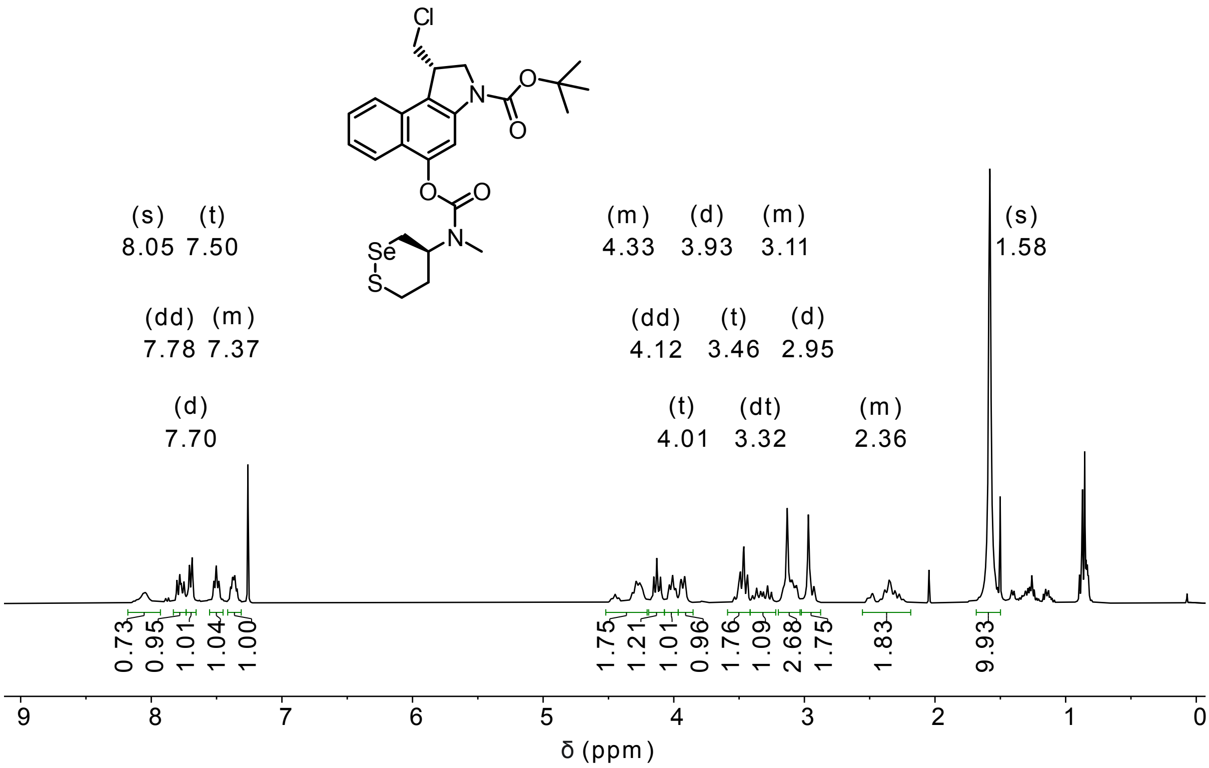


$^{13}$C-NMR

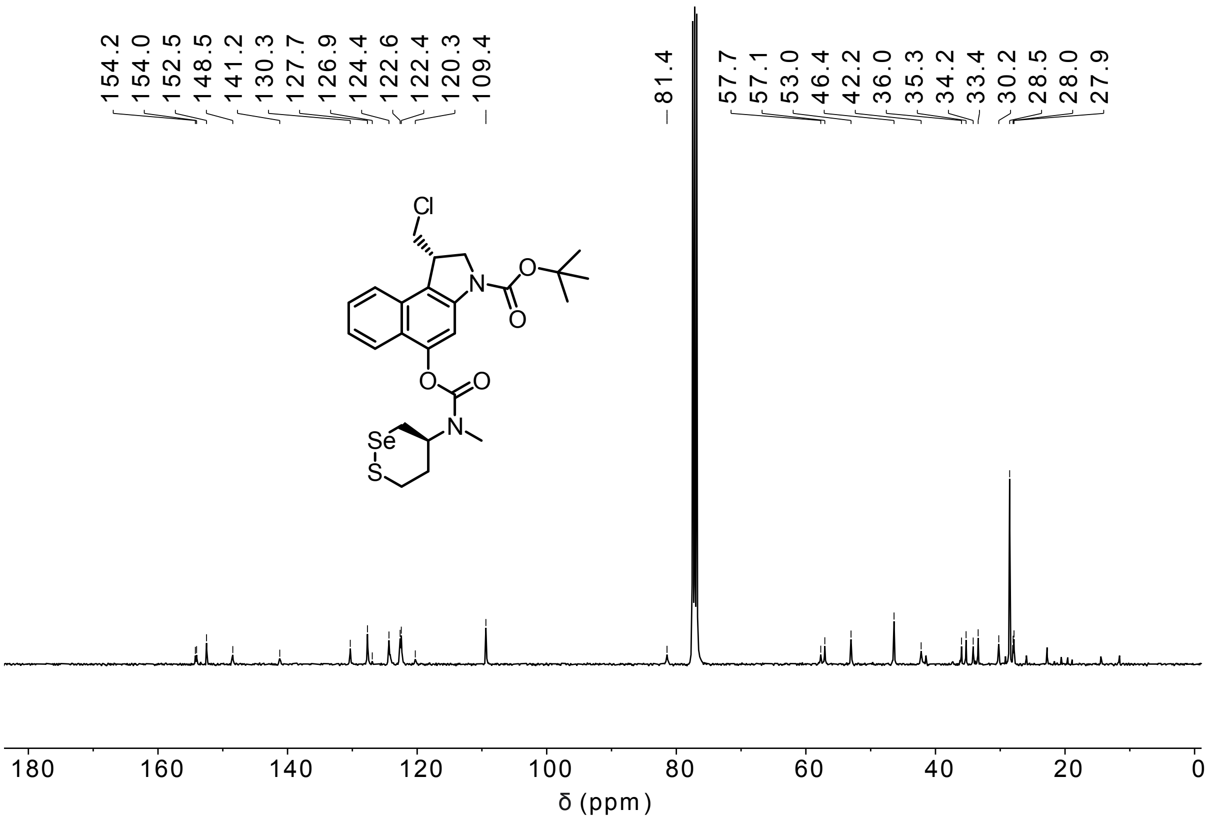

**H-SS66T-CBI-Boc**

[1]H-NMR

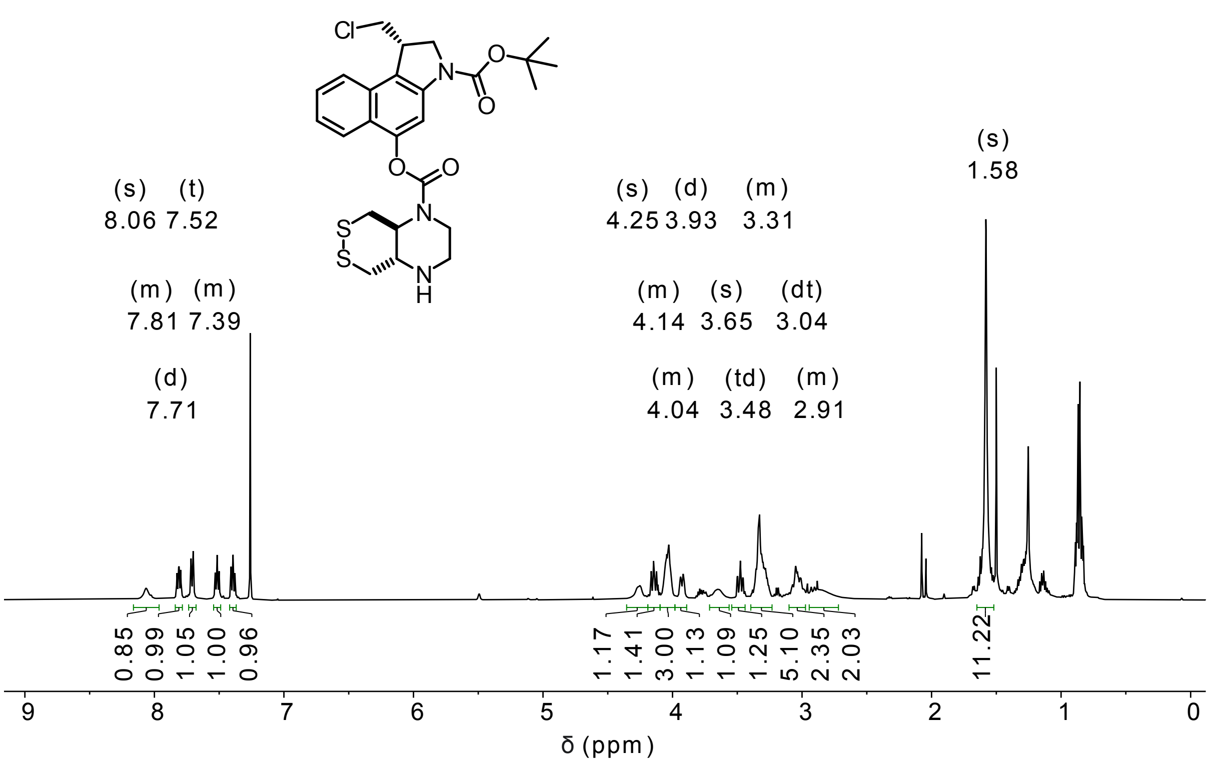


[13]C-NMR

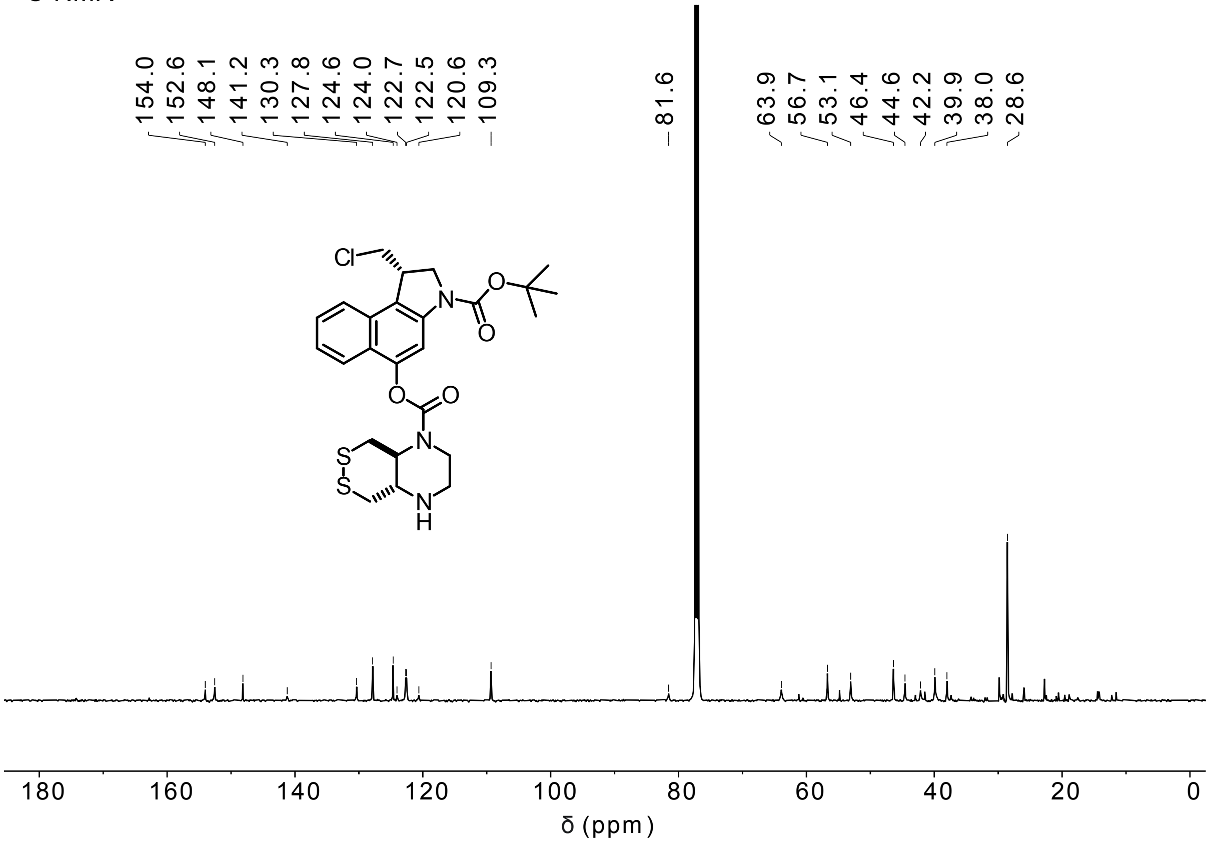

**Me-SS66T-CBI-Boc**

$^{1}$H-NMR

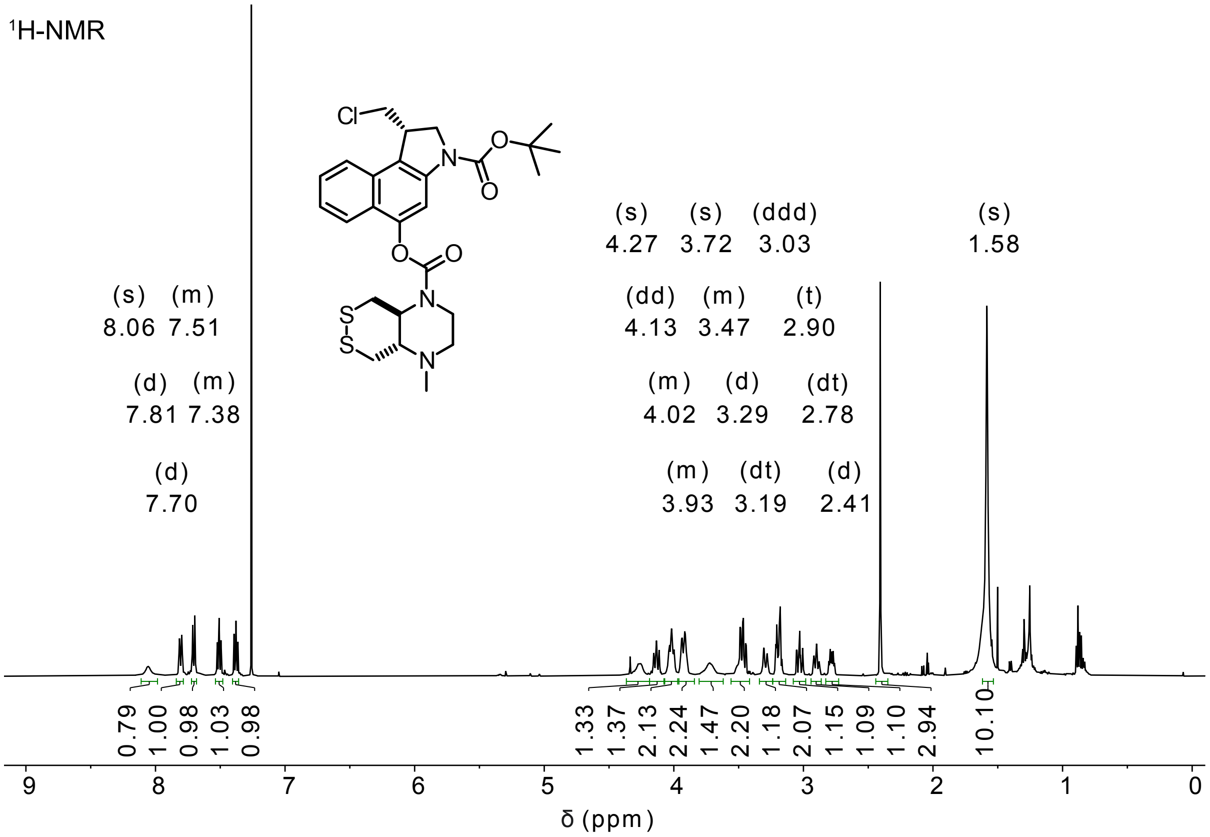


$^{13}$C-NMR

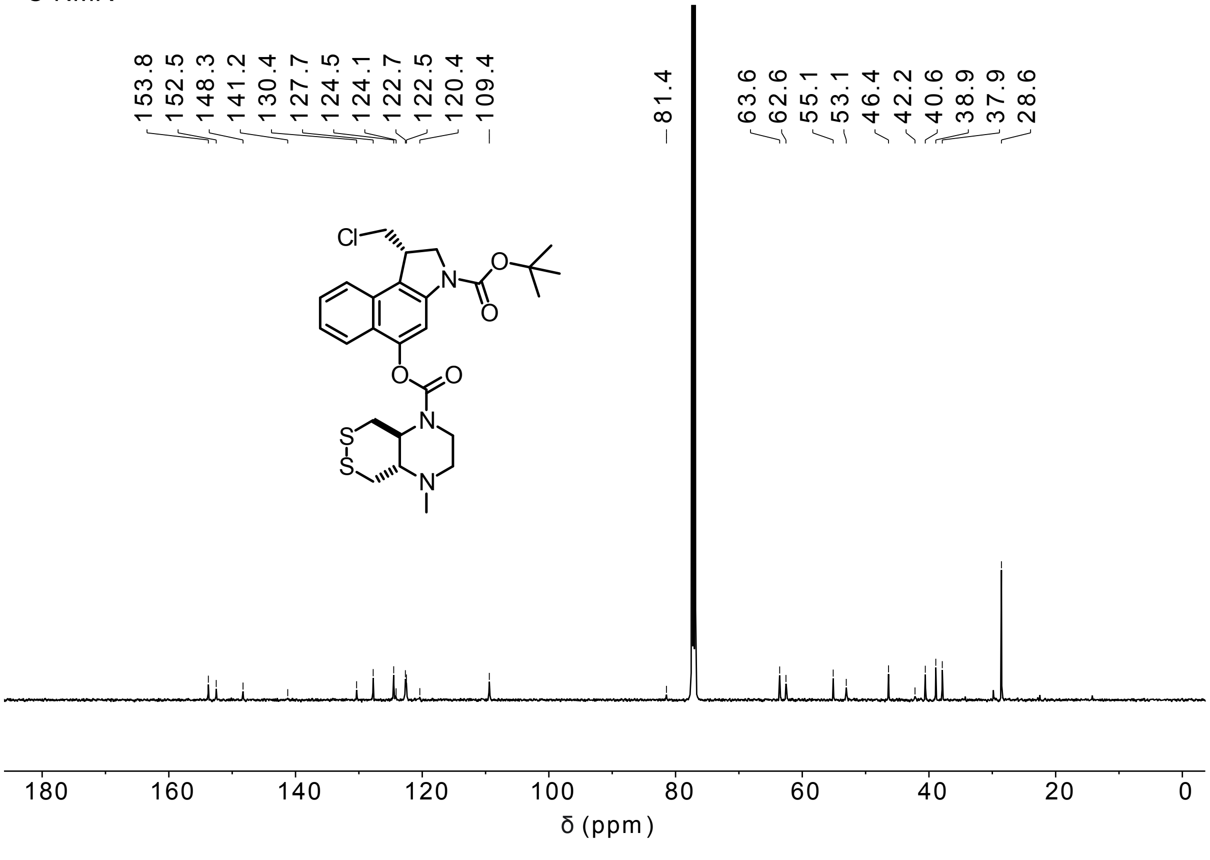

#### H-SS66C-CBI-Boc

$^{1}$H-NMR

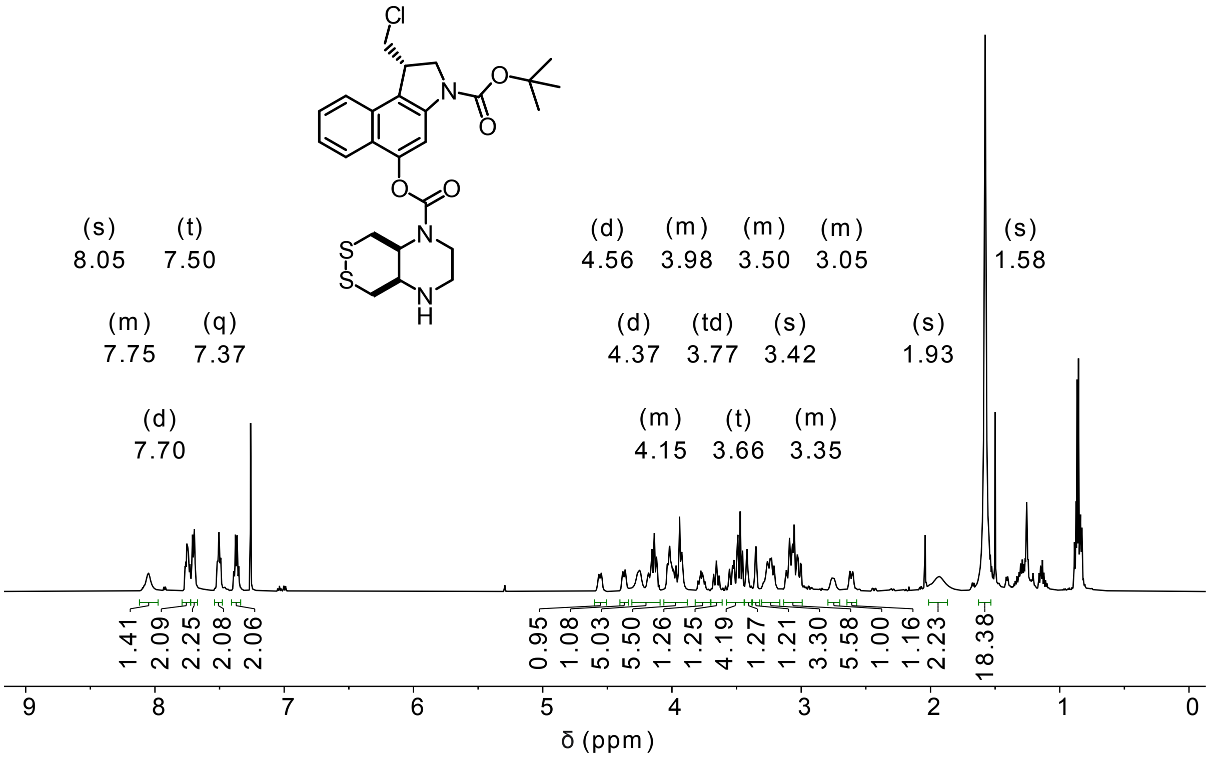


$^{13}$C-NMR

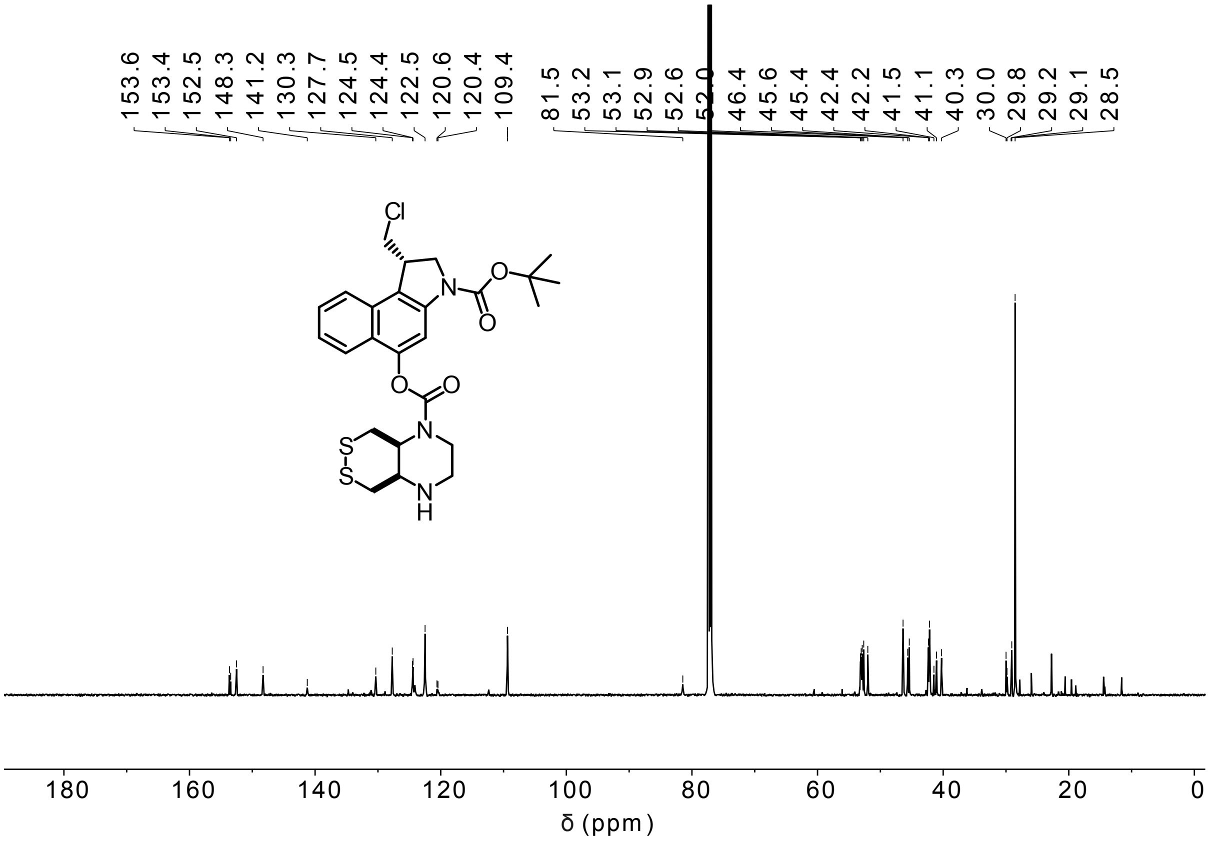

**Me-SS66C-CBI-Boc**

$^{1}$H-NMR

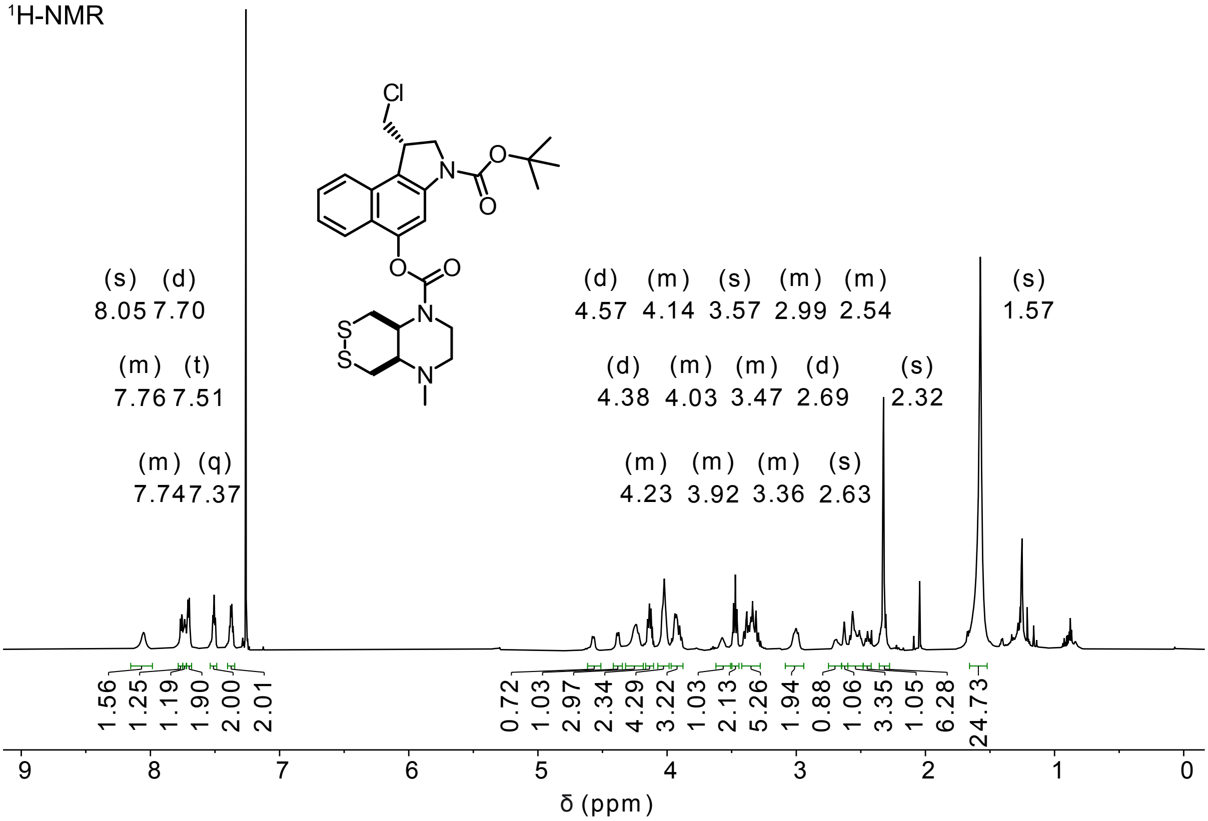


$^{13}$C-NMR

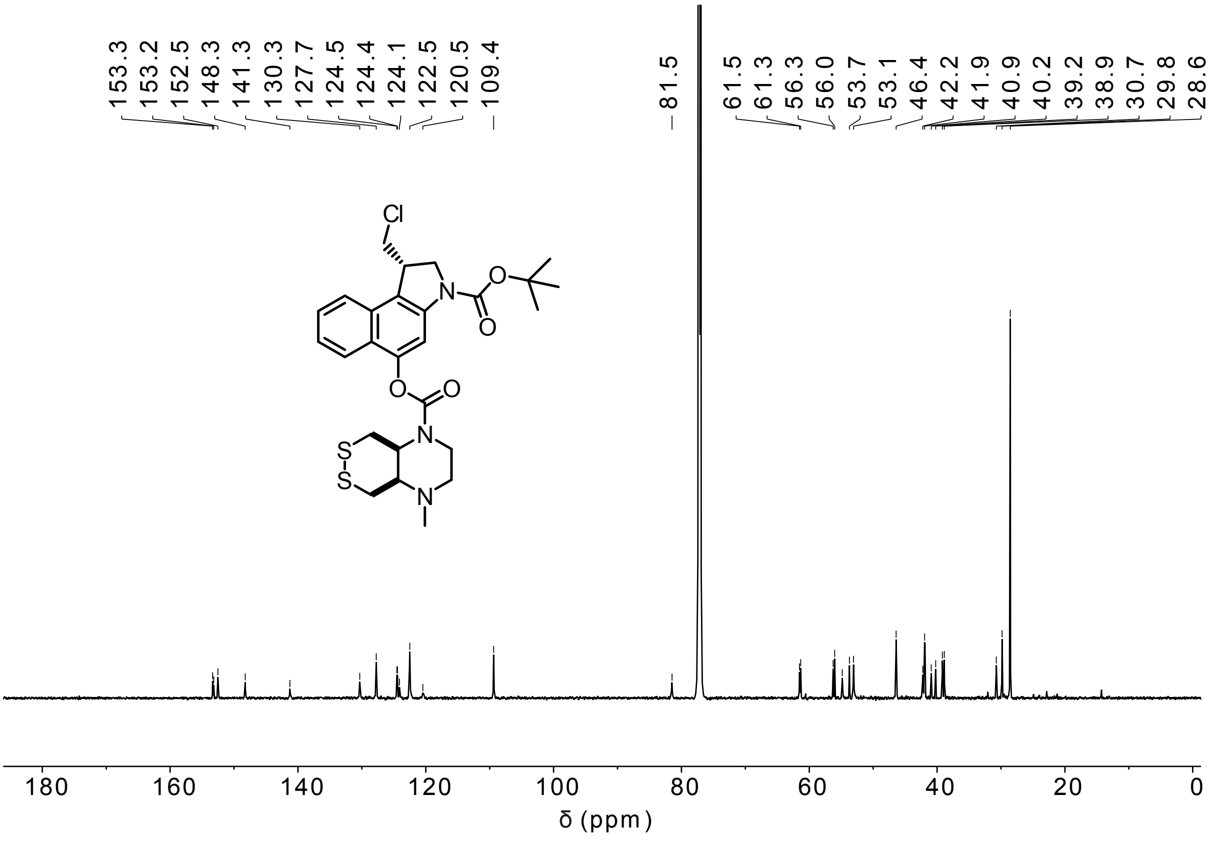

**P-SS66C-CBI-Boc**

$^{1}$H-NMR

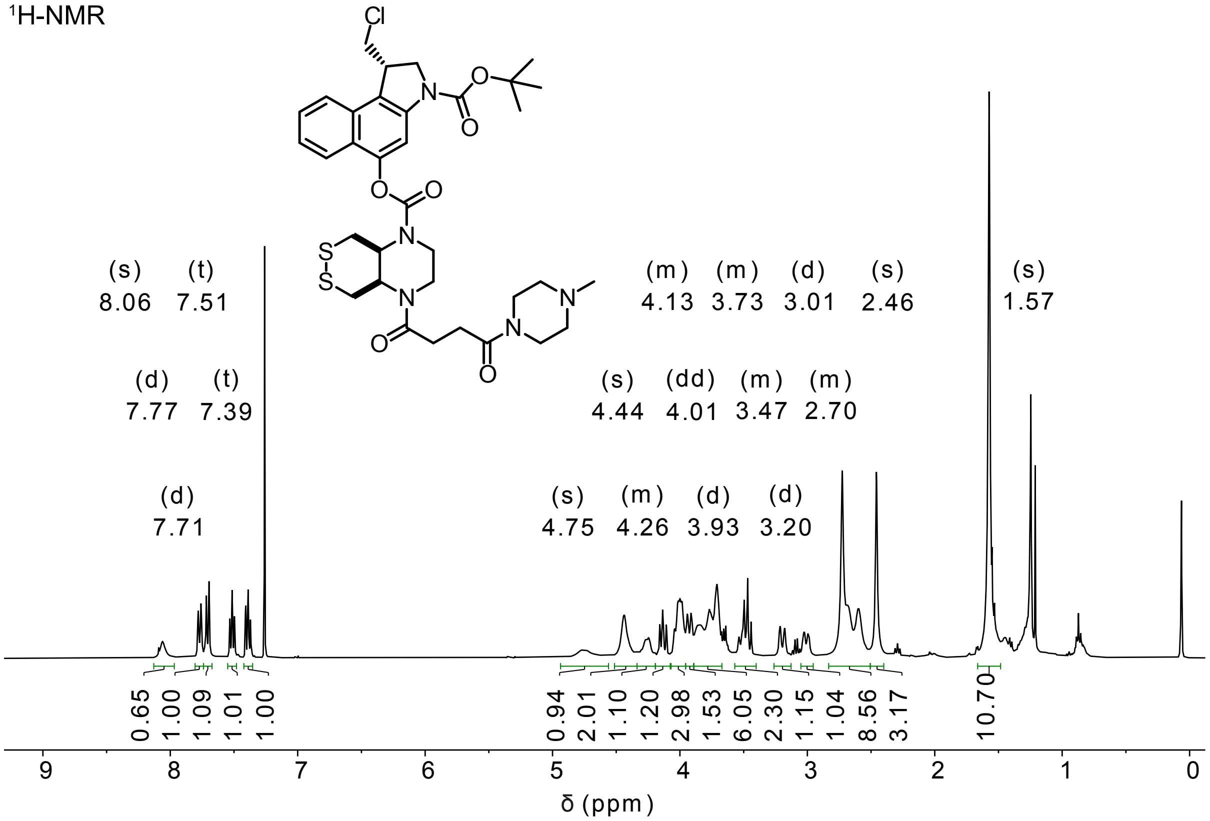


$^{13}$C-NMR

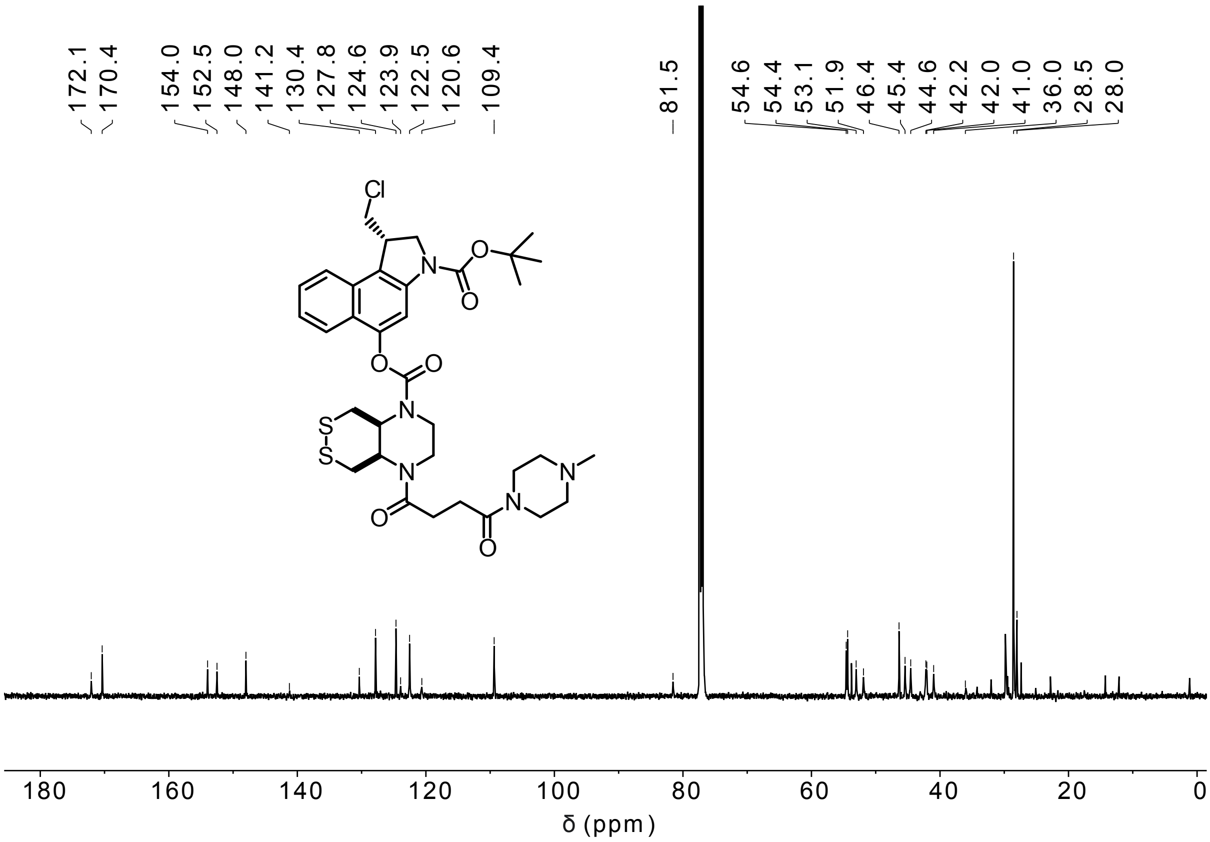

**SS66T-CBI-Boc**

$^{1}$H-NMR

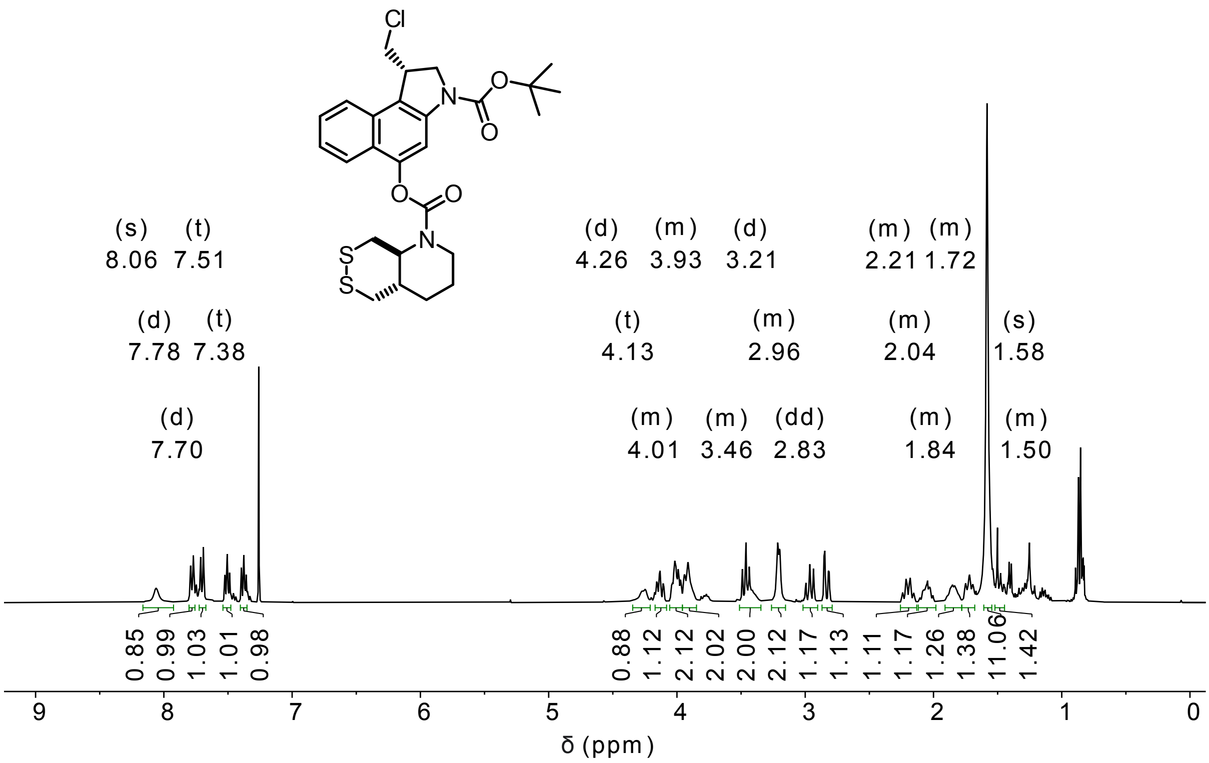


$^{13}$C-NMR

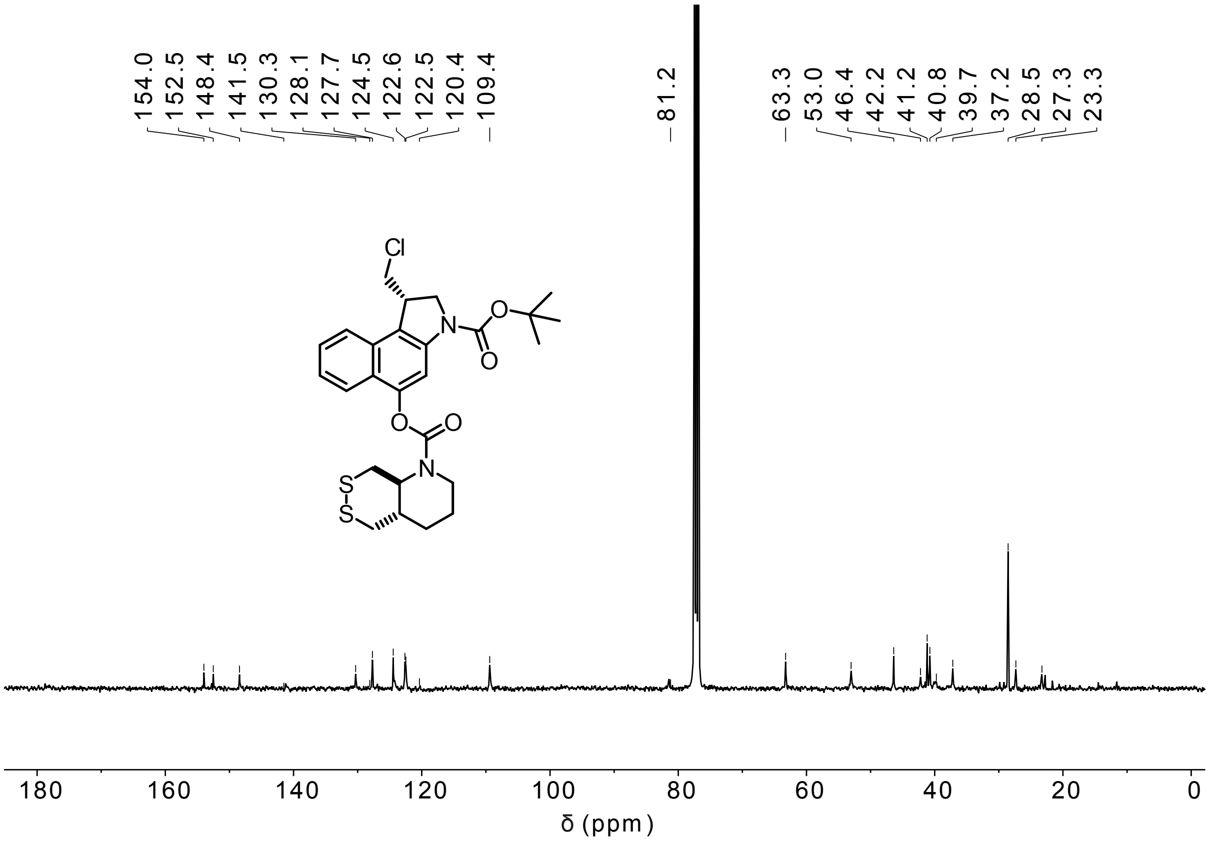

**SS66-CBI-Boc (1:1 mixture cis:trans)**

$^{1}$H-NMR

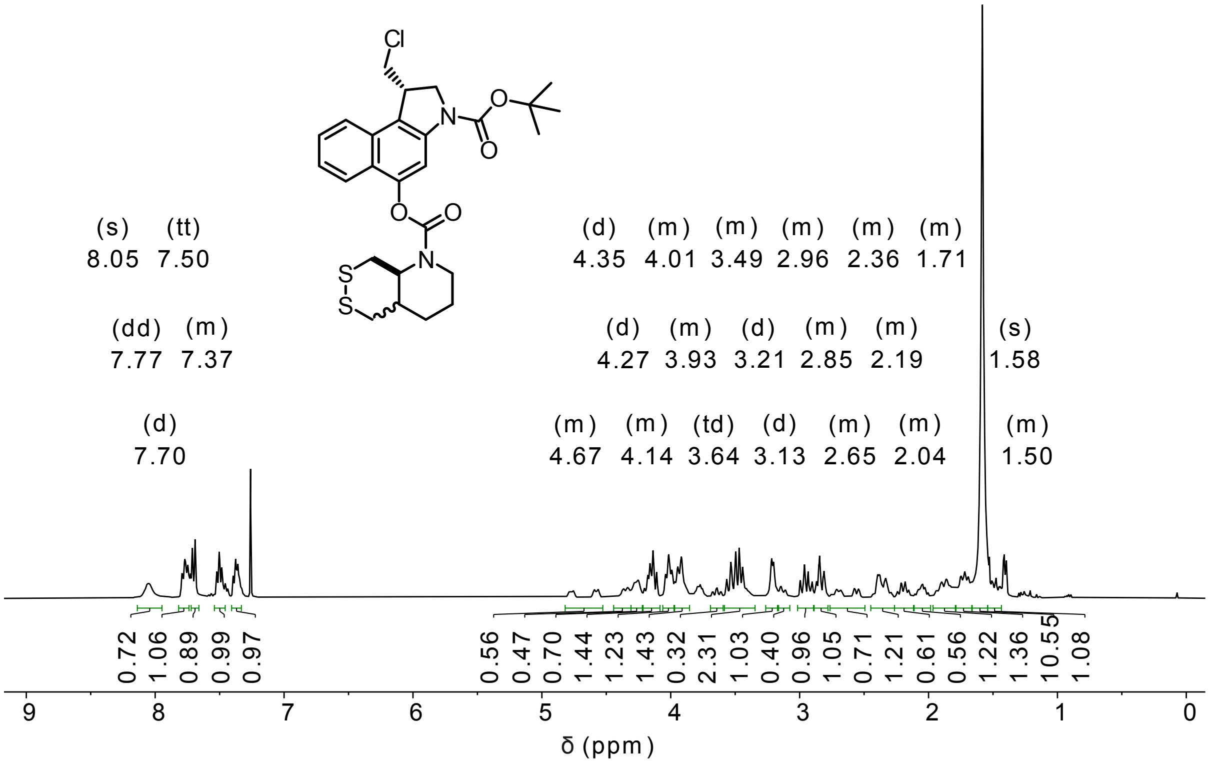


$^{13}$C-NMR

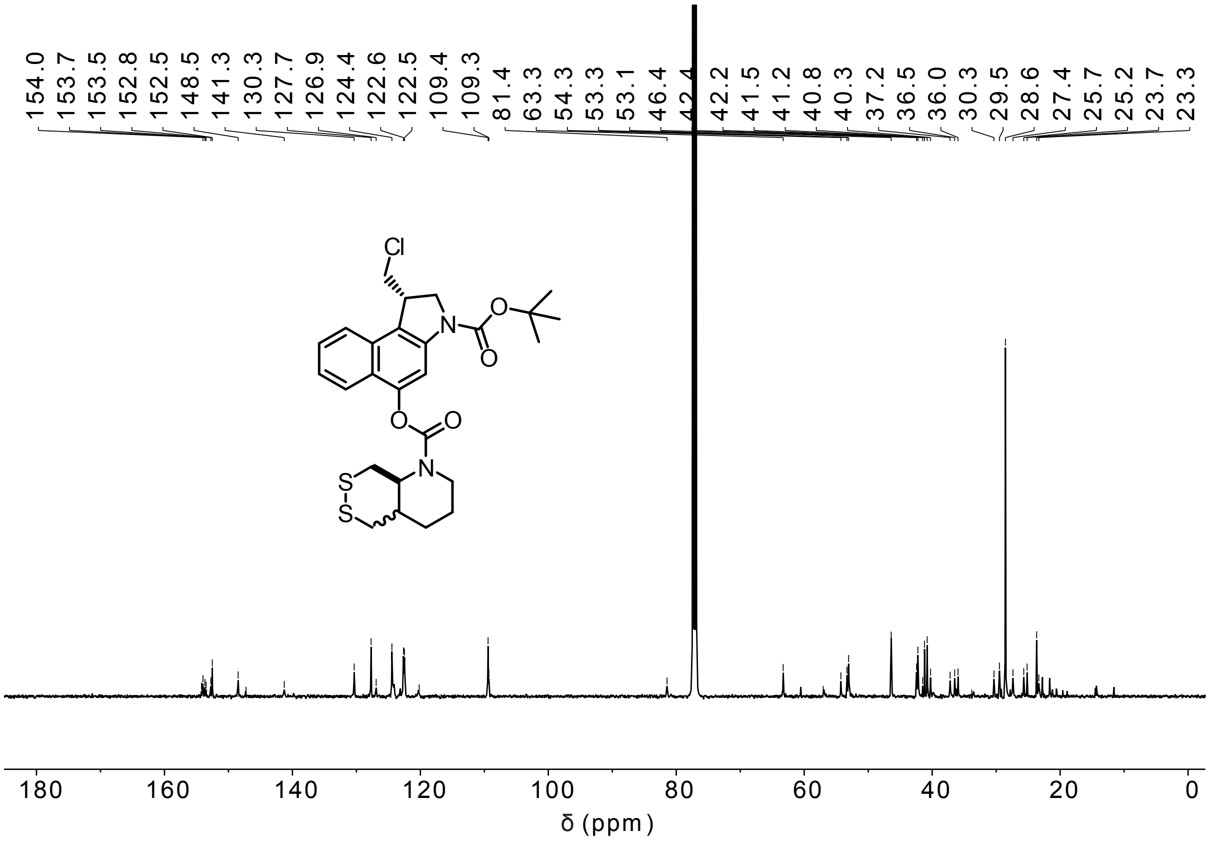

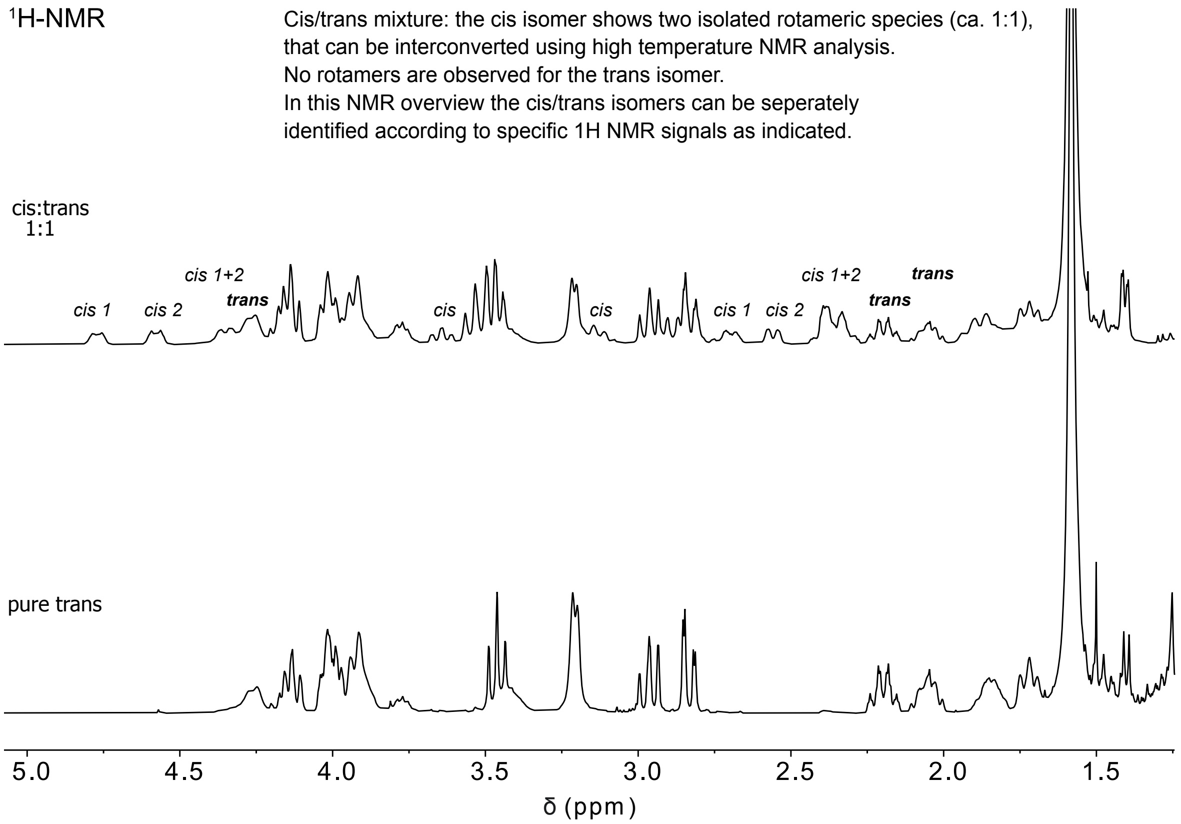
1H-NMR
Cis/trans mixture: the cis isomer shows two isolated rotameric species (ca. 1:1),
that can be interconverted using high temperature NMR analysis.
No rotamers are observed for the trans isomer.
In this NMR overview the cis/trans isomers can be seperately
identified according to specific 1H NMR signals as indicated.
cis:trans
1:1
cis 1
cis 2
cis 1+2
trans
cis
cis
cis 1
cis 2
cis 1+2
trans
trans
pure trans
5.0
4.5
4.0
3.5
3.0
2.5
2.0
1.5
δ (ppm)

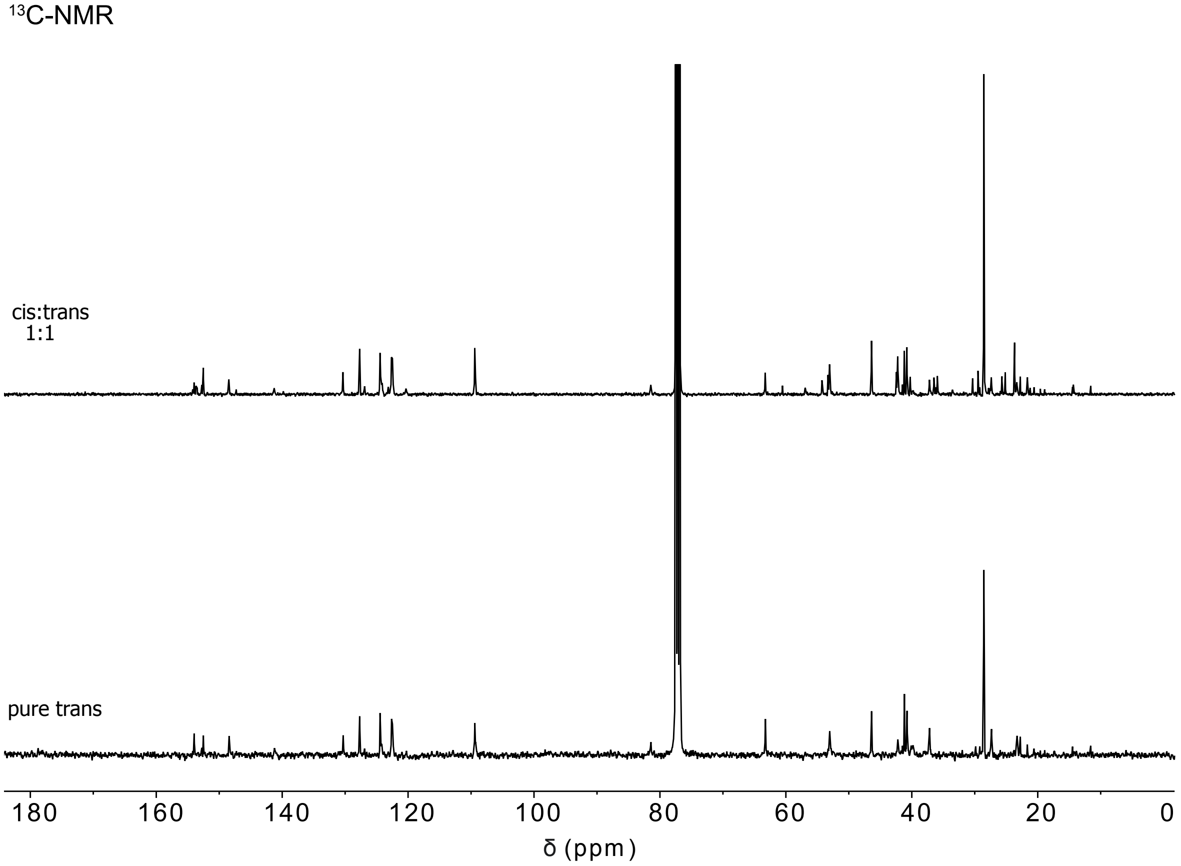
13C-NMR
cis:trans
1:1
pure trans
180
160
140
120
100
80
60
40
20
0
δ (ppm)

#### SS00$_M$-CBI-Boc

$^1$H-NMR

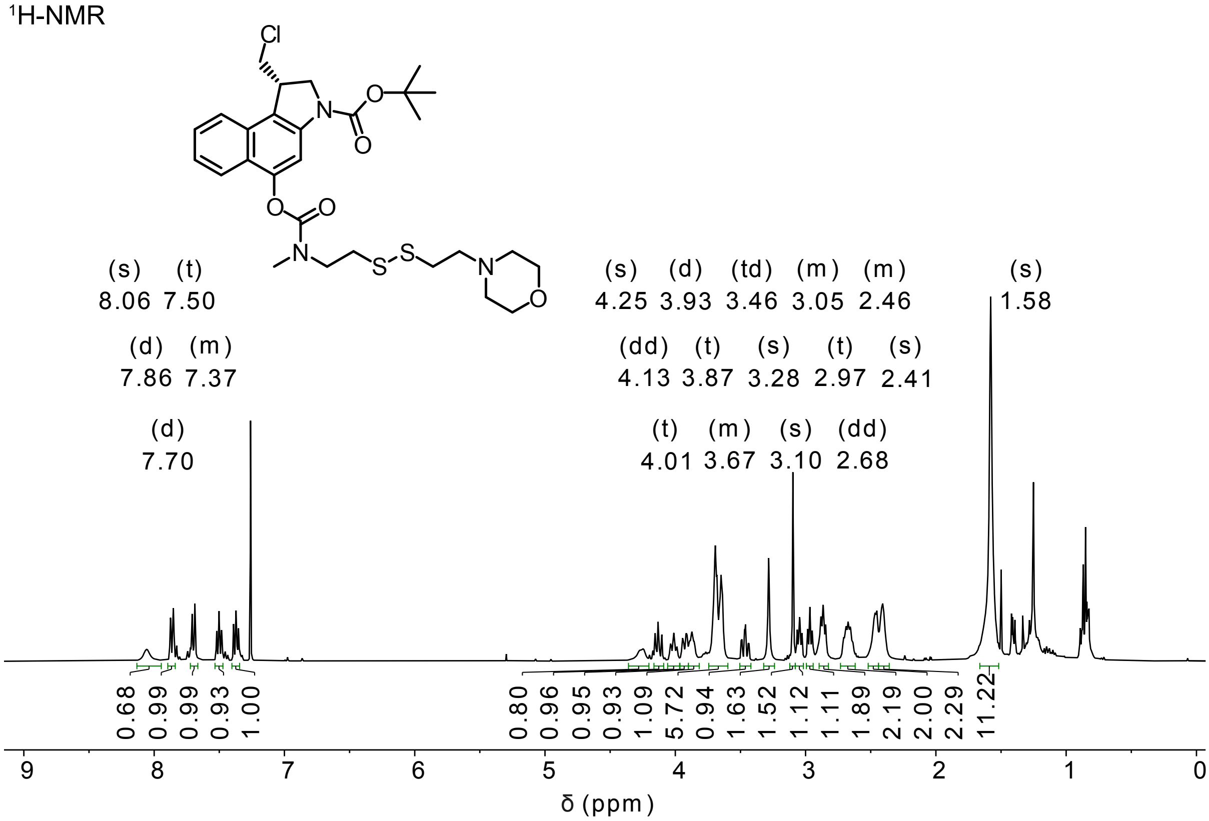


$^{13}$C-NMR

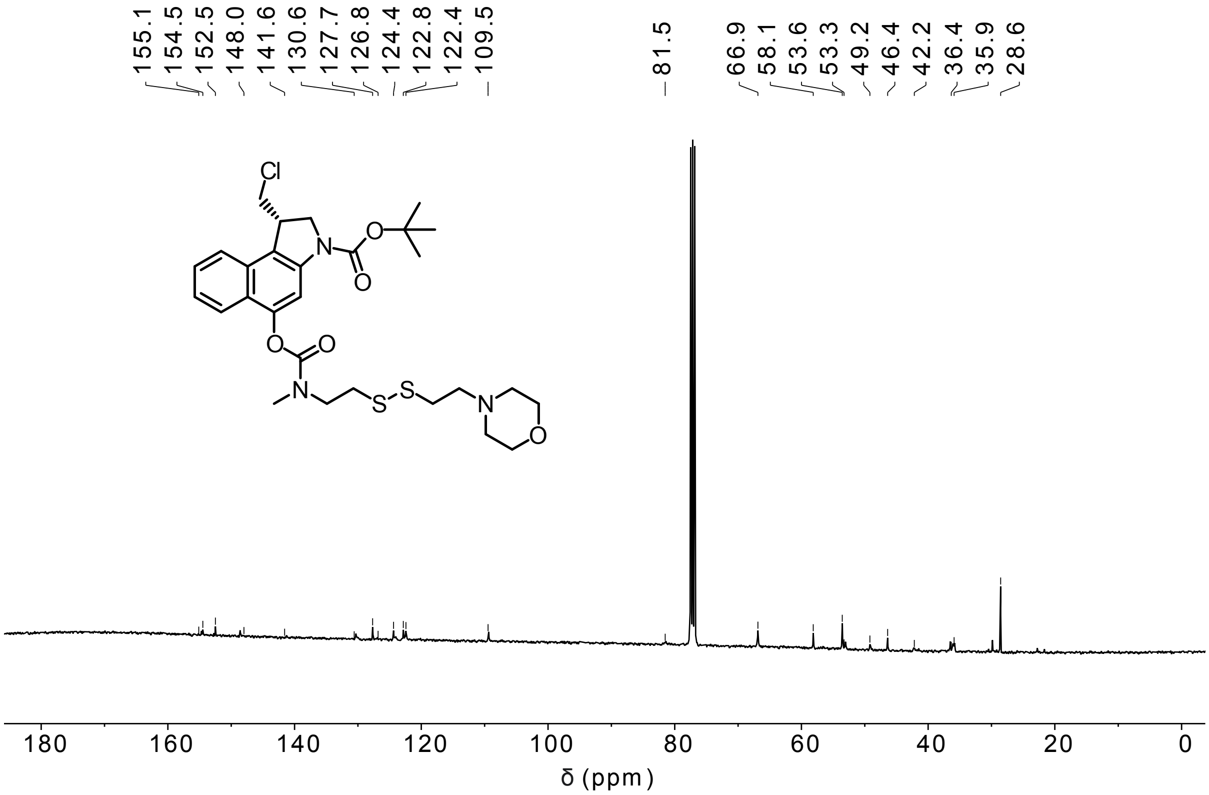

**MC-CBI-Boc**

$^1$H-NMR

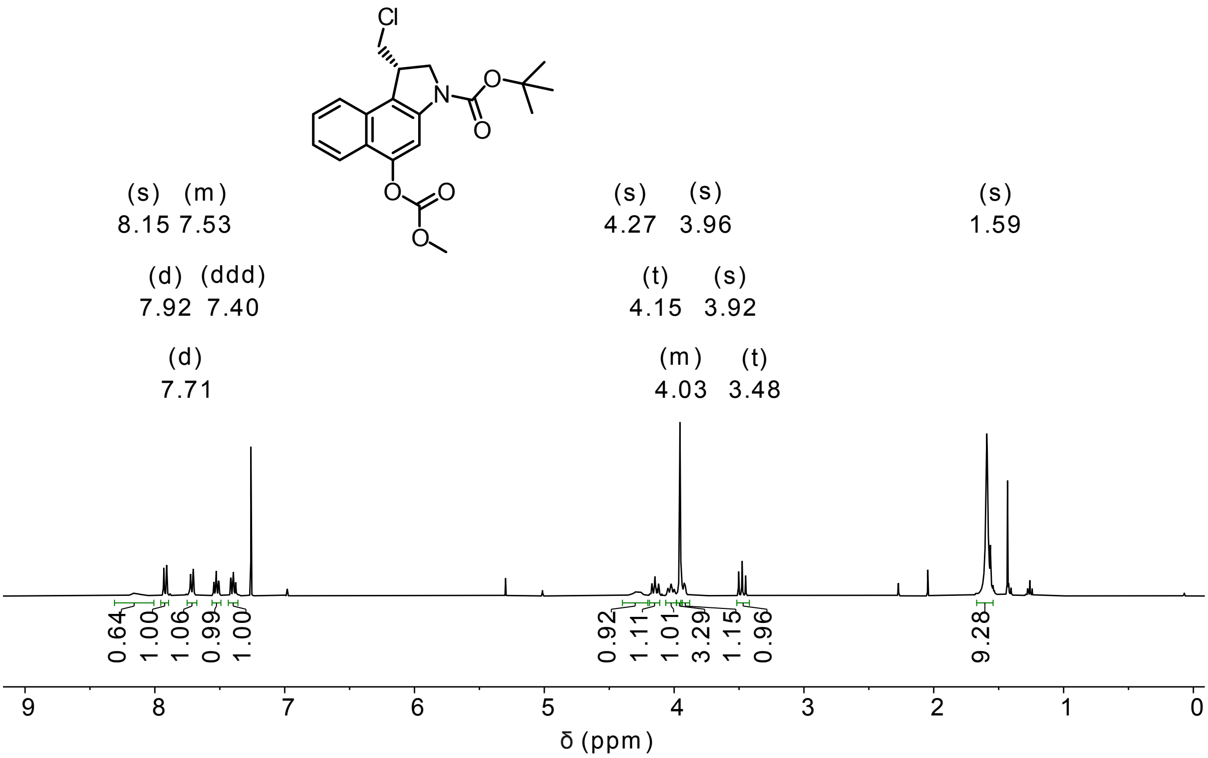


$^{13}$C-NMR

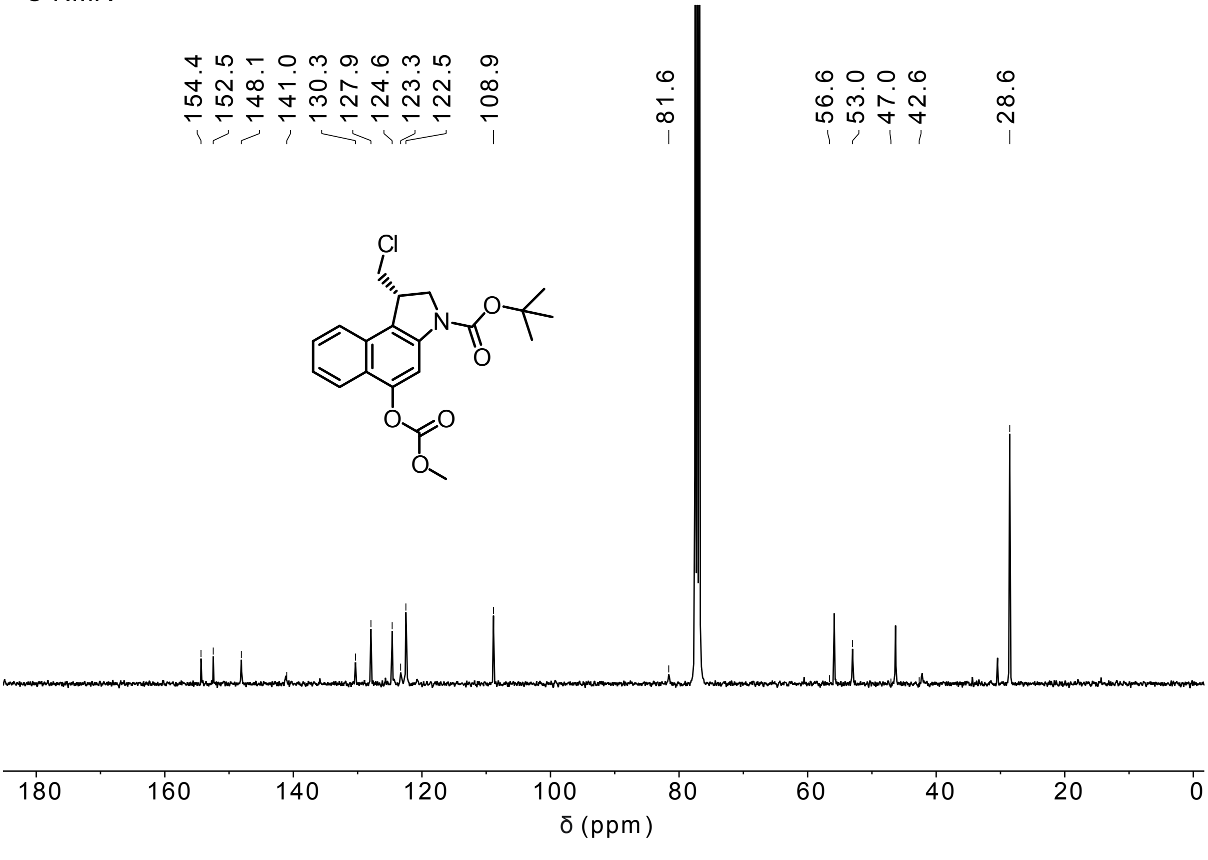

**O56-CBI-Boc**

$^1$H-NMR

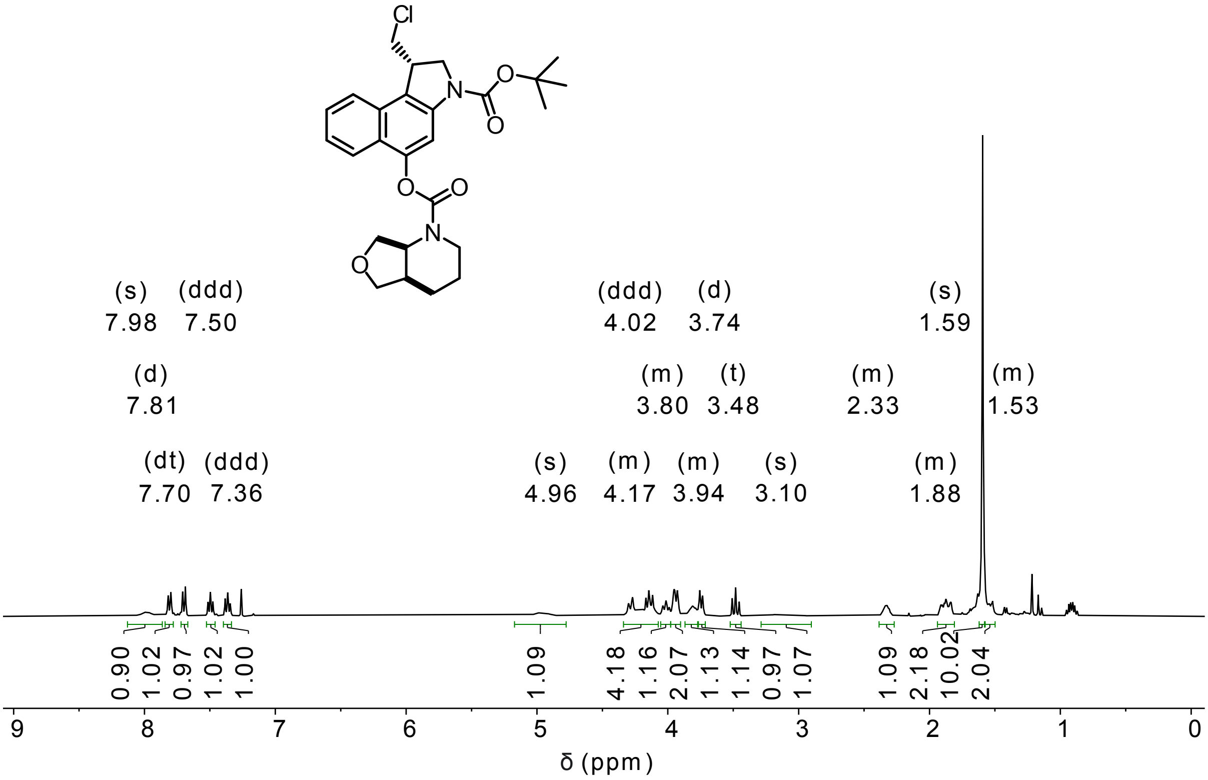


$^{13}$C-NMR

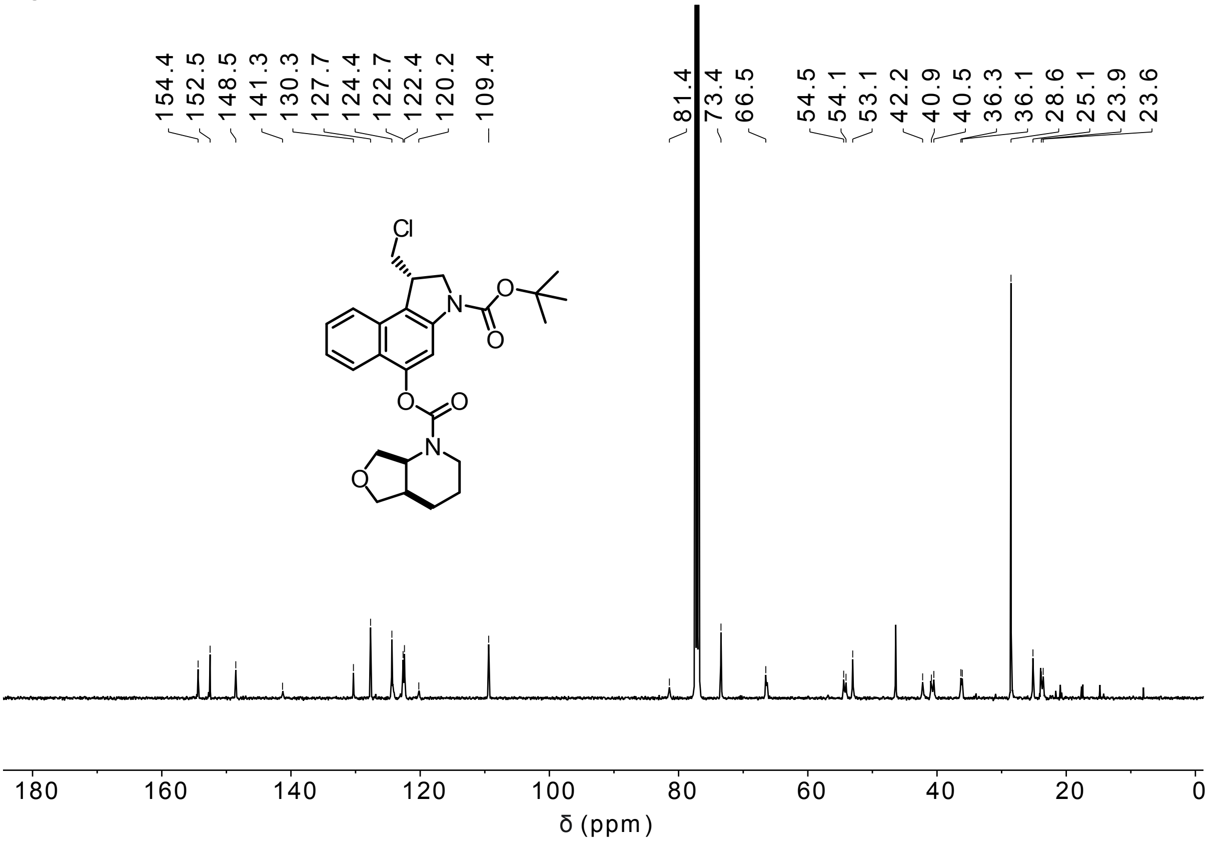

**SS60-CBI-AZI**

$^{1}$H-NMR

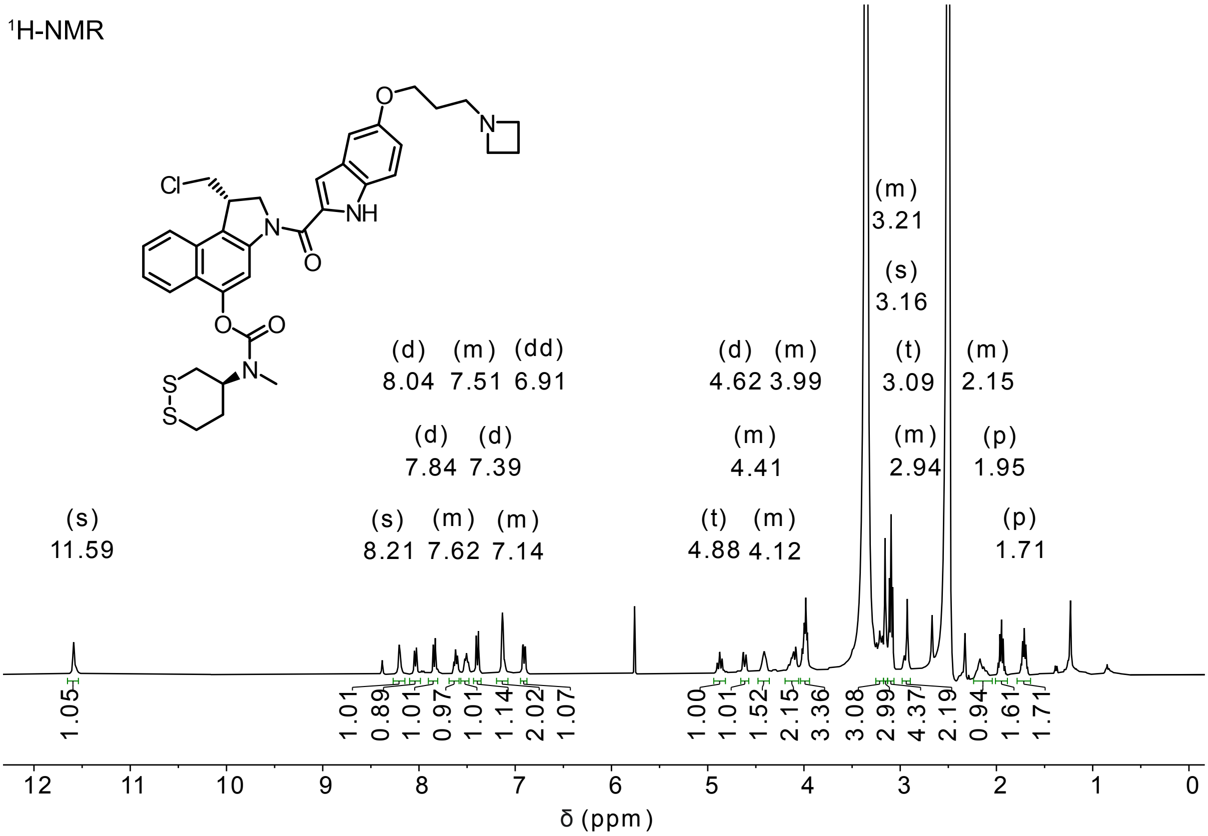


$^{13}$C-NMR

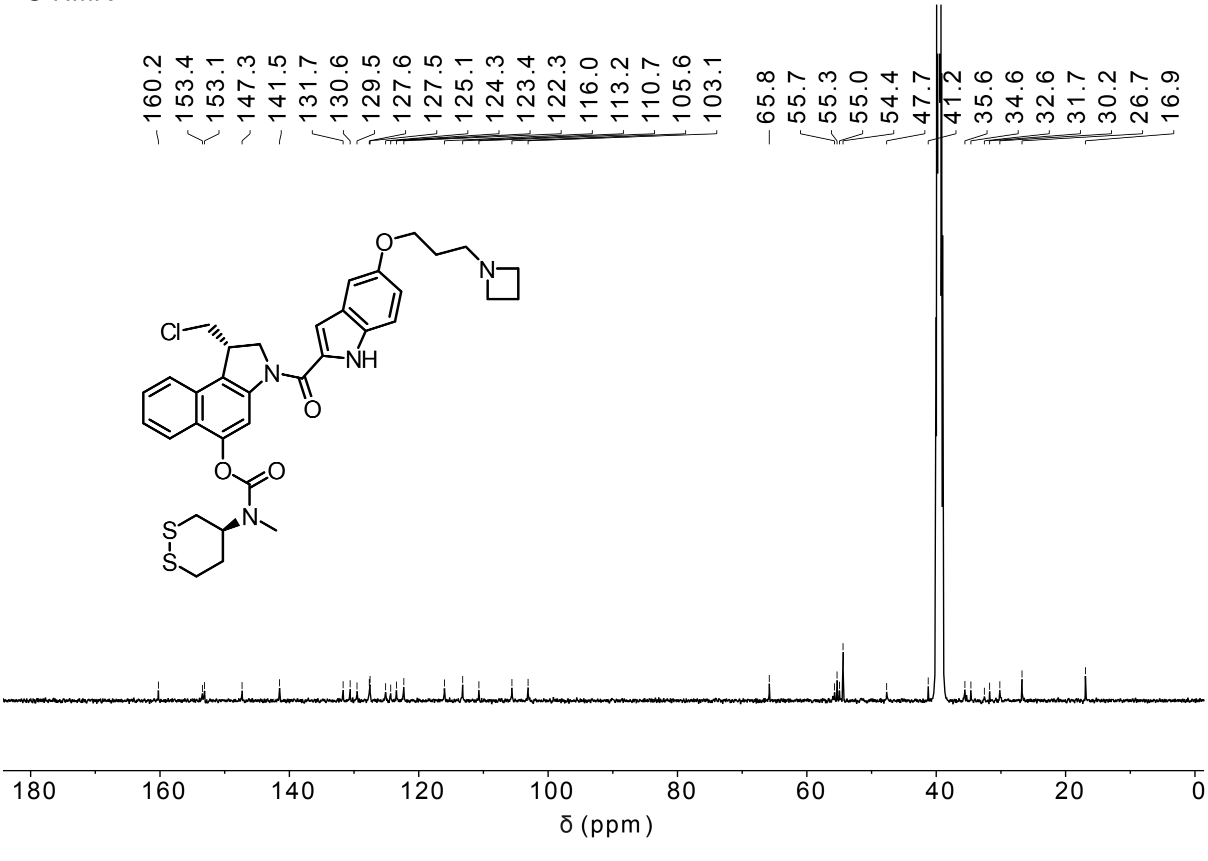

**P-SS60-CBI-AZI**

$^{1}$H-NMR

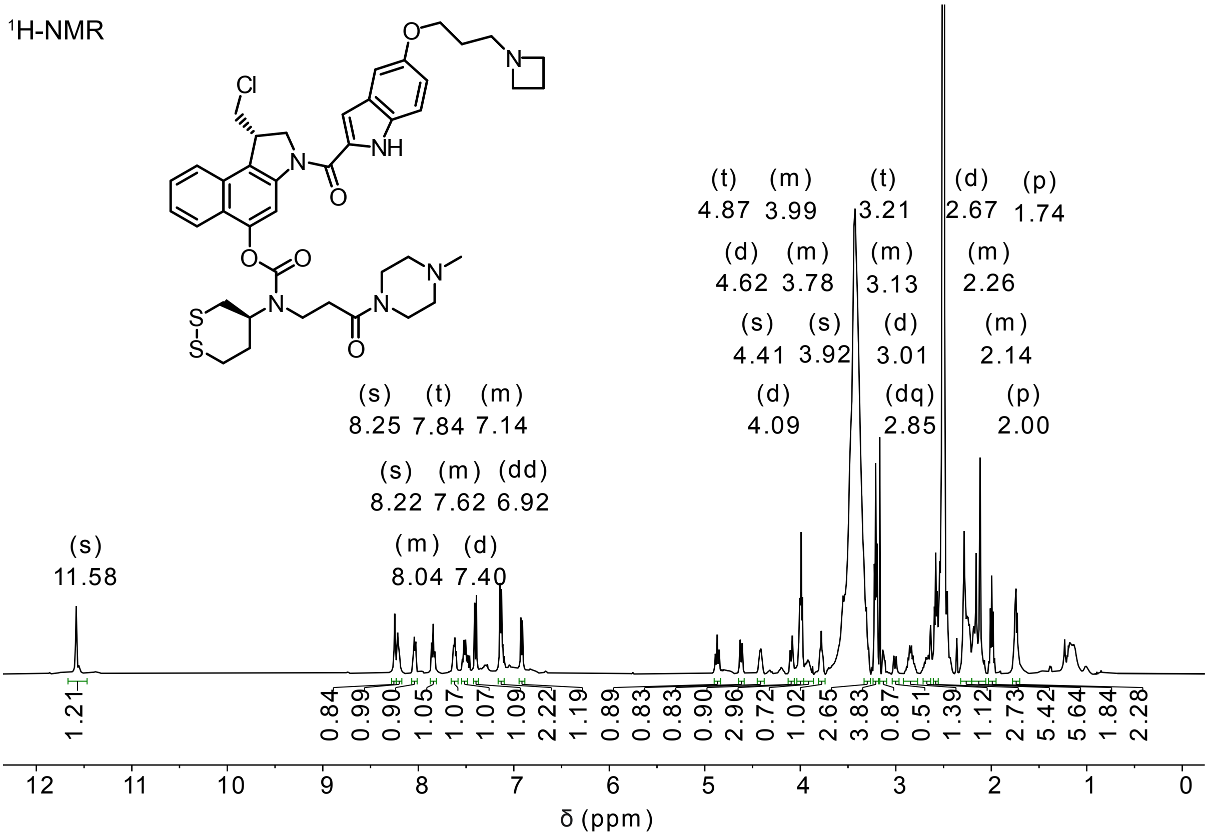


$^{13}$C-NMR

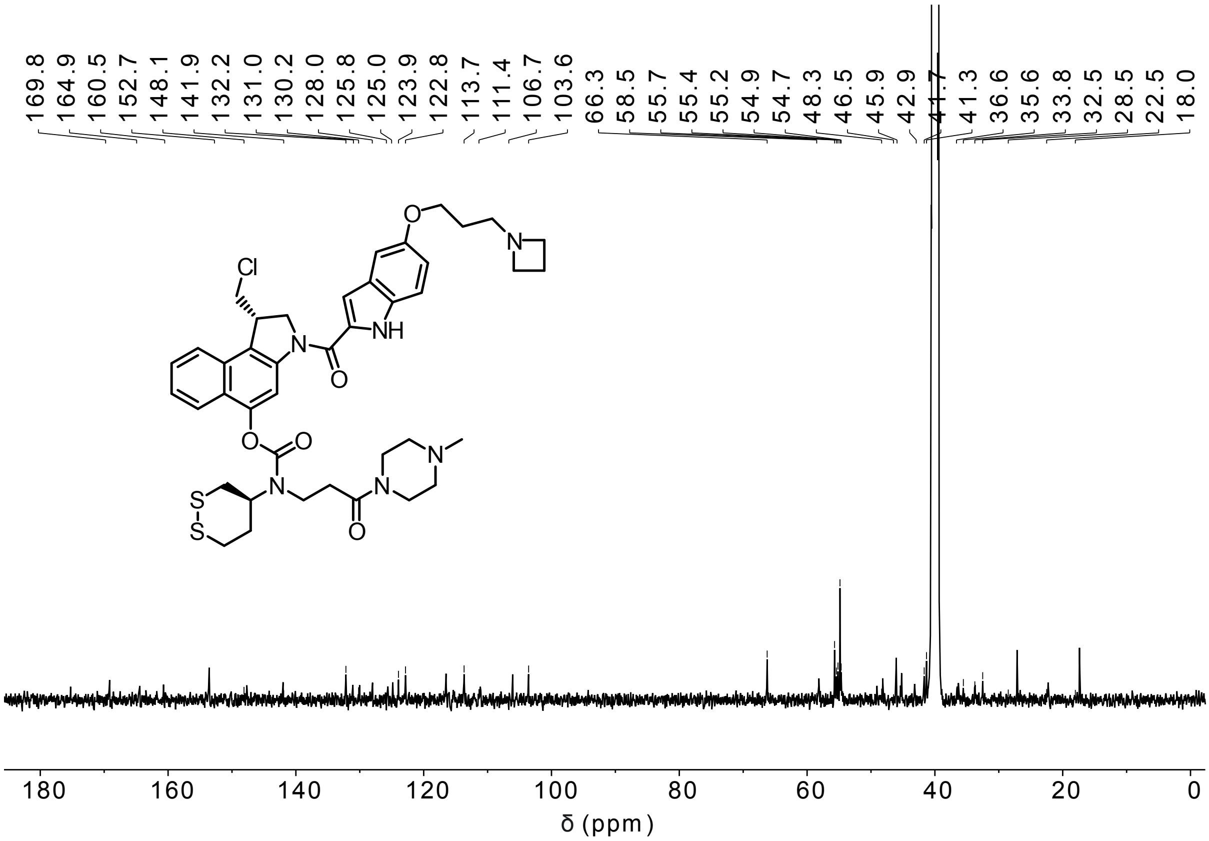

**SeS60-CBI-AZI**

$^{1}$H-NMR

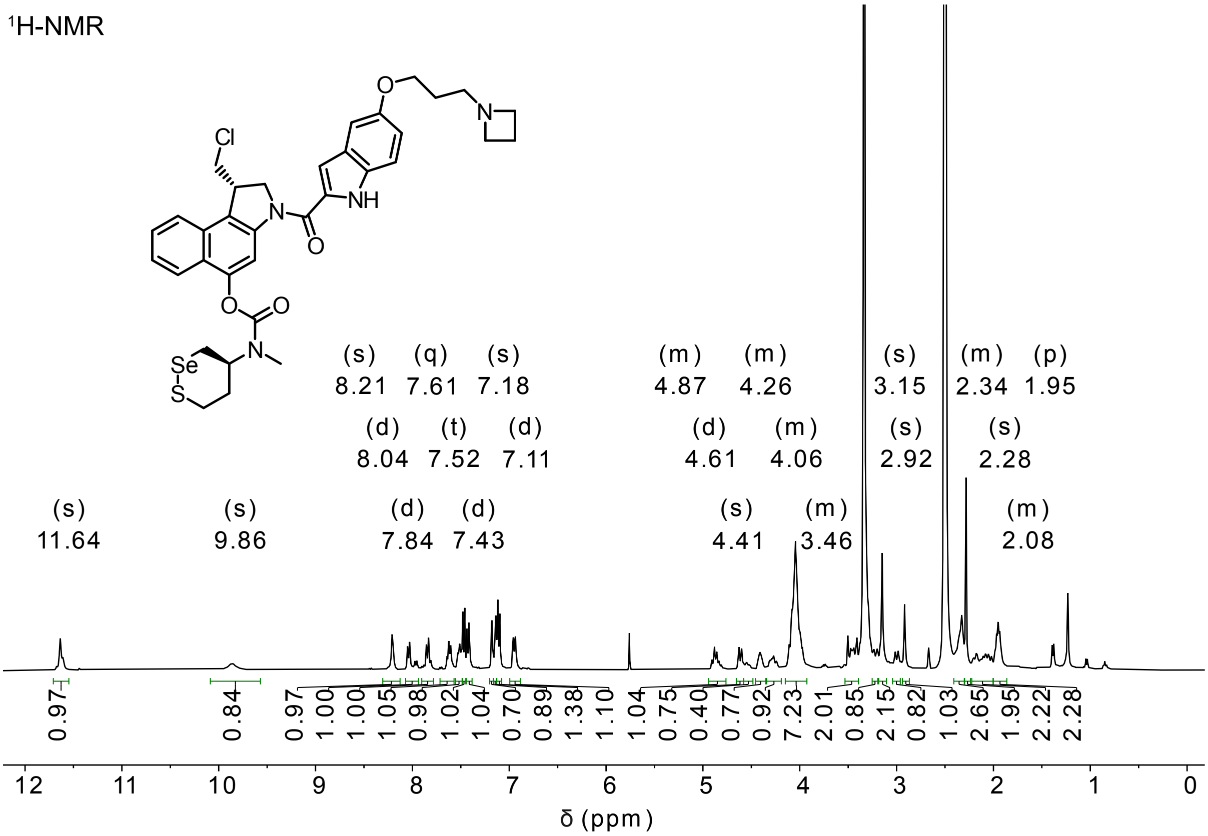


$^{13}$C-NMR

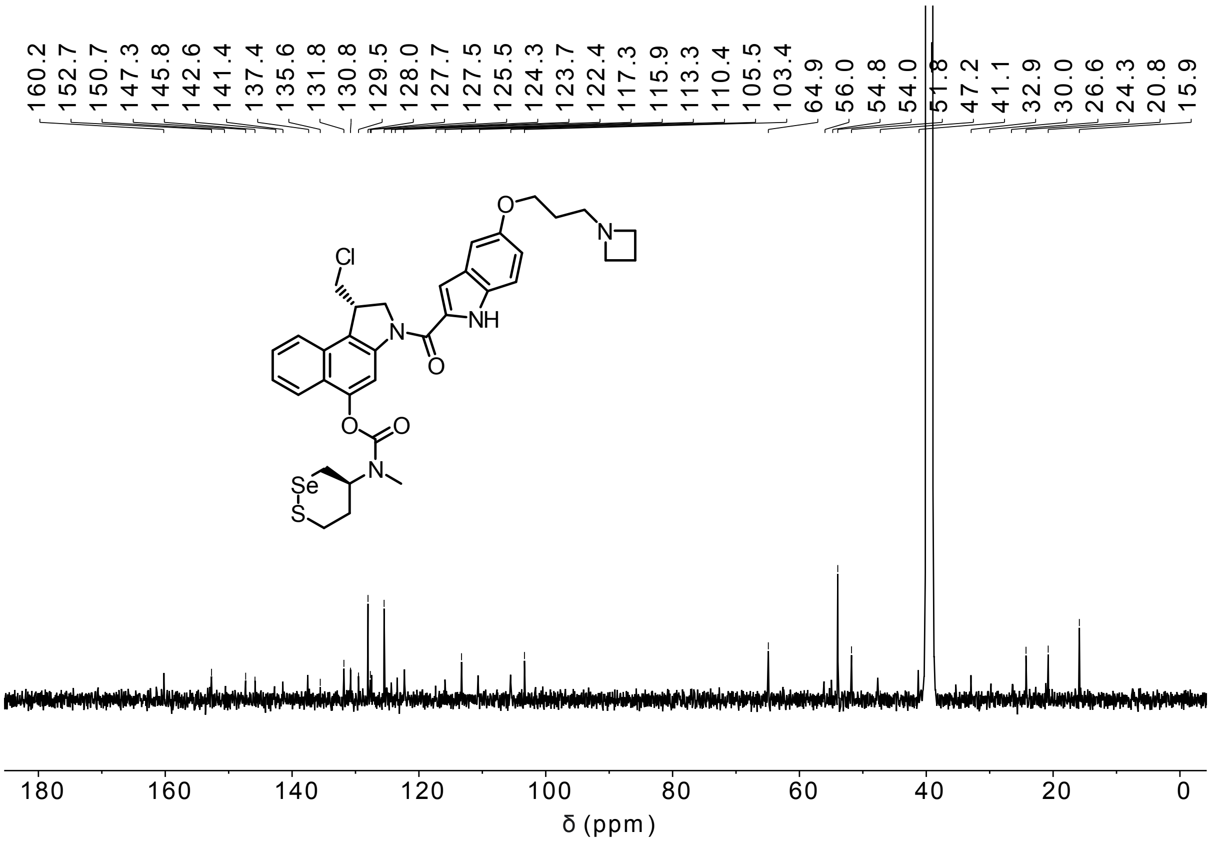

**Me-SS66T-CBI-AZI**

$^{1}$H-NMR

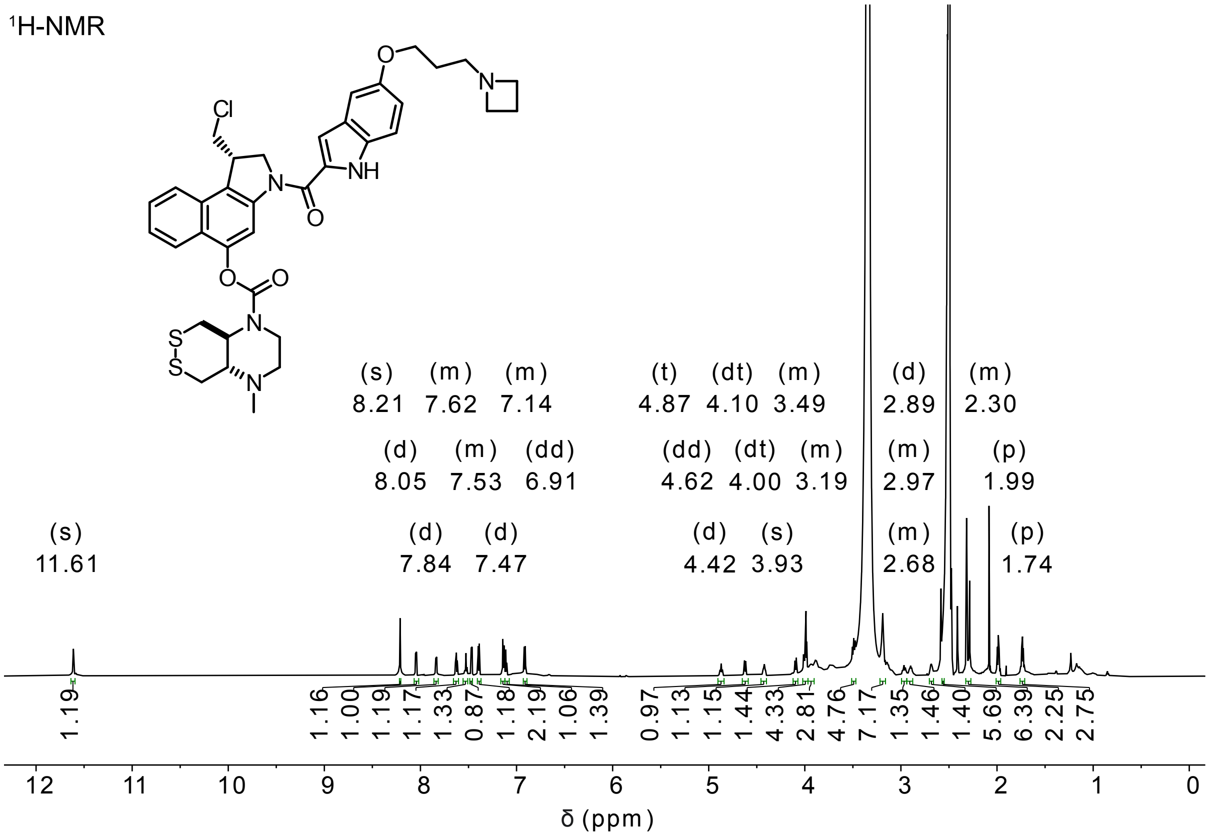


$^{13}$C-NMR

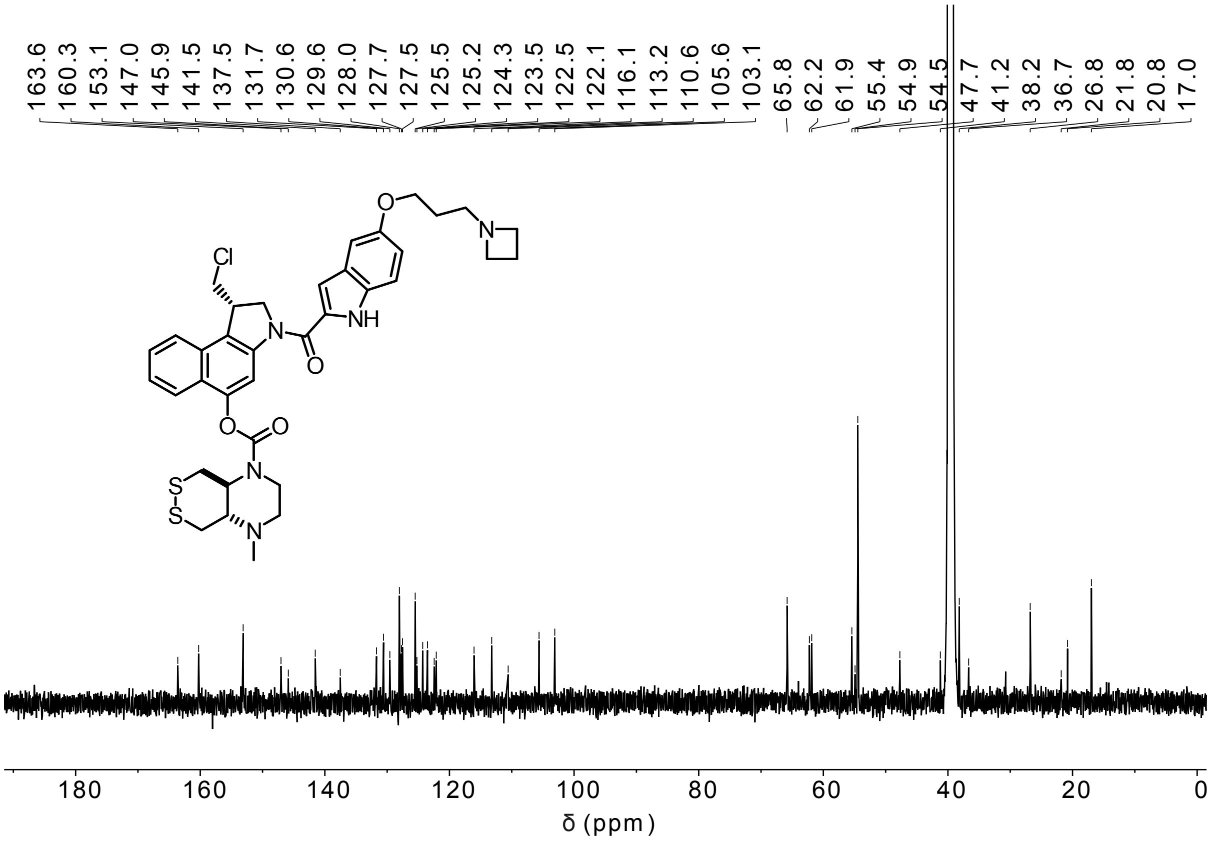

**Me-SS66C-CBI-AZI**

$^{1}$H-NMR

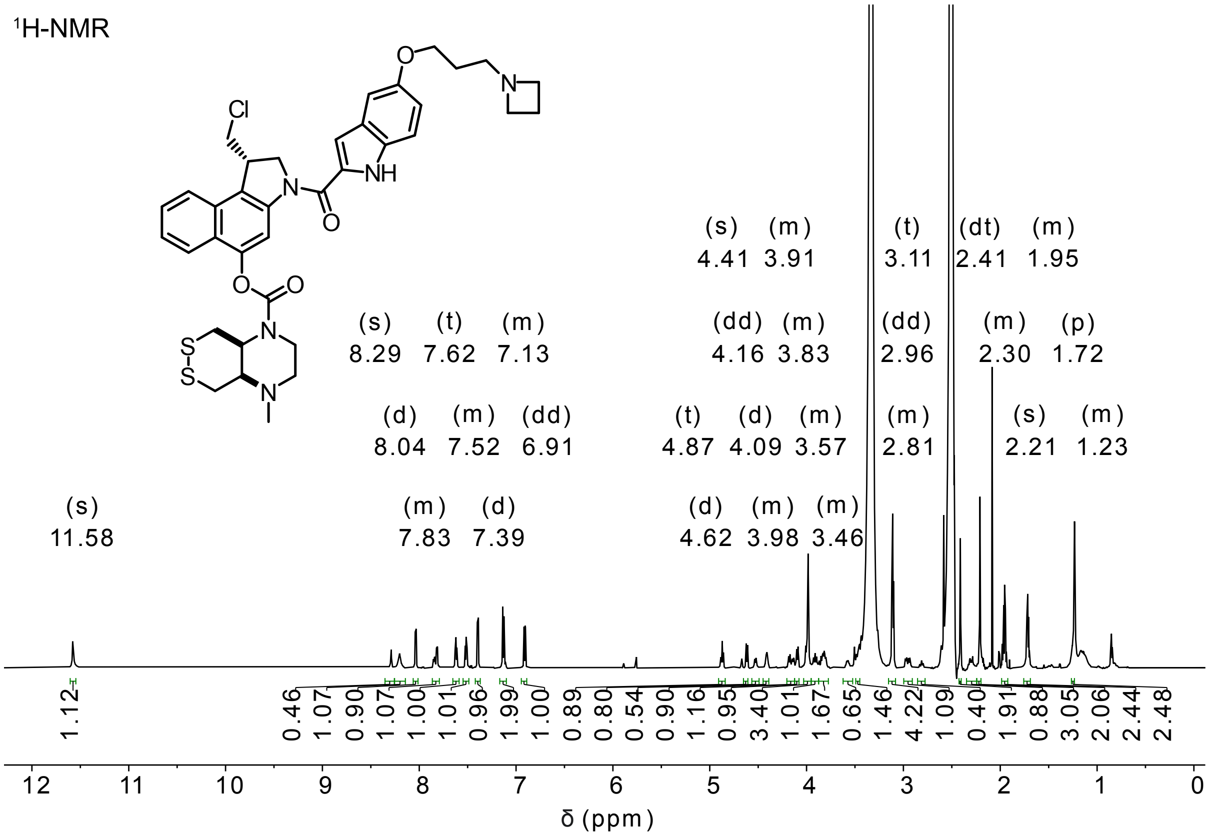


$^{13}$C-NMR

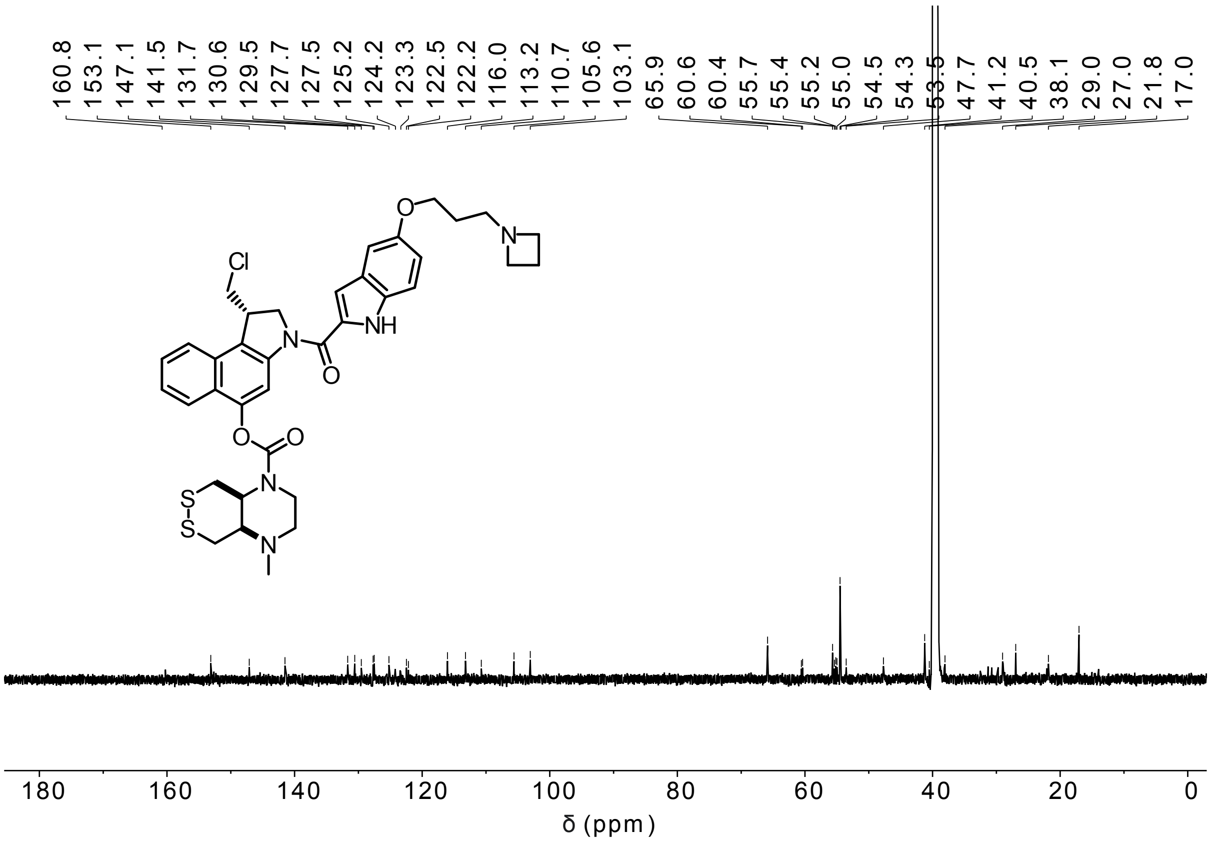

**SS00$_M$-CBI-AZI**

$^1$H-NMR

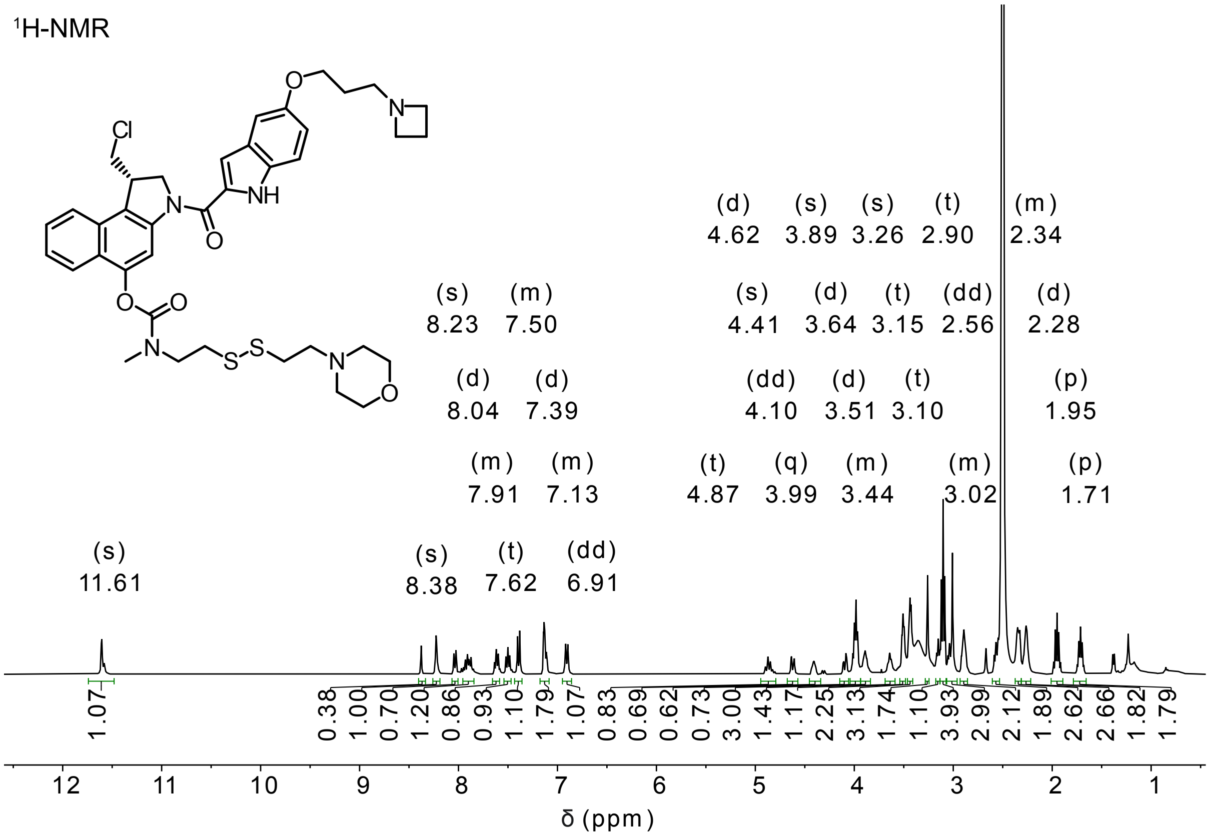


$^1$H/$^{13}$C-HSQC

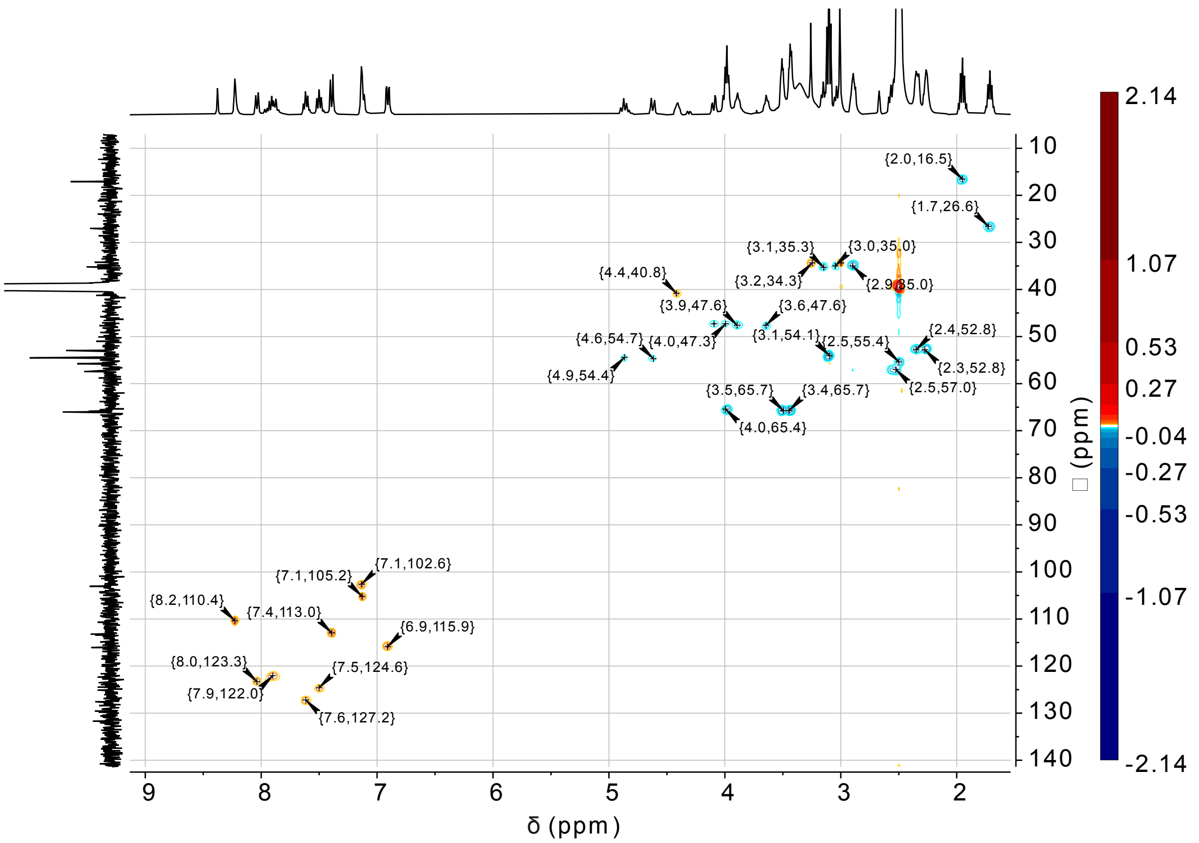

**MC-CBI-AZI**

[1]H-NMR

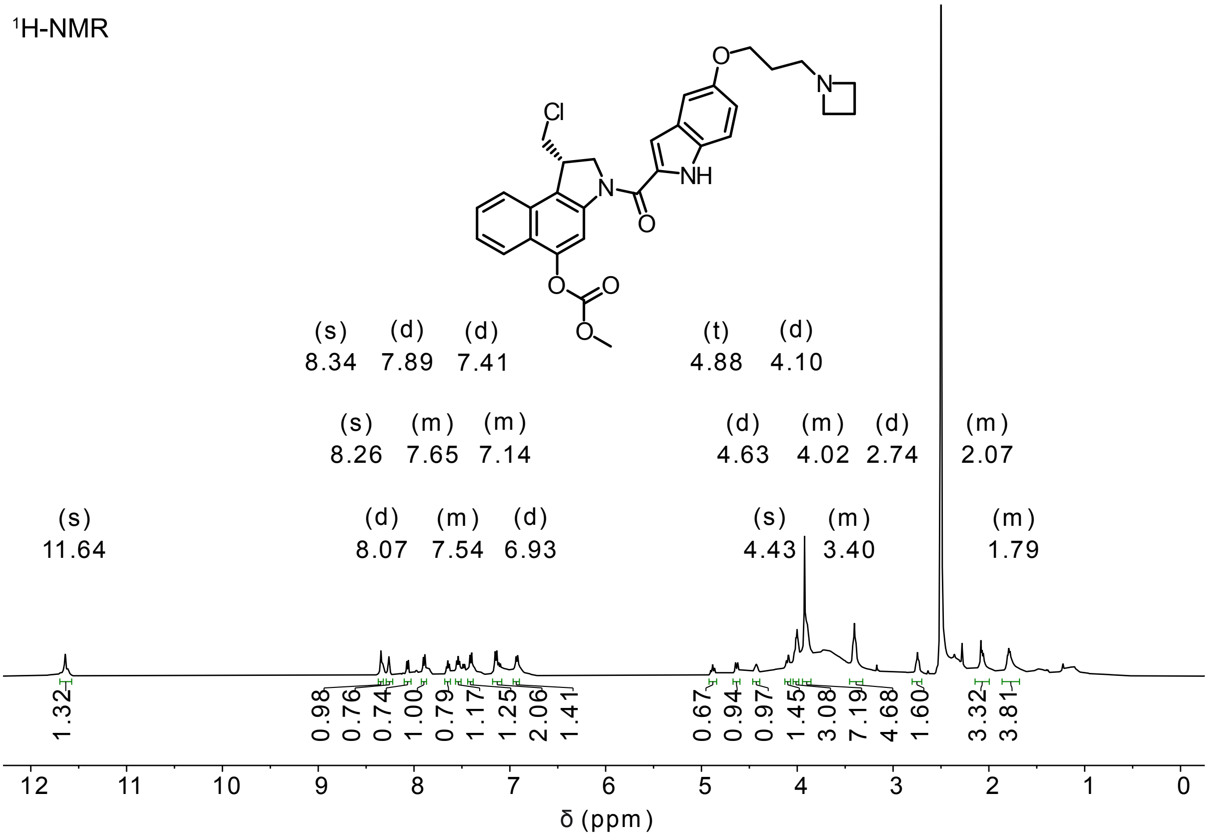


[13]C-NMR

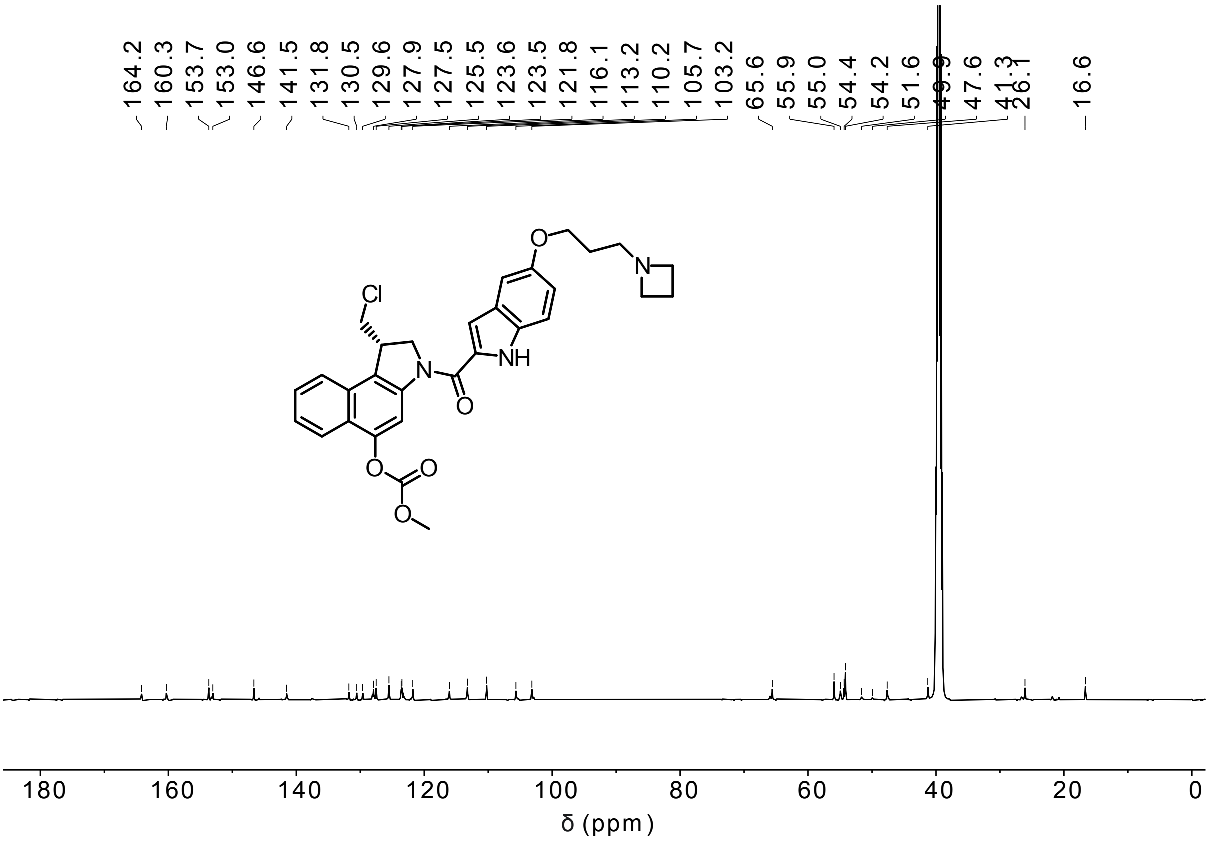

**O56-CBI-AZI**

[1]H-NMR

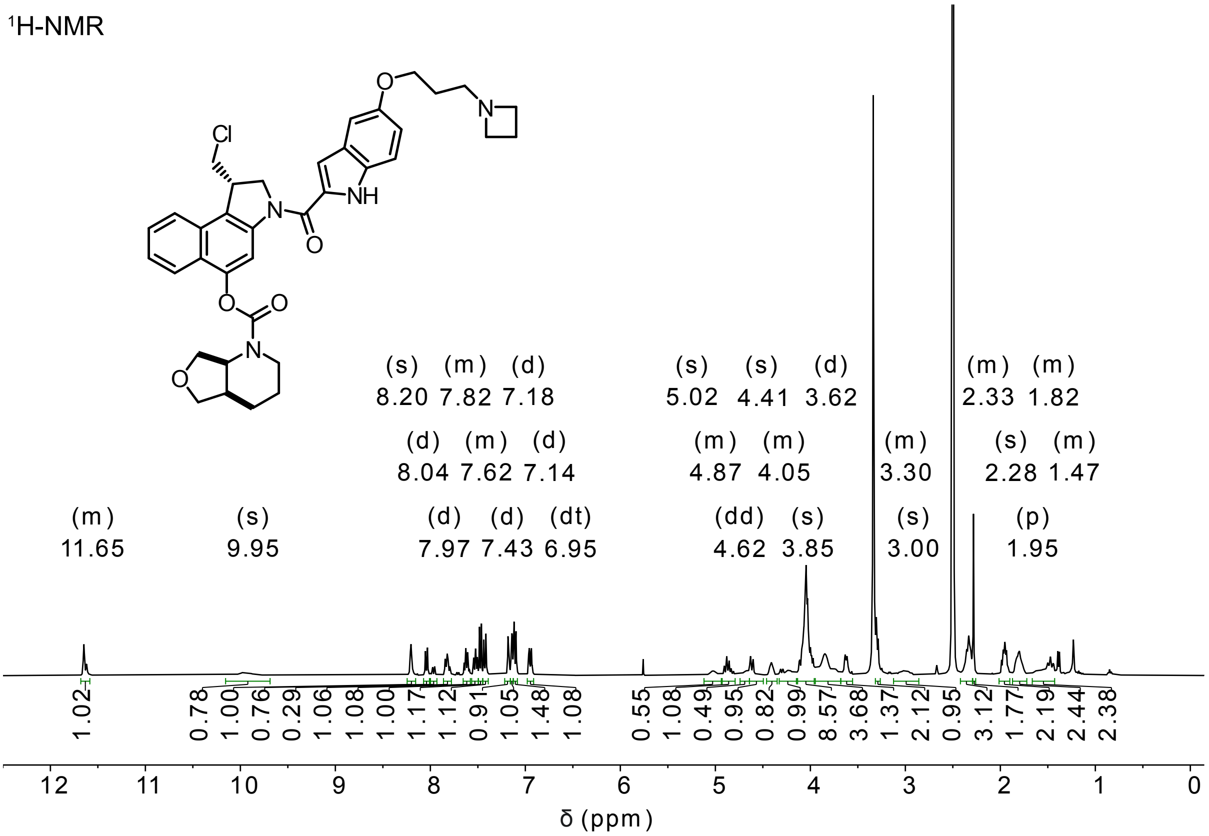


[13]C-NMR

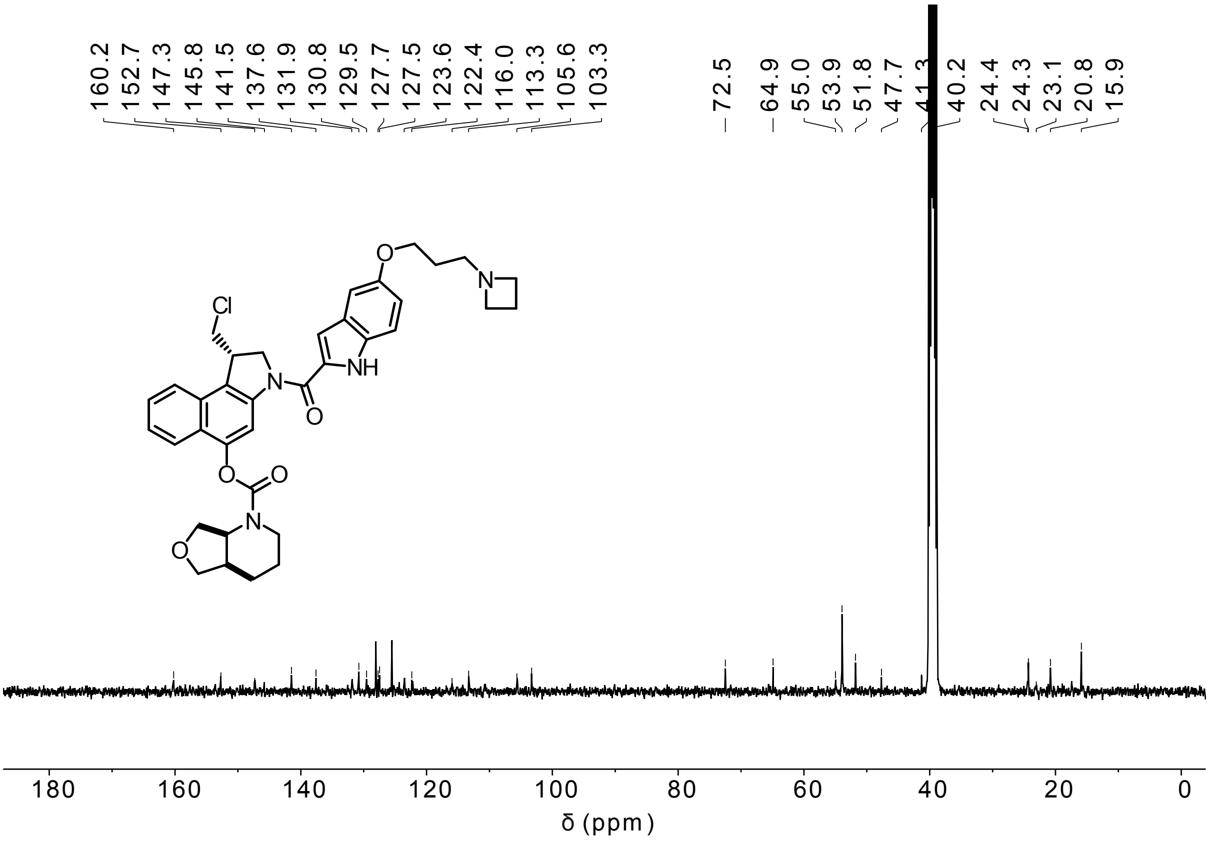

**OBn-CBI-AZI**

[1]H-NMR

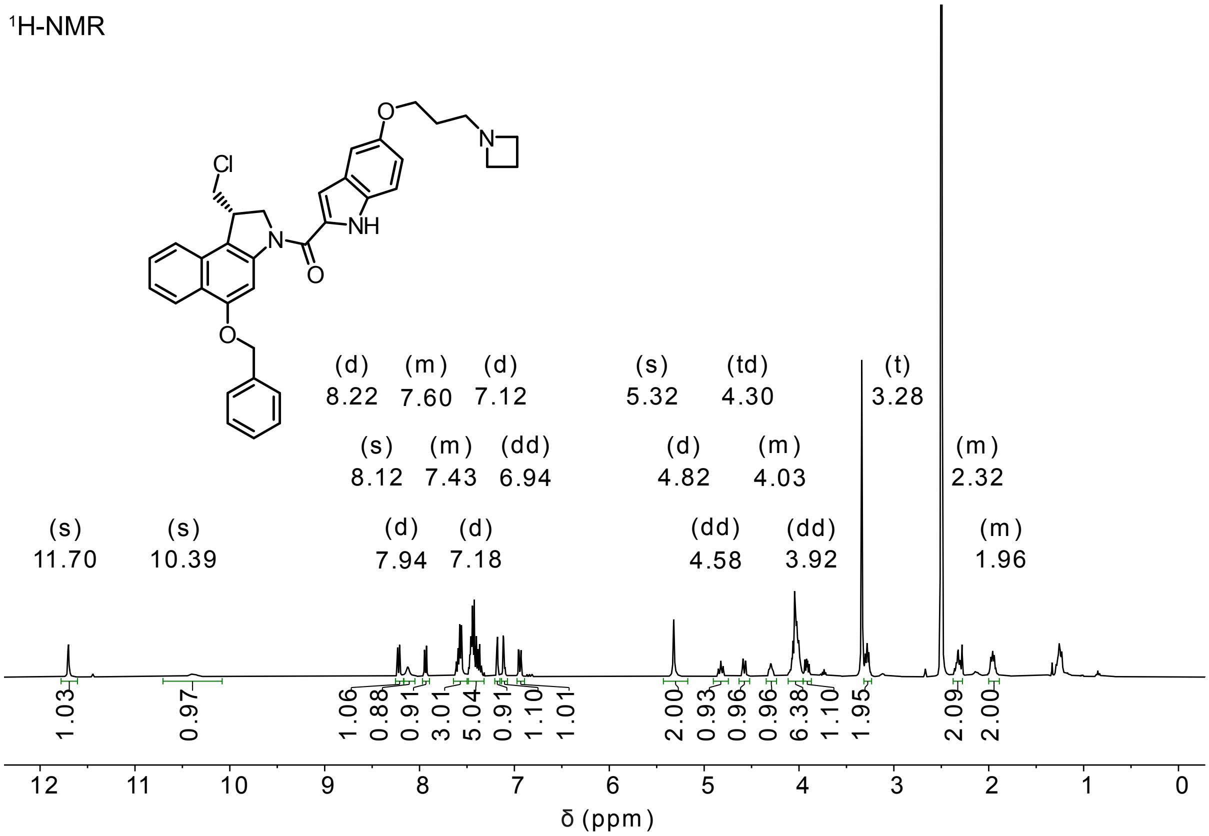


[13]C-NMR

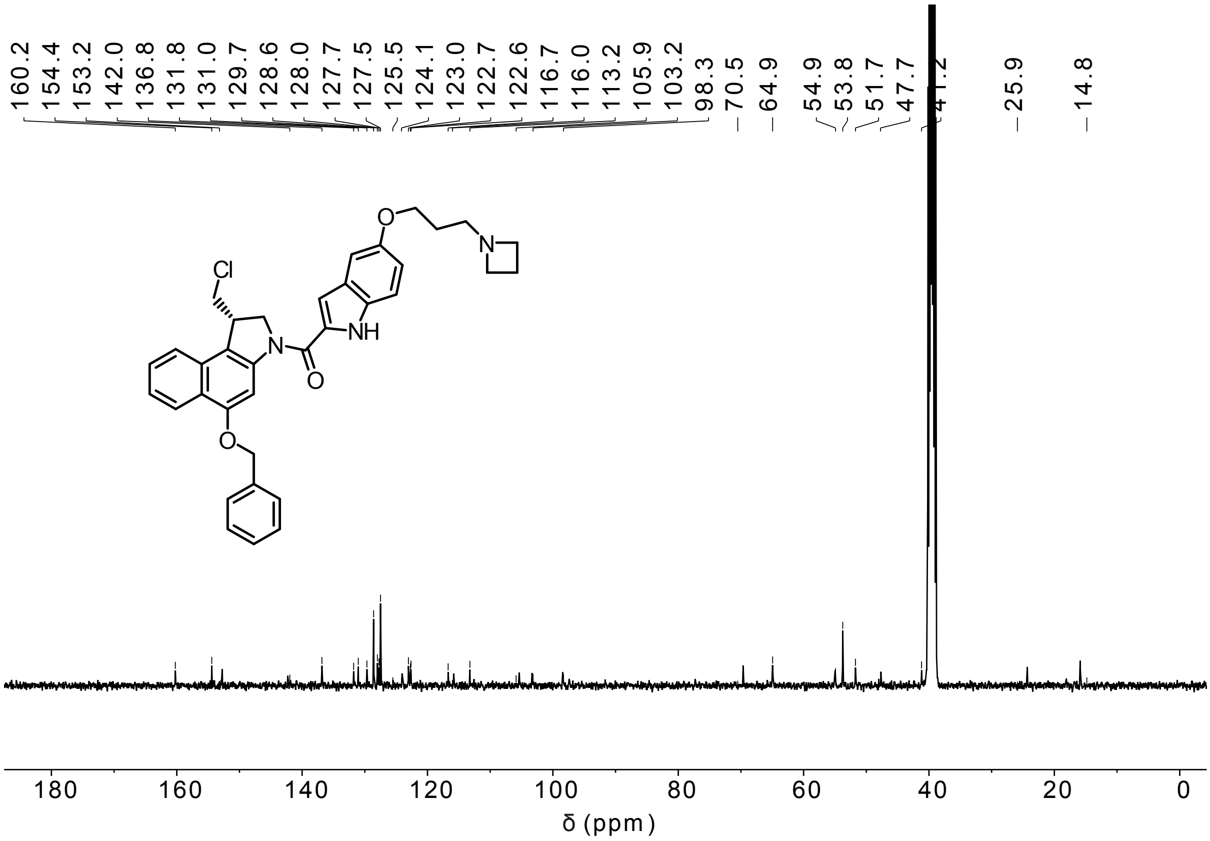

**P-SS60-CBI-TMI**

$^1$H-NMR

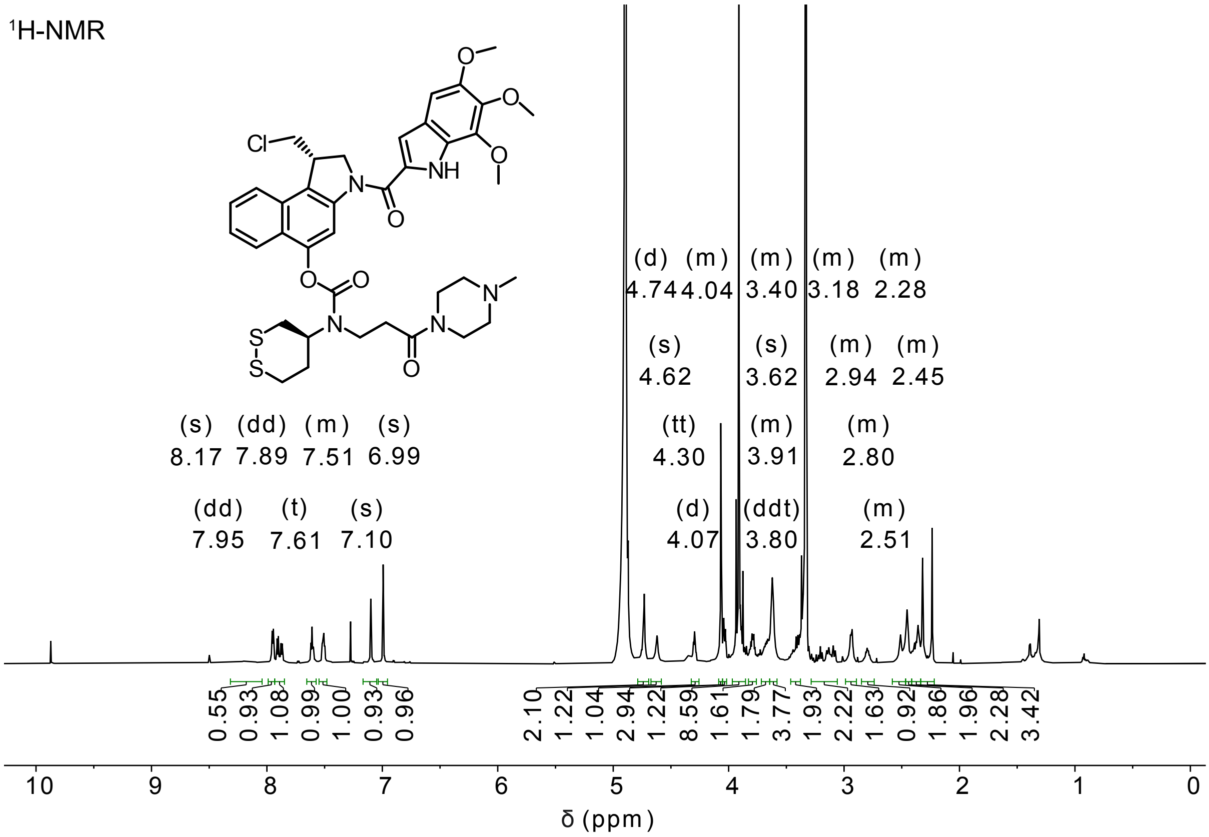


$^{13}$C-NMR

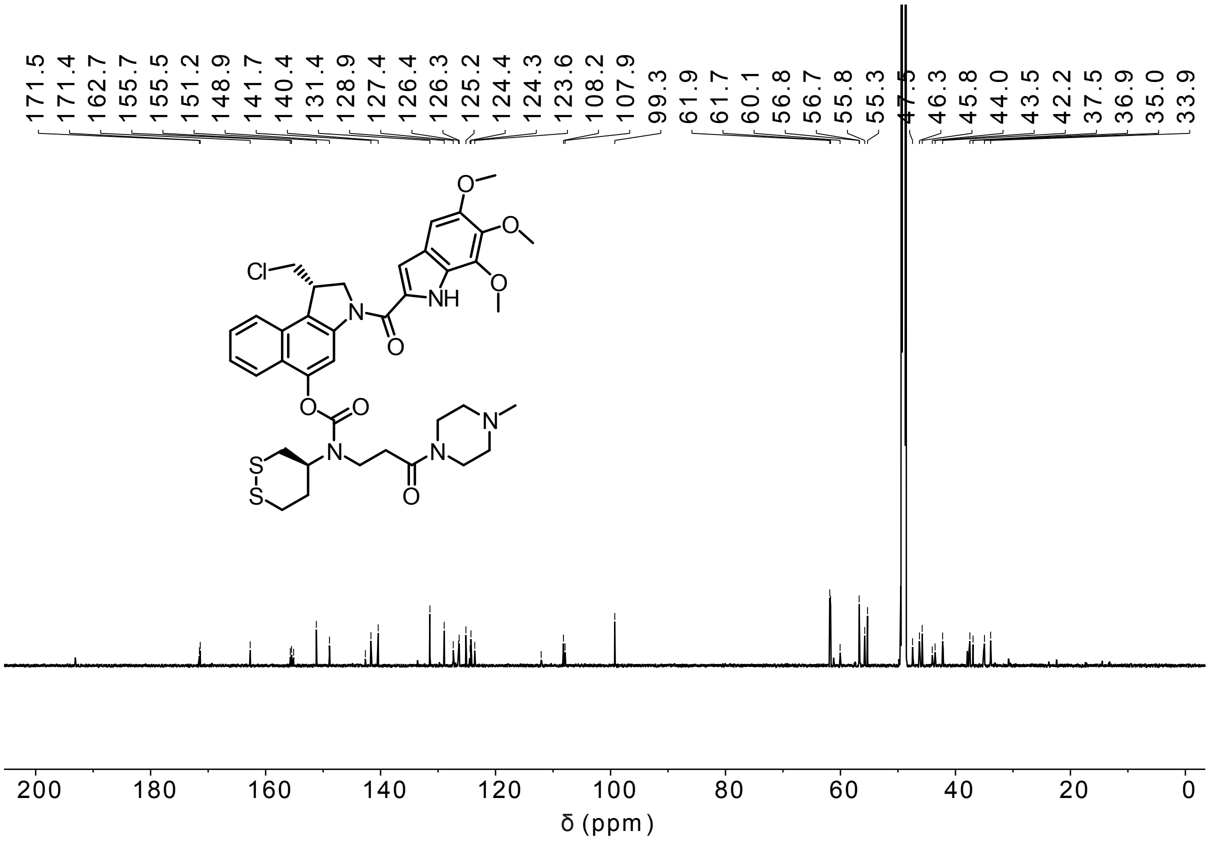

**Me-SS66T-CBI-TMI**

[1]H-NMR

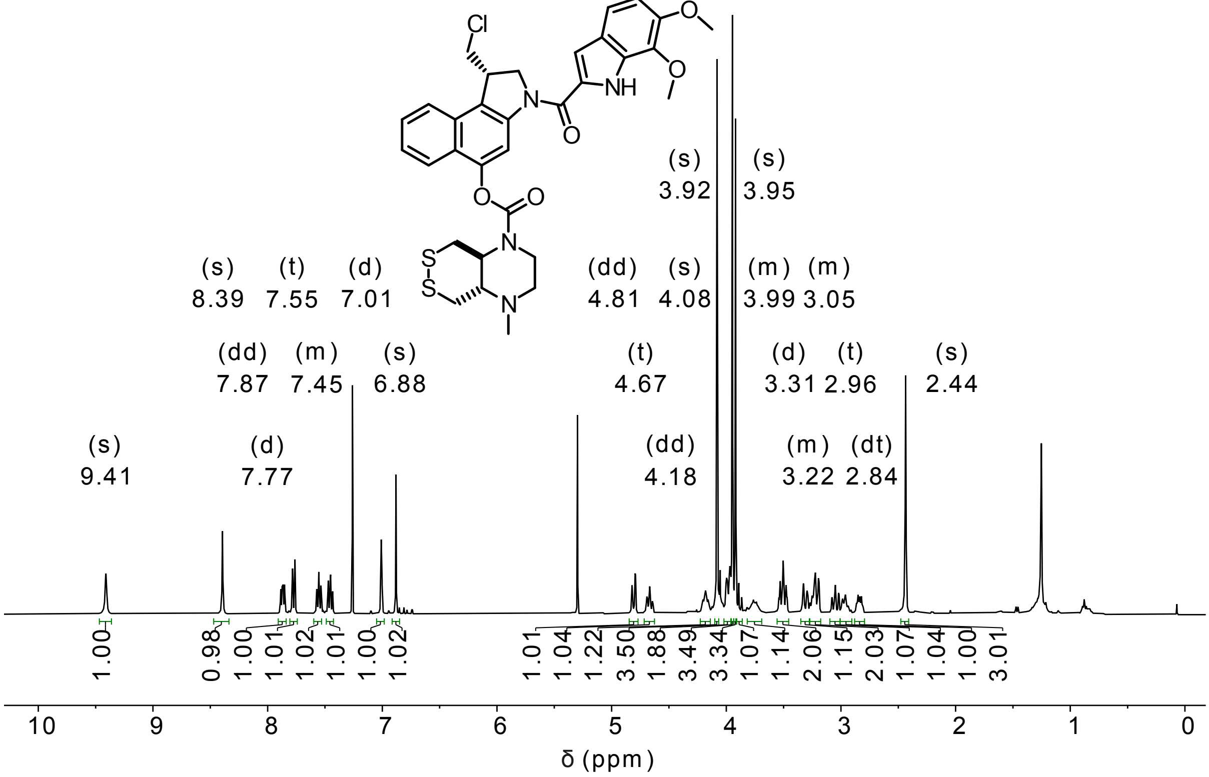


[13]C-NMR

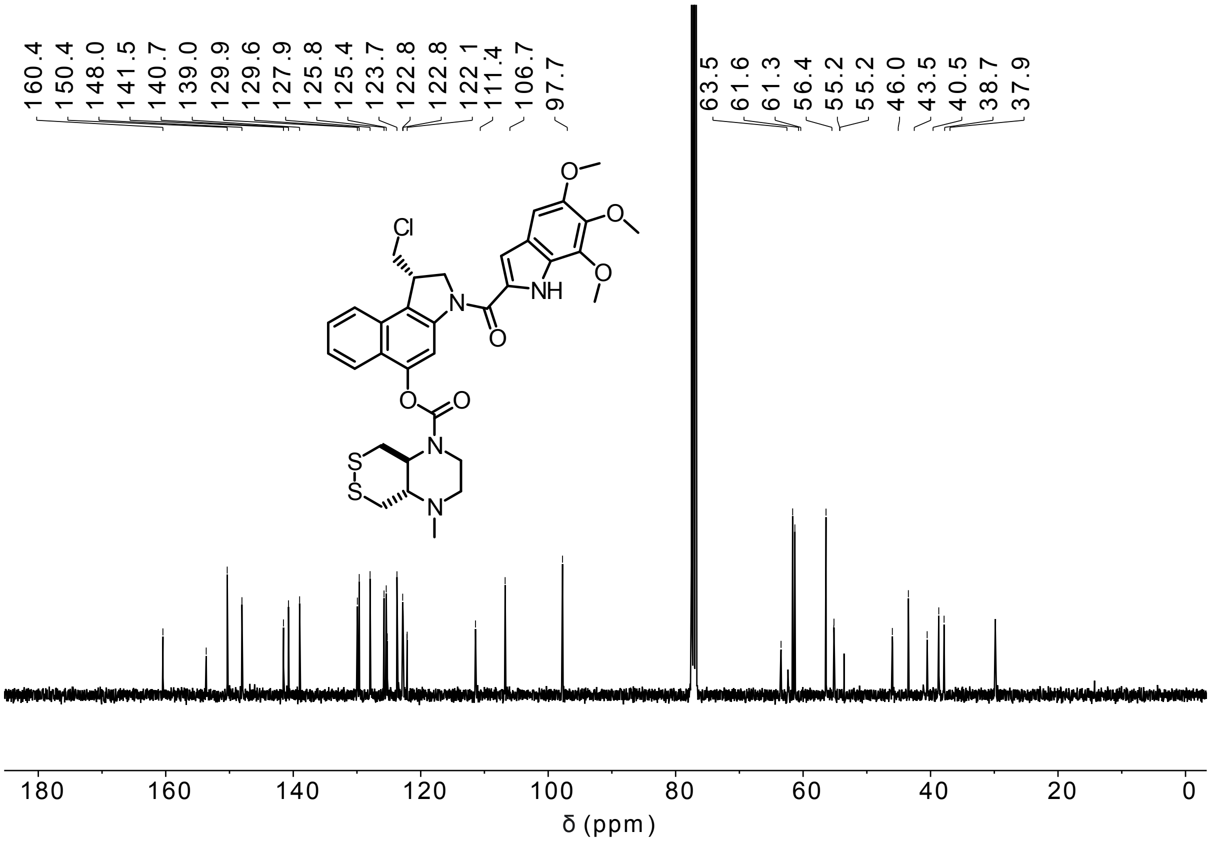

HSQC - aromatic signals

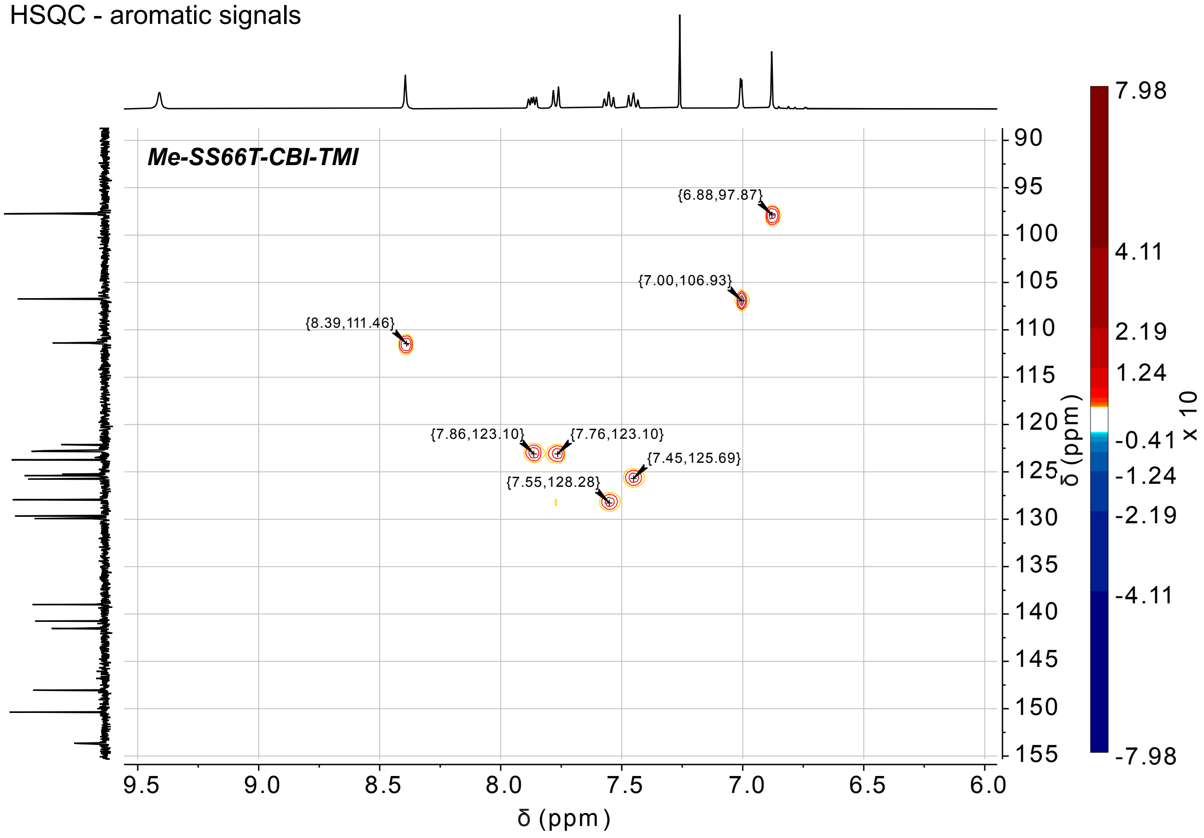


HSQC - aliphatic signals

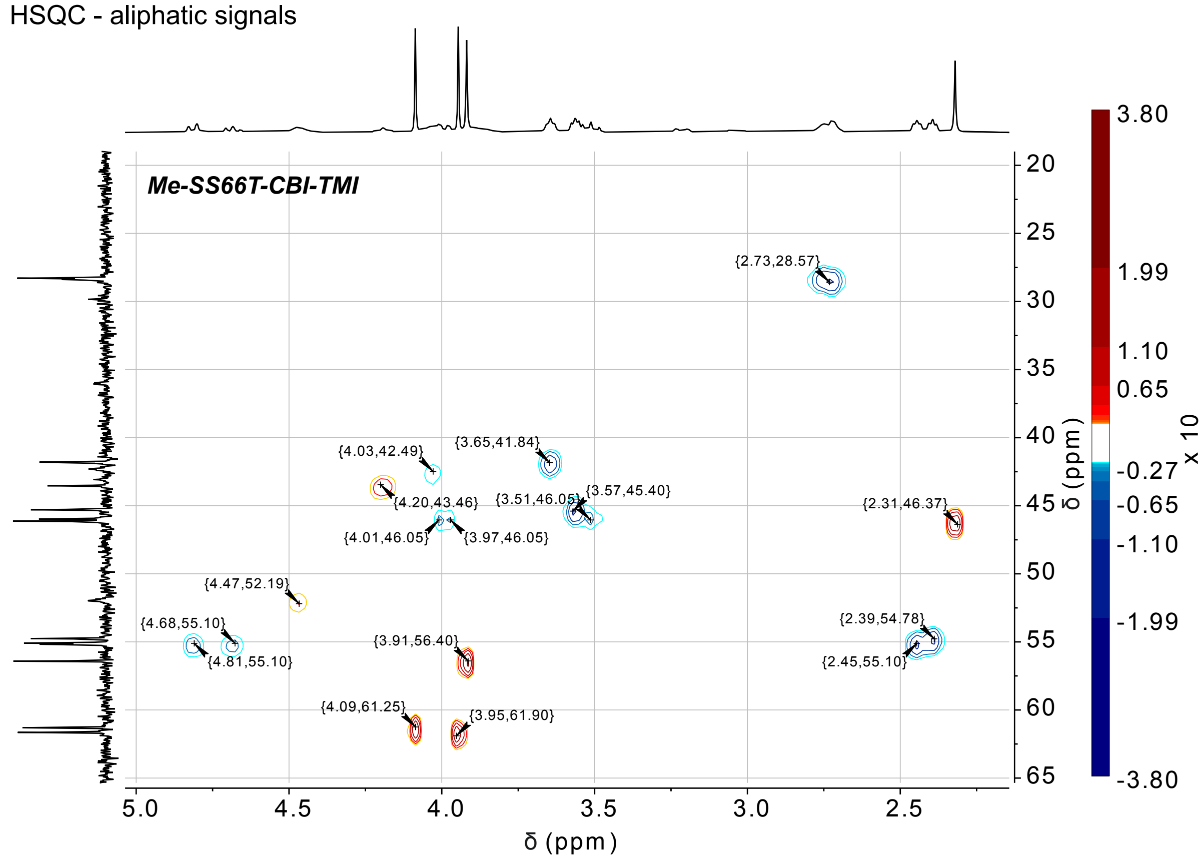

**Me-SS66C-CBI-TMI**

¹H-NMR

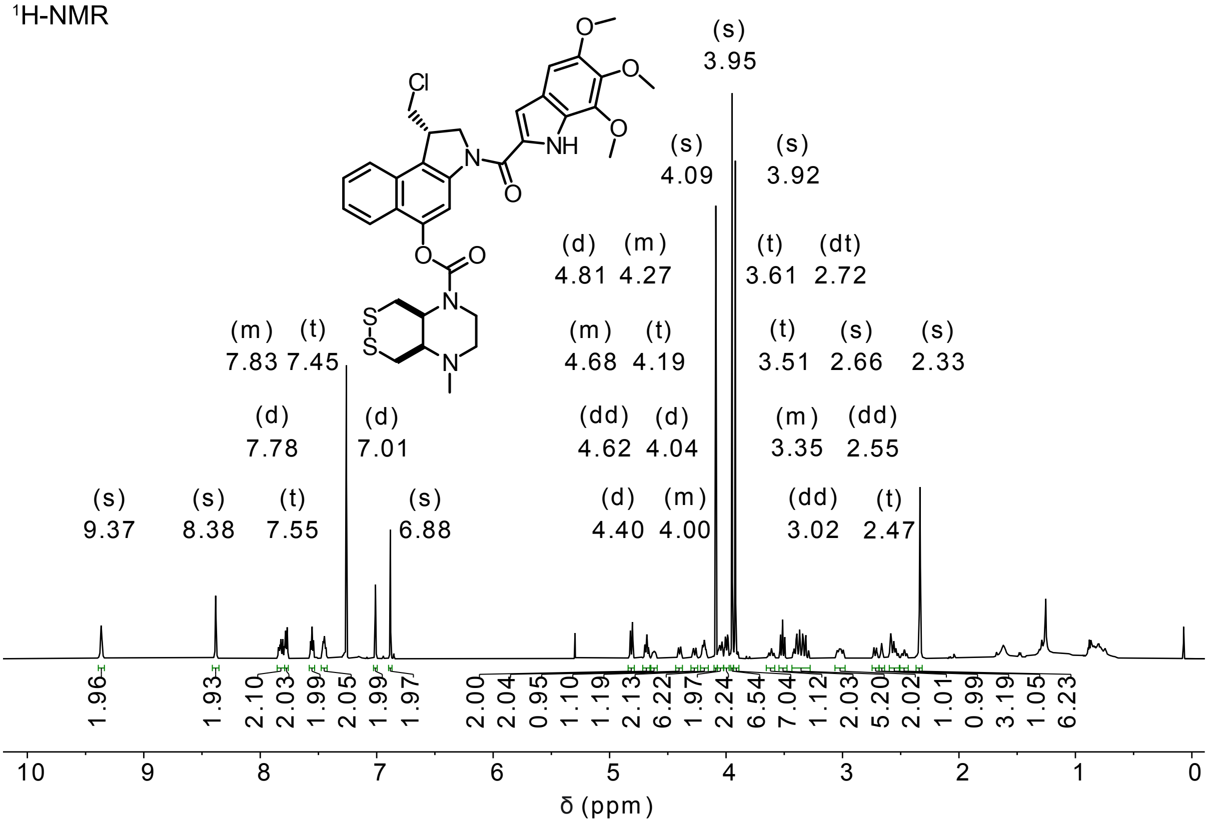


¹³C-NMR

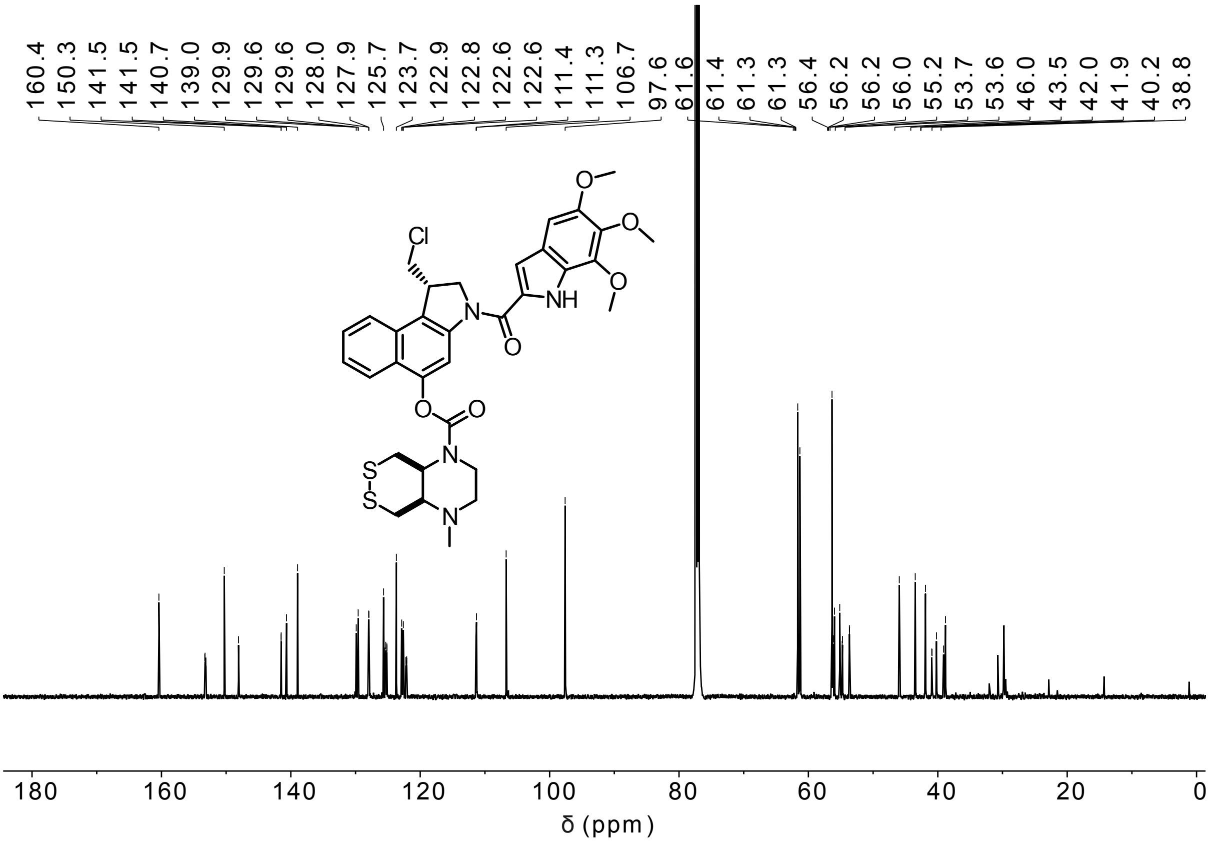

HSQC - aromatic signals

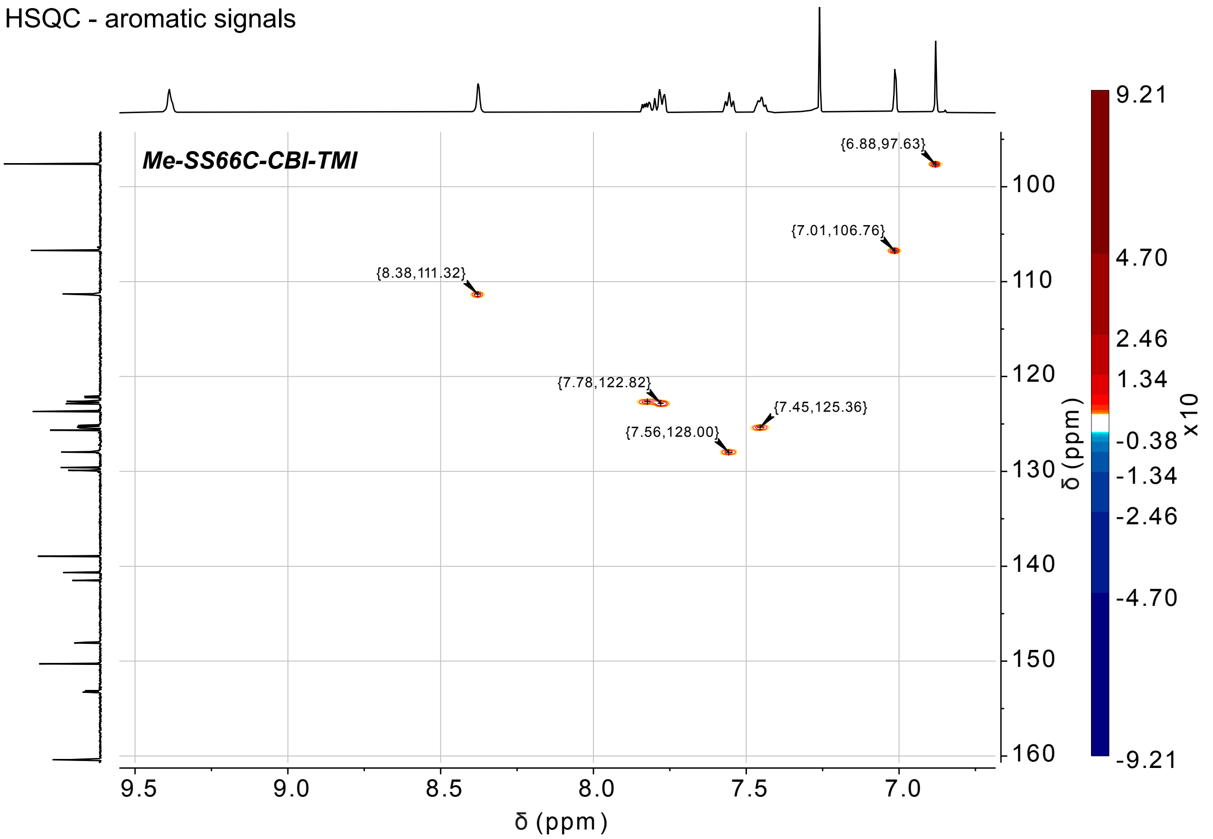


HSQC - aliphatic signals

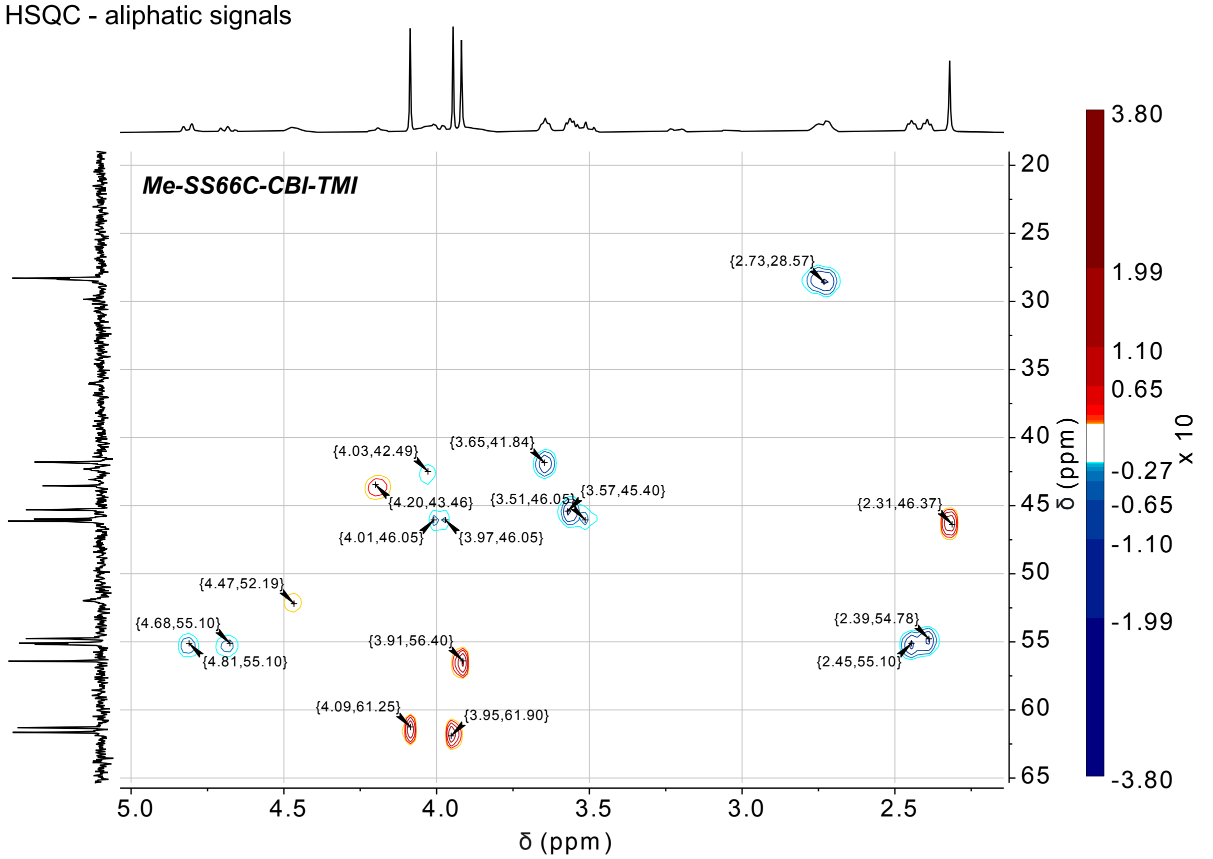

**P-SS66C-CBI-TMI**

[1]H-NMR

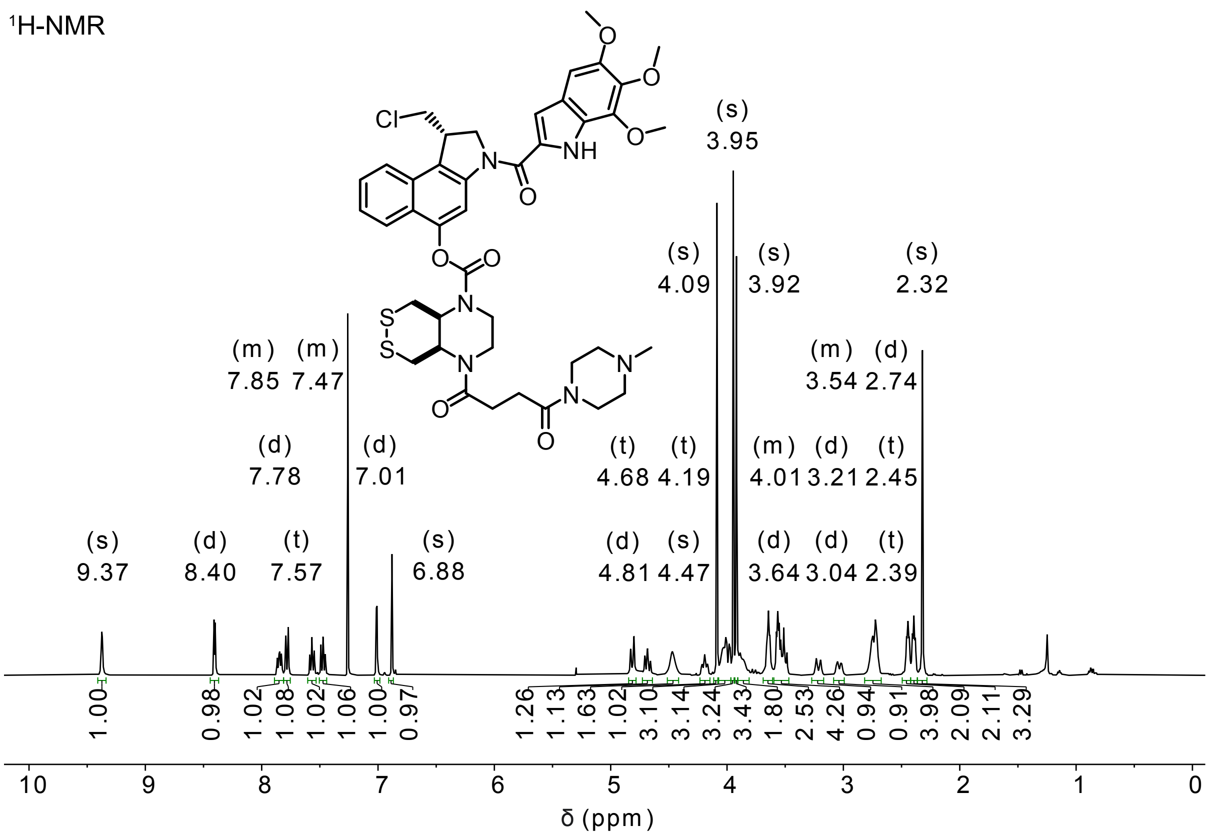


[13]C-NMR

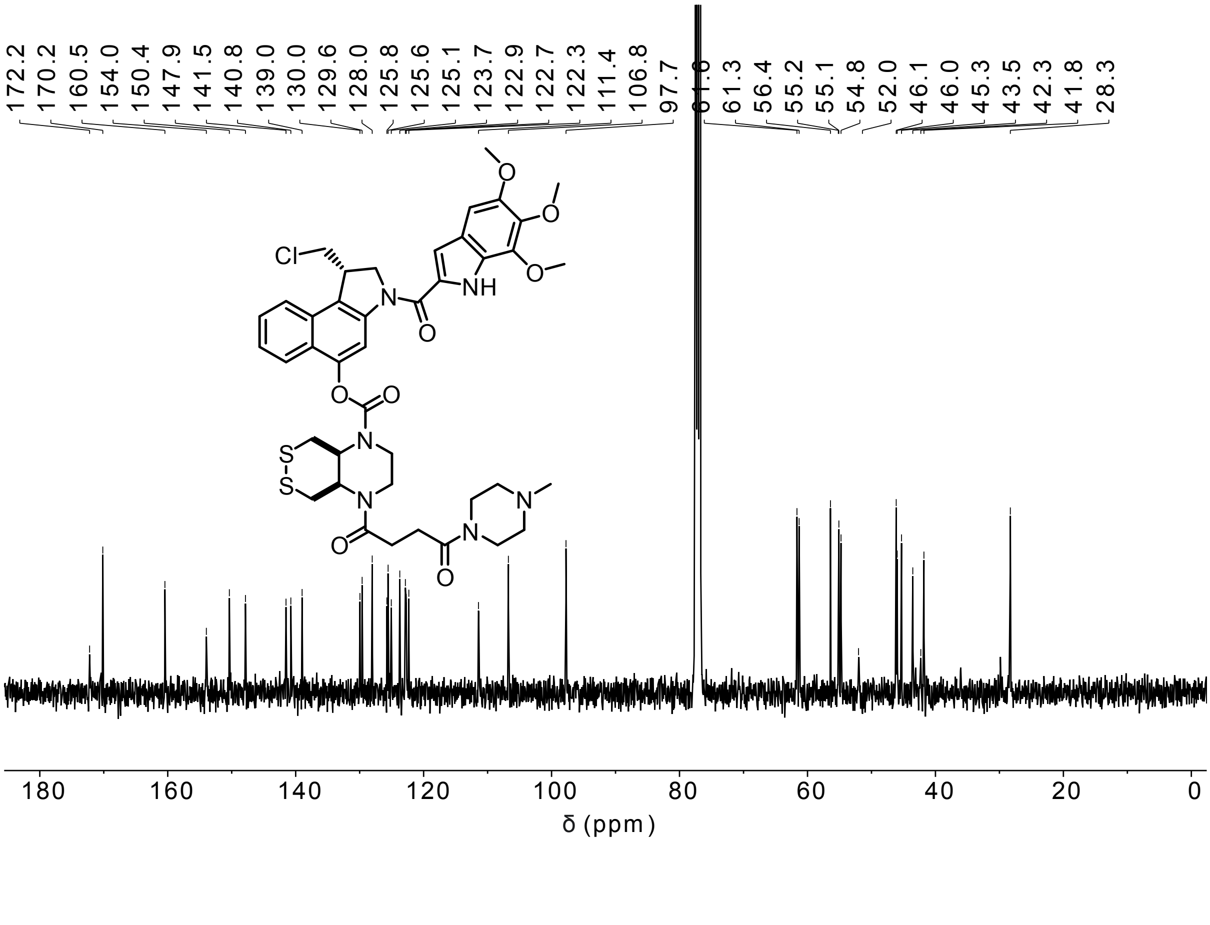

COSY

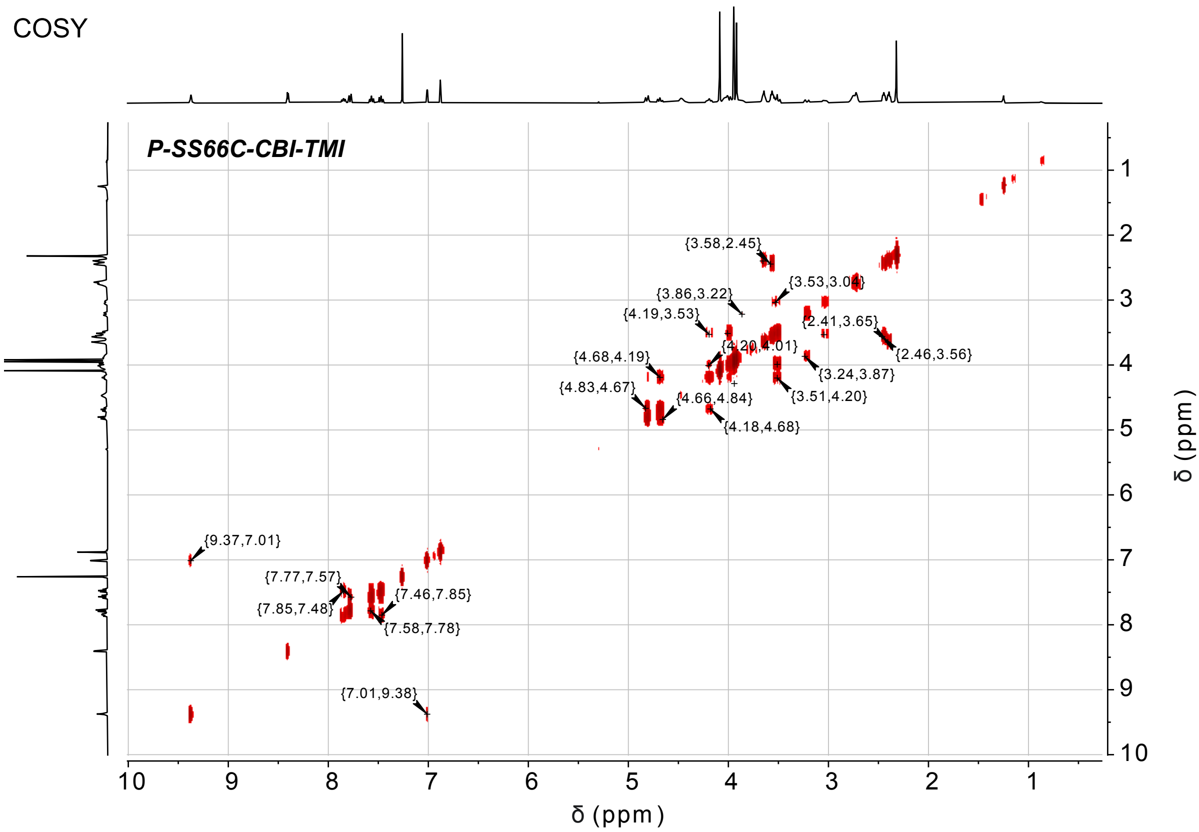
P-SS66C-CBI-TMI
{3.58,2.45}
{3.53,3.04}
{3.86,3.22}
{4.19,3.53}
{2.41,3.65}
{4.20,4.01}
{2.46,3.56}
{4.68,4.19}
{3.24,3.87}
{4.83,4.67}
{4.66,4.84}
{3.51,4.20}
{4.18,4.68}
{9.37,7.01}
{7.77,7.57}
{7.46,7.85}
{7.85,7.48}
{7.58,7.78}
{7.01,9.38}
δ (ppm)
δ (ppm)

HMQC

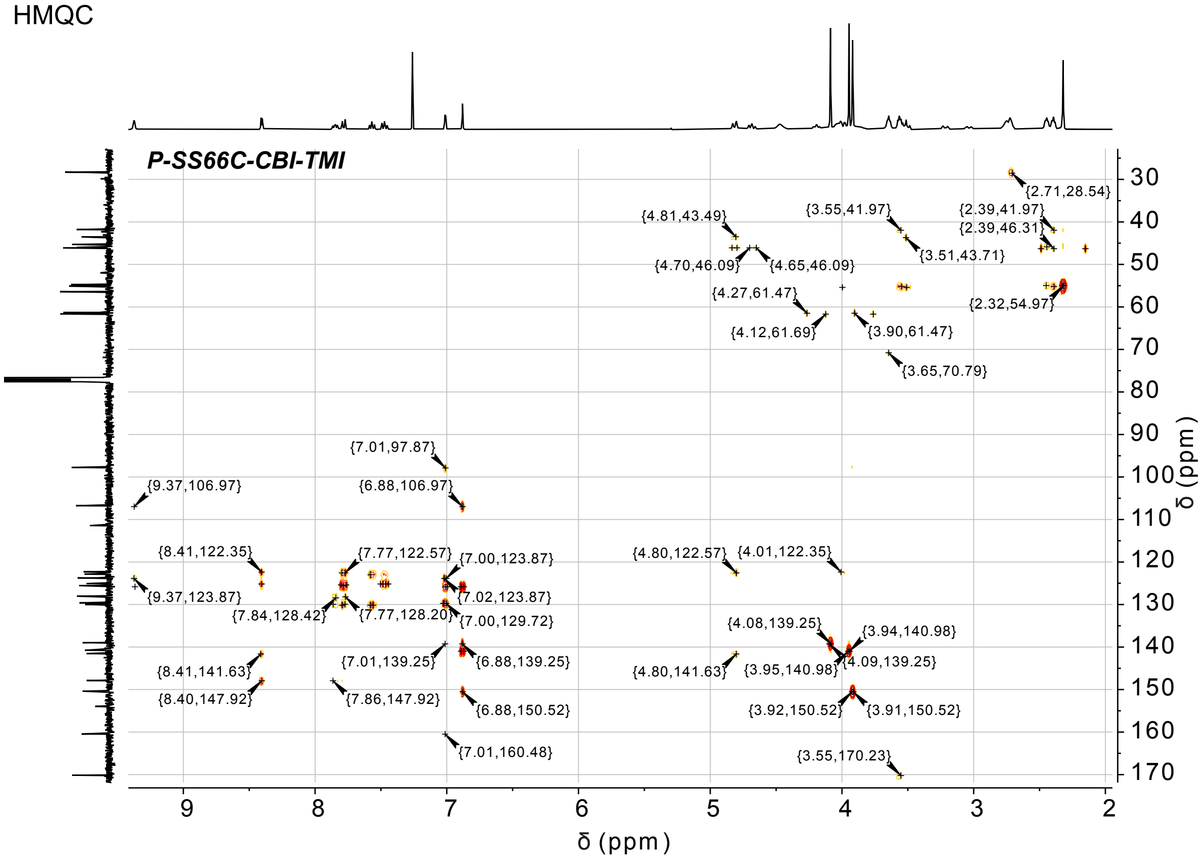
P-SS66C-CBI-TMI
{2.71,28.54}
{4.81,43.49}
{3.55,41.97}
{2.39,41.97}
{2.39,46.31}
{3.51,43.71}
{4.70,46.09}
{4.65,46.09}
{4.27,61.47}
{2.32,54.97}
{4.12,61.69}
{3.90,61.47}
{3.65,70.79}
{7.01,97.87}
{9.37,106.97}
{6.88,106.97}
{8.41,122.35}
{7.77,122.57}
{7.00,123.87}
{4.80,122.57}
{4.01,122.35}
{9.37,123.87}
{7.02,123.87}
{7.84,128.42}
{7.77,128.20}
{7.00,129.72}
{4.08,139.25}
{3.94,140.98}
{7.01,139.25}
{6.88,139.25}
{8.41,141.63}
{4.80,141.63}
{3.95,140.98}
{4.09,139.25}
{8.40,147.92}
{7.86,147.92}
{6.88,150.52}
{3.92,150.52}
{3.91,150.52}
{7.01,160.48}
{3.55,170.23}
δ (ppm)
δ (ppm)

HSQC - aromatic signals

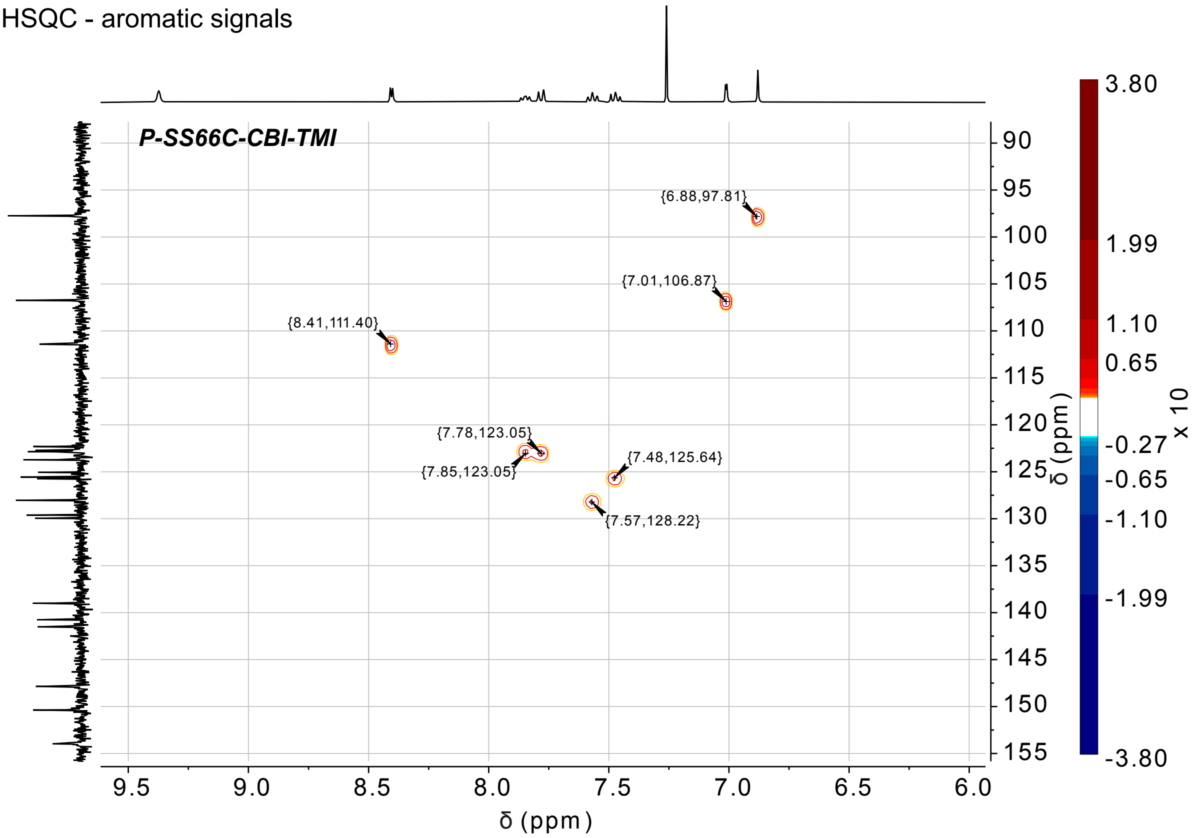


HSQC - aliphatic signals

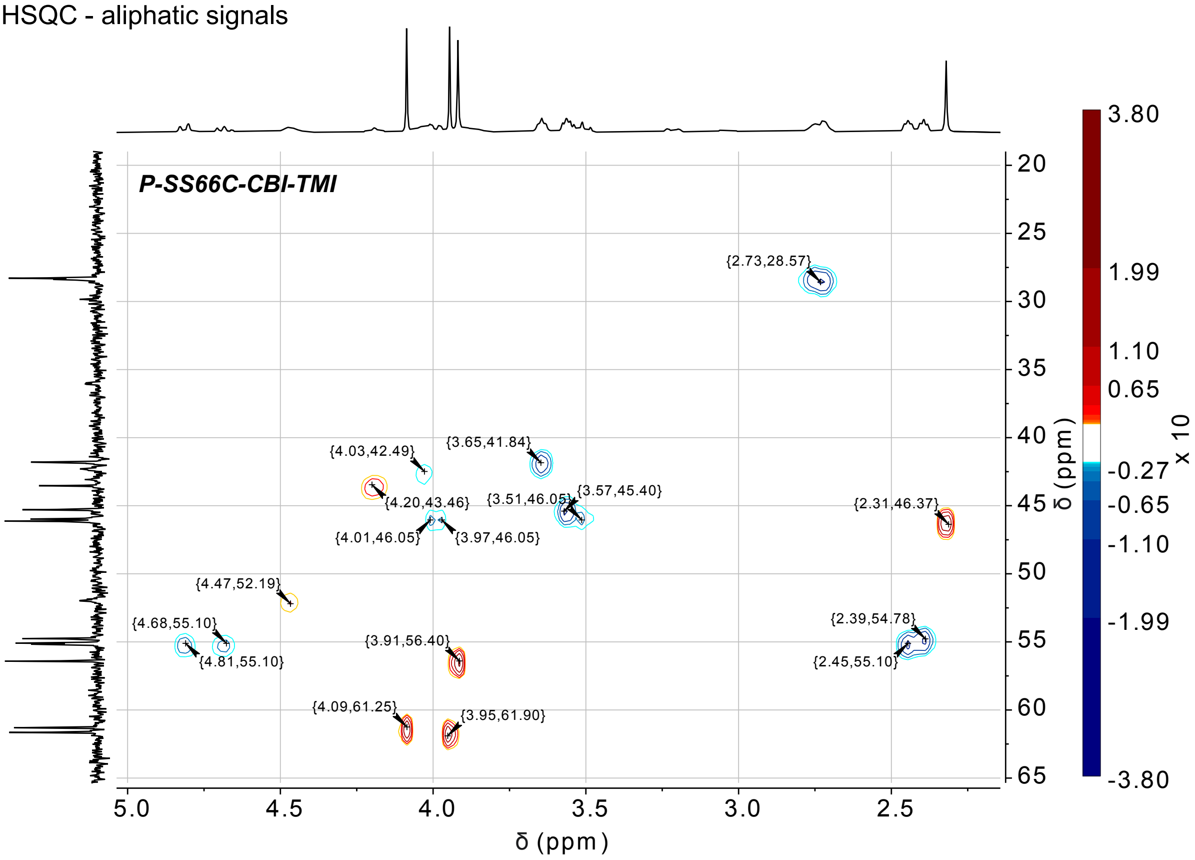

**SS00$_M$-CBI-TMI**

[1]H-NMR

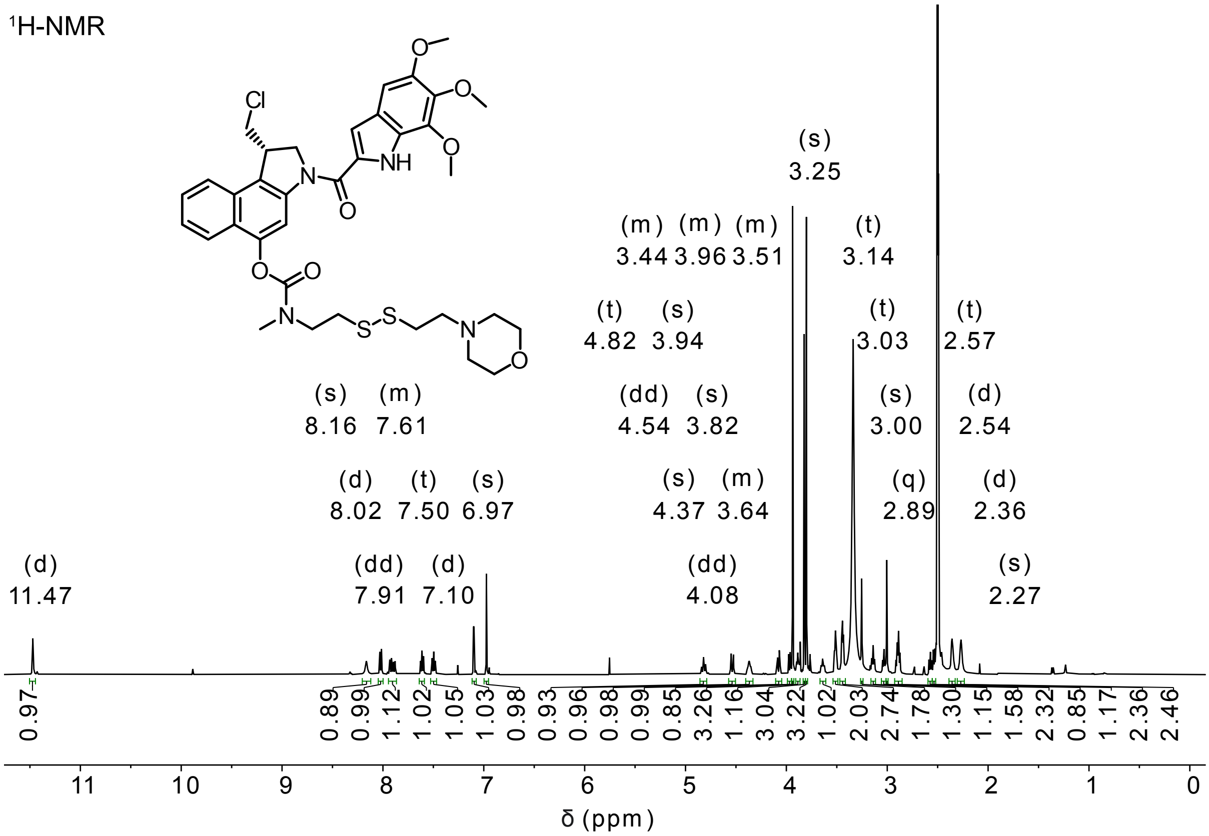


[13]C-NMR

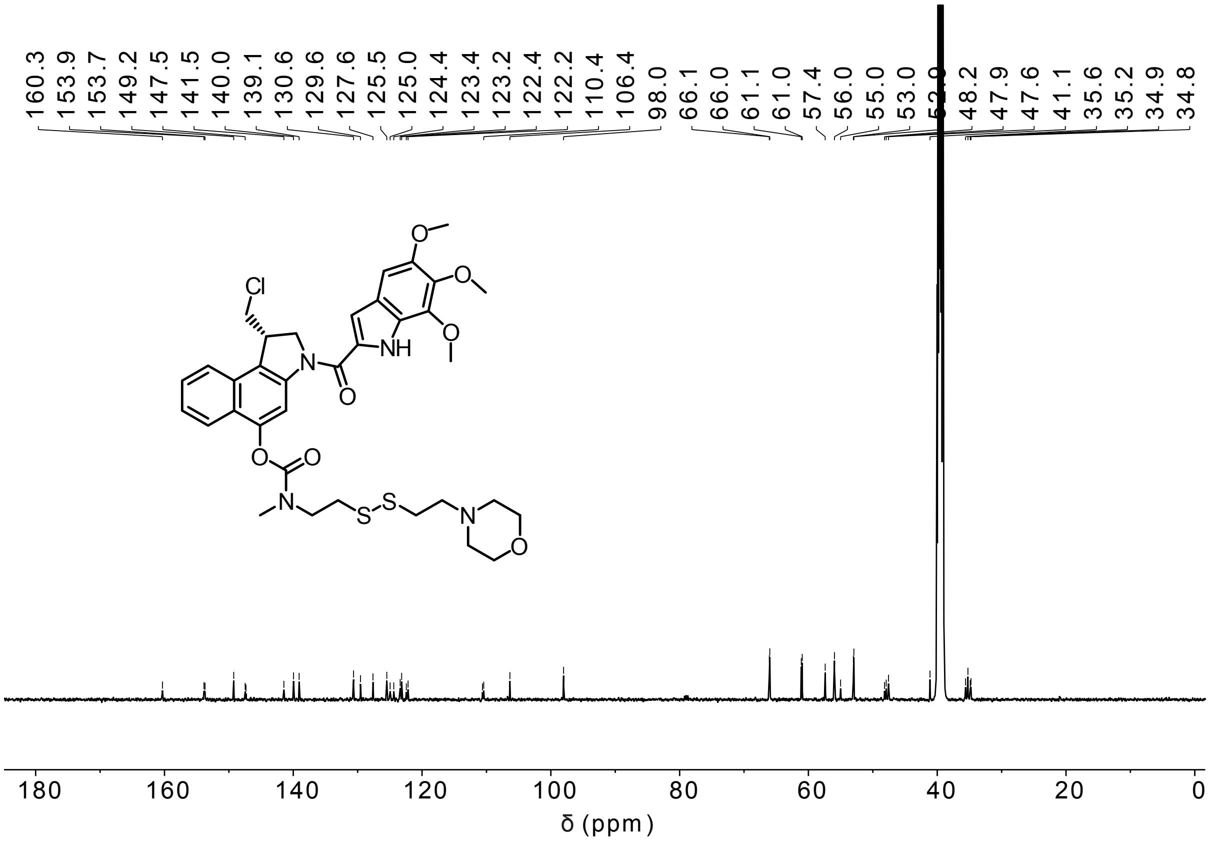

**MC-CBI-TMI**

[1]H-NMR

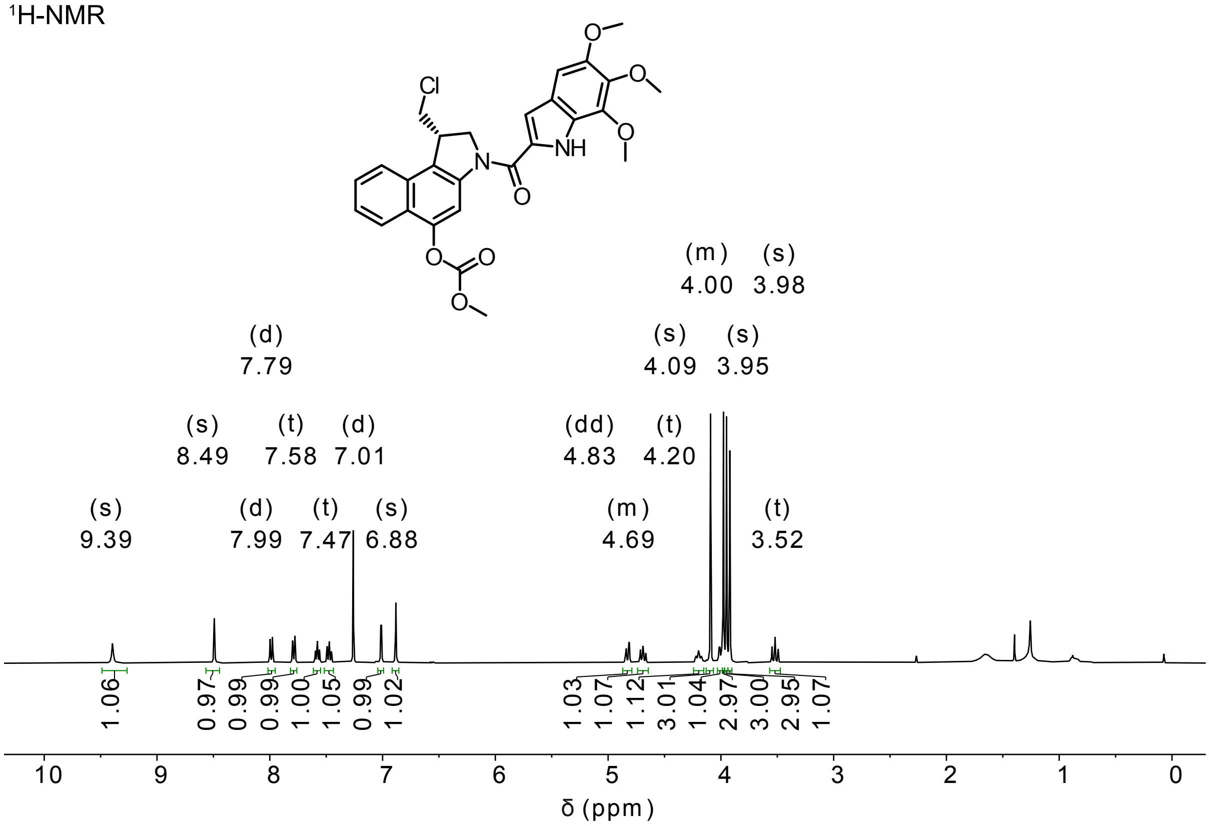


[13]C-NMR

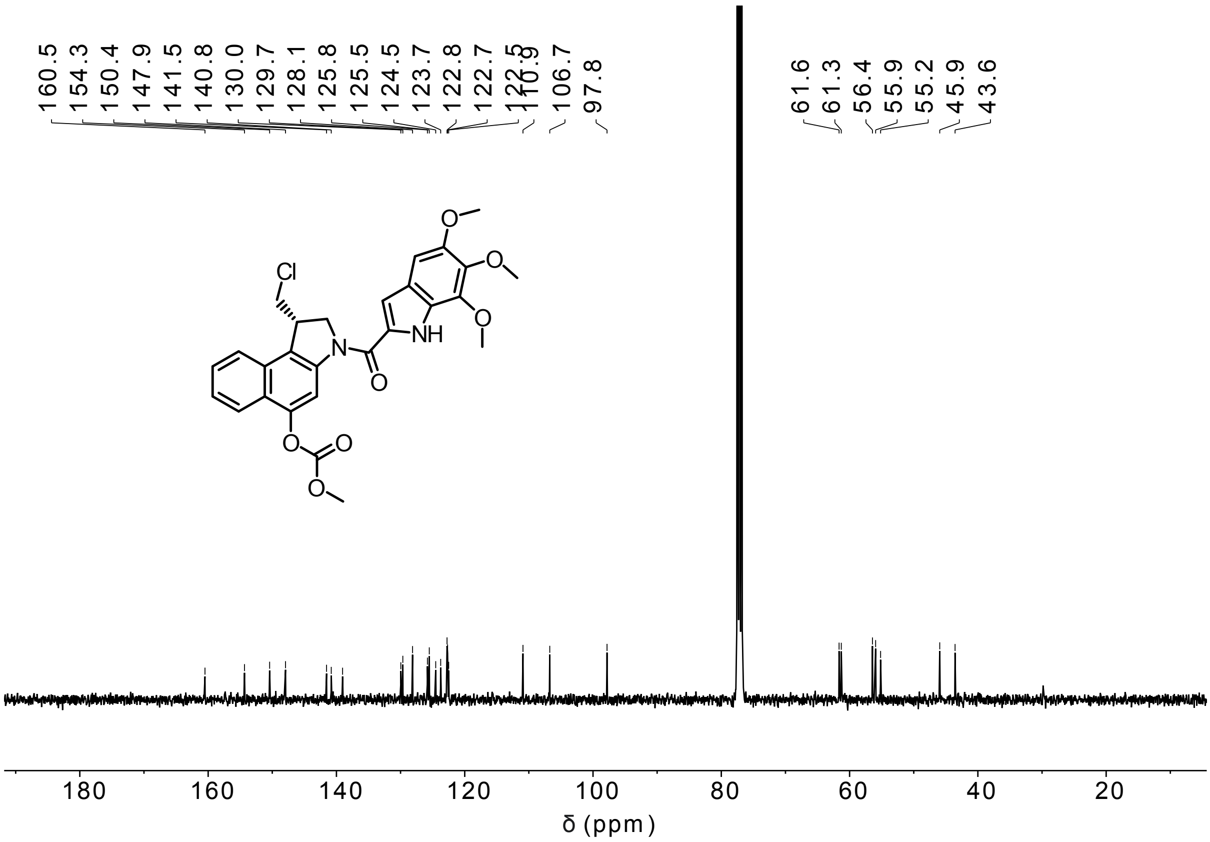

**P-CC60-CBI-TMI**

[1]H-NMR

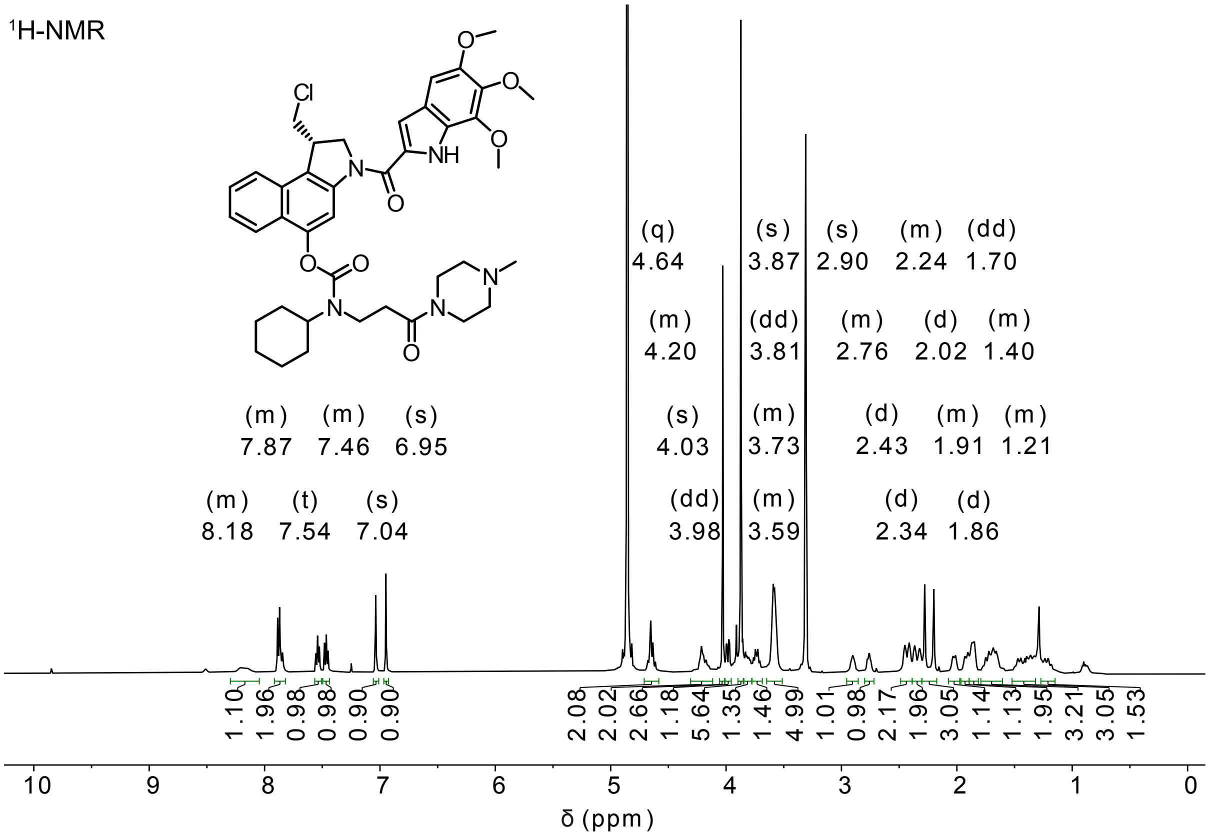


[13]C-NMR

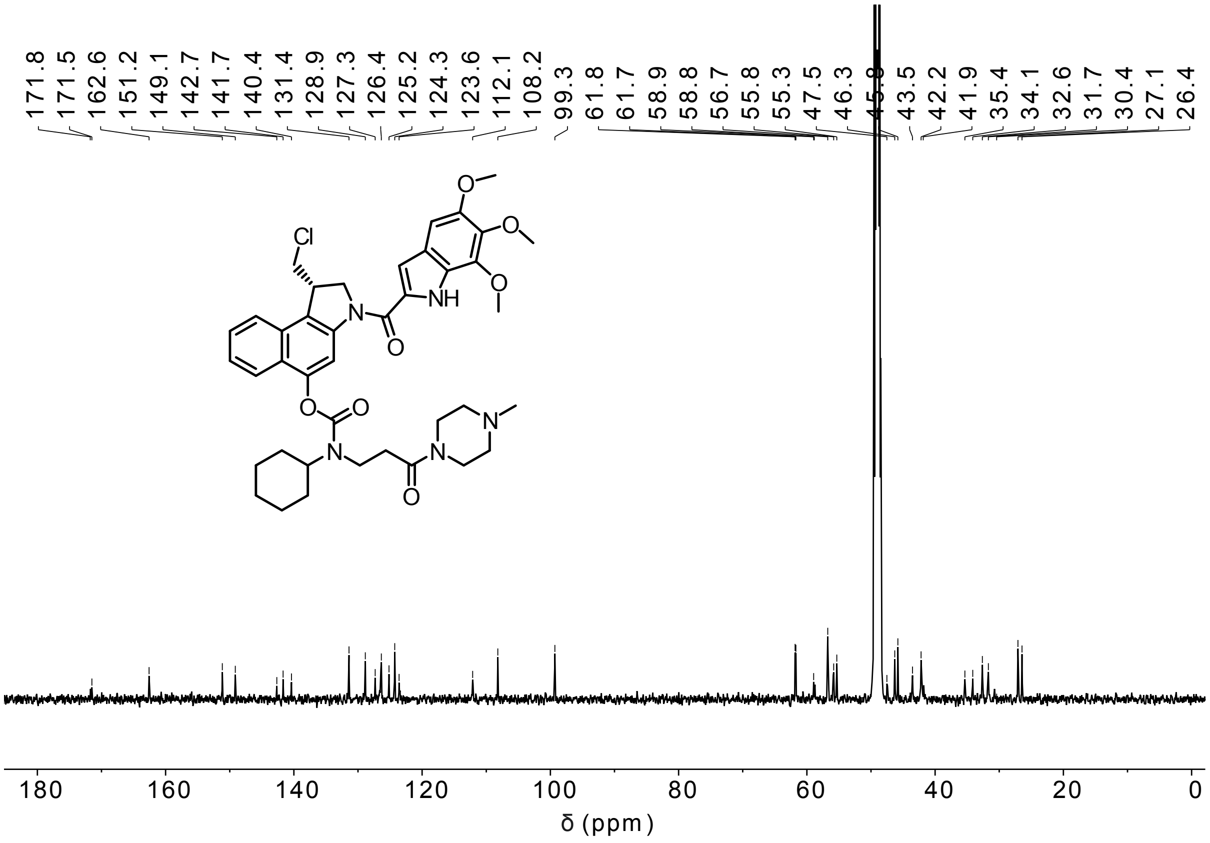

**Table S5.** Crystallographic data. **BnO-CBI-Boc**

| | 1 |
|---|---|
| net formula | $C_{25}H_{26}ClNO_3$ |
| $M_r$/g mol$^{-1}$ | 423.92 |
| crystal size/mm | 0.080 × 0.050 × 0.030 |
| $T$/K | 102.(2) |
| radiation | MoKα |
| diffractometer | 'Bruker D8 Venture TXS' |
| crystal system | orthorhombic |
| space group | 'P 21 21 21' |
| $a$/Å | 7.8910(4) |
| $b$/Å | 12.8528(6) |
| $c$/Å | 21.6817(11) |
| α/° | 90 |
| β/° | 90 |
| γ/° | 90 |
| $V$/Å$^3$ | 2198.99(19) |
| Z | 4 |
| calc. density/g cm$^{-3}$ | 1.280 |
| μ/mm$^{-1}$ | 0.200 |
| absorption correction | Multi-Scan |
| transmission factor range | 0.97–0.99 |
| refls. measured | 39132 |
| $R_{int}$ | 0.0394 |
| mean σ($I$)/$I$ | 0.0261 |
| θ range | 3.029–28.275 |
| observed refls. | 5214 |
| $x$, $y$ (weighting scheme) | 0.0401, 0.5584 |
| hydrogen refinement | constr |
| Flack parameter | 0.012(15) |
| refls in refinement | 5446 |
| parameters | 274 |
| restraints | 0 |
| $R(F_{obs})$ | 0.0324 |
| $R_w(F^2)$ | 0.0838 |
| S | 1.094 |
| shift/error$_{max}$ | 0.001 |
| max electron density/e Å$^{-3}$ | 0.248 |
| min electron density/e Å$^{-3}$ | −0.199 |

Anomalous dispersion effects (Flack test) applied in order to determine the correct structure.
